# Environment and taxonomy shape the genomic signature of prokaryotic extremophiles

**DOI:** 10.1101/2023.05.24.542097

**Authors:** Pablo Millán Arias, Joseph Butler, Gurjit S. Randhawa, Maximillian P. M. Soltysiak, Kathleen A. Hill, Lila Kari

**Author notes:** These first authors contributed computer science, respectively biology, aspects of this work (Pablo Millán Arias, Joseph Butler).

## Abstract

This study provides comprehensive quantitative evidence suggesting that adaptations to extreme temperatures and pH imprint a discernible environmental component in the genomic signature of microbial extremophiles. Both supervised and unsupervised machine learning algorithms were used to analyze genomic signatures, each computed as the *k*-mer frequency vector of a 500 kbp DNA fragment arbitrarily selected to represent a genome. Computational experiments classified/clustered genomic signatures extracted from a curated dataset of ∼700 extremophile (temperature, pH) bacteria and archaea genomes, at multiple scales of analysis, 1 ≤*k* ≤6. The supervised learning resulted in high accuracies for taxonomic classifications at 2 ≤*k* ≤6, and medium to medium-high accuracies for environment category classifications of the same datasets at 3 ≤*k* ≤6. For *k* = 3, our findings were largely consistent with amino acid compositional biases and codon usage patterns in coding regions, previously attributed to extreme environment adaptations. The unsupervised learning of unlabelled sequences identified several exemplars of hyperthermophilic organisms with large similarities in their genomic signatures, in spite of belonging to different domains in the Tree of Life.

## Introduction

Life exists in the most unusual and extreme environments on our planet. Biodiversity exists in environments such as volcanoes, deep-sea trenches, and polar regions, that are characterized by extreme physical conditions (temperature, radiation, pressure, salinity, pH, chemical, etc.), and pose unique challenges to life^1^. Organisms that are able to survive, and sometimes thrive, in extreme conditions are known as *extremophiles*, characterized by their peculiar phenotypic adaptations and yet underexplored in their genome composition. The study of the genome organization and diversity of extremophiles is of particular interest, as it can provide insights into the mechanisms of adaptation and the evolution of biodiversity in extreme environmental conditions^2–4^. In particular, the study of microbial extremophiles has yielded important research reagents and gained popularity more recently due to their potential applications in biorefineries^5^, as sources of industrially-relevant biocatalysts^6^, and due to explorations of microbial dormancy and survivability in outer space^7, 8^. This study uses several machine learning algorithms and an alignment-free methodology to uncover evidence suggesting that microbial adaptations to extreme temperatures and pH conditions imprint a discernible environmental component in their genomic signatures.

As it has been previously observed, extremophiles have developed a wide range of structural, biochemical and metabolic strategies that support cell viability in high-stress environments, and there is evidence that the adaptive mechanisms arising with convergent evolution of extremophilic taxa can be observed at the proteomic and genomic levels^9^. At the proteomic level, diverse organisms living in certain extreme environments have a strong amino acid compositional bias, attributed in part to convergent proteomic adaptations^3, 10, 11^. At the genomic level, codon usage patterns in the genomes of different extremophilic organisms are linked to the physiochemical characteristics imposed by selective pressures experienced in their respective environments^2, 12, 13^. In addition to the localized influences of the selection pressures upon the proteome, open reading frame sequence composition is also influenced by nucleic acid level adaptations associated with structural functions^14^. In particular, high Guanine+Cytosine (G+C) content was observed to be correlated with DNA stability in thermophiles of diverse taxa^15^ and is the major factor influencing tRNA stability in hyperthermophiles^16^, while the fraction of Adenine+Guanine (A+G) content in coding DNA was observed to be correlated with the optimal growth temperature (OGT) in thermophiles^17–19^. These observations all suggest the hypothesis of a correlation between adaptation to extreme environments and specific genome composition patterns.

One way to approach the concept of genome composition is to study “genomic signatures”^20^, a general term used for a variety of quantitative measures, pervasive along a genome, that can be used to discriminate between genomes of different species^21^. In the last two decades, numerous studies have confirmed the effectiveness of alignment-free methods that use genomic signatures for the purpose of genome analysis^22^, comparison^23^, and sensitive taxonomic classification^24–26^, even without supervision^27^. These findings confirmed the existence of a strong phylogenetic signal that is present in (genome-wide, pervasive) genomic signatures, and offered a different perspective complementing alignment-based taxonomic comparisons and distinctions. In particular, genomic signatures based on *k*-mer (subwords of length *k*) frequency profiles have been widely used to classify organisms at different taxonomic levels, from Kingdom to species subtypes^24–26, 28–31^.

These findings suggested that differences and similarities in genomic signatures can be attributed to phylogeny. However, the possibility exists that there are other contributors to the differentiating power of genomic signatures^32^. Of special interest are the genomes of extremophile microbes of diverse taxa, which present a unique case marked by convergent adaptations to extreme physical environments. As shared phenotypic adaptations can also impact genome sequence composition (i.e., codon usage, nucleotide biases), genomic signature analyses may identify instances of convergent evolution. Prior work on thermophiles has also suggested some genomic composition biases, such as dinucleotide^20^ or tetranucleotide^33^ frequencies, that are pervasive across the genome. In addition, other research suggests that amino acid compositional biases and codon usage patterns in coding regions may contribute to the existence of a detectable pervasive genomic signature, strong enough to differentiate between taxonomically related organisms that live at opposite environmental extremes^10, 19^.

This paper provides comprehensive quantitative evidence suggesting that adaptation to extreme temperatures or pH introduces a discernible environmental component in the genomic signature of microbial extremophiles. Herein, a genomic signature is defined as the *k*-mer frequency vector of a 500 kbp DNA fragment, arbitrarily selected to represent a genome, where *k* is a fixed positive integer 1 ≤*k* ≤6. This hypothesis was tested using both supervised and unsupervised machine learning algorithms, on a prokaryote dataset comprising 693 high-quality genomes from bacterial or archaeal organisms adapted to extreme temperature conditions, or extreme pH conditions. Supervised machine learning has proved effective in using genomic signatures for taxonomic classification, on data that was problematic for traditional alignment-based algorithms due to its sparseness, complexity, and high dimensionality^25, 26, 29^. Thus, the first approach was to use supervised machine learning methods to learn the taxonomic and environmental components of genomic signatures. To this end, several supervised learning algorithms were trained on genomic signatures labelled with either taxonomic labels or with environment category labels. Each classifier was then used to obtain a taxonomic classification (if it was trained using taxonomic labels), or an environment category classification (if it was trained using environment category labels). The classification accuracies obtained were high for the taxonomic classification, and medium to medium-high for the environment category classification, suggesting the presence of an environmental component in the genomic signature in addition to its taxonomic component. For further insight, interpretability tools of supervised learning were used to determine the features of the genomic signatures (specific *k*-mers) that were most relevant to the environment category classification, and our findings were compared with existing literature on codon usage and amino acid compositional bias in extremophiles.

The presence of an environmental component in the genomic signature was independently confirmed by unsupervised clustering analysis of data, whereby the first step was to assess several unsupervised learning algorithms for their ability to learn the taxonomic structure of unlabelled data. The most performant clustering algorithms were then used to identify several candidate organisms, with similar genomic signatures in spite of large taxonomic differences. Of these, additional stringent tests based on supervised learning classifications, in challenging scenarios, identified exemplars of hyperthemophile bacteria and archaea whose genomic signatures were grouped together under all classification and clustering scenarios, by all machine learning algorithms used.

The main contributions of this paper are:

- An extensive supervised machine learning analysis of a dataset, augmented with literature references and annotations, of ∼700 high-quality microbial extremophile genomes (temperature and pH), at various scales. The results suggest the presence of an environmental component in the genomic signature of microbial extremophiles (temperature, pH) for values 3 ≤*k* ≤6, in addition to a strong taxonomic component for values 2 ≤*k* ≤6. Subsets of 3-mers that contribute to this enviromental component are also identified, together with an assessment of the relative importance of their contribution.
- An unsupervised clustering-based analysis of the aforementioned dataset, providing independent support of the hypothesis of the presence of an environmental component in the genomic signature of these extremophiles. This analysis also identified a hyperthemophile bacterium, *Thermocrinis ruber*, and three hyperthermophile archaea, *Pyrococcus furiosus, Thermococcus litoralis*, and *Pyrococcus chitonophagus*, as exemplars of genomic signatures grouped together by all machine learning algorithms used, in spite of their vast taxonomic differences.

Overall, the results of machine learning analyses, corroborated in the exemplar cases by observations of shared characteristics of the isolating environments, suggest the existence of an *environmental component* that co-exists with a strong *taxonomic component* in the genomic signatures of organisms living in extreme temperatures or extreme pH conditions. To the best of our knowledge, this study is the most comprehensive examination to date of the genomic signature of prokaryotic extremophiles, at various scales, of a substantial, well-curated, dataset of extremophile genomes.

## Materials and Methods

### Datasets

The data in this study were collected through a systematic literature search focused on identifying extremophilic microbes adapted to environments of extreme temperature and extreme pH. The search was conducted on the PubMed Database (accessed September 2022) and Google Scholar (accessed September 2022) for primary research articles and reviews, and identified 768 microbial species or strains for which extremophilic characteristics were recorded. Subsequently, these species/strains were identified in the Genome Taxonomy Database (GTDB; release R207 April 8, 2022, Accessed February 2023), the gold-standard database for taxonomy^34^, and only GTDB species representative genomes with reported completeness of over 95%, and contamination of under 5% were selected. Species/strains were mapped to their identified extremophilic characteristic(s), along with genome assembly numbers provided by GTDB for each given organism. The extremophilic characteristic(s) was validated for each organism by searching PubMed with the given strain/species name, and identifying a primary article/review and/or reliable BacDive database (accessed February 2023) entry to confirm the accuracy of the characteristic(s). Entries lacking consistent observations related to the growth characteristics of the respective microbe were removed from the dataset.

For this study, we used the following definitions, based on the Optimal Growth Temperature (OGT), respectively Optimal Growth pH (OGpH): Psychrophile (OGT of < 20ºC)^32^, mesophile (OGT of 20-45ºC)^32^, thermophile (OGT of 45-80ºC)^32^, and hyperthermophile (OGT of > 80ºC)^32^, acidophile (OgpH < pH 5)^32^ and alkaliphile (OGpH > pH 9)^32^. The dataset was then curated for 154 descriptors, so as to be in accordance with the temperature and pH intervals used in the above definitions. Fourteen entries could not be validated and were discarded from the dataset.

This selection process resulted in 693 annotated high-quality extremophile microbial genome assemblies. These high-quality assemblies were then used to form two datasets according to two extremophilic characteristic(s), as follows. The first dataset, called the Temperature Dataset, is composed of 148 psychrophile genomes (8 archaeal, 140 bacterial), 190 mesophile genomes (84 archaeal, 106 bacterial), 183 thermophile genomes (67 archaeal, 116 bacteria), and 77 hyperthermophile genomes (70 archaeal, 7 bacterial) for a total of 598 organism genomes (229 archaeal, 369 bacterial) (Table 1). The second dataset, called the pH Dataset, is composed of 100 acidophile genomes (39 archaeal, 61 bacterial) and 86 alkaliphile genomes (30 archaeal, 56 bacterial), for a total of 186 organisms (69 archaeal, 117 bacterial) (Table 2). Note that 91 organisms were identified to belong to both the Temperature Dataset and the pH Dataset. The datasets are described in Supplementary Table S1, with assembly metadata provided in Supplementary Table S2. The proportions of both the Temperature Dataset and the pH Dataset are described in terms of genus, organized by domain and by environment category, and are described in greater detail in Supplementary Data S1. As well, phylogenetic trees organized by domain (bacteria, archaea) and environment category (temperature, pH) are accessible in Supplementary Data S2.

**Table 1.** Composition of the Temperature Dataset: 598 DNA fragments from microbial genomes/species (369 DNA fragments from bacterial genomes, and 229 DNA fragments from archaeal genomes).

| Domain | Temperature Category | # Phyla | # Classes | # Orders | # Families | # Genera | # Species |
| --- | --- | --- | --- | --- | --- | --- | --- |
| Archaea | Psychrophiles | 2 | 4 | 4 | 5 | 7 | 8 |
|  | Mesophiles | 4 | 6 | 7 | 20 | 45 | 84 |
|  | Thermophiles | 6 | 11 | 14 | 21 | 41 | 67 |
|  | Hyperthermophiles | 5 | 6 | 8 | 15 | 31 | 70 |
| Bacteria | Psychrophiles | 4 | 4 | 6 | 13 | 19 | 140 |
|  | Mesophiles | 3 | 3 | 6 | 10 | 14 | 106 |
|  | Thermophiles | 15 | 19 | 24 | 27 | 47 | 116 |
|  | Hyperthermophiles | 5 | 5 | 5 | 5 | 5 | 7 |

**Table 2.** Composition of the pH Dataset: 186 DNA fragments from microbial genomes/species (117 DNA fragments from bacterial genomes, and 69 DNA fragments from archaeal genomes).

| Domain | pH Category | # Phyla | # Classes | # Orders | # Families | # Genera | # Species |
| --- | --- | --- | --- | --- | --- | --- | --- |
| Archaea | Acidophiles | 4 | 5 | 7 | 11 | 24 | 39 |
|  | Alkaliphiles | 2 | 5 | 5 | 9 | 18 | 30 |
| Bacteria | Acidophiles | 10 | 12 | 13 | 13 | 32 | 61 |
|  | Alkaliphiles | 12 | 14 | 25 | 30 | 36 | 56 |

The selection of a genomic fragment *s* to represent the genome of an organism is a process that has to consider several factors, including fragment length, taxonomic level, and computational complexity of the algorithms used. For methods that rely on *k*-mer frequency for sequence classification, some studies^35, 36^ suggest the relation *k* = *log*_4_(|*s*|), where |*s*| is the minimum length of sequence *s* that is necessary, in theory, to obtain statistical significance. However, in practice, longer sequences are needed. For example, another study^27^ used sequence length of 500 kbp in conjunction with *k* = 6 to cluster bacterial sequences at the family level, even though in theory a length of 4,096 bp would have sufficed for this value of *k*.

In this study, each genome (assembly) was represented by a single, arbitrarily selected, 500 kbp DNA fragment. The values for *k* used in this study, namely 1 ≤*k* ≤6, were empirically selected so as to balance the trade-off between classification accuracy and computational complexity, and to explore multiple scales of the *k*-mer analysis.

More precisely, a DNA fragment was arbitrarily selected to represent each DNA genome/assembly, as follows. First, the contigs of the assembly were sorted by length, from the longest to the shortest. Then, if the longest contig was longer than 500 kbp, then a 500 kbp fragment was randomly selected to be the *representative DNA sequence* for that genome. Otherwise, the sorted contigs were concatenated one by one, until the desired length of 500 kbp was reached, and this became the DNA representative sequence for that genome. The *k*-mers were counted starting from the beginning of the representative DNA sequence, by using a sliding window with step size 1. To avoid spurious *k*-mers that could arise from the concatenation of contigs, the *N* character was added as a separator between contigs or contig fragments, but no *k*-mers that contain *N* were considered when calculating *k*-mer counts. Also note that the inserted letters *N* were not counted towards the length of the DNA sequence representing each genome/assembly. To eliminate the variable of the strand orientation of the uploaded DNA sequences, the final *k*-mer frequency vector of a sequence was computed as the sum between the vector of its *k*-mer counts and the corresponding vector of *k*-mer counts of its reverse complement.^37^ In the remainder of this paper, a *k*-mer and its reverse complement will be considered to be indistinguishable, and only the *canonical k*-mer of a pair (the first, in alphabetical order, of the two reverse complementary *k*-mers) will be listed.

### Sequence Classification Using Supervised Machine Learning

To test the hypothesis of the existence of an environmental component in the genomic signature of microbial extremophiles, the two previously described datasets (Temperature, and pH) were classified using supervised machine learning algorithms, and the average accuracy of each classification was computed. For each dataset, computational experiments were performed using six different classifiers, and different values of *k*, as detailed below. In addition, for each computational experiment, three different scenarios for labelling the training dataset were analyzed, as follows:

1. All DNA sequences used in training were labelled taxonomically, by their domain (bacteria or archaea),
2. All DNA sequences used in training were labelled by their environment category (psychrophile, mesophiles, acidophile, etc.),
3. All DNA sequences used in training were labelled with pseudo-labels sampled from a discrete uniform distribution. The discrete uniform distribution was *Unif*(0, 3) in the case of the Temperature Dataset (four possible labels), and respectively *Unif*(0, 1) in the case of the pH Dataset (two possible labels). This third scenario was introduced as a control, and it was expected to result in predictions of the correct pseudo-labels with probabilities equal to the sampling probability for each dataset. Note that an alternative sampling strategy would be to sample the pseudo-labels according to the distribution of the environment category labels in the dataset. The results associated with this alternative sampling strategy can be found in Supplementary Table S3.

The six different classifiers used for these classification tasks were selected as being representative algorithms of four main categories in the classification of DNA sequences. Support Vector Machines (SVM) were selected as a representative of *Kernel Methods*, with a radial basis function kernel^38^. Random Forest was selected as a representative of *Tree-Based Methods*, with the Gini index as the classification criteria^39^. The third algorithm was an *Artificial Neural Network (ANN)*, with a simple and versatile architecture consisting of two fully connected layers, Linear (512 neurons) and Linear (64 neurons), each one followed by a Rectified Linear Unit (ReLU) and a Dropout layer with a dropout rate of 0.5. Lastly, a *Digital Signal Processing* framework^26^ was considered, whereby pairwise distances between numerical representations of DNA sequences are computed and then used in conjunction with *Linear Discriminant* (MLDSP-1), with *Quadratic SVM* (MLDSP-2), or with *Subspace Discriminant* (MLDSP-3) machine learning algorithms.

Two different types of computational experiments were performed for each of the two datasets (Temperature and pH), supervised machine learning classifier (six classifiers), value of *k* (1 ≤*k* ≤6), and training data labelling (taxonomy, environment category, random).

In the first type of tests, called *restriction-free*, the predictive power of the algorithms was tested using standard stratified 10-fold cross-validation, as follows. The dataset was split into 10 distinct subsets, called *folds*, and a model was trained using 9 of the folds as training data; the resulting model was validated on the remaining part of the data (i.e., it was used as a test set to compute a performance measure such as accuracy). The performance measure reported by 10-fold cross-validation was calculated as the average of the classification accuracy for each of the 10 possible test sets.

The second type of tests, called *restricted*, or *non-overlapping genera*, was designed to address the possibility that a correct environment category label classification may be influenced by a contributing taxonomic component. For example, one goal was to ensure that a DNA sequence was not classified as a hyperthermophile simply due to its similarity to DNA sequences of the same genus, that happened to belong to the same hyperthermophile category. To this end, we adopted a modified 10-fold cross-validation approach, whereby all sequences of the same genus appeared in exactly one fold. At the same time, to align with the principles of stratified cross-validation, the distribution of the labels in each fold was kept the same as the distribution of the corresponding labels in the entire dataset. In this *restricted* (*non-overlapping genera*) scenario, if a DNA sequence is in the test set, then no other sequence of the same genus is present in the training set. This approach attempts to disentangle, at the genus level, the taxonomic component from the environmental component of the genomic signature.

As an independent method for assessing the environmental component of the genomic signature, we employed interpretability tools for machine learning methods. Global interpretability tools were preferred as they are useful in understanding the general mechanisms in the data through a global importance measure. Given the high correlation between the *k*-mers, the mean decrease in impurity (MDI) for Random Forest was selected as a *k*-mer global importance measure, and then used to learn the actual *k*-mers that were relevant to the environment category classification. (This measure was preferred over the widely adopted global-agnostic Permutation Feature Importance method, as that method is not suitable for handling highly correlated features^40^.) The methodology used to determine the relevant *k*-mers is as follows. First, a one-vs-all classifier was trained for each environment category present in the dataset, using stratified 10-fold cross-validation. Second, the MDI algorithm was used to compute the global importance of each *k*-mer in each fold, and the average taken over all folds was used to create a ranked list of *k*-mers, in decreasing order of their contribution. Finally, for each environment category, the “most relevant subset of *k*-mers” was computed, defined as the subset of the ranked *k*-mer list that was sufficient to classify the dataset with the same classification accuracy as when *all k*-mers were used in that classification.

### Unsupervised Learning for Sequence Clustering

In unsupervised learning, no labels are provided for the DNA sequences in the dataset, and various algorithms are used to cluster similar genomic signatures, and to explore the structure of the space of the genomic signatures in the dataset.

Two groups of tests with unsupervised learning algorithms were performed in this study: parametric clustering algorithms (that take the number of expected clusters as an input parameter), and non-parametric clustering algorithms (that determine automatically the number of clusters). In the first group, four parametric clustering algorithms were used: K-means, Gaussian Mixture Model, K-medoids, and DeLUCS^27^. The computation of the cluster label assignments for each sequence in the Temperature and the pH Datasets was performed with various values of the parameter `n_clusters` (the expected number of clusters) in each algorithm, `n_clusters` ∈{2, 4, 8} for the Temperature Dataset, and respectively `n_clusters` ∈{2, 4} for the pH Dataset, based on the number of potential true clusters in each dataset.

For each dataset, the strength of each of the two components of the signature (taxonomic, environmental) was assessed by comparing the clustering accuracies in two scenarios, the first where the clustering was assessed against the true taxonomic groups, and the second when the clustering was assessed against the true environment category groups. In each case, the performance was evaluated using the unsupervised clustering accuracy metric^41^, defined as:

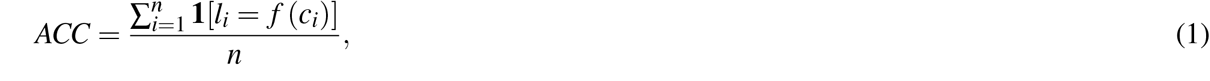

where *n* is the total number of sequences and, for each 1 ≤*i* ≤*n* and DNA sequence *x*_*i*_ we have that: The true taxonomic label of *x*_*i*_ is denoted by *l*_*i*_; the numerical cluster label that the algorithm assigns to *x*_*i*_ is denoted by *c*_*i*_; an optimal mapping *f* is calculated by the Hungarian algorithm^42^, that maps numerical cluster labels to true taxonomic labels (specifically, *f* (*c*_*i*_) denotes the true taxonomic label assigned by *f* to the cluster labelled *c*_*i*_); and **1**[*k* = *i*] ∈{0, 1} is an indicator function, equal to 1 if and only if *k* = *i*.

In the second type of test, we assessed whether the clusters of each dataset at the lowest possible taxonomic level (genus) can be discovered by non-parametric clustering algorithms. For this purpose, we used two non-parametric clustering algorithms, HDBSCAN^43^ and iterative medoids^28^, combined with three different dimensionality reduction techniques: Variational autoencoders (VAE), Deep Contrastive Learning (CL) and Uniform Map Approximation (UMAP). We also used *i*DeLUCS, which is semi-parametric, in the sense that its parameter `n_clusters` (herein = 300) represents an upper limit of the number of clusters found by the algorithm. These seven clustering algorithms were used to recover the lowest taxonomic groups. The following metrics were defined to assess the quality of the found clusters: the *completeness* of each cluster (defined as the number of occurrences of the most common genus present in the cluster, divided by the total number of sequences of that genus in the dataset), and the *contamination* of each cluster (defined as the number of sequences that belong to the most common genus in the cluster, divided by the cluster size). The overall quality of each clustering algorithm was then calculated as the total number of clusters that are at least 50% complete, and at most 50% contaminated.

## Results

### Supervised Machine Learning Analysis of the Temperature Dataset and the pH Dataset *Supervised classification by taxonomy, environment category, and random label assignment*

Several supervised machine learning computational tests were performed to classify the Temperature Dataset and the pH Dataset, respectively, using *(1)* taxonomy labels (domain), *(2)* environment category labels, and *(3)* randomly assigned environment category labels (four for the Temperature Dataset, respectively two for the pH Dataset). More specifically, six supervised machine learning algorithms were used to classify the two datasets, for several *k*-mer lengths, 1 ≤*k* ≤6. The classification tests were performed under two scenarios, *(a) restriction-free*, using stratified 10-fold cross-validation, and *(b) restricted*, using stratified 10-fold cross-validation with non-overlapping genera.

The classification accuracies for the *restriction-free* case are summarized in Table 3. For *k* = 6, classifications using taxonomy labels for training resulted in high classification accuracies of over 97.49% for the Temperature Dataset and over 94.18% for the pH Dataset, across all six classification models. Classifications using environment category labels resulted in medium-high classification accuracies of over 77.59% for the Temperature Dataset and over 84.95% for the pH Dataset, across all six classification models. Classifications using randomly assigned labels resulted in the expected low accuracies of at most 28.09% for the Temperature Dataset and at most 50.06% for the pH Dataset, across all six classification models.

**Table 3.** Classification accuracies of six supervised learning classifiers trained on the Temperature Dataset and pH Dataset, in the *restriction-free* scenario, for three different label assignments (taxonomy, environment category, and random label assignment), and values of 1 ≤*k* ≤6. The classification accuracy in each cell is calculated using standard stratified 10-fold cross-validation.

| Dataset | $k$-value | Class Labelling Type | Classification Model Accuracy (%) | | | | | |
| --- | --- | --- | --- | --- | --- | --- | --- | --- |
|  |  |  | RBF SVM | Random Forest | ANN | MLDSP-1 | MLDSP-2 | MLDSP-3 |
| Temperature | $k = 1$ | Taxonomy | 62.88 | 53.87 | 62.21 | 47.99 | 54.85 | 59.03 |
|  |  | by Environment | 39.97 | 35.29 | 38.65 | 26.92 | 32.27 | 31.44 |
|  |  | Random | 22.26 | 29.42 | 31.77 | 27.59 | 26.92 | 27.59 |
| | $k = 2$ | Taxonomy | 96.65 | 95.14 | 96.14 | 86.79 | 92.64 | 86.79 |
|  |  | by Environment | 74.58 | 76.91 | 74.42 | 46.49 | 68.06 | 46.32 |
|  |  | Random | 23.25 | 28.10 | 27.09 | 26.42 | 25.08 | 25.75 |
| | $k = 3$ | Taxonomy | 98.82 | 97.99 | 97.32 | 92.64 | 96.82 | 92.64 |
|  |  | Environment | 82.11 | 81.59 | 75.41 | 71.91 | 74.58 | 71.24 |
|  |  | Random | 23.58 | 25.08 | 27.76 | 25.59 | 26.09 | 24.58 |
| | $k = 4$ | Taxonomy | 99.50 | 98.33 | 98.66 | 98.16 | 97.16 | 98.16 |
|  |  | Environment | 83.29 | 84.11 | 82.28 | 78.43 | 75.08 | 80.43 |
|  |  | Random | 25.06 | 23.74 | 27.59 | 25.42 | 26.92 | 23.58 |
| | $k = 5$ | Taxonomy | 99.50 | 98.16 | 99.33 | 97.32 | 97.32 | 98.16 |
|  |  | Environment | 83.27 | 84.76 | 83.29 | 69.23 | 77.26 | 81.77 |
|  |  | Random | 24.08 | 20.23 | 23.07 | 26.09 | 25.42 | 24.25 |
| | $k = 6$ | Taxonomy | 99.50 | 98.50 | 99.33 | 99.16 | 97.49 | 98.83 |
|  |  | Environment | 83.46 | 83.94 | 84.12 | 79.60 | 77.59 | 82.44 |
|  |  | Random | 27.24 | 22.91 | 26.58 | 28.09 | 25.59 | 24.25 |
| pH | $k = 1$ | Taxonomy | 65.2 | 66.70 | 62.37 | 52.69 | 56.99 | 58.06 |
|  |  | by Environment | 56.52 | 58.10 | 51.14 | 54.30 | 53.23 | 54.30 |
|  |  | Random | 51.20 | 53.39 | 50.53 | 49.46 | 53.23 | 50.54 |
| | $k = 2$ | Taxonomy | 95.15 | 93.48 | 95.09 | 84.95 | 91.40 | 84.41 |
|  |  | by Environment | 87.72 | 83.33 | 85.00 | 80.65 | 82.26 | 81.72 |
|  |  | Random | 51.14 | 52.72 | 51.67 | 54.84 | 52.69 | 55.91 |
| | $k = 3$ | Taxonomy | 97.34 | 94.09 | 96.78 | 94.62 | 96.24 | 94.62 |
|  |  | Environment | 90.94 | 90.94 | 90.38 | 81.18 | 83.87 | 80.11 |
|  |  | Random | 44.15 | 52.72 | 55.91 | 54.84 | 46.77 | 44.62 |
| | $k = 4$ | Taxonomy | 97.87 | 96.29 | 96.81 | 93.01 | 95.16 | 97.85 |
|  |  | Environment | 90.44 | 88.80 | 91.58 | 84.95 | 86.02 | 89.78 |
|  |  | Random | 49.42 | 47.84 | 49.01 | 44.62 | 44.62 | 47.85 |
| | $k = 5$ | Taxonomy | 98.42 | 96.81 | 95.79 | 95.70 | 96.24 | 98.92 |
|  |  | Environment | 91.55 | 88.30 | 87.81 | 88.17 | 86.02 | 90.32 |
|  |  | Random | 55.35 | 53.77 | 52.13 | 48.39 | 46.24 | 46.24 |
| | $k = 6$ | Taxonomy | 98.42 | 94.71 | 94.18 | 98.92 | 96.77 | 98.39 |
|  |  | Environment | 91.99 | 88.30 | 86.70 | 92.47 | 84.95 | 92.47 |
|  |  | Random | 47.81 | 49.06 | 50.06 | 50.00 | 45.70 | 46.77 |

The classification accuracies for the *restricted (non-overlapping genera)* case are summarized in Table 4. For *k* = 6, classifications using taxonomy labels for training resulted in high classification accuracies of over 95.30% for the Temperature Dataset and over 91.90% for the pH Dataset, across all six classification models. Classifications using environment category labels resulted in medium classification accuracies of over 61.90% for the Temperature Dataset and medium-high accuracies of over 79.24% for the pH Dataset, across all six classification models. Classifications using randomly assigned labels resulted in the expected low accuracies of at most 27.90% for the Temperature Dataset and at most 55.62% for the pH Dataset, across all six classification models.

**Table 4.** Classification accuracies of six supervised learning classifiers trained on the Temperature Dataset and pH Dataset, in the *restricted* scenario, for three different label assignments (taxonomy, environment category, and random label assignment), and values of 1 ≤*k* ≤6. The classification accuracy in each cell is calculated using stratified 10-fold cross-validation with *non-overlapping genera*.

| Dataset | $k$-value | Class Labelling Type | Classification Model Accuracy (%) | | | | | |
| --- | --- | --- | --- | --- | --- | --- | --- | --- |
|  |  |  | RBF SVM | Random Forest | ANN | MLDSP-1 | MLDSP-2 | MLDSP-3 |
| Temperature | $k = 1$ | Taxonomy | 60.05 | 49.49 | 58.99 | 50.20 | 53.30 | 58.50 |
|  |  | by Environment | 30.87 | 29.72 | 26.38 | 23.70 | 30.80 | 31.30 |
|  |  | Random | 23.91 | 25.12 | 25.15 | 24.20 | 23.40 | 28.30 |
| | $k = 2$ | Taxonomy | 94.11 | 91.12 | 93.79 | 85.80 | 90.80 | 85.60 |
|  |  | by Environment | 57.30 | 53.75 | 54.99 | 33.30 | 48.20 | 33.10 |
|  |  | Random | 22.59 | 27.56 | 25.80 | 24.20 | 24.40 | 24.10 |
| | $k = 3$ | Taxonomy | 98.82 | 95.13 | 97.14 | 87.00 | 94.60 | 87.00 |
|  |  | Environment | 65.57 | 63.25 | 58.10 | 44.80 | 53.30 | 44.50 |
|  |  | Random | 24.93 | 21.40 | 26.12 | 26.60 | 27.10 | 26.60 |
| | $k = 4$ | Taxonomy | 99.16 | 96.13 | 97.81 | 95.00 | 94.50 | 97.20 |
|  |  | Environment | 70.55 | 63.75 | 63.29 | 54.00 | 56.70 | 59.90 |
|  |  | Random | 25.94 | 26.60 | 27.22 | 26.40 | 26.90 | 25.40 |
| | $k = 5$ | Taxonomy | 99.16 | 96.13 | 98.82 | 92.50 | 94.50 | 97.20 |
|  |  | Environment | 72.21 | 64.13 | 66.89 | 50.00 | 62.70 | 65.40 |
|  |  | Random | 26.74 | 23.23 | 22.49 | 24.20 | 26.80 | 26.40 |
| | $k = 6$ | Taxonomy | 99.16 | 96.47 | 97.81 | 99.20 | 95.30 | 98.00 |
|  |  | Environment | 74.17 | 65.48 | 67.88 | 61.90 | 64.70 | 67.90 |
|  |  | Random | 24.20 | 26.74 | 24.59 | 24.90 | 24.10 | 27.90 |
| pH | $k = 1$ | Taxonomy | 65.09 | 67.31 | 62.37 | 51.10 | 50.50 | 58.60 |
|  |  | by Environment | 53.30 | 49.91 | 47.75 | 51.10 | 52.70 | 59.10 |
|  |  | Random | 41.78 | 55.89 | 48.60 | 47.80 | 50.00 | 52.70 |
| | $k = 2$ | Taxonomy | 92.98 | 90.29 | 94.09 | 79.60 | 86.00 | 79.60 |
|  |  | by Environment | 75.09 | 75.15 | 82.66 | 80.60 | 79.60 | 81.20 |
|  |  | Random | 51.52 | 54.14 | 45.79 | 55.90 | 46.80 | 55.90 |
| | $k = 3$ | Taxonomy | 97.37 | 93.54 | 96.78 | 88.70 | 92.50 | 88.20 |
|  |  | Environment | 79.24 | 86.73 | 84.04 | 73.70 | 76.30 | 74.20 |
|  |  | Random | 54.91 | 48.57 | 55.96 | 43.50 | 54.30 | 44.10 |
| | $k = 4$ | Taxonomy | 97.37 | 96.20 | 96.78 | 88.20 | 92.50 | 94.10 |
|  |  | Environment | 81.43 | 83.51 | 85.61 | 73.10 | 79.60 | 80.60 |
|  |  | Random | 46.83 | 41.74 | 47.31 | 52.70 | 48.40 | 49.50 |
| | $k = 5$ | Taxonomy | 97.89 | 97.28 | 96.23 | 94.60 | 92.50 | 96.80 |
|  |  | Environment | 80.91 | 88.83 | 83.01 | 77.40 | 79.60 | 83.90 |
|  |  | Random | 46.73 | 54.83 | 52.44 | 46.80 | 50.00 | 50.00 |
| | $k = 6$ | Taxonomy | 98.42 | 96.23 | 95.73 | 97.30 | 91.90 | 96.80 |
|  |  | Environment | 83.54 | 86.70 | 79.24 | 81.70 | 80.10 | 86.60 |
|  |  | Random | 53.15 | 48.69 | 55.62 | 47.30 | 50.50 | 52.70 |

In both the restriction-free and the restricted cases, the classification of genomic signatures for *k* = 1 corresponds exactly to a classification based on the G+C content of the sequences (this is due to *k*-mers being counted from a DNA fragment together with its reverse complement). As seen from Table 3, the supervised classification accuracies for *k* = 1 were relatively low for taxonomic classifications, and even lower for the environment category classifications. These results suggest that previous observations^16^ of high G+C content of archaeal tRNA sequences being correlated with DNA stability in high temperature environments (≥60°*C*) may not generalize to pervasive genomic signatures and to larger datasets. This inference is also supported by the single nucleotide composition summary for the datasets in this study, see Supplementary Data S3.

Overall, we first note that for both the Temperature Dataset and the pH Dataset, the classification accuracies improved with higher values of *k*. Second, we observe that for both datasets, the classification accuracies obtained when using a random label assignment were approximately equal to the probabilities that a sequence had one of the environment category labels (around 25% in the case of the four temperature labels, and around 50% in the case of the two pH labels). Third, note that the classification accuracies in the restricted scenario were slightly lower than in the restriction-free scenario, for both the taxonomic and the environment category classifications. This decrease could be partly attributed to the decrease in the amount of training data in the restricted scenario. This being said, even in the restricted scenario, the environment category classification accuracies were significantly higher than those for the random label assignment scenario.

Most importantly, these supervised machine learning classification experiments suggest the presence of an environmental component in the genomic signature of temperature and pH microbial extremophiles, able to provide discriminating power for *k*-mer values 3 ≤*k* ≤6. This environmental component of the genomic signature appears to co-exist with a stronger taxonomic component, able to provide discriminating power for *k*-mer values 2 ≤ *k* ≤ 6.

### Sets of k-mers relevant to environment category classifications computed by interpretablity tool of supervised learning algorithm

Of the six supervised classifiers used in the previous section, in this section we use the Mean Decrease in Impurity (MDI) algorithm for the Random Forest classifier to compute a global measure of feature importance. This serves as an interpretability tool that provides insight into the relative contribution of each feature (*k*-mer) to the successful classification.

To this end, we first conducted 10-fold cross-validation on a one-vs-all Random Forest classifier, which achieves a specific accuracy for each environment category. In the four computational experiments associated with the Temperature Dataset, the psychrophile category was correctly separated from the other sequences in the Temperature Dataset with 86.31% accuracy, the mesophile category with 71.14% accuracy, the thermophile category with 75.22% accuracy, and the hyperthermophile category with 89.62% accuracy. Similarly, in the two computational experiments associated with the pH Dataset, the alkaliphile category was classified with 86.45% accuracy, and the acidophile category with 83.76% accuracy.

We then used the trained models obtained in these computational experiments in conjunction with Random Forest’s interpretability tool, the MDI algorithm, to compute a *global importance* measure for each *k*-mer (for *k* = 6, the maximum value analyzed) to determine their relative contribution to the one-vs-all environment category classification. This global importance can then be visualized using the Frequency Chaos Game Representation (*f CGR*_*k*_) to identify potential patterns, as seen Figure 1. A visual inspection of Figure 1 suggests that the set of 6-mers that is relevant in distinguishing DNA sequences from a given environment category from the rest of the dataset is specific to that environment category.

**Figure 1.**
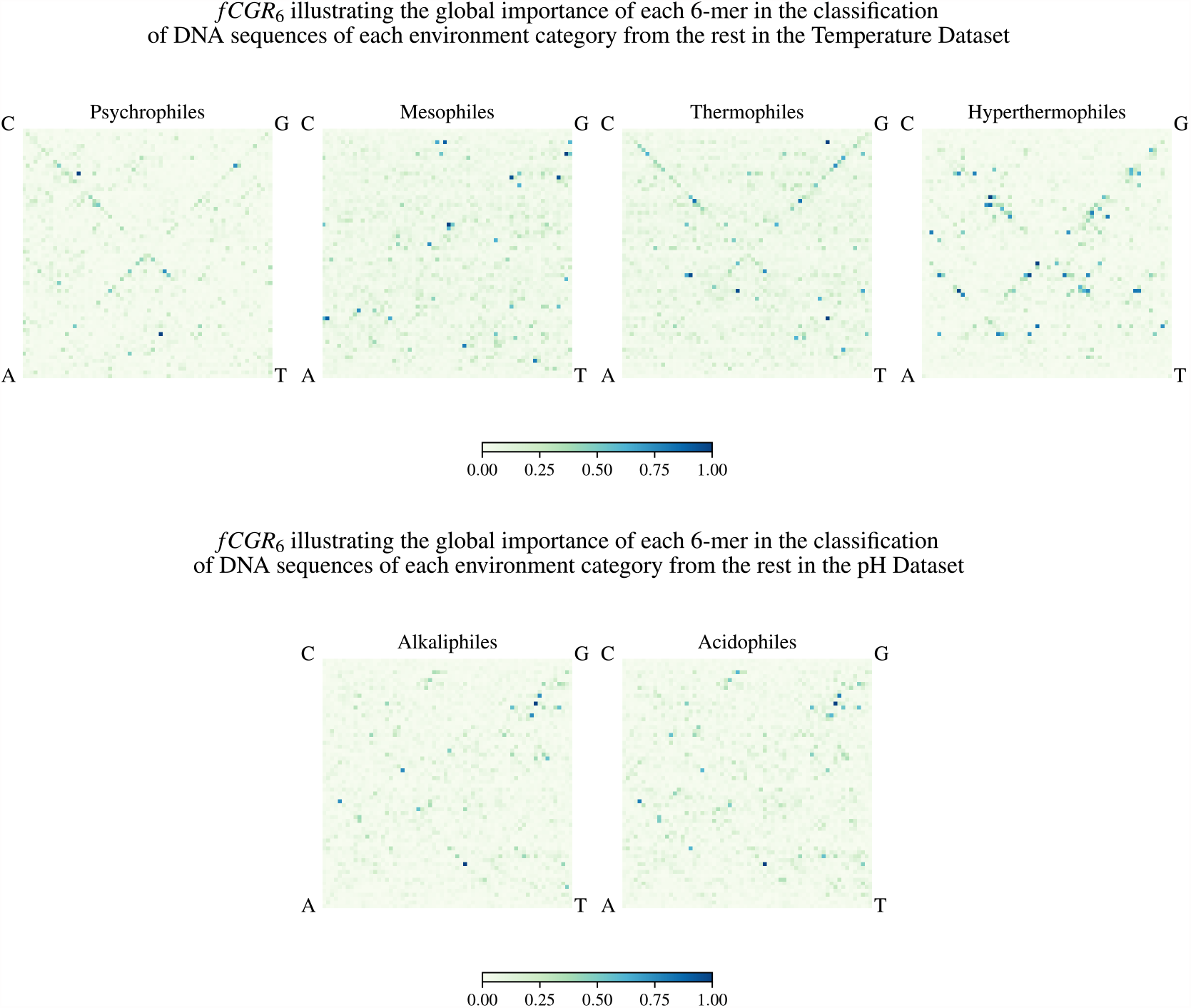
Frequency Chaos Game Representation (*f CGR*_*k*_) of the global importance of various 6-mers in the classification of DNA sequences of each environment category from the rest of the dataset. The top panel shows the *f CGR*_*k*_ for the Temperature Dataset, and the bottom panel shows the *f CGR*_*k*_ for the pH Dataset, both for *k* = 6. The colour and intensity of each pixel represent the relative importance (relevance) of its corresponding 6-mer (dark blue pixels represent the most relevant 6-mers, etc., as described in the colour bar legend).

To confirm these findings and supplement the analysis with previous observations on codon usage patterns and amino acid compositional biases in extremophiles, we also examined the value *k* = 3. Note that not all the 3-mers identified by our method as relevant to the classification are codons, because 3-mers are not counted only from coding sequences or translation frames. For each environment category, the MDI algorithm was used to identify the specific 3-mers that are relevant for each of the one-vs-all Random Forest environment category classifications.

To investigate further the concept of “relevance” and explore its connection with the over-representation and under-representation of codons/amino acids as described in the literature, we computed the histograms of the 3-mers’ deviation from the dataset mean, for each dataset and environment category. Figures 2 and 3 display these histograms, and single out (in green) the 3-mers relevant for each environment category in the Temperature Dataset (Figure 2) and the pH Dataset (Figure 3). To complement this analysis, Tables 5 and 6 list the sets of relevant 3-mers displayed in Figures 2 and 3, respectively, alongside with the relevant literature on biological observations of codon/amino acid compositional biases associated with extreme temperature and pH environments. Note that each set of relevant 3-mers listed in an environment category panel in Figure 2 (Figure 3), ordered left-to-right alphabetically on the *x*-axis of the panel, corresponds to a set of relevant 3-mers in a matching environment category column in Table 5 (Table 6), ordered top-to-bottom alphabetically by the abbreviation of the amino acid they would encode if they were codons.

**Table 5.**
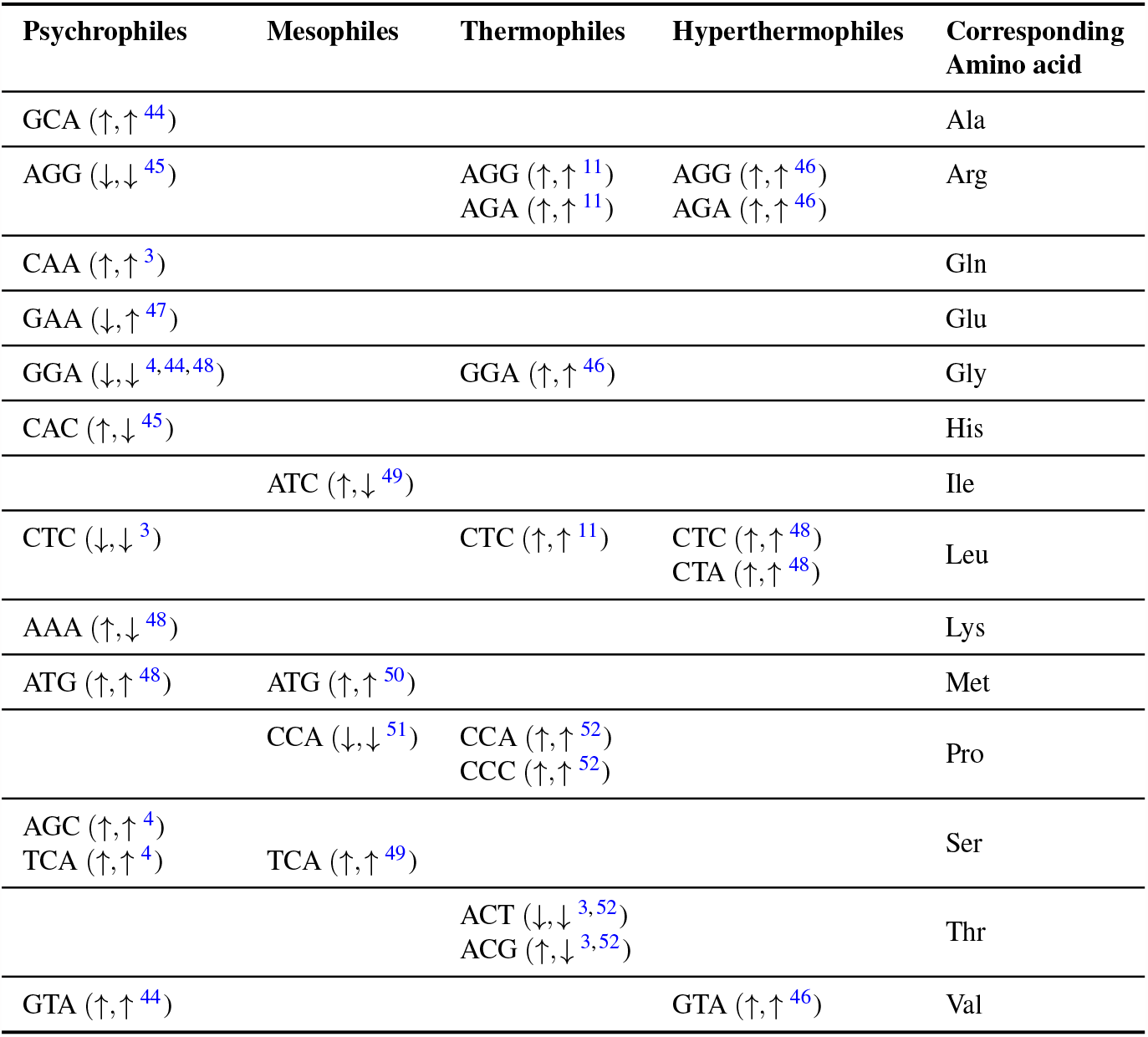
Over- and under-representation of the relevant 3-mers, found by our method to be collectively associated with genomic signatures of temperature-adapted prokaryotic extremophiles. The symbol ↑ (↓) indicates over-representation (under-representation) of a 3-mer/codon. Matched arrows, e.g., (↑, ↑^*re f*^) indicate that both our method and reference *re f* agree in their finding. Mismatched arrows indicate disagreement. See Supplementary Table S4 for details on the observations in biological literature.

**Table 6.** Over- and under-representation of the relevant 3-mers, found by our method to be collectively associated with genomic signatures of pH-adapted prokaryotic extremophiles. The symbol ↑ (↓) indicates over-representation (under-representation) of a 3-mer/codon. Matched arrows, e.g., (↓, ↓^*re f*^) indicate that both our method and reference *ref* agree in their finding. Mismatched arrows indicate disagreement. See Supplementary Table S5 for details of observations in biological literature.

| Alkaliphiles | Acidophiles | Corresponding Amino Acid |
| --- | --- | --- |
| AGG ($\downarrow, \uparrow^{53}$)<br>CGA ($\uparrow, \uparrow^{19}$) | AGG ($\uparrow, \downarrow^{19}$) | Arg |
| AAC ($\uparrow, \uparrow^{19,54}$)<br>AAT ($\downarrow, \uparrow^{19}$) | | Asn |
| GAC ($\uparrow, \uparrow^{19}$) | | Asp |
| CAG ($\uparrow, \uparrow^{19}$) | | Gln |
| CAA ($\downarrow, \downarrow^{19}$) | CAA ($\uparrow, \uparrow^{19}$) | Glu |
| GGA ($\downarrow, \downarrow^{19}$) | GGA ($\uparrow, \uparrow^{19}$) | Gly |
| ATA ($\downarrow, \downarrow^{19}$) | | Ile |
| CTC ($\uparrow, \uparrow^{19}$) | CTC ($\downarrow, \downarrow^{19}$) | Leu |
| ATG ($\downarrow, \downarrow^{19}$) | | Met |
| CCA ($\downarrow, \uparrow^{19}$) | CCA ($\downarrow, \uparrow^{54}$) | Pro |
| TCA ($\uparrow, \downarrow^{19}$) | TCA ($\downarrow, \downarrow^{19}$) | Ser |
| ACG ($\uparrow, \uparrow^{19}$) | | Thr |

**Figure 2.**
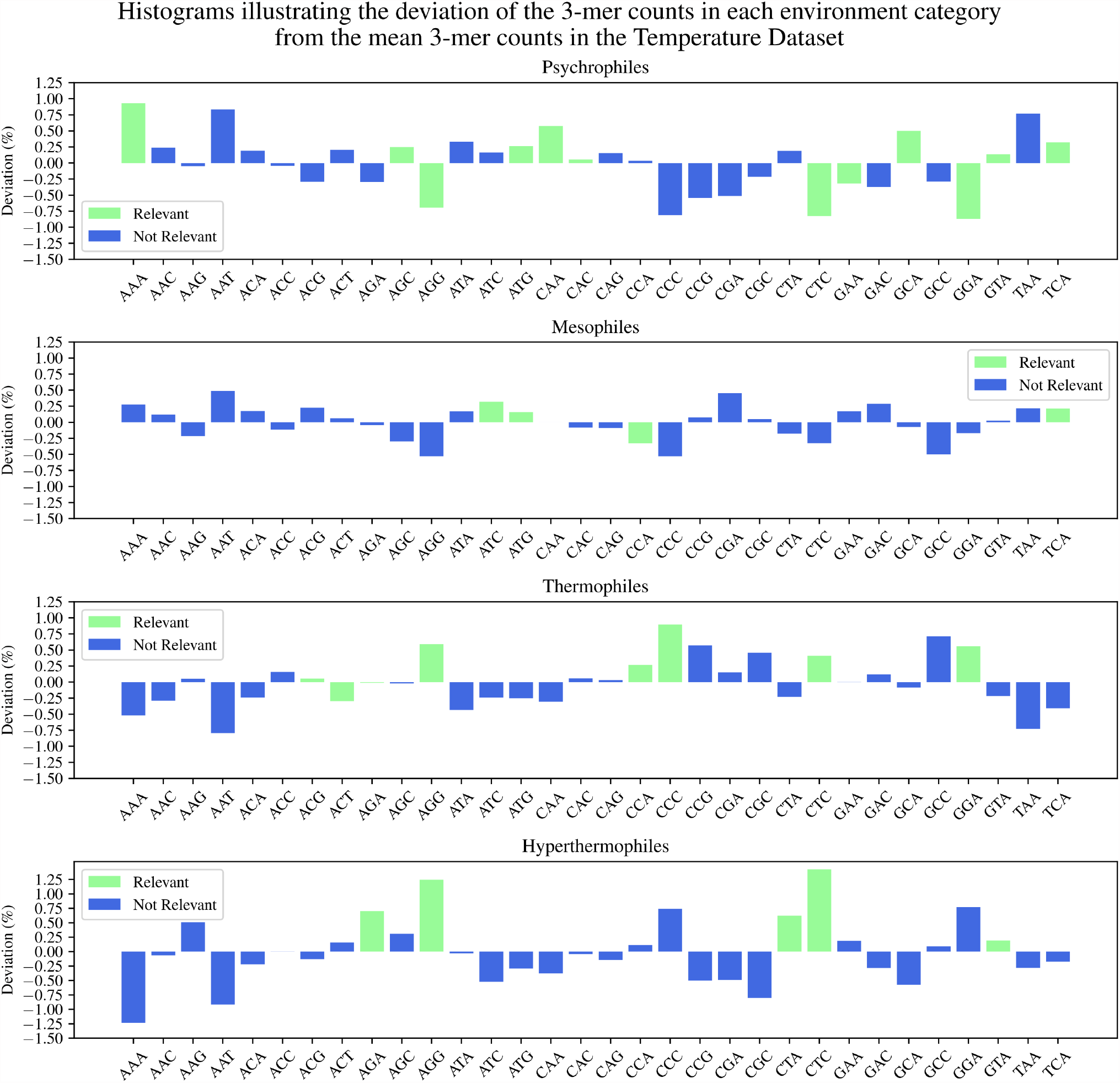
Histograms of the deviation of 3-mer counts in each environment category from the Temperature Dataset mean. A 3-mer and its reverse complement are considered to be indistinguishable, and only canonical 3-mers are listed. Relevant 3-mers for the one-vs-all classification are highlighted in green. The height of each bar represents the difference between a 3-mer’s count in that temperature category and the mean of that 3-mer’s counts over the entire Temperature Dataset (in percentage points).

**Figure 3.**
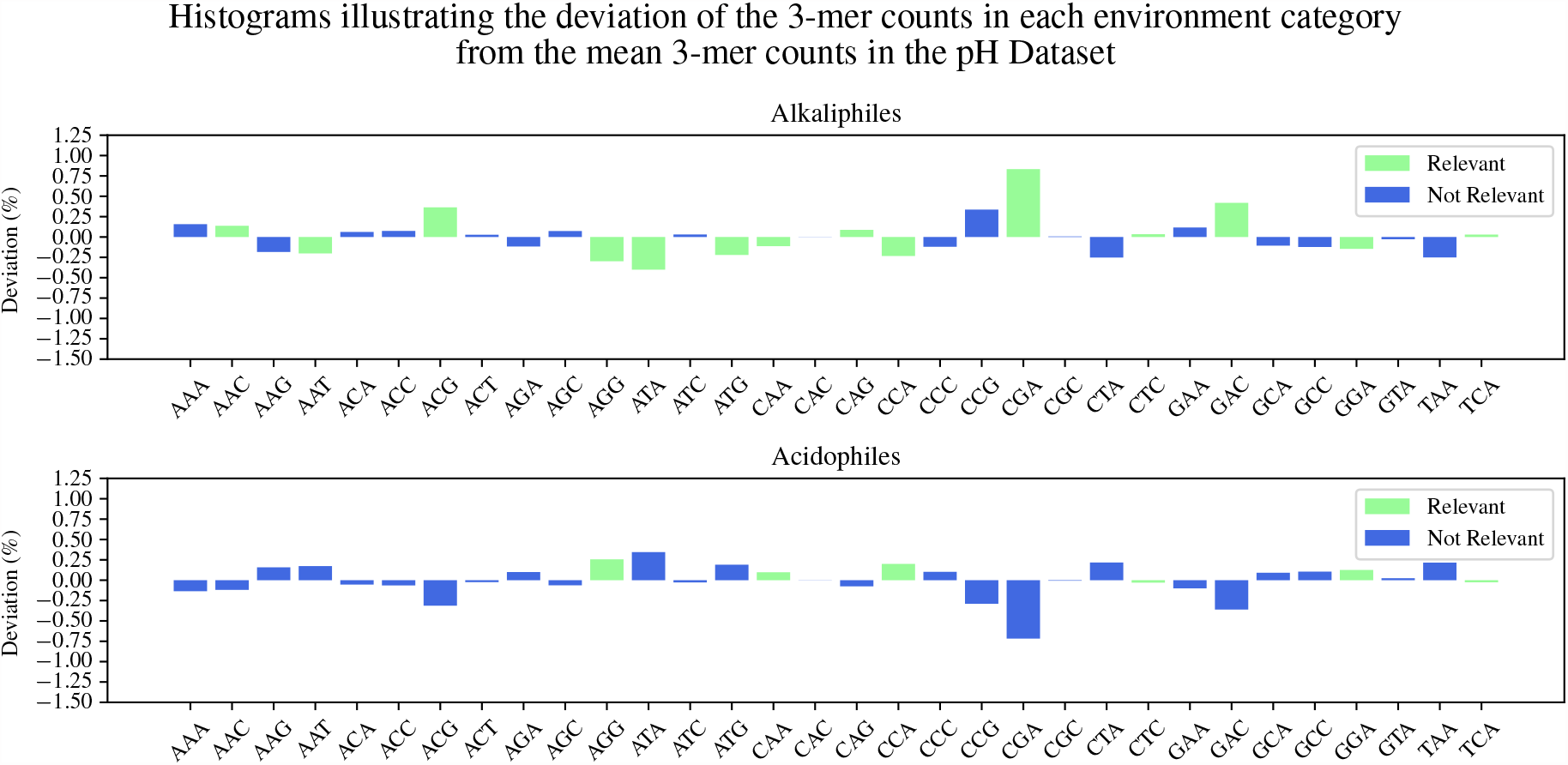
Histograms of the deviation of 3-mer counts in each environment category from the pH Dataset mean. A 3-mer and its reverse complement are considered to be indistinguishable, and only canonical 3-mers are listed. Relevant 3-mers for the one-vs-all classification are highlighted in green. The height of each bar represents the difference between a 3-mer’s count in that pH category and the mean of that 3-mer’s counts over the entire pH Dataset (in percentage points).

As seen in Tables 5 and 6, the majority of our findings regarding over- and under-representation of 3-mers match existing observations in the literature about codon/amino acid bias in extremophiles’ genomic sequences. Disagreements could be due to several factors. First, the 3-mers are not codons: They are counted from an arbitrarily selected 500 kbp DNA fragment representing a genome, and their frequency profile (the genomic signature) has been shown to be quasi-constant along a genome. Thus, some 3-mers could be relevant for the one-vs-all classification of a temperature/pH category in ways that are unrelated to transcriptional or proteomic adaptations. Second, the fact that a 3-mer is found to be relevant for a temperature/pH category indicates that it belongs to a set of 3-mers that *collectively* contribute to distinguishing sequences in that temperature/pH category from the rest of the dataset. In this sense, the concept of “relevant *k*-mer set” is more general, and the fact that a *k*-mer belongs to the relevant set of *k*-mers for a classification does not necessarily imply that it is over- or under-represented in the genomic sequences of that environment category.

### Unsupervised Clustering of the Temperature Dataset and the pH Dataset

The supervised learning computational experiments suggested the existence of an environmental component in the genomic signature of microbial extremophiles, in both a restriction-free scenario and a restricted scenario where sequences from the same genus as the test sequence were absent from training. It should be noted that the datasets considered in this study are not comprehensive, since the discovery and sequencing of genomes of extremophilic organisms is an ongoing difficult process given the challenging environments in which they are found, which are difficult to reproduce in order to culture and further characterize microbial extremophiles^55^. In particular, the datasets’ sparsity and sampling bias do not allow computational experiments in restricted scenarios at taxonomic levels higher than the genus level. This is because such restrictions could eliminate many of the labelled sequences from the cross-validation training sets, rendering them insufficient in size for supervised learning purposes.

To address this challenge, in this section we explore the genomic signatures of the Temperature Dataset and pH Dataset through an unsupervised clustering approach. In unsupervised clustering, no taxonomic or environment category labels for DNA sequences are used during the entire process of learning, and ground-truth labels are used exclusively for the evaluation of the quality of clustering (if applicable). In a first set of tests, we applied *parametric* unsupervised algorithms for the task of clustering both datasets with different values for the parameter `n_cluster` (the number of clusters). When compared to the highest taxonomic level (domain), the ACC measure (Eq.1) for the clustering assignments computed by each algorithm suggest that for the Temperature Dataset, all algorithms can partially cluster sequences according to their real taxonomic labels at `n_clusters` = 2, with *i*DeLUCS (68%) outperforming the others by a small margin (see Table 7 for accuracies). For the pH Dataset, all algorithms are unsuccessful at separating by domain (see Table 8 for accuracies). For values of the parameter `n_clusters` greater than 2, the accuracy increases for both datasets, but the increase is more significant for the pH Dataset

**Table 7.** Accuracies (ACC) of the unsupervised clustering of the Temperature Dataset, for several parametric clustering algorithms, and several values of the pre-specified number of clusters. For each value of the number of clusters parameter, the unsupervised clustering accuracies are computed using the taxonomic labels as ground truth (top row), respectively the environment category labels as ground truth (bottom row).

| No. Clusters | Labelling | Unsupervised Clustering Accuracy (%) |  |  |  |
| --- | --- | --- | --- | --- | --- |
|  |  | K-means | K-medoids | GMM | iDeLUCS |
| 2 | Tax | 63.84 | 63.92 | 63.23 | 68.97 |
|  | Env | 36.27 | 36.50 | 36.26 | 38.23 |
| 4 | Tax | 63.99 | 77.65 | 68.45 | 75.44 |
|  | Env | 34.37 | 40.81 | 38.30 | 48.31 |
| 8 | Tax | 87.81 | 82.13 | 77.79 | 81.48 |
|  | Env | 50.99 | 49.63 | 53.74 | 56.77 |

**Table 8.**
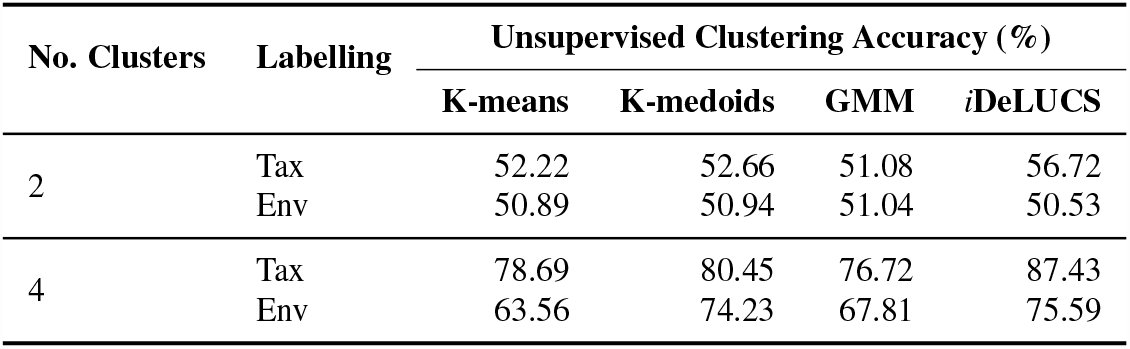
Accuracies (ACC) of the unsupervised clustering of the pH Dataset, for several parametric clustering algorithms, and several values of the pre-specified number of clusters. For each value of the number of clusters parameter, the unsupervised clustering accuracies are computed using the taxonomic labels as ground truth (top row), respectively the environment category labels as ground truth (bottom row).

where the ACC increases by ∼30%, which suggests that there is a good separation by environment category within each domain in the pH Dataset. Overall, the unsupervised clustering accuracy computed using taxonomic labels as ground truth, is higher than when computed using environment category labels as ground truth. This confirms the supervised machine learning results in the previous section, suggesting that the taxonomic component is stronger than the environmental component of genomic signatures.

In a second set of tests, six different *non-parametric* algorithms (the number of clusters is discovered by the algorithm instead of being given as a parameter) and the semi-parametric algorithm *i*DeLUCS were employed to cluster both datasets. Subsequently, all clusters obtained from each algorithm were compared with GTDB labels at the genus level, hereafter referred to as *true genera*, and only those clusters meeting the predefined quality criteria (*>* 50% completeness, and *<* 50% contamination, see Methods) were selected for evaluation. The outcomes, presented in Figure 4, speak to the effectiveness of deep learning clustering methodologies in accurately recovering the true genera, as well as illustrate the importance of choosing appropriate algorithms for specific datasets. For the datasets in this study, the VAE with Iterative Medoids method (VAE+IM) demonstrated superior performance in recovering clusters that meet the predefined quality criteria. Specifically, VAE+IM successfully recovered 61 out of a total of 93 true genera represented by more than two sequences in the Temperature Dataset, and 31 out of a total of 37 true genera represented by more than two sequences in the pH Dataset.

**Figure 4.**
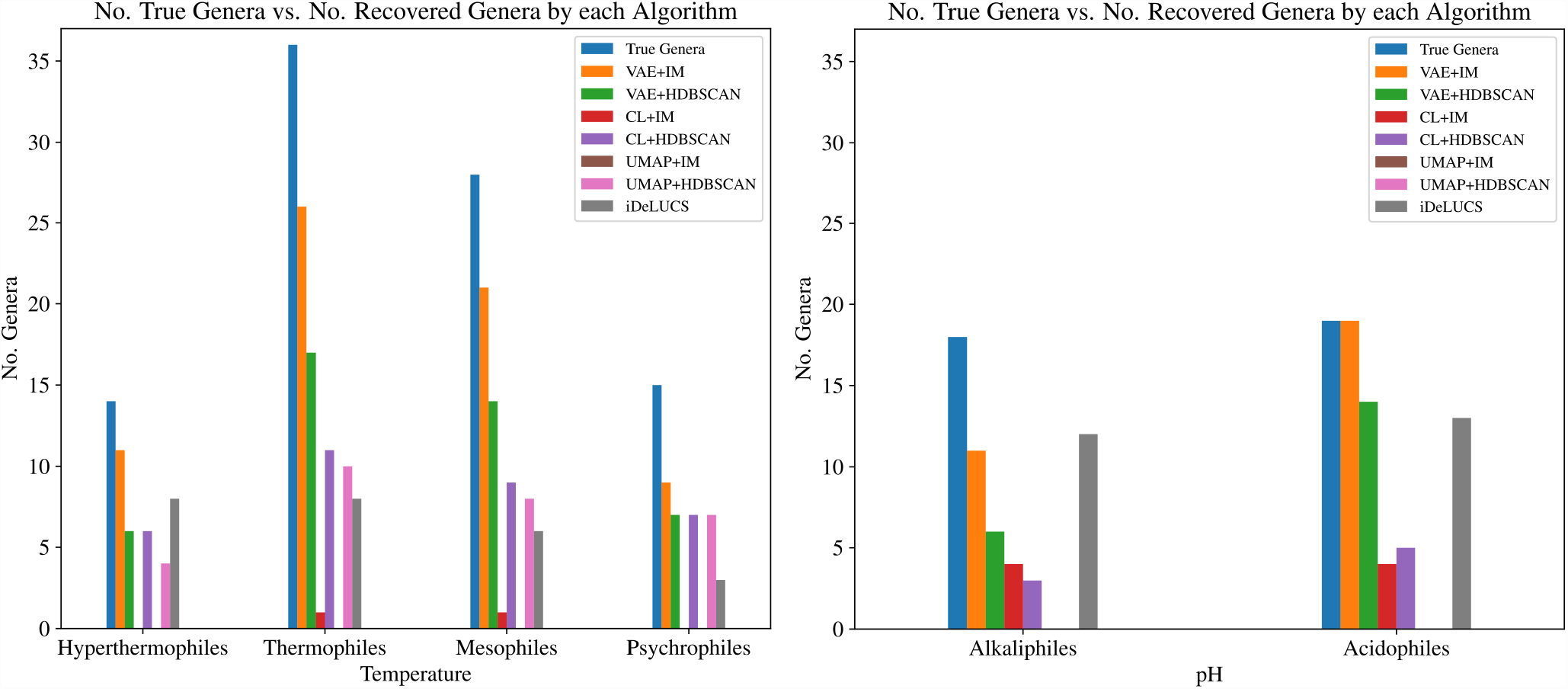
Number of true genera (blue) vs. the number of genera identified by seven clustering algorithms, for each environment category in the Temperature Dataset (left), respectively the pH Dataset (right). Only true genera that are represented by more than two sequences in the respective dataset (Temperature or pH) are considered, and only clusters meeting the quality criteria are counted.

Based on this analysis, the five algorithms that were able to recover at least 20% of the total number of true genera were VAE+HDBSCAN, CL+HDBSCAN, VAE+IM, UMAP+HDBSCAN, and *i*DeLUCS for the Temperature Dataset, respectively VAE+HDBSCAN, CL+HDBSCAN, VAE+IM, CL+IM, and *i*DeLUCS for the pH Dataset. These five algorithms were thus selected as source of information for subsequent analysis, since they performed best when compared to true genera groupings.

### Microbial Extremophiles from Different Domains, with Similar Genomic Signatures

Following the selection of the five top performing unsupervised clustering algorithms in the previous section, the clusters discovered by these algorithms were used in conjunction with a majority voting scheme, to determine concrete “candidates,” that is, concrete exemplars of taxonomically different organisms that were clustered together presumably due to the environmental component of their genomic signatures. This computational process identified a list of pairs of hyperthermophilic, alkaliphilic, and acidophilic *candidate sequences*, each belonging to a different taxonomic domain, which were nevertheless grouped together by the majority of the aforementioned unsupervised clustering algorithms.

Of these candidates, we then proceeded to select sequences for which the unexpected results of the clustering could be independently confirmed by *(i)* supervised machine learning for the prediction of environment category, by *(ii)* supervised machine learning for the prediction of the taxonomic labels, and by *(iii)* observations of shared characteristics of their isolating environments. In these experiments, both thermophiles and hyperthermophiles were treated as part of a single environment category called “high-temperature,” so as to enhance the rigour of the confirmation procedure, given the lack of definitive knowledge of the precise threshold that separates these two environment categories from each other.

The experimental design was aimed to devise challenging scenarios that would clearly demonstrate the presence of the environmental component in the genomic signature of each candidate. To this end, for each candidate sequence to be tested, a challenge training set was created by selecting all DNA sequences of organisms from the opposite domain (i.e., archaea or bacteria), as well as sequences within the same domain but under a different environment category. The classifiers were then trained to perform two different tasks.

In experiments *(i)*, a classifier was trained to predict the environment category of a candidate test sequence, as follows. For instance, if the test sequence was of a hyperthermophilic bacteria, the training set comprised all archaeal sequences (different domain), together with all the mesophilic and psychrophilic bacterial sequences (same domain, different environment category). The objective was to determine if the hyperthermophilic bacterial test sequence would be assigned the correct label “high-temperature,” despite the absence of high-temperature bacterial sequences in the training set. If this were the case, it would indicate that the correct temperature label assignment was due to the similarity of this bacterial sequence to other high-temperature archaeal sequences in the dataset, further suggesting that the environmental component overrides the taxonomical component in the genomic signature of the candidate sequence.

In experiments *(ii)*, a classifier was trained to predict the domain of each candidate test sequence, as follows. For example, if the candidate test sequence was of a hyperthermophilic archaea, the training set comprised all bacteria sequences (different domain), together with all the mesophilic and psychrophilic archaeal sequences (same domain, different environment category). The objective was to determine if the hyperthermophilic archaeal sequence would be assigned the incorrect label “Bacteria.” If this were indeed the case, it would indicate that the assignment of this archaeal sequence to domain Bacteria was likely due to its similarity to the high-temperature bacterial sequences, further suggesting that the environmental component overrides the taxonomic component of the candidate sequence.

All candidate sequences generated by the unsupervised clustering experiment underwent both computational experiments *(i)* and *(ii)*. Of these, the following four sequences were assigned by the majority of the classifiers (SVM, Random Forest, ANN, MLDSP) to the correct environment category in experiment *(i)*, and to the incorrect domain in experiment *(ii)*: the bacterial sequence *Thermocrinis ruber* – Accession ID: GCA_000512735.1, and the three archaeal sequences, *Pyrococcus furiosus DSM 3638 (formerly Pyrococcus sp000211475)* – Accession ID: GCA_000007305.1, *Thermococcus litoralis DSM 5473 (formerly Thermococcus litoralis NS-C)* – Accession ID: GCA_000246985.3, and *Pyrococcus chitonophagus (formerly known as Thermococcus chitonophagus)* – Accession ID: GCA_002214605.1. Note that the current release of Genome Taxonomy Database (GTDB release R214 April 28, 2023) defines *Thermococcus litoralis* as a strain type of species *Thermococcus alcaliphilus*. In this study, we refer to it as “*Thermococcus litoralis*,” given its classification in the database version used for creation of the dataset. Indeed, in experiments *(i)*, all environment-trained classifiers correctly predicted these four microbial sequences as belonging to the high-temperature environment category, in spite of the fact that all genomic sequences used to train the classifier to predict temperature conditions were from a different domain than that of the test sequence. Moreover, in experiments *(ii)* all taxonomy-trained classifiers erroneously predicted the genomic sequences of these microbial extremophiles as belonging to a different domain, likely due to their environmental characteristic.

For biological corroboration *(iii)*, a literature search was undertaken in an attempt to correlate the candidate species to the context of phenotypic traits and the characteristics of the isolating environments. It was determined that few phenotypic traits were congruent between candidates, including gram negative cell walls, OGpH falling within the neutrophilic range (pH 5.0 to 9.0) for each candidate, presence of intergenic sequences, and emissions of light hydrocarbons from the nearby environment^56–63^. However, several more phenotypic traits display dissimilarities between organisms, as described in Supplementary Table S6. The particular environments each of the species was initially isolated from were analyzed in greater detail, and it was found that two Joint Genome Institute’s Genomes OnLine Database-derived (JGI-GOLD) ecosystem classifiers describe the isolating environment of all 4 species, as follows: ID 4027 for *P. furiosus* and *P. chitonophagus*, and ID 3991 for *T. litoralis* and *T. ruber*^57, 64, 65^. The descriptors for these classifiers are “aquatic marine hydrothermal vent” and “aquatic thermal hot springs” respectively^64^. Although these environments are classified differently by JGI-GOLD, as ID 3991 and ID 4027 respectively, the descriptors accurately describe these environments due to the presence of hydrothermal systems^61, 66^. Note that *T. litoralis* (ID 3991) has been recently isolated from the Guaymas Basin, albeit from a geographic site of the Guaymas basin that was different from the isolation site of *P. chitonophagus*^67^ (Supplementary Table S7).

For additional insight, the pairwise distance matrix of all genomic signatures generated by ML-DSP^26^ for each dataset, with *k* = 6, was analyzed. The pairwise distance matrix of the Temperature Dataset revealed that the DNA fragment with the shortest distance from that of *Thermocrinis ruber* (bacterium) belonged to *Thermococcus_A litoralis* (archaea) with a distance value of 0.0327 (the distance ranges between 0 and 1, with 0 the minimum distance, between identical sequences, and 1 the maximum distance).

## Discussion

We note that the six supervised machine learning algorithms produced highly accurate taxonomic classifications of extremophile prokaryotic genome sequences, and medium to medium-high accurate environment category classifications of the same sequences. These results suggest that, in addition to the taxonomic information present in the genomic signatures of extremophiles, a distinct *k*-mer frequency profile associated with each environment category also exists. Thus, if the bacteria and archaea sequences in the training set are labelled by environment category, then the supervised learning algorithms will likely assign a new sequence to its correct environment category, regardless of its taxonomy. Also note that the classification accuracies obtained when the datasets were taxonomy-labelled and environment category-labelled were both significantly higher than those obtained when the same datasets were assigned random labels. These findings are consistent with the claim that these taxonomic and environment category classifications are not due to chance, and support the hypothesis of the presence of both a taxonomic and an environmental component in the genomic signatures of microbial extremophiles.

Additional analyses revealed that the classification accuracies obtained in restriction-free supervised classification scenarios were higher than those obtained in the restricted (non-overlapping genera) supervised classification scenarios. However, even in the restricted scenario, the accuracies of classifications by the environment category were higher than those in the control “random label” scenario. Together, these findings suggest that the taxonomic component of the genomic signature is stronger than the environmental component, but that the latter is discernible and it provides discriminating power.

Note that, while the subsets of 3-mers relevant for the environment category classification that were identified by the MDI algorithm provide insights into the relations between genomic signatures and extreme environmental conditions, caution should be taken when interpreting the results. This is because the experiment prioritized classification accuracy, and the identified subsets of relevant 3-mers may partially reflect a correlation between taxonomy and environment. In other words, especially due to the bias and sparsity of both datasets, it is likely that some taxonomic information may also have influenced the process of computational discovery of these subsets of relevant 3-mers. This being said, the overlap between the aforementioned subsets of relevant 3-mers and codon usage patterns and amino acid compositional biases found to be associated with extreme environments in the biological literature, still suggest a detectable environmental component of genomic signatures in temperature and pH-adapted microbial extremophiles. Future work is needed to explore the possibility of multiple environmental components influencing the genomic signatures of polyextremophiles.

The use of unsupervised learning algorithms for exploring the space of genomic signatures holds significant value, as these algorithms effectively discover clusters of genomic fragments possessing similar genomic signatures, free from the influence of any human annotations. Since the precise definition of the term “genomic signature” entails differentiation of genetically distant organisms from each other, a high-performing clustering algorithm should primarily yield clusters corresponding to the true genera within the dataset. That being said, ascertaining causality for fragments assigned to erroneous clusters proves challenging, given the potential for similar genomic signatures to coincide with taxonomic information at a lower level, as well as the inherent systematic errors in each algorithm. For that reason, in the present study, the identification of pairs exhibiting a similar environmental component in their genomic signature based on the clustering assignments, relies predominantly on the consensus of the high-performing clustering algorithms. Furthermore, only pairs of fragments originating from organisms in different domains were retained. Additional confirmation steps by supervised learning in challenging scenarios were applied to the remaining pairs, and four hyperthermopilic exemplars successfully passed all these stringent tests. It is thus possible that other candidates from the list identified by unsupervised clustering could be viable, such as pairs for which only some of the supervised tests yielded successful results. One such example is the pair of acidophilic organisms *Thermoanaerobacterium thermosaccharolyticum* (bacterium) and *Caldisphaera lagunensis* (archaea) in the pH Dataset, which were clustered together in spite of their domain-level taxonomic differences. Further analysis is needed to confirm such additional pairs, by, e.g., an analysis that utilizes, as a genome representative, multiple DNA fragments combined into a single genomic signature.

The dataset in this paper is, to the best of our knowledge, the largest and most comprehensive to date for the study of the genomic signatures of extremophilic microbes. Larger and more balanced datasets, combined with extensive literature and database searches, could yield more nuanced bioinformatic analyses in future studies.

## Conclusion

This paper demonstrates the successful application of supervised machine learning algorithms for highly accurate taxonomic classifications of extremophile prokaryotic genome sequences, and medium to medium-high accurate classifications of the same sequences based on their environment category (hyperthermophile, psychrophile, acidophile, alkaliphile, etc). The use of *k*-mer frequency vectors of arbitrarily selected 500 kbp DNA fragments as genomic signatures, reveals a strong taxonomic component for 2≤*k*≤6, and a discernible environmental component for 3≤*k*≤6. Furthermore, specific *k*-mer profiles associated with distinct environment categories are identified, with partial agreement with previous observations in literature using alignment-based analyses. Finally, these findings are confirmed using unsupervised learning clustering algorithms, which also reveal specific exemplar organisms whose environmental component appears to be as strong as the taxonomic component of their genomic signature. This multi-pronged approach, applied to a substantial dataset, significantly strengthens the hypothesis of an environmental component in the genomic signature of microbial extremophiles adapted to extreme temperature or pH environmental conditions.

## Supporting information

Supplementary Information Guide

## Acknowledgements

The authors thank Flora Li and Mohammed Essa for their assistance in the process of data collection, and the anonymous Reviewers for their insightful and constructive suggestions. This work has been supported by the Natural Sciences and Engineering Research Council of Canada Discovery Grants [RGPIN-2023-03663 to L.K, R3511A12 to K.A.H, and RGPIN-2022-03547 to G.R], and Compute Canada Research Platforms & Portals Grant [616 to K.A.H].

## Author contributions statement

L.K, K.A.H., M.S. designed the computational experiments. J.B. collected and curated the data, P.M.A. and G.R. conducted the computational experiments. J.B, K.A.H, M.S. provided biological interpretation of the findings. P.M.A., L.K., K.A.H., J.B., and G.R. contributed to the writing of the manuscript, and all authors analysed the results and reviewed the manuscript.

## Additional information

**Supplementary Information** accompanying this paper will be made available online, upon publication.

### Data Availability

All sequence data used in this paper is publicly available for download at NCBI. The unique assembly accession ids of all the sequences and their respective labels used in this study are listed in Supplementary Material, Table S1. The representative DNA fragments used in this paper are available at https://github.com/Kari-Genomics-Lab/Extreme_Env/.

