## Supplementary Information Guide for "Environment and taxonomy shape the genomic signature of prokaryotic extremophiles"

**Supplementary Data S1**: Describes the proportions of both the temperature and pH dataset described in terms of genus, organized by domain (bacteria, archaea) and by extremophile class (psychrophiles, mesophiles, thermophiles, hyperthermophiles, acidophiles, alkaliphiles)

**Supplementary Data S2**: Describes the phylogenetic tree for species included in both the temperature and pH dataset, organized by domain (bacteria, archaea)

**Supplementary Data S3**: Describes the single nucleotide (DNA) composition of extreme temperature and extreme pH adapted microbial genomes, organized by genera. Purple colored genera describe bacteria, while black colored genera describe archaea.

**Supplementary Table S1**: Describes the complete dataset composition, including assembly numbers, extremophilic classification, genome size, domain, phylum, class, order, family, genus, species name, NCBI Taxonomic ID, NCBI species name, and link to referenced literature for each species entry. Similar tables are also accessible as .tsv files through the Temperature and pH links.

**Supplementary Table S2.** Describes the assembly metadata associated with each genome assembly included in the curated dataset.

**Supplementary Table S3**: Describes accuracies obtained by computational experiments for supervised training with an alternative sampling of the random pseudo-labels.

**Supplementary Table S4**: Describes observations of codon usage bias or amino acid compositional patterns in extremophiles and link to paper for each reference named in Table 5.

**Supplementary Table S5**: Describes observations of codon usage bias or amino acid compositional patterns in extremophiles and link to paper for each reference named in Table 6.

**Supplementary Table S6**: Compares several phenotypic traits between candidate species of convergent genomic signatures due to environment rather than taxonomic relationships.

**Supplementary Table S7**: Compares the initial independent isolation and co-isolation of *P. chitonophagus* and *T. litoralis* in Guaymas Basin.
