## Supplementary Data S1 for "Environment and taxonomy shape the genomic signature of prokaryotic extremophiles"

### Supplementary Data S1
### Distribution of Genera–Temperature Dataset

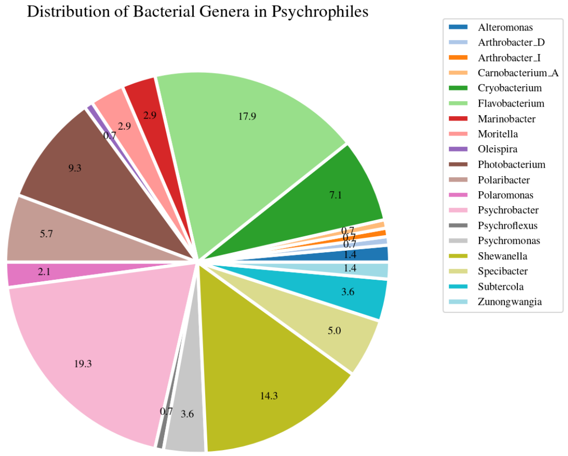


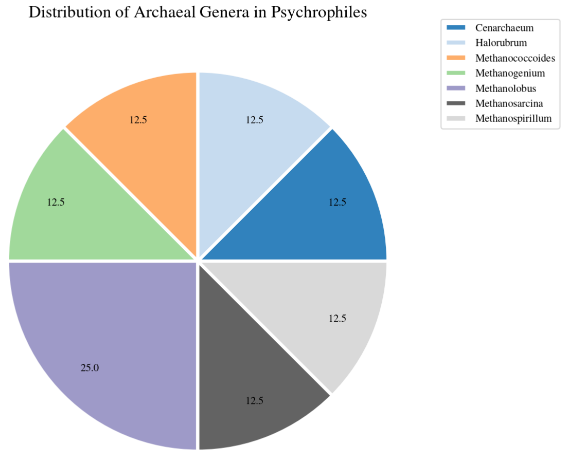


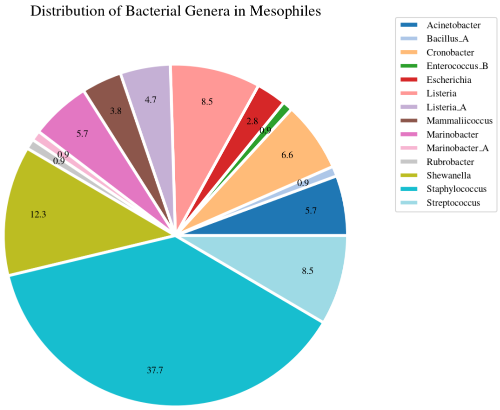


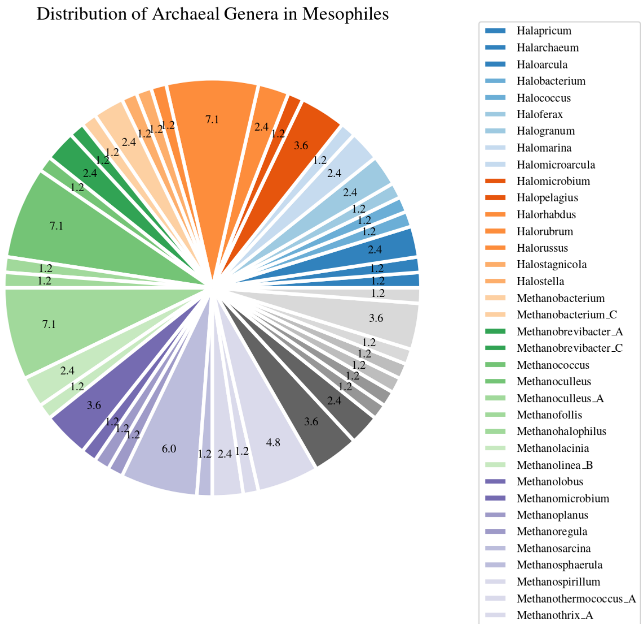

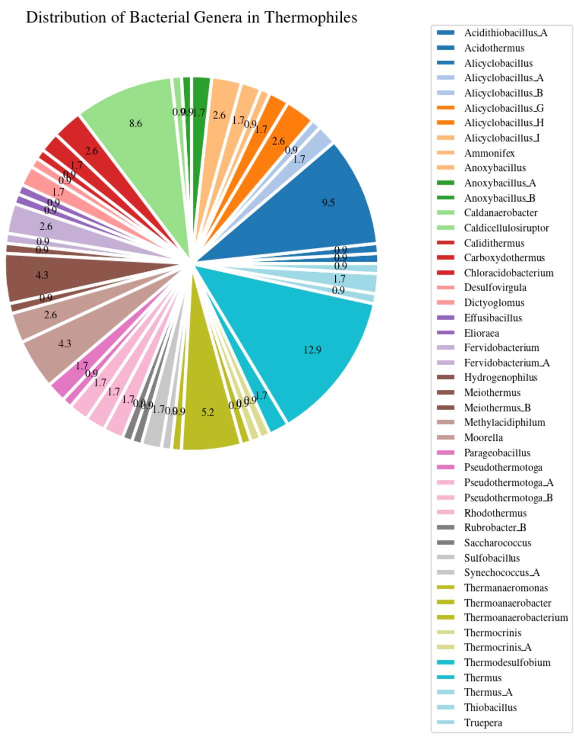
Distribution of Bacterial Genera in Thermophiles
Acidithiobacillus_A
Acidothermus
Alicyclobacillus
Alicyclobacillus_A
Alicyclobacillus_B
Alicyclobacillus_G
Alicyclobacillus_H
Alicyclobacillus_I
Ammonifex
Anoxybacillus
Anoxybacillus_A
Anoxybacillus_B
Caldanaerobacter
Caldicellulosiruptor
Calidithermus
Carboxydothermus
Chloracidobacterium
Desulfovirgula
Dictyoglomus
Effusibacillus
Elioraea
Fervidobacterium
Fervidobacterium_A
Hydrogenophilus
Meiothermus
Meiothermus_B
Methylacidiphilum
Moorella
Parageobacillus
Pseudothermotoga
Pseudothermotoga_A
Pseudothermotoga_B
Rhodothermus
Rubrobacter_B
Saccharococcus
Sulfobacillus
Synechococcus_A
Thermanaeromonas
Thermoanaerobacter
Thermoanaerobacterium
Thermocrinis
Thermocrinis_A
Thermodesulfobium
Thermus
Thermus_A
Thiobacillus
Truepera

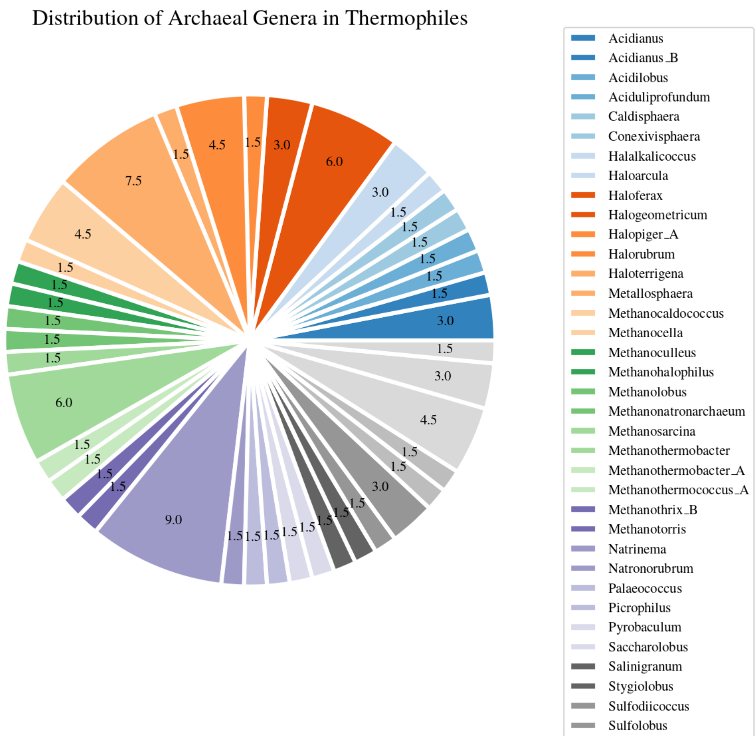
Distribution of Archaeal Genera in Thermophiles
Acidianus
Acidianus_B
Acidilobus
Aciduliprofundum
Caldisphaera
Conexivisphaera
Halalkalicoccus
Haloarcula
Haloferax
Halogeometricum
Halopiger_A
Halorubrum
Haloterrigena
Metallosphaera
Methanocaldococcus
Methanocella
Methanoculleus
Methanohalophilus
Methanolobus
Methanonatronarchaeum
Methanosarcina
Methanothermobacter
Methanothermobacter_A
Methanothermococcus_A
Methanothrix_B
Methanotorris
Natrinema
Natronorubrum
Palaeococcus
Picrophilus
Pyrobaculum
Saccharolobus
Salinigranum
Stygiolobus
Sulfodiicoccus
Sulfolobus

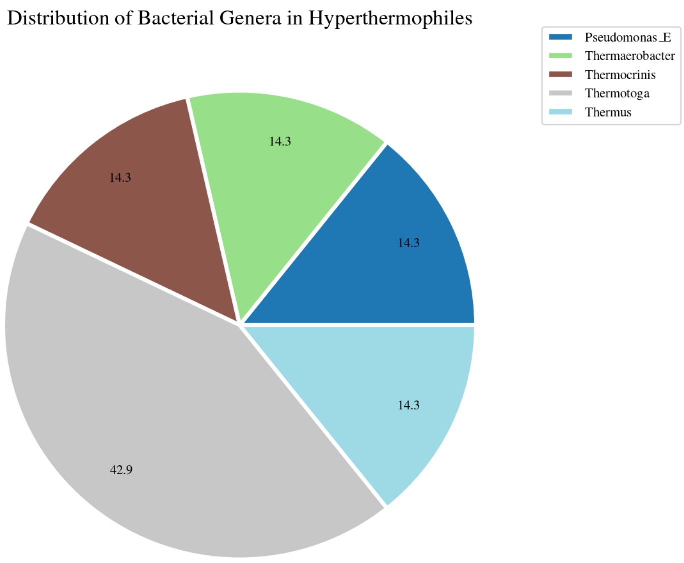
Distribution of Bacterial Genera in Hyperthermophiles
14.3
14.3
14.3
14.3
42.9
Pseudomonas_E
Thermaerobacter
Thermocrinis
Thermotoga
Thermus

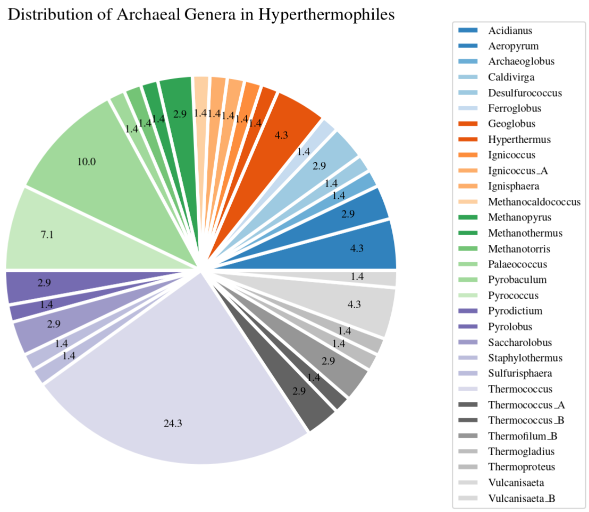
Distribution of Archaeal Genera in Hyperthermophiles
Acidianus
Aeropyrum
Archaeoglobus
Caldivirga
Desulfurococcus
Ferroglobus
Geoglobus
Hyperthermus
Ignicoccus
Ignicoccus_A
Ignisphaera
Methanocaldococcus
Methanopyrus
Methanothermus
Methanotorris
Palaeococcus
Pyrobaculum
Pyrococcus
Pyrodictium
Pyrolobus
Saccharolobus
Staphylothermus
Sulfurisphaera
Thermococcus
Thermococcus_A
Thermococcus_B
Thermofilum_B
Thermogladius
Thermoproteus
Vulcanisaeta
Vulcanisaeta_B

### Distribution of Genera–pH Dataset

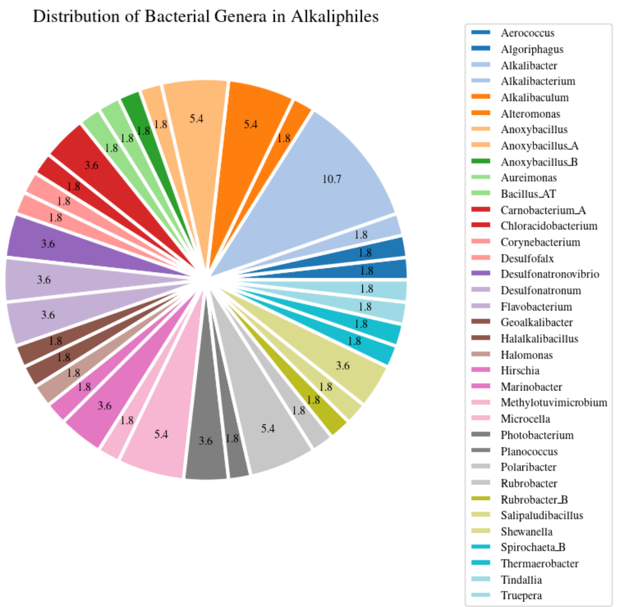


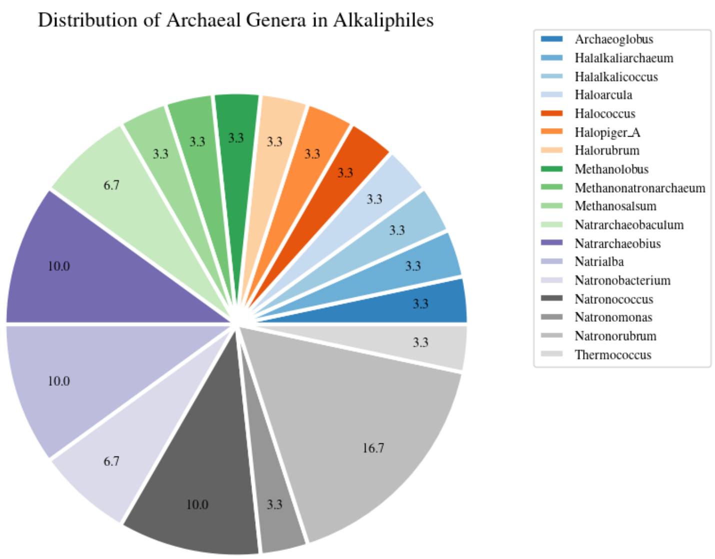


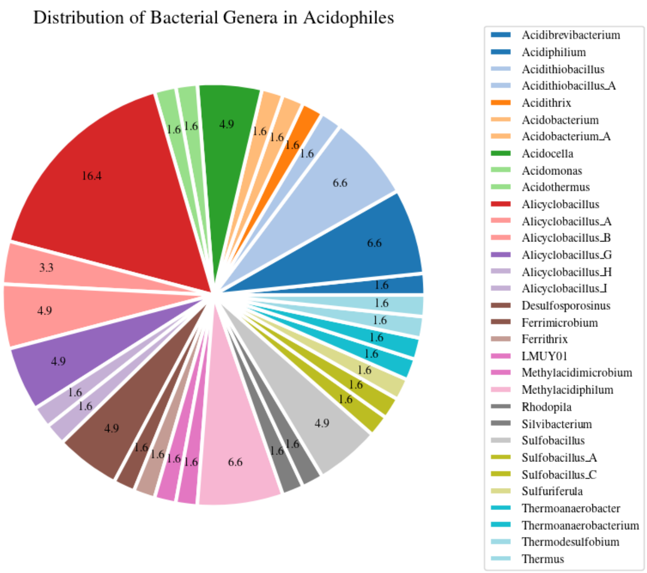


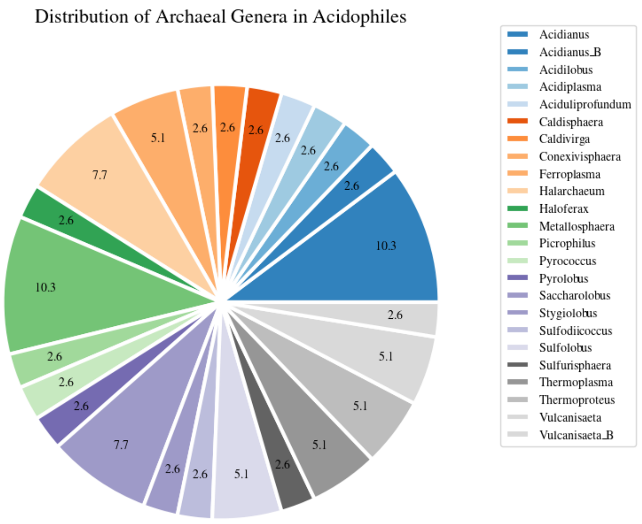
