## Supplementary Data S2 for "Environment and taxonomy shape the genomic signature of prokaryotic extremophiles"

Unrooted Phylogenetic Tree Reconstructed from GTDB
Bacteria - Temperature Dataset

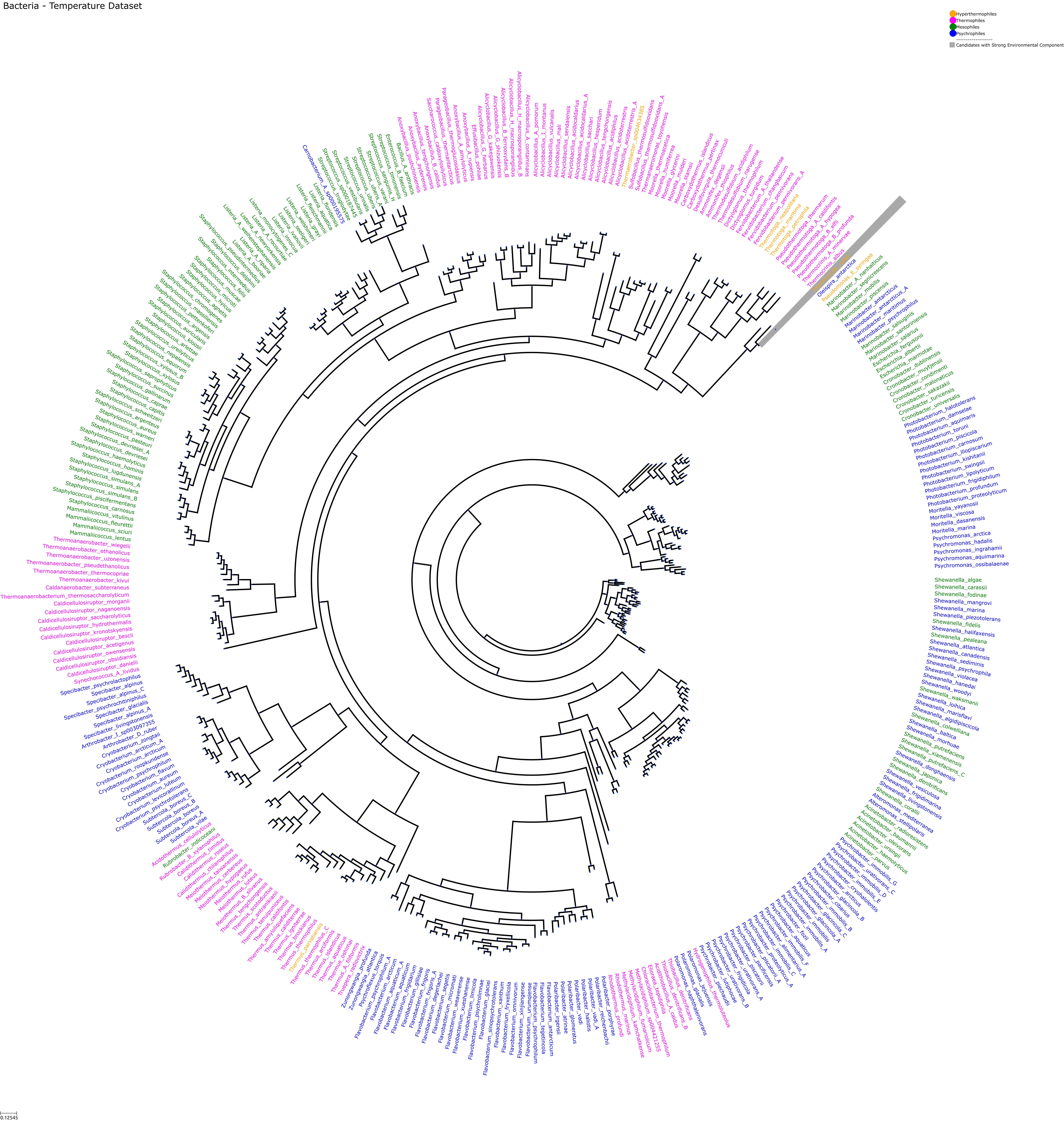

### Unrooted Phylogenetic Tree Reconstructed from GTDB
### Archaea - Temperature Dataset

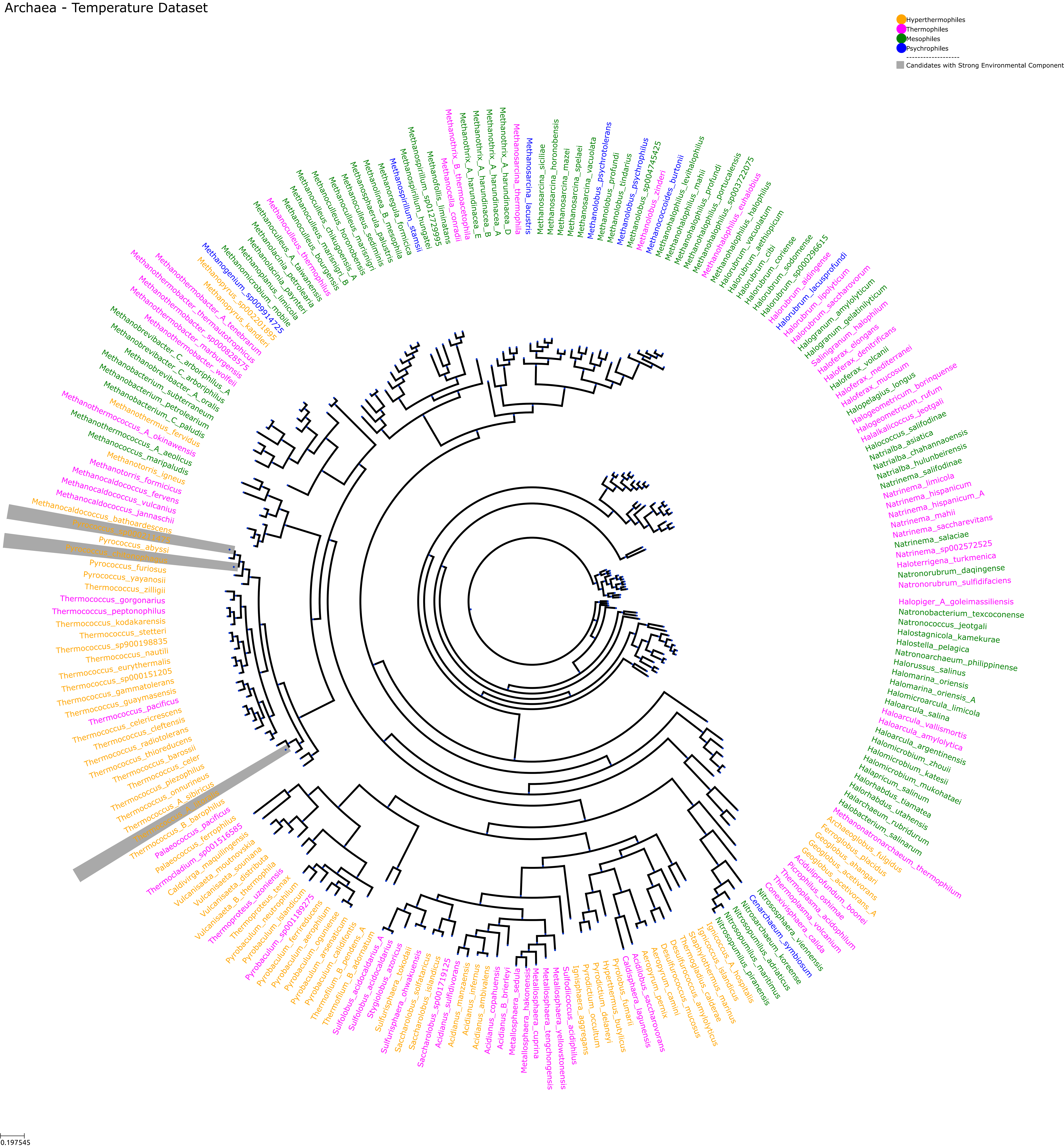

### Unrooted Phylogenetic Tree Reconstructed from GTDB Bacteria - pH Dataset

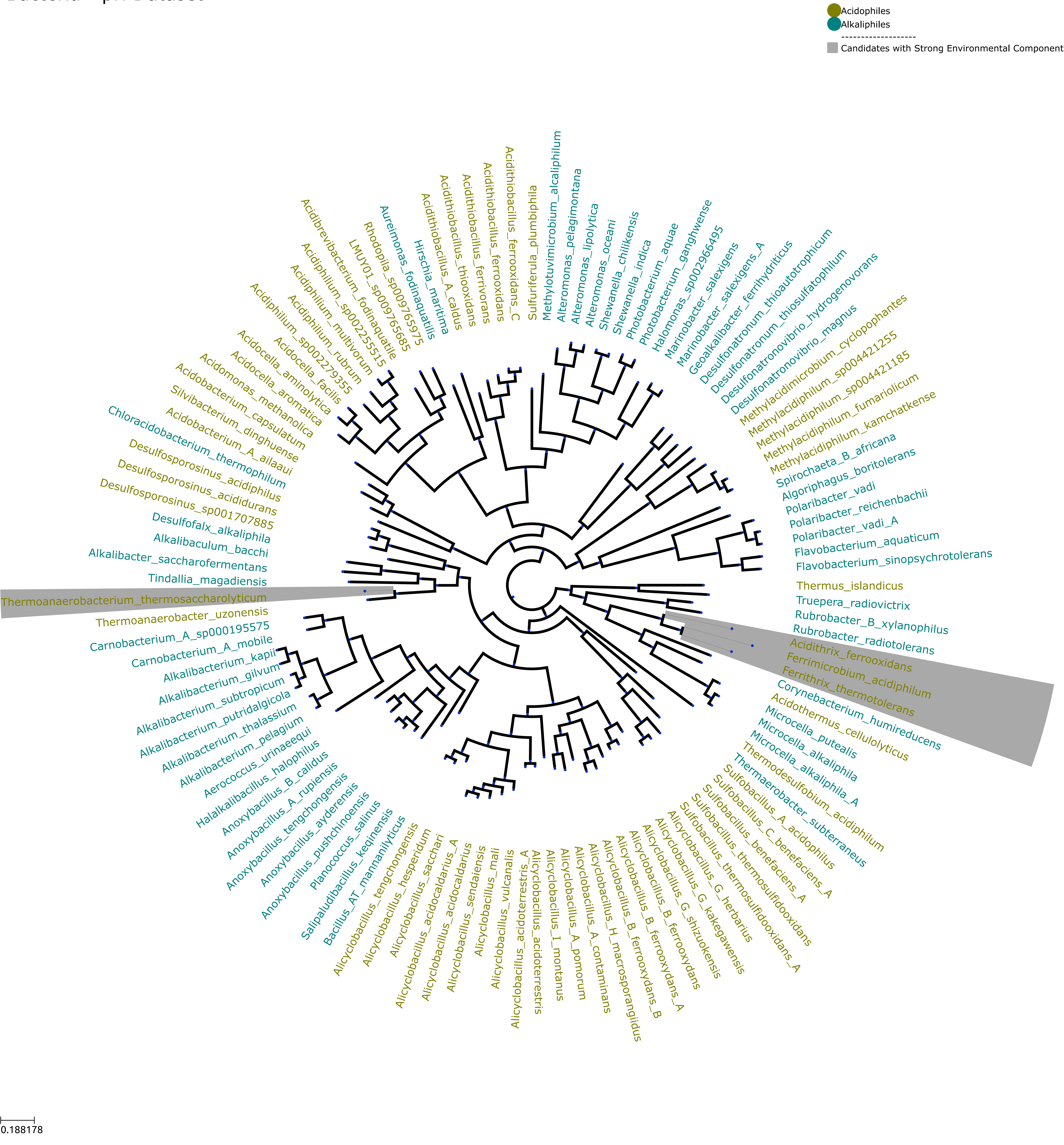

### Unrooted Phylogenetic Tree Reconstructed from GTDB Archaea - pH Dataset

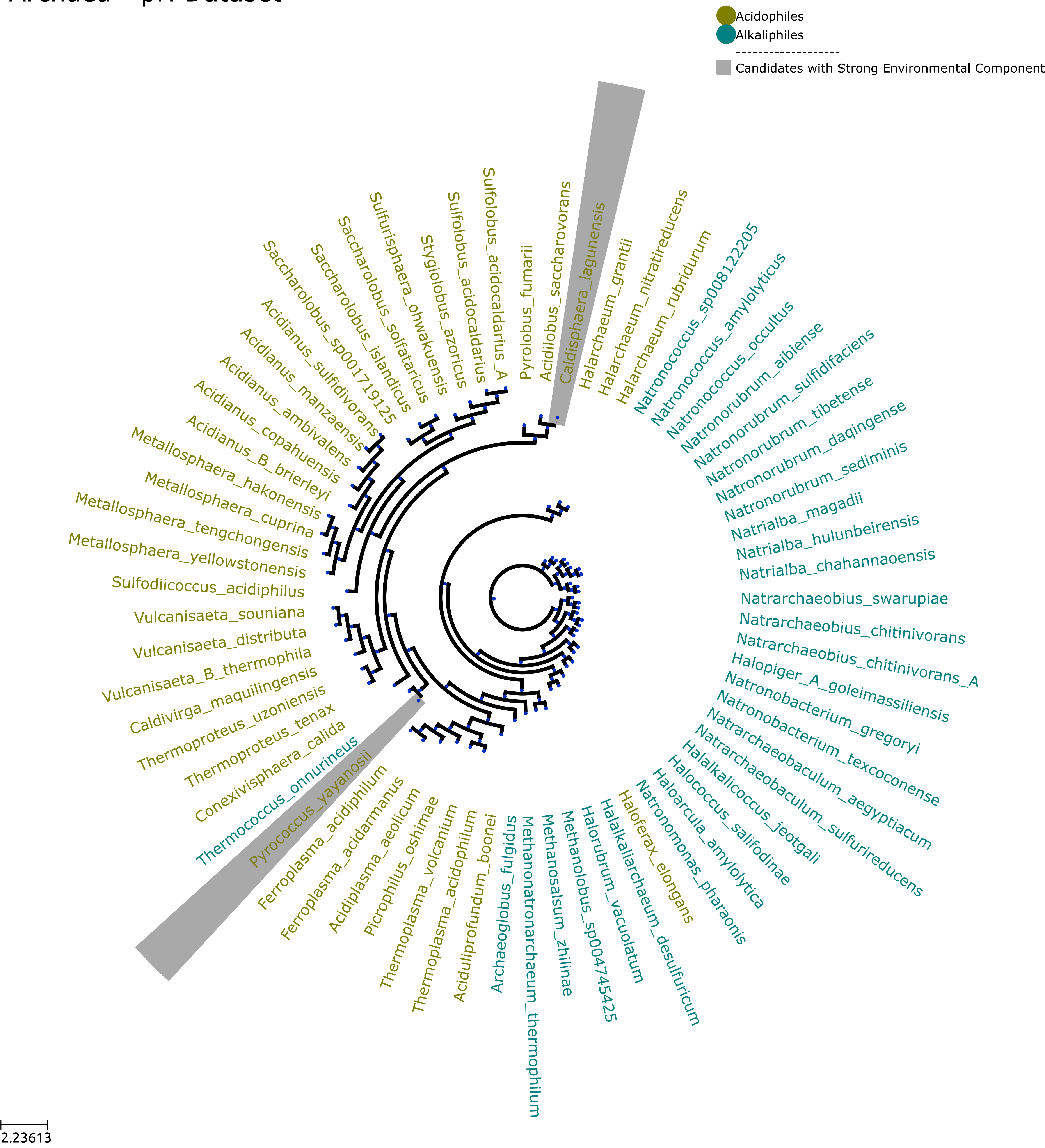
