## Supplementary Data S3 for "Environment and taxonomy shape the genomic signature of prokaryotic extremophiles"

### Supplementary Data S3
### Single Nucleotide Composition - Temperature Dataset

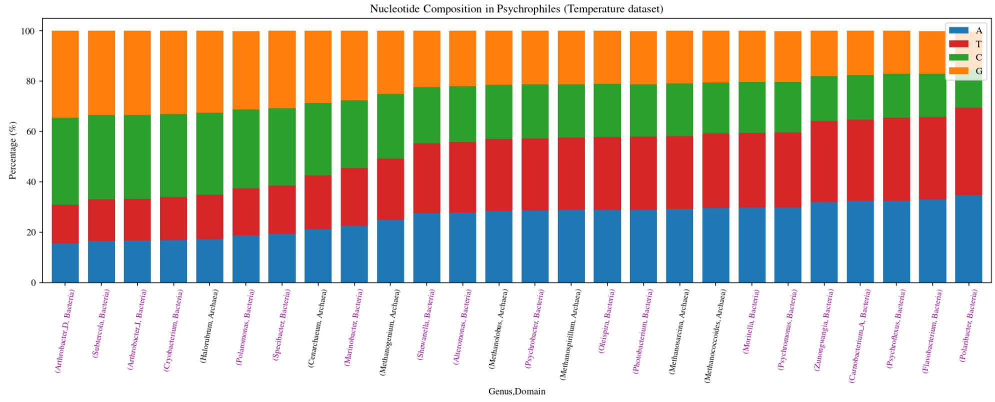


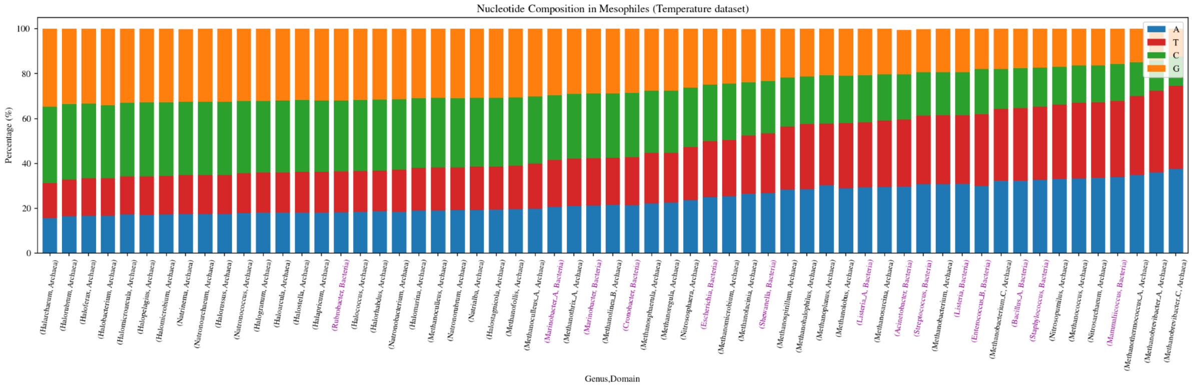


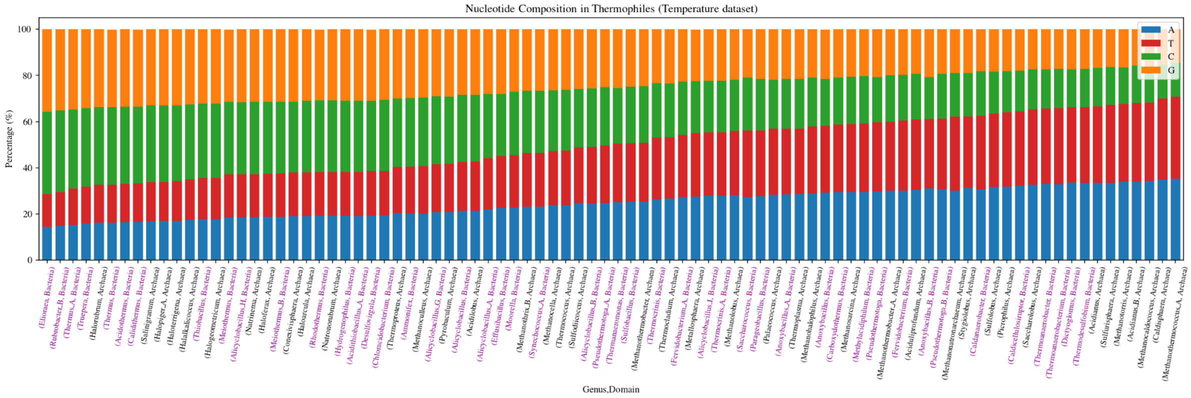


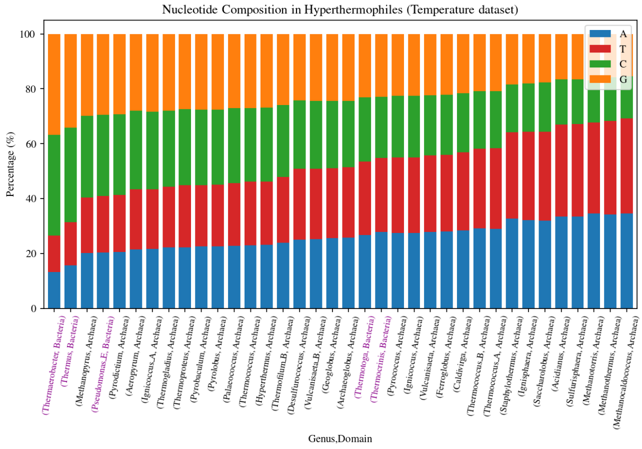

### Single Nucleotide Composition - pH Dataset

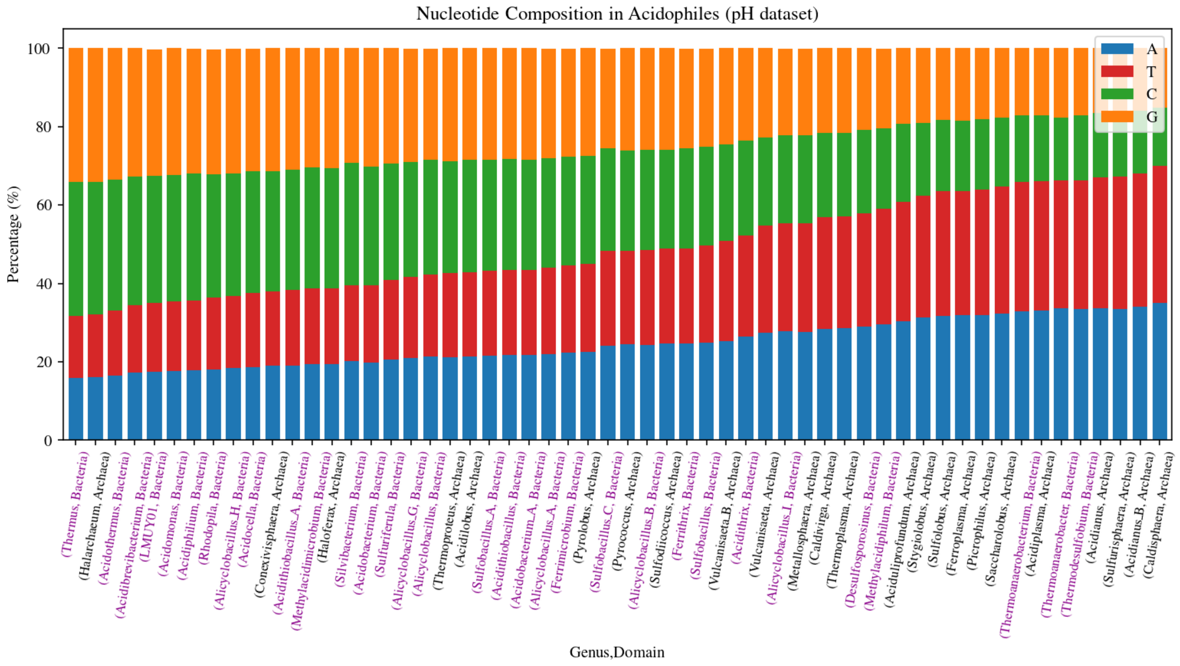


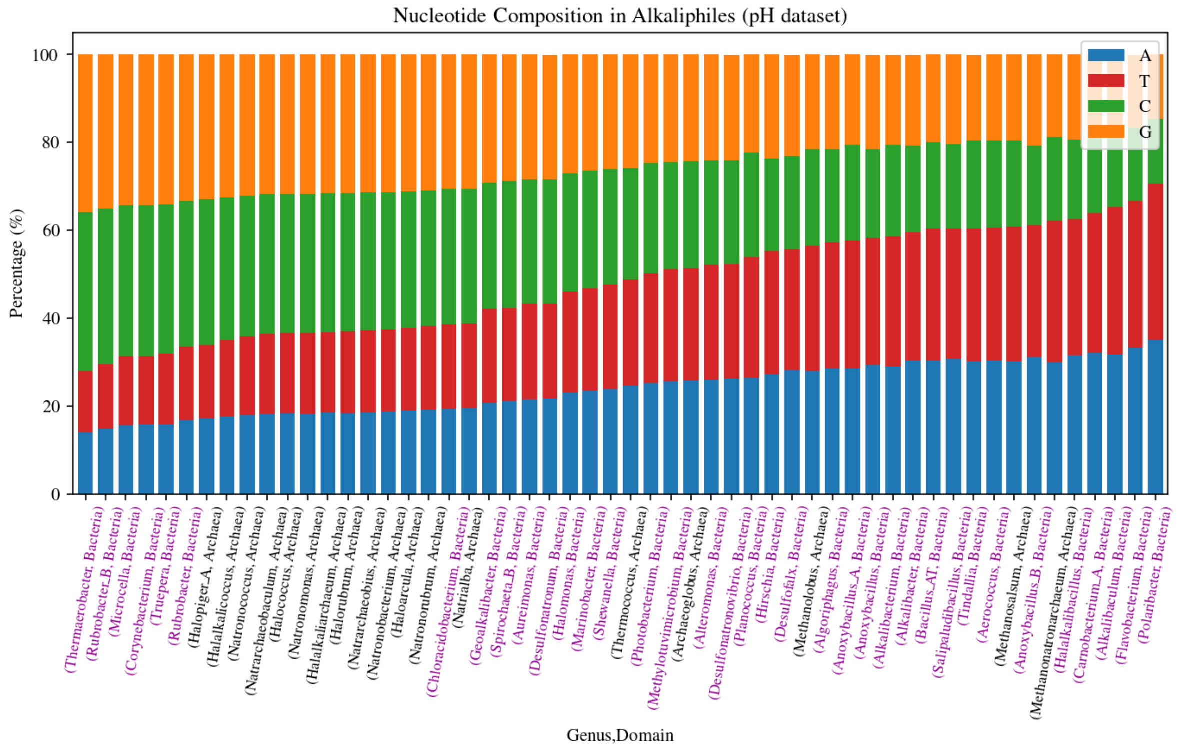
