## Supplementary Table S1 for "Environment and taxonomy shape the genomic signature of prokaryotic extremophiles"

### Supplementary Table S1 - Dataset Composition

| Assembly | Temperature | pH | Genome Size | Domain | Phylum | Class | Order | Family | Genus | Species | NCBI tax ID | *NCBI Species* | Reference Link |
|---|---|---|---|---|---|---|---|---|---|---|---|---|---|
| GCA_009729015.1 | Hyperthermophiles | Acidophiles | 2252027 | Archaea | Thermoproteota | Thermoproteia | Sulfolobales | Sulfolobaceae | Acidianus | *Acidianus ambivalens* | 2283 | *Acidianus ambivalens* | https://doi.org/10.1016/j.bbabio.2003.08.011 / |
| GCA_000632495.1 | Thermophiles | Acidophiles | 2454023 | Archaea | Thermoproteota | Thermoproteia | Sulfolobales | Sulfolobaceae | Acidianus | *Acidianus copahuensis* | 1160895 | *Candidatus Acidianus copahuensis* | https://doi.org/10.1007/s00248-012-0129-4 |
| GCA_009729545.1 | Hyperthermophiles | | 2222414 | Archaea | Thermoproteota | Thermoproteia | Sulfolobales | Sulfolobaceae | Acidianus | *Acidianus infernus* | 12915 | *Acidianus infernus So4a* | https://bacdive.dsmz.de/strain/16642 |
| GCA_002116695.1 | Hyperthermophiles | Acidophiles | 2687463 | Archaea | Thermoproteota | Thermoproteia | Sulfolobales | Sulfolobaceae | Acidianus | *Acidianus manzaensis* | 282676 | *Acidianus manzaensis* | https://doi.org/10.1007/s00284-006-0151-1 |
| GCA_003201765.2 | Thermophiles | Acidophiles | 2265309 | Archaea | Thermoproteota | Thermoproteia | Sulfolobales | Sulfolobaceae | Acidianus | *Acidianus sulfidivorans* | 619593 | *Acidianus sulfidivorans JP7* | https://doi.org/10.1099/ijs.0.64846-0 |
| GCA_003201835.2 | Thermophiles | Acidophiles | 2947244 | Archaea | Thermoproteota | Thermoproteia | Sulfolobales | Sulfolobaceae | Acidianus_B | *Acidianus_B brierleyi* | 41673 | *Acidianus brierleyi* | https://www.sciencedirect.com/science/article/pii/B9780123739445002819 |
| GCA_003352165.1 | | Acidophiles | 4037611 | Bacteria | Proteobacteria | Alphaproteobacteria | Acetobacterales | Acetobacteraceae | Acidibrevibacterium | *Acidibrevibacterium fodinaquatile* | 1969806 | *Acidibrevibacterium fodinaquatile* | https://doi.org/10.1099/ijsem.0.003618 |
| GCA_000144915.1 | Thermophiles | Acidophiles | 1496453 | Archaea | Thermoproteota | Thermoproteia | Sulfolobales | Acidilobaceae | Acidilobus | *Acidilobus saccharovorans* | 666510 | *Acidilobus saccharovorans 345-15* | https://doi.org/10.1128%2FAEM.00599-10 |
| GCA_000202835.1 | | Acidophiles | 4214744 | Bacteria | Proteobacteria | Alphaproteobacteria | Acetobacterales | Acetobacteraceae | Acidiphilium | *Acidiphilium multivorum* | 349163 | *Acidiphilium cryptum JF-5* | https://www.nite.go.jp/en/nbrc/genome/project/annotation/am1.html |
| GCA_900156265.1 | | Acidophiles | 3976993 | Bacteria | Proteobacteria | Alphaproteobacteria | Acetobacterales | Acetobacteraceae | Acidiphilium | *Acidiphilium rubrum* | 1408418 | *Acidiphilium angustum ATCC 35903* | https://doi.org/10.1128/mSystems.00867-20 |
| GCA_902712915.1 | | Acidophiles | 3851483 | Bacteria | Proteobacteria | Alphaproteobacteria | Acetobacterales | Acetobacteraceae | Acidiphilium | *Acidiphilium sp002255515* | 1970291 | *Acidiphilium sp. 20-67-58* | https://doi.org/10.1128%2FmSystems.00867-20 |
| GCA_002279355.1 | | Acidophiles | 3069660 | Bacteria | Proteobacteria | Alphaproteobacteria | Acetobacterales | Acetobacteraceae | Acidiphilium | *Acidiphilium sp002279355* | 1970292 | *Acidiphilium sp. 21-60-14* | https://doi.org/10.1128%2FmSystems.00867-20 |
| GCA_001402945.1 | | Acidophiles | 1778901 | Archaea | Thermoplasmatota | Thermoplasmata | Thermoplasmatales | Thermoplasmataceae | Acidiplasma | *Acidiplasma aeolicum* | 507754 | *Acidiplasma aeolicum* | https://doi.org/10.1099/ijs.0.009639-0 |
| GCA_018853935.1 | | Acidophiles | 4623392 | Bacteria | Proteobacteria | Gammaproteobacteria | Acidithiobacillales | Acidithiobacillaceae | Acidithiobacillus | *Acidithiobacillus ferrivorans* | 160808 | *Acidithiobacillus ferrivorans* | https://doi.org/10.1007/s00792-009-0282-y |
| GCA_000021485.1 | | Acidophiles | 2982397 | Bacteria | Proteobacteria | Gammaproteobacteria | Acidithiobacillales | Acidithiobacillaceae | Acidithiobacillus | *Acidithiobacillus ferrooxidans* | 920 | *Acidithiobacillus ferrooxidans* | https://doi.org/10.1074%2Fmcp.M700042-MCP200 |
| GCA_018854495.1 | | Acidophiles | 3011393 | Bacteria | Proteobacteria | Gammaproteobacteria | Acidithiobacillales | Acidithiobacillaceae | Acidithiobacillus | *Acidithiobacillus ferrooxidans_C* | 920 | *Acidithiobacillus ferrooxidans* | https://doi.org/10.1074%2Fmcp.M700042-MCP200 |
| GCA_009662475.1 | | Acidophiles | 3415726 | Bacteria | Proteobacteria | Gammaproteobacteria | Acidithiobacillales | Acidithiobacillaceae | Acidithiobacillus | *Acidithiobacillus thiooxidans* | 930 | *Acidithiobacillus thiooxidans* | https://doi.org/10.1128/jb.06281-11 |
| GCA_000175575.2 | Thermophiles | Acidophiles | 2987045 | Bacteria | Proteobacteria | Gammaproteobacteria | Acidithiobacillales | Acidithiobacillaceae | Acidithiobacillus_A | *Acidithiobacillus_A caldus* | 637389 | *Acidithiobacillus caldus ATCC 51756* | https://bacdive.dsmz.de/strain/126 |
| GCA_000949295.1 | | Acidophiles | 4019867 | Bacteria | Actinobacteriota | Acidimicrobiia | Acidimicrobiales | Acidimicrobiaceae | Acidithrix | *Acidithrix ferrooxidans* | 1280514 | *Acidithrix ferrooxidans* | https://doi.org/10.1016/j.resmic.2015.01.003 |
| GCA_000022565.1 | | Acidophiles | 4127356 | Bacteria | Acidobacteriota | Acidobacteriae | Acidobacteriales | Acidobacteriaceae | Acidobacterium | *Acidobacterium capsulatum* | 240015 | *Acidobacterium capsulatum ATCC 51196* | https://doi.org/10.1128/AEM.02294-08 |
| GCA_000688455.1 | | Acidophiles | 3686523 | Bacteria | Acidobacteriota | Acidobacteriae | Acidobacteriales | Acidobacteriaceae | Acidobacterium_A | *Acidobacterium_A ailaaui* | 1382359 | *Pseudacidobacterium ailaaui* | https://doi.org/10.1099/ijsem.0.005415 |
| GCA_900129125.1 | | Acidophiles | 3964820 | Bacteria | Proteobacteria | Alphaproteobacteria | Acetobacterales | Acetobacteraceae | Acidocella | *Acidocella aminolytica* | 1120923 | *Acidocella aminolytica 101 = DSM 11237* | https://doi.org/10.1016/S0723-2020(11)80453-4 |
| GCA_014201825.1 | | Acidophiles | 2877351 | Bacteria | Proteobacteria | Alphaproteobacteria | Acetobacterales | Acetobacteraceae | Acidocella | *Acidocella aromatica* | 1303579 | *Acidocella aromatica* | https://doi.org/10.1007/s00792-013-0566-0 |
| GCA_000687875.1 | | Acidophiles | 3403931 | Bacteria | Proteobacteria | Alphaproteobacteria | Acetobacterales | Acetobacteraceae | Acidocella | *Acidocella facilis* | 1214225 | *Acidocella sp. MX-AZ02* | http://dx.doi.org/10.1264/jsme2.2002.98 |
| GCA_004346035.1 | | Acidophiles | 3681251 | Bacteria | Proteobacteria | Alphaproteobacteria | Acetobacterales | Acetobacteraceae | Acidomonas | *Acidomonas methanolica* | 437 | *Acidomonas methanolica* | https://doi.org/10.1099/00207713-39-1-50 |
| GCA_000015025.1 | Thermophiles | Acidophiles | 2443540 | Bacteria | Actinobacteriota | Actinomycetia | Acidothermales | Acidothermaceae | Acidothermus | *Acidothermus cellulolyticus* | 351607 | *Acidothermus cellulolyticus 11B* | https://www.atcc.org/products/43068 |
| GCA_000025665.1 | Thermophiles | Acidophiles | 1486778 | Archaea | Thermoplasmatota | Thermoplasmata | Aciduliprofundales | Aciduliprofundaceae | Aciduliprofundum | *Aciduliprofundum boonei* | 439481 | *Aciduliprofundum boonei* | https://doi.org/10.1007%2Fs00792-007-0111-0 |
| GCA_009759685.1 | Mesophiles | | 3990388 | Bacteria | Proteobacteria | Gammaproteobacteria | Pseudomonadales | Moraxellaceae | Acinetobacter | *Acinetobacter baumannii* | 509170 | *Acinetobacter baumannii SDF* | https://doi.org/10.2166/wst.2018.409 |
| GCA_000369065.1 | Mesophiles | | 3513082 | Bacteria | Proteobacteria | Gammaproteobacteria | Pseudomonadales | Moraxellaceae | Acinetobacter | *Acinetobacter haemolyticus* | 707232 | *Acinetobacter haemolyticus ATCC 19194* | https://bacdive.dsmz.de/strain/137522 |
| GCA_000196795.1 | Mesophiles | | 4152543 | Bacteria | Proteobacteria | Gammaproteobacteria | Pseudomonadales | Moraxellaceae | Acinetobacter | *Acinetobacter oleivorans* | 436717 | *Acinetobacter oleivorans DR1* | https://bacdive.dsmz.de/strain/157136 |
| GCA_000368025.1 | Mesophiles | | 2912642 | Bacteria | Proteobacteria | Gammaproteobacteria | Pseudomonadales | Moraxellaceae | Acinetobacter | *Acinetobacter parvus* | 981333 | *Acinetobacter parvus DSM 16617 = CIP 108168* | https://bacdive.dsmz.de/strain/8139 |
| GCA_006757745.1 | Mesophiles | | 3433938 | Bacteria | Proteobacteria | Gammaproteobacteria | Pseudomonadales | Moraxellaceae | Acinetobacter | *Acinetobacter radioresistens* | 575589 | *Acinetobacter radioresistens SH164* | https://bacdive.dsmz.de/strain/155470 |
| GCA_000368825.1 | Mesophiles | | 3535739 | Bacteria | Proteobacteria | Gammaproteobacteria | Pseudomonadales | Moraxellaceae | Acinetobacter | *Acinetobacter ursingii* | 981336 | *Acinetobacter ursingii DSM 16037 = CIP 107286* | https://bacdive.dsmz.de/strain/155369 |
| GCA_001543205.1 | | Alkaliphiles | 2013339 | Bacteria | Firmicutes | Bacilli | Lactobacillales | Aerococcaceae | Aerococcus | *Aerococcus urinaeequi* | 1120952 | *Aerococcus urinaeequi DSM 20341 = CCUG 28094* | https://bacdive.dsmz.de/strain/222 |
| GCA_000591035.1 | Hyperthermophiles | | 1595994 | Archaea | Thermoproteota | Thermoproteia | Sulfolobales | Acidilobaceae | Aeropyrum | *Aeropyrum camini* | 1198449 | *Aeropyrum camini SY1 = JCM 12091* | https://doi.org/10.1099/ijs.0.02826-0 |
| GCA_000011125.1 | Hyperthermophiles | | 1669696 | Archaea | Thermoproteota | Thermoproteia | Sulfolobales | Acidilobaceae | Aeropyrum | *Aeropyrum pernix* | 56636 | *Aeropyrum pernix* | https://doi.org/10.1099/00207713-46-4-1070 |
| GCA_900108085.1 | | Alkaliphiles | 4834448 | Bacteria | Bacteroidota | Bacteroidia | Cytophagales | Cyclobacteriaceae | Algoriphagus | *Algoriphagus boritolerans* | 1120964 | *Algoriphagus boritolerans DSM 17298 = JCM 18970* | https://bacdive.dsmz.de/strain/17702 |
| GCA_001544355.1 | Thermophiles | | 3865262 | Bacteria | Firmicutes | Bacilli | Alicyclobacillales | Alicyclobacillaceae | Alicyclobacillus | *Alicyclobacillus acidiphilus* | 1255277 | *Alicyclobacillus acidiphilus NBRC 100859* | https://doi.org/10.1016/B978-0-12-802230-6.00016-3 |
| GCA_000024285.1 | Thermophiles | Acidophiles | 3205686 | Bacteria | Firmicutes | Bacilli | Alicyclobacillales | Alicyclobacillaceae | Alicyclobacillus | *Alicyclobacillus acidocaldarius* | 521098 | *Alicyclobacillus acidocaldarius subsp. acidocaldarius DSM 446* | https://doi.org/10.2174/092986608785849209 |
| GCA_000219875.1 | Thermophiles | Acidophiles | 3124048 | Bacteria | Firmicutes | Bacilli | Alicyclobacillales | Alicyclobacillaceae | Alicyclobacillus | *Alicyclobacillus acidocaldarius_A* | 1048834 | *Alicyclobacillus acidocaldarius subsp. acidocaldarius Tc-4-1* | https://doi.org/10.1016/j.fm.2008.07.008 |
| GCA_000444055.1 | Thermophiles | Acidophiles | 4063548 | Bacteria | Firmicutes | Bacilli | Alicyclobacillales | Alicyclobacillaceae | Alicyclobacillus | *Alicyclobacillus acidoterrestris* | 1450 | *Alicyclobacillus acidoterrestris* | https://doi.org/10.1016/B978-0-12-384730-0.00404-3 |
| GCA_007991715.1 | Thermophiles | Acidophiles | 4033884 | Bacteria | Firmicutes | Bacilli | Alicyclobacillales | Alicyclobacillaceae | Alicyclobacillus | *Alicyclobacillus acidoterrestris_A* | 1450 | *Alicyclobacillus acidoterrestris* | https://doi.org/10.1016/B978-0-12-384730-0.00404-3 |
| GCA_900107035.1 | Thermophiles | Acidophiles | 2859702 | Bacteria | Firmicutes | Bacilli | Alicyclobacillales | Alicyclobacillaceae | Alicyclobacillus | *Alicyclobacillus hesperidum* | 89784 | *Alicyclobacillus hesperidum* | https://doi.org/10.1128%2FJB.01612-12 |
| GCA_001570745.1 | Thermophiles | Acidophiles | 2786970 | Bacteria | Firmicutes | Bacilli | Alicyclobacillales | Alicyclobacillaceae | Alicyclobacillus | *Alicyclobacillus mali* | 1314748 | *Alicyclobacillus mali NBRC 102425* | https://doi.org/10.3389/fmicb.2021.639697 |
| GCA_004366795.1 | Thermophiles | Acidophiles | 2946319 | Bacteria | Firmicutes | Bacilli | Alicyclobacillales | Alicyclobacillaceae | Alicyclobacillus | *Alicyclobacillus sacchari* | 392010 | *Alicyclobacillus sacchari* | https://doi.org/10.1099/ijs.0.64692-0 |
| GCA_001552675.1 | Thermophiles | Acidophiles | 2793850 | Bacteria | Firmicutes | Bacilli | Alicyclobacillales | Alicyclobacillaceae | Alicyclobacillus | *Alicyclobacillus sendaiensis* | 1220572 | *Alicyclobacillus sendaiensis NBRC 100866* | http://dx.doi.org/10.1099/ijs.0.02409-0 |
| GCA_001447355.1 | Thermophiles | Acidophiles | 2809442 | Bacteria | Firmicutes | Bacilli | Alicyclobacillales | Alicyclobacillaceae | Alicyclobacillus | *Alicyclobacillus tengchongensis* | 368812 | *Alicyclobacillus tengchongensis* | https://doi.org/10.1007/s12275-014-3625-z |
| GCA_900156755.1 | Thermophiles | Acidophiles | 2994480 | Bacteria | Firmicutes | Bacilli | Alicyclobacillales | Alicyclobacillaceae | Alicyclobacillus | *Alicyclobacillus vulcanalis* | 252246 | *Alicyclobacillus vulcanalis* | https://doi.org/10.1099/ijs.0.03012-0 |
| GCA_000429525.1 | Thermophiles | Acidophiles | 3272318 | Bacteria | Firmicutes | Bacilli | Alicyclobacillales | Alicyclobacillaceae | Alicyclobacillus_A | *Alicyclobacillus_A contaminans* | 1120971 | *Alicyclobacillus contaminans DSM 17975* | https://doi.org/10.3390/microorganisms3040625 |
| GCA_000472905.1 | Thermophiles | Acidophiles | 3398558 | Bacteria | Firmicutes | Bacilli | Alicyclobacillales | Alicyclobacillaceae | Alicyclobacillus_A | *Alicyclobacillus_A pomorum* | 1111479 | *Alicyclobacillus pomorum DSM 14955* | https://bacmedia.dsmz.de/medium/402?ccno=DSM+14955 / https://bacdive.dsmz.de/strain/424 |
| GCA_001399675.1 | | Acidophiles | 5044184 | Bacteria | Firmicutes | Bacilli | Alicyclobacillales | Alicyclobacillaceae | Alicyclobacillus_B | *Alicyclobacillus_B ferrooxydans* | 471514 | *Alicyclobacillus ferrooxydans* | http://dx.doi.org/10.1099/ijs.0.2008/000562-0 |
| GCA_017298655.1 | | Acidophiles | 6027356 | Bacteria | Firmicutes | Bacilli | Alicyclobacillales | Alicyclobacillaceae | Alicyclobacillus_B | *Alicyclobacillus_B ferrooxydans_A* | 471514 | *Alicyclobacillus ferrooxydans (Alicyclobacillus curvatus)* | https://doi.org/10.1099/ijsem.0.005285 |
| GCA_017298635.1 | Thermophiles | Acidophiles | 5293537 | Bacteria | Firmicutes | Bacilli | Alicyclobacillales | Alicyclobacillaceae | Alicyclobacillus_B | *Alicyclobacillus_B ferrooxydans_B* | 471514 | *Alicyclobacillus ferrooxydans (Alicyclobacillus mengziensis)* | https://doi.org/10.1093%2Fg3journal%2Fjkac225 |
| GCA_000430585.1 | Thermophiles | Acidophiles | 3310509 | Bacteria | Firmicutes | Bacilli | Alicyclobacillales | Alicyclobacillaceae | Alicyclobacillus_G | *Alicyclobacillus_G herbarius* | 1120972 | *Alicyclobacillus herbarius DSM 13609* | https://doi.org/10.1099/00207713-52-1-109 |
| GCA_001552655.1 | Thermophiles | Acidophiles | 3421583 | Bacteria | Firmicutes | Bacilli | Alicyclobacillales | Alicyclobacillaceae | Alicyclobacillus_G | *Alicyclobacillus_G kakegawensis* | 1220571 | *Alicyclobacillus kakegawensis NBRC 103104* | https://doi.org/10.1099/ijs.0.64692-0 |
| GCA_001552255.1 | Thermophiles | Acidophiles | 3628929 | Bacteria | Firmicutes | Bacilli | Alicyclobacillales | Alicyclobacillaceae | Alicyclobacillus_G | *Alicyclobacillus_G shizuokensis* | 1220573 | *Alicyclobacillus shizuokensis NBRC 103103* | https://doi.org/10.1099/ijs.0.64692-0 |

| | | | | | | | | | | | | | |
|---|---|---|---|---|---|---|---|---|---|---|---|---|---|
| GCA_900116805.1 | Thermophiles | Acidophiles | 3789420 | Bacteria | Firmicutes | Bacilli | Alicyclobacillales | Alicyclobacillaceae | Alicyclobacillus_H | *Alicyclobacillus_H macrosporangiidus* | 392015 | *Alicyclobacillus macrosporangiidus* | https://doi.org/10.1099/ijs.0.64692-0 |
| GCA_000702485.1 | Thermophiles | | 4091649 | Bacteria | Firmicutes | Bacilli | Alicyclobacillales | Alicyclobacillaceae | Alicyclobacillus_H | *Alicyclobacillus_H macrosporangiidus_B* | 392015 | *Alicyclobacillus macrosporangiidus* | https://doi.org/10.3390%2Fmicroorganisms10030656 |
| GCA_900142255.1 | Thermophiles | Acidophiles | 3045912 | Bacteria | Firmicutes | Bacilli | Alicyclobacillales | Alicyclobacillaceae | Alicyclobacillus_I | *Alicyclobacillus_I montanus* | 1830138 | *Alicyclobacillus montanus* | https://bacdive.dsmz.de/strain/158721 |
| GCA_900128885.1 | | Alkaliphiles | 2331365 | Bacteria | Firmicutes_A | Clostridia | Eubacteriales | Alkalibacteraceae | Alkalibacter | *Alkalibacter saccharofermentans* | 1120975 | *Alkalibacter saccharofermentans DSM 14828* | https://doi.org/10.1007/s00792-004-0390-7 |
| GCA_900109085.1 | | Alkaliphiles | 2181898 | Bacteria | Firmicutes | Bacilli | Lactobacillales | Carnobacteriaceae | Alkalibacterium | *Alkalibacterium gilvum* | 1130080 | *Alkalibacterium gilvum* | https://doi.org/10.1099/ijs.0.042556-0 |
| GCA_007988865.1 | | Alkaliphiles | 1991267 | Bacteria | Firmicutes | Bacilli | Lactobacillales | Carnobacteriaceae | Alkalibacterium | *Alkalibacterium kapii* | 426704 | *Alkalibacterium kapii* | https://bacdive.dsmz.de/strain/2243 |
| GCA_900109325.1 | | Alkaliphiles | 2638902 | Bacteria | Firmicutes | Bacilli | Lactobacillales | Carnobacteriaceae | Alkalibacterium | *Alkalibacterium pelagium* | 426702 | *Alkalibacterium pelagium* | https://bacdive.dsmz.de/strain/2246 |
| GCA_900109825.1 | | Alkaliphiles | 2600402 | Bacteria | Firmicutes | Bacilli | Lactobacillales | Carnobacteriaceae | Alkalibacterium | *Alkalibacterium putridalgicola* | 426703 | *Alkalibacterium putridalgicola* | https://doi.org/10.1099/ijs.0.65602-0 |
| GCA_900112455.1 | | Alkaliphiles | 2491452 | Bacteria | Firmicutes | Bacilli | Lactobacillales | Carnobacteriaceae | Alkalibacterium | *Alkalibacterium subtropicum* | 753702 | *Alkalibacterium subtropicum* | https://doi.org/10.1099/ijs.0.027953-0 |
| GCA_900101165.1 | | Alkaliphiles | 2364962 | Bacteria | Firmicutes | Bacilli | Lactobacillales | Carnobacteriaceae | Alkalibacterium | *Alkalibacterium thalassium* | 426701 | *Alkalibacterium thalassium* | https://doi.org/10.1099/ijs.0.65602-0 |
| GCA_003317055.1 | | Alkaliphiles | 3098941 | Bacteria | Firmicutes_A | Clostridia | Eubacteriales | Alkalibacteraceae | Alkalibaculum | *Alkalibaculum bacchi* | 645887 | *Alkalibaculum bacchi* | http://dx.doi.org/10.1099/ijs.0.018507-0 |
| GCA_001758465.1 | | Alkaliphiles | 5018451 | Bacteria | Proteobacteria | Gammaproteobacteria | Enterobacterales | Alteromonadaceae | Alteromonas | *Alteromonas lipolytica* | 1856405 | *Alteromonas lipolytica* | https://bacdive.dsmz.de/strain/133345 |
| GCA_000020585.3 | Psychrophiles | | 4480937 | Bacteria | Proteobacteria | Gammaproteobacteria | Enterobacterales | Alteromonadaceae | Alteromonas | *Alteromonas mediterranea* | 1300253 | *Alteromonas mediterranea 615* | https://bacdive.dsmz.de/strain/23563 |
| GCA_003731635.1 | | Alkaliphiles | 4987953 | Bacteria | Proteobacteria | Gammaproteobacteria | Enterobacterales | Alteromonadaceae | Alteromonas | *Alteromonas oceani* | 2071609 | *Alteromonas oceani* | https://bacdive.dsmz.de/strain/158616#ref65338 |
| GCA_002499975.2 | | Alkaliphiles | 4314189 | Bacteria | Proteobacteria | Gammaproteobacteria | Enterobacterales | Alteromonadaceae | Alteromonas | *Alteromonas pelagimontana* | 1858656 | *Alteromonas pelagimontana* | https://bacdive.dsmz.de/strain/140746 |
| GCA_001562115.1 | Psychrophiles | | 4904192 | Bacteria | Proteobacteria | Gammaproteobacteria | Enterobacterales | Alteromonadaceae | Alteromonas | *Alteromonas stellipolaris* | 233316 | *Alteromonas stellipolaris* | https://bacdive.dsmz.de/strain/447#ref6058 |
| GCA_000024605.1 | Thermophiles | | 2157067 | Bacteria | Firmicutes_B | Desulfotomaculia | Ammonifexales | Ammonificaceae | Ammonifex | *Ammonifex degensii* | 429009 | *Ammonifex degensii KC4* | https://bacdive.dsmz.de/strain/16789 |
| GCA_003368535.1 | Thermophiles | | 2111834 | Bacteria | Firmicutes_B | Desulfotomaculia | Ammonifexales | Ammonificaceae | Ammonifex | *Ammonifex thiophilus* | 444093 | *Ammonifex thiophilus* | https://bacdive.dsmz.de/strain/16790 |
| GCA_000833605.1 | Thermophiles | Alkaliphiles | 2832347 | Bacteria | Firmicutes | Bacilli | Bacillales | Anoxybacillaceae | Anoxybacillus | *Anoxybacillus ayderensis* | 265546 | *Anoxybacillus ayderensis* | https://pubmed.ncbi.nlm.nih.gov/15388701/ |
| GCA_900111795.1 | Thermophiles | Alkaliphiles | 2625421 | Bacteria | Firmicutes | Bacilli | Bacillales | Anoxybacillaceae | Anoxybacillus | *Anoxybacillus pushchinoensis* | 150248 | *Anoxybacillus pushchinoensis* | https://doi.org/10.1099/00207713-50-6-2109 |
| GCA_014201585.1 | Thermophiles | Alkaliphiles | 2727467 | Bacteria | Firmicutes | Bacilli | Bacillales | Anoxybacillaceae | Anoxybacillus | *Anoxybacillus tengchongensis* | 576944 | *Anoxybacillus tengchongensis* | https://doi.org/10.1099/ijs.0.020834-0 |
| GCA_001634285.1 | Thermophiles | | 3158269 | Bacteria | Firmicutes | Bacilli | Bacillales | Anoxybacillaceae | Anoxybacillus_A | *Anoxybacillus_A amylolyticus* | 294699 | *Anoxybacillus amylolyticus* | https://doi.org/10.1016/j.syapm.2005.10.003 |
| GCA_014196195.1 | Thermophiles | Alkaliphiles | 3705654 | Bacteria | Firmicutes | Bacilli | Bacillales | Anoxybacillaceae | Anoxybacillus_A | *Anoxybacillus_A rupiensis* | 1895648 | *Anoxybacillus sp. UARK-01* | https://doi.org/10.1007/s00284-017-1239-5 |
| GCA_013760845.1 | Thermophiles | Alkaliphiles | 3405528 | Bacteria | Firmicutes | Bacilli | Bacillales | Anoxybacillaceae | Anoxybacillus_B | *Anoxybacillus_B calidus* | 575178 | *Anoxybacillus calidus* | https://pubmed.ncbi.nlm.nih.gov/24052627/ |
| GCA_000008665.1 | Hyperthermophiles | Alkaliphiles | 2178400 | Archaea | Halobacteriota | Archaeoglobi | Archaeoglobales | Archaeoglobaceae | Archaeoglobus | *Archaeoglobus fulgidus* | 2234 | *Archaeoglobus fulgidus* | https://doi.org/10.1007/s13205-019-1693-9 |
| GCA_002954225.1 | Psychrophiles | | 3655669 | Bacteria | Actinobacteriota | Actinomycetia | Actinomycetales | Micrococcaceae | Arthrobacter_D | *Arthrobacter_D ruber* | 1258893 | *Arthrobacter ruber* | https://doi.org/10.1099/ijsem.0.002719 |
| GCA_003097355.1 | Psychrophiles | | 4708612 | Bacteria | Actinobacteriota | Actinomycetia | Actinomycetales | Micrococcaceae | Arthrobacter_I | *Arthrobacter_I sp003097355* | 2703675 | *Arthrobacter sp. H-02-3* | https://doi.org/10.1186/s40168-021-01084-z |
| GCA_008369445.1 | | Alkaliphiles | 3936980 | Bacteria | Proteobacteria | Alphaproteobacteria | Rhizobiales | Rhizobiaceae | Aureimonas | *Aureimonas fodinaquatilis* | 2565783 | *Aureimonas fodinaquatilis* | https://doi.org/10.1007/s00203-020-01988-8 |
| GCA_000742895.1 | Mesophiles | | 5506189 | Bacteria | Firmicutes | Bacilli | Bacillales | Bacillaceae_G | Bacillus_A | *Bacillus_A anthracis* | 572264 | *Bacillus cereus 03BB102* | https://edis.ifas.ufl.edu/publication/FS269 |
| GCA_000615945.1 | | Alkaliphiles | 4534596 | Bacteria | Firmicutes | Bacilli | Caldalkalibacillales | JCM-10596 | Bacillus_AT | *Bacillus_AT mannanilyticus* | 1236954 | *Caldalkalibacillus mannanilyticus JCM 10596* | https://doi.org/10.1007/s00203-022-02789-x |
| GCA_004345675.1 | Thermophiles | | 2575920 | Bacteria | Firmicutes_A | Thermoanaerobacteria | Thermoanaerobacterales | Thermoanaerobacteraceae | Caldanaerobacter | *Caldanaerobacter subterraneus* | 911092 | *Caldanaerobacter subterraneus* | https://doi.org/10.1128%2FAEM.05190-11 |
| GCA_000421725.1 | Thermophiles | | 2589957 | Bacteria | Firmicutes_A | Thermoanaerobacteria | Caldicellulosiruptorales | Caldicellulosiruptoraceae | Caldicellulosiruptor | *Caldicellulosiruptor acetigenus* | 1121259 | *Caldicellulosiruptor acetigenus DSM 7040* | https://doi.org/10.1099/ijs.0.63723-0 |
| GCA_000022325.1 | Thermophiles | | 2931662 | Bacteria | Firmicutes_A | Thermoanaerobacteria | Caldicellulosiruptorales | Caldicellulosiruptoraceae | Caldicellulosiruptor | *Caldicellulosiruptor bescii* | 31899 | *Caldicellulosiruptor bescii* | https://doi.org/10.1099/ijs.0.017731-0 |
| GCA_000955725.1 | Thermophiles | | 2834482 | Bacteria | Firmicutes_A | Thermoanaerobacteria | Caldicellulosiruptorales | Caldicellulosiruptoraceae | Caldicellulosiruptor | *Caldicellulosiruptor danielii* | 1387557 | *Caldicellulosiruptor danielii* | https://doi.org/10.1128/aem.02694-17 |
| GCA_000166355.1 | Thermophiles | | 2770676 | Bacteria | Firmicutes_A | Thermoanaerobacteria | Caldicellulosiruptorales | Caldicellulosiruptoraceae | Caldicellulosiruptor | *Caldicellulosiruptor hydrothermalis* | 632292 | *Caldicellulosiruptor hydrothermalis 108* | https://www.microbiologyresearch.org/content/journal/ijsem/10.1099/ijs.0.65236-0 |
| GCA_000166775.1 | Thermophiles | | 2843785 | Bacteria | Firmicutes_A | Thermoanaerobacteria | Caldicellulosiruptorales | Caldicellulosiruptoraceae | Caldicellulosiruptor | *Caldicellulosiruptor kronotskyensis* | 632348 | *Caldicellulosiruptor kronotskyensis 2002* | https://doi.org/10.1099/ijs.0.65236-0 |
| GCA_000955745.1 | Thermophiles | | 2488483 | Bacteria | Firmicutes_A | Thermoanaerobacteria | Caldicellulosiruptorales | Caldicellulosiruptoraceae | Caldicellulosiruptor | *Caldicellulosiruptor morganii* | 1387555 | *Caldicellulosiruptor morganii* | https://doi.org/10.1128/genomeA.00440-15 |
| GCA_000955735.1 | Thermophiles | | 2514985 | Bacteria | Firmicutes_A | Thermoanaerobacteria | Caldicellulosiruptorales | Caldicellulosiruptoraceae | Caldicellulosiruptor | *Caldicellulosiruptor naganoensis* | 1387569 | *Caldicellulosiruptor naganoensis NA10* | https://doi.org/10.1128/mra.01292-22 |
| GCA_000145215.1 | Thermophiles | | 2532343 | Bacteria | Firmicutes_A | Thermoanaerobacteria | Caldicellulosiruptorales | Caldicellulosiruptoraceae | Caldicellulosiruptor | *Caldicellulosiruptor obsidiansis* | 608506 | *Caldicellulosiruptor obsidiansis OB47* | https://doi.org/10.1128/aem.01903-09 |
| GCA_000166335.1 | Thermophiles | | 2428903 | Bacteria | Firmicutes_A | Thermoanaerobacteria | Caldicellulosiruptorales | Caldicellulosiruptoraceae | Caldicellulosiruptor | *Caldicellulosiruptor owensensis* | 632518 | *Caldicellulosiruptor owensensis OL* | https://doi.org/10.1128/jb.01515-10 |
| GCA_000016545.1 | Thermophiles | | 2970275 | Bacteria | Firmicutes_A | Thermoanaerobacteria | Caldicellulosiruptorales | Caldicellulosiruptoraceae | Caldicellulosiruptor | *Caldicellulosiruptor saccharolyticus* | 351627 | *Caldicellulosiruptor saccharolyticus DSM 8903* | https://doi.org/10.1007/s00253-006-0783-x |
| GCA_000317795.1 | Thermophiles | Acidophiles | 1546846 | Archaea | Thermoproteota | Thermoproteia | Sulfolobales | Acidilobaceae | Caldisphaera | *Caldisphaera lagunensis* | 1056495 | *Caldisphaera lagunensis* | https://doi.org/10.1099/ijs.0.02580-0 |
| GCA_000018305.1 | Hyperthermophiles | Acidophiles | 2077567 | Archaea | Thermoproteota | Thermoproteia | Thermoproteales | Thermocladiaceae | Caldivirga | *Caldivirga maquilingensis* | 397948 | *Caldivirga maquilingensis IC-167* | https://doi.org/10.1099/00207713-49-3-1157 |
| GCA_000430045.1 | Thermophiles | | 4688964 | Bacteria | Deinococcota | Deinococci | Deinococcales | Thermaceae | Calidithermus | *Calidithermus chliarophilus* | 926560 | *Calidithermus chliarophilus DSM 9957* | https://www.microbiologyresearch.org/content/journal/ijsem/10.1099/ijsem.0.003270 |
| GCA_003574095.1 | Thermophiles | | 3675477 | Bacteria | Deinococcota | Deinococci | Deinococcales | Thermaceae | Calidithermus | *Calidithermus roseus* | 1644118 | *Calidithermus roseus* | https://www.microbiologyresearch.org/content/journal/ijsem/10.1099/ijsem.0.003270 |
| GCA_000373205.1 | Thermophiles | | 3190628 | Bacteria | Deinococcota | Deinococci | Deinococcales | Thermaceae | Calidithermus | *Calidithermus timidus* | 1122223 | *Calidithermus timidus DSM 17022* | https://www.microbiologyresearch.org/content/journal/ijsem/10.1099/ijsem.0.003270 |
| GCA_001950325.1 | Thermophiles | | 2386596 | Bacteria | Firmicutes_B | Z-2901 | Carboxydothermales | Carboxydothermaceae | Carboxydothermus | *Carboxydothermus islandicus* | 661089 | *Carboxydothermus islandicus* | https://doi.org/10.1099/ijs.0.030288-0 |
| GCA_001950255.1 | Thermophiles | | 2465639 | Bacteria | Firmicutes_B | Z-2901 | Carboxydothermales | Carboxydothermaceae | Carboxydothermus | *Carboxydothermus pertinax* | 870242 | *Carboxydothermus pertinax* | https://doi.org/10.1099/ijs.0.031583-0 |
| GCA_000744825.1 | | Alkaliphiles | 2632867 | Bacteria | Firmicutes | Bacilli | Lactobacillales | Carnobacteriaceae | Carnobacterium_A | *Carnobacterium_A mobile* | 1449342 | *Carnobacterium mobile DSM 4848* | https://hal.univ-lorraine.fr/tel-01749410/document |
| GCA_000195575.1 | Psychrophiles | Alkaliphiles | 2685399 | Bacteria | Firmicutes | Bacilli | Lactobacillales | Carnobacteriaceae | Carnobacterium_A | *Carnobacterium_A sp000195575* | 208596 | *Carnobacterium sp. 17-4* | https://link.springer.com/article/10.1007/s00792-011-0377-0 |
| GCA_000200715.1 | Psychrophiles | | 2045086 | Archaea | Thermoproteota | Nitrososphaeria | Nitrososphaerales | Nitrosopumilaceae | Cenarchaeum | *Cenarchaeum symbiosum* | 414004 | *Cenarchaeum symbiosum A* | https://doi.org/10.1073/pnas.93.13.6241 |
| GCA_000226295.1 | Thermophiles | Alkaliphiles | 3695372 | Bacteria | Acidobacteriota | Blastocatellia | Chloracidobacteriales | Chloracidobacteriaceae | Chloracidobacterium | *Chloracidobacterium thermophilum* | 458033 | *Chloracidobacterium thermophilum* | https://doi.org/10.1099/ijs.0.000113 |
| GCA_013340765.1 | Thermophiles | Acidophiles | 1593902 | Archaea | Thermoproteota | Nitrososphaeria | Conexivisphaerales | Conexivisphaeraceae | Conexivisphaera | *Conexivisphaera calida* | 1874277 | *Conexivisphaera calida* | https://doi.org/10.1099/ijsem.0.004595 |
| GCA_000819445.1 | | Alkaliphiles | 2681312 | Bacteria | Actinobacteriota | Actinomycetia | Mycobacteriales | Mycobacteriaceae | Corynebacterium | *Corynebacterium humireducens* | 1223515 | *Corynebacterium humireducens NBRC 106098 = DSM 45392* | https://doi.org/10.1099/ijs.0.020909-0 |
| GCA_001277255.1 | Mesophiles | | 4499482 | Bacteria | Proteobacteria | Gammaproteobacteria | Enterobacterales | Enterobacteriaceae | Cronobacter | *Cronobacter condimenti* | 1073999 | *Cronobacter condimenti 1330* | https://www.microbiologyresearch.org/content/journal/ijsem/10.1099/ijs.0.032292-0 |
| GCA_001277235.1 | Mesophiles | | 4628405 | Bacteria | Proteobacteria | Gammaproteobacteria | Enterobacterales | Enterobacteriaceae | Cronobacter | *Cronobacter dublinensis* | 1208656 | *Cronobacter dublinensis 1210* | https://www.microbiologyresearch.org/content/journal/ijsem/10.1099/ijs.0.65577-0 |
| GCA_001277215.2 | Mesophiles | | 4473761 | Bacteria | Proteobacteria | Gammaproteobacteria | Enterobacterales | Enterobacteriaceae | Cronobacter | *Cronobacter malonaticus* | 413503 | *Cronobacter malonaticus* | https://www.microbiologyresearch.org/content/journal/ijsem/10.1099/ijs.0.65577-0 |
| GCA_001277195.1 | Mesophiles | | 4364114 | Bacteria | Proteobacteria | Gammaproteobacteria | Enterobacterales | Enterobacteriaceae | Cronobacter | *Cronobacter muytjensii* | 1159613 | *Cronobacter muytjensii ATCC 51329* | https://www.microbiologyresearch.org/content/journal/ijsem/10.1099/ijs.0.65577-0 |
| GCA_000982825.1 | Mesophiles | | 4663565 | Bacteria | Proteobacteria | Gammaproteobacteria | Enterobacterales | Enterobacteriaceae | Cronobacter | *Cronobacter sakazakii* | 1208592 | *Cronobacter sakazakii 680* | https://www.microbiologyresearch.org/content/journal/ijsem/10.1099/ijs.0.65577-0 |
| GCA_000027065.2 | Mesophiles | | 4599092 | Bacteria | Proteobacteria | Gammaproteobacteria | Enterobacterales | Enterobacteriaceae | Cronobacter | *Cronobacter turicensis* | 1208590 | *Cronobacter turicensis 564* | https://www.microbiologyresearch.org/content/journal/ijsem/10.1099/ijs.0.65577-0 |

| | | | | | | | | | | | | | |
|---|---|---|---|---|---|---|---|---|---|---|---|---|---|
| GCA_001277175.1 | Mesophiles | | 4436873 | Bacteria | Proteobacteria | Gammaproteobacteria | Enterobacterales | Enterobacteriaceae | Cronobacter | *Cronobacter universalis* | 1074000 | *Cronobacter universalis NCTC 9529* | https://www.microbiologyresearch.org/content/journal/ijsem/10.1099/ijs.0.65577-0 |
| GCA_003185895.1 | Psychrophiles | | 4305516 | Bacteria | Actinobacteriota | Actinomycetia | Actinomycetales | Microbacteriaceae | Cryobacterium | *Cryobacterium arcticum* | 670052 | *Cryobacterium arcticum* | https://doi.org/10.1099/ijs.0.027128-0 |
| GCA_001679725.1 | Psychrophiles | | 4351229 | Bacteria | Actinobacteriota | Actinomycetia | Actinomycetales | Microbacteriaceae | Cryobacterium | *Cryobacterium arcticum_A* | 670052 | *Cryobacterium arcticum* | https://doi.org/10.1099/ijs.0.027128-0 |
| GCA_002954245.1 | Psychrophiles | | 4314199 | Bacteria | Actinobacteriota | Actinomycetia | Actinomycetales | Microbacteriaceae | Cryobacterium | *Cryobacterium aureum* | 995037 | *Cryobacterium aureum* | https://www.microbiologyresearch.org/content/journal/ijsem/10.1099/ijsem.0.002647 |
| GCA_900103805.1 | Psychrophiles | | 4040838 | Bacteria | Actinobacteriota | Actinomycetia | Actinomycetales | Microbacteriaceae | Cryobacterium | *Cryobacterium flavum* | 1424659 | *Cryobacterium flavum* | https://doi.org/10.1099/ijs.0.033738-0 |
| GCA_004402405.1 | Psychrophiles | | 3753316 | Bacteria | Actinobacteriota | Actinomycetia | Actinomycetales | Microbacteriaceae | Cryobacterium | *Cryobacterium levicorallinum* | 995038 | *Cryobacterium levicorallinum* | https://doi.org/10.1099/ijs.0.046896-0 |
| GCA_900110125.1 | Psychrophiles | | 3834069 | Bacteria | Actinobacteriota | Actinomycetia | Actinomycetales | Microbacteriaceae | Cryobacterium | *Cryobacterium luteum* | 1424661 | *Cryobacterium luteum* | https://doi.org/10.1099/ijs.0.033738-0 |
| GCA_004365915.1 | Psychrophiles | | 3677616 | Bacteria | Actinobacteriota | Actinomycetia | Actinomycetales | Microbacteriaceae | Cryobacterium | *Cryobacterium psychrophilum* | 41988 | *Cryobacterium psychrophilum* | https://doi.org/10.1099/00207713-47-2-474 |
| GCA_900101115.1 | Psychrophiles | | 3247111 | Bacteria | Actinobacteriota | Actinomycetia | Actinomycetales | Microbacteriaceae | Cryobacterium | *Cryobacterium psychrotolerans* | 386301 | *Cryobacterium psychrotolerans* | https://doi.org/10.1099/ijs.0.64750-0 |
| GCA_014200405.1 | Psychrophiles | | 4531652 | Bacteria | Actinobacteriota | Actinomycetia | Actinomycetales | Microbacteriaceae | Cryobacterium | *Cryobacterium roopkundense* | 1001240 | *Cryobacterium roopkundense* | https://doi.org/10.1099/ijs.0.011775-0 |
| GCA_002909375.1 | Psychrophiles | | 4048390 | Bacteria | Actinobacteriota | Actinomycetia | Actinomycetales | Microbacteriaceae | Cryobacterium | *Cryobacterium zongtaii* | 1259217 | *Cryobacterium zongtaii* | https://doi.org/10.1016/j.syapm.2018.10.005 |
| GCA_000711975.1 | | Alkaliphiles | 2622239 | Bacteria | Firmicutes_B | Desulfotomaculia | Desulfotomaculales | Desulfohalotomaculaceae | Desulfofalx | *Desulfofalx alkaliphila* | 1121423 | *Desulfofalx alkaliphila DSM 12257* | 10.1002/9781118960608.gbm01777 |
| GCA_000686525.1 | | Alkaliphiles | 2939696 | Bacteria | Desulfobacterota_I | Desulfovibrionia | Desulfovibrionales | Desulfonatronovibrionaceae | Desulfonatronovibrio | *Desulfonatronovibrio hydrogenovorans* | 1121413 | *Desulfonatronovibrio hydrogenovorans DSM 9292* | https://doi.org/10.1099/00207713-47-1-144 |
| GCA_000934755.1 | | Alkaliphiles | 4809662 | Bacteria | Desulfobacterota_I | Desulfovibrionia | Desulfovibrionales | Desulfonatronovibrionaceae | Desulfonatronovibrio | *Desulfonatronovibrio magnus* | 698827 | *Desulfonatronovibrio magnus* | https://doi.org/10.1007/s00792-011-0370-7 |
| GCA_000934745.1 | | Alkaliphiles | 4634603 | Bacteria | Desulfobacterota_I | Desulfovibrionia | Desulfovibrionales | Desulfonatronaceae | Desulfonatronum | *Desulfonatronum thioautotrophicum* | 617001 | *Desulfonatronum thioautotrophicum* | https://doi.org/10.1007/s00792-011-0370-7 |
| GCA_900104215.1 | | Alkaliphiles | 3602125 | Bacteria | Desulfobacterota_I | Desulfovibrionia | Desulfovibrionales | Desulfonatronaceae | Desulfonatronum | *Desulfonatronum thiosulfatophilum* | 617002 | *Desulfonatronum thiosulfatophilum* | GCA_900104215.1 |
| GCA_001029285.1 | | Acidophiles | 4637866 | Bacteria | Firmicutes_B | Desulfitobacteriia | Desulfitobacteriales | Desulfitobacteriaceae | Desulfosporosinus | *Desulfosporosinus acididurans* | 476652 | *Desulfosporosinus acididurans* | https://doi.org/10.1007/s00792-014-0701-6 |
| GCA_000255115.3 | | Acidophiles | 4991181 | Bacteria | Firmicutes_B | Desulfitobacteriia | Desulfitobacteriales | Desulfitobacteriaceae | Desulfosporosinus | *Desulfosporosinus acidiphilus* | 646529 | *Desulfosporosinus acidiphilus SJ4* | https://doi.org/10.1007/s00792-014-0701-6 |
| GCA_001707885.1 | | Acidophiles | 4523251 | Bacteria | Firmicutes_B | Desulfitobacteriia | Desulfitobacteriales | Desulfitobacteriaceae | Desulfosporosinus | *Desulfosporosinus sp001707885* | 1633135 | *Desulfosporosinus sp. BG* | https://doi.org/10.1016/j.gdata.2016.12.014 |
| GCA_000429345.1 | Thermophiles | | 3060447 | Bacteria | Firmicutes_B | Desulfotomaculia | Desulfotomaculales | Desulfovirgulaceae | Desulfovirgula | *Desulfovirgula thermocuniculi* | 1121468 | *Desulfovirgula thermocuniculi DSM 16036* | https://doi.org/10.1099/ijs.0.64655-0 |
| GCA_000513855.1 | Hyperthermophiles | | 1307099 | Archaea | Thermoproteota | Thermoproteia | Sulfolobales | Desulfurococcaceae | Desulfurococcus | *Desulfurococcus amylolyticus* | 490899 | *Desulfurococcus amylolyticus 1221n* | https://doi.org/10.1099/ijsem.0.000747 |
| GCA_000186365.1 | Hyperthermophiles | | 1314639 | Archaea | Thermoproteota | Thermoproteia | Sulfolobales | Desulfurococcaceae | Desulfurococcus | *Desulfurococcus mucosus* | 765177 | *Desulfurococcus mucosus 07/1, DSM 2162* | https://doi.org/10.1099/ijsem.0.000747 |
| GCA_000020965.1 | Thermophiles | | 1959987 | Bacteria | Dictyoglomota | Dictyoglomia | Dictyoglomales | Dictyoglomaceae | Dictyoglomus | *Dictyoglomus thermophilum* | 309799 | *Dictyoglomus thermophilum H-6-12* | https://doi.org/10.1099/00207713-35-3-253 |
| GCA_000021645.1 | Thermophiles | | 1855560 | Bacteria | Dictyoglomota | Dictyoglomia | Dictyoglomales | Dictyoglomaceae | Dictyoglomus | *Dictyoglomus turgidum* | 515635 | *Dictyoglomus turgidum DSM 6724* | https://doi.org/10.3389/fmicb.2016.01979 |
| GCA_000376225.1 | Thermophiles | | 3363524 | Bacteria | Firmicutes | Bacilli | Tumebacillales | Effusibacillaceae | Effusibacillus | *Effusibacillus pohliae* | 1120973 | *Effusibacillus pohliae DSM 22757* | https://doi.org/10.1099/ijs.0.055814-0 |
| GCA_000378465.1 | Thermophiles | | 4304237 | Bacteria | Proteobacteria | Alphaproteobacteria | Acetobacterales | Acetobacteraceae | Elioraea | *Elioraea tepidiphila* | 1121861 | *Elioraea tepidiphila DSM 17972* | https://doi.org/10.1099/ijs.0.65294-0 |
| GCA_001544255.1 | Mesophiles | | 2484851 | Bacteria | Firmicutes | Bacilli | Lactobacillales | Enterococcaceae | Enterococcus_B | *Enterococcus_B faecium* | 1442605 | *Enterococcus faecium VSE1036* | https://doi.org/10.1016/j.resmic.2005.11.006 |
| GCA_000759775.1 | Mesophiles | | 4422416 | Bacteria | Proteobacteria | Gammaproteobacteria | Enterobacterales | Enterobacteriaceae | Escherichia | *Escherichia albertii* | 910238 | *Escherichia albertii TW15818* | https://www.sciencedirect.com/science/article/pii/S0963996913000896 |
| GCA_000026225.1 | Mesophiles | | 4643861 | Bacteria | Proteobacteria | Gammaproteobacteria | Enterobacterales | Enterobacteriaceae | Escherichia | *Escherichia fergusonii* | 564 | *Escherichia fergusonii* | https://www.sciencedirect.com/science/article/pii/S0378113514002168?via%3Dihub |
| GCA_002900365.1 | Mesophiles | | 4896291 | Bacteria | Proteobacteria | Gammaproteobacteria | Enterobacterales | Enterobacteriaceae | Escherichia | *Escherichia marmotae* | 1499973 | *Escherichia marmotae* | https://www.microbiologyresearch.org/content/journal/ijsem/10.1099/ijs.0.000228 |
| GCA_000745905.1 | | Acidophiles | 2928893 | Bacteria | Actinobacteriota | Acidimicrobiia | Acidimicrobiales | Acidimicrobiaceae | Ferrimicrobium | *Ferrimicrobium acidiphilum* | 1121877 | *Ferrimicrobium acidiphilum DSM 19497* | https://doi.org/10.1099/ijs.0.65409-0 |
| GCA_900128965.1 | | Acidophiles | 2489535 | Bacteria | Actinobacteriota | Acidimicrobiia | Acidimicrobiales | Acidimicrobiaceae | Ferrithrix | *Ferrithrix thermotolerans* | 1121881 | *Ferrithrix thermotolerans DSM 19514* | https://doi.org/10.1099/ijs.0.65409-0 |
| GCA_000025505.1 | Hyperthermophiles | | 2196266 | Archaea | Halobacteriota | Archaeoglobi | Archaeoglobales | Archaeoglobaceae | Ferroglobus | *Ferroglobus placidus* | 589924 | *Ferroglobus placidus DSM 10642* | https://doi.org/10.1007/s002030050388 |
| GCA_000152265.2 | | Acidophiles | 1935211 | Archaea | Thermoplasmatota | Thermoplasmata | Thermoplasmatales | Thermoplasmataceae | Ferroplasma | *Ferroplasma acidarmanus* | 333146 | *Ferroplasma acidarmanus fer1* | https://www.microbiologyresearch.org/content/journal/micro/10.1099/mic.0.28076-0 |
| GCA_002078355.1 | | Acidophiles | 1826943 | Archaea | Thermoplasmatota | Thermoplasmata | Thermoplasmatales | Thermoplasmataceae | Ferroplasma | *Ferroplasma acidiphilum* | 74969 | *Ferroplasma acidiphilum* | https://doi.org/10.1099/00207713-50-3-997 |
| GCA_004117075.1 | Thermophiles | | 2266449 | Bacteria | Thermotogota | Thermotogae | Thermotogales | Fervidobacteriaceae | Fervidobacterium | *Fervidobacterium changbaicum* | 310769 | *Fervidobacterium changbaicum* | https://doi.org/10.1099/ijs.0.64758-0 |
| GCA_000235405.3 | Thermophiles | | 2166381 | Bacteria | Thermotogota | Thermotogae | Thermotogales | Fervidobacteriaceae | Fervidobacterium | *Fervidobacterium pennivorans* | 93466 | *Fervidobacterium pennivorans* | https://doi.org/10.3390/microorganisms11010022 |
| GCA_001644665.1 | Thermophiles | | 2061852 | Bacteria | Thermotogota | Thermotogae | Thermotogales | Fervidobacteriaceae | Fervidobacterium | *Fervidobacterium pennivorans_A* | 93466 | *Fervidobacterium pennivorans* | https://doi.org/10.3390/microorganisms11010022 |
| GCA_001719065.1 | Thermophiles | | 2040210 | Bacteria | Thermotogota | Thermotogae | Thermotogales | Fervidobacteriaceae | Fervidobacterium_A | *Fervidobacterium_A thailandense* | 1008305 | *Fervidobacterium thailandense* | https://doi.org/10.1099/ijsem.0.001463 |
| GCA_000419685.1 | Psychrophiles | | 3079036 | Bacteria | Bacteroidota | Bacteroidia | Flavobacteriales | Flavobacteriaceae | Flavobacterium | *Flavobacterium antarcticum* | 1111730 | *Flavobacterium antarcticum DSM 19726* | https://doi.org/10.1099/ijs.0.63423-0 |
| GCA_003259835.1 | Psychrophiles | Alkaliphiles | 2821499 | Bacteria | Bacteroidota | Bacteroidia | Flavobacteriales | Flavobacteriaceae | Flavobacterium | *Flavobacterium aquaticum* | 1236486 | *Flavobacterium aquaticum* | https://link.springer.com/article/10.1007/s12275-013-2293-8 |
| GCA_015223105.1 | Psychrophiles | | 2830228 | Bacteria | Bacteroidota | Bacteroidia | Flavobacteriales | Flavobacteriaceae | Flavobacterium | *Flavobacterium aquaticum_A* | 1236486 | *Flavobacterium proteolyticum* | https://doi.org/10.1007/s00203-021-02744-2 |
| GCA_003344925.1 | Psychrophiles | | 2970356 | Bacteria | Bacteroidota | Bacteroidia | Flavobacteriales | Flavobacteriaceae | Flavobacterium | *Flavobacterium arcticum* | 1784713 | *Flavobacterium arcticum* | https://doi.org/10.1099/ijsem.0.001804 |
| GCA_900106645.1 | Psychrophiles | | 3856409 | Bacteria | Bacteroidota | Bacteroidia | Flavobacteriales | Flavobacteriaceae | Flavobacterium | *Flavobacterium degerlachei* | 229203 | *Flavobacterium degerlachei* | https://doi.org/10.1099/ijs.0.02857-0 |
| GCA_000425505.1 | Psychrophiles | | 3631041 | Bacteria | Bacteroidota | Bacteroidia | Flavobacteriales | Flavobacteriaceae | Flavobacterium | *Flavobacterium frigidarium* | 1121890 | *Flavobacterium frigidarium DSM 17623* | https://doi.org/10.1099/00207713-51-4-1235 |
| GCA_900111075.1 | Psychrophiles | | 4045707 | Bacteria | Bacteroidota | Bacteroidia | Flavobacteriales | Flavobacteriaceae | Flavobacterium | *Flavobacterium frigoris* | 229204 | *Flavobacterium frigoris* | https://doi.org/10.1099/ijs.0.02857-0 |
| GCA_000252125.2 | Psychrophiles | | 3934101 | Bacteria | Bacteroidota | Bacteroidia | Flavobacteriales | Flavobacteriaceae | Flavobacterium | *Flavobacterium frigoris_A* | 229204 | *Flavobacterium frigoris* | https://doi.org/10.1099/ijs.0.02857-0 |
| GCA_900143245.1 | Psychrophiles | | 3709863 | Bacteria | Bacteroidota | Bacteroidia | Flavobacteriales | Flavobacteriaceae | Flavobacterium | *Flavobacterium fryxellicola* | 249352 | *Flavobacterium fryxellicola* | https://doi.org/10.1099/ijs.0.03056-0 |
| GCA_900107635.1 | Psychrophiles | | 4377959 | Bacteria | Bacteroidota | Bacteroidia | Flavobacteriales | Flavobacteriaceae | Flavobacterium | *Flavobacterium gillisiae* | 150146 | *Flavobacterium gillisiae* | https://doi.org/10.1099/00207713-50-3-1055 |
| GCA_003350545.1 | Psychrophiles | | 3143612 | Bacteria | Bacteroidota | Bacteroidia | Flavobacteriales | Flavobacteriaceae | Flavobacterium | *Flavobacterium glaciei* | 386300 | *Flavobacterium glaciei* | https://doi.org/10.1099/ijs.0.64564-0 |
| GCA_003634755.1 | Psychrophiles | | 3305187 | Bacteria | Bacteroidota | Bacteroidia | Flavobacteriales | Flavobacteriaceae | Flavobacterium | *Flavobacterium limicola* | 180441 | *Flavobacterium limicola* | https://doi.org/10.1099/ijs.0.02369-0 |
| GCA_900129585.1 | Psychrophiles | | 3692790 | Bacteria | Bacteroidota | Bacteroidia | Flavobacteriales | Flavobacteriaceae | Flavobacterium | *Flavobacterium micromati* | 229205 | *Flavobacterium micromati* | https://doi.org/10.1099/ijs.0.02857-0 |
| GCA_900099915.1 | Psychrophiles | | 3808360 | Bacteria | Bacteroidota | Bacteroidia | Flavobacteriales | Flavobacteriaceae | Flavobacterium | *Flavobacterium omnivorum* | 178355 | *Flavobacterium omnivorum* | https://doi.org/10.1099/ijs.0.02310-0 |
| GCA_003312425.1 | Psychrophiles | | 3652041 | Bacteria | Bacteroidota | Bacteroidia | Flavobacteriales | Flavobacteriaceae | Flavobacterium | *Flavobacterium psychrolimnae* | 249351 | *Flavobacterium psychrolimnae* | https://doi.org/10.1099/ijs.0.03056-0 |
| GCA_002217405.1 | Psychrophiles | | 2638051 | Bacteria | Bacteroidota | Bacteroidia | Flavobacteriales | Flavobacteriaceae | Flavobacterium | *Flavobacterium psychrophilum* | 96345 | *Flavobacterium psychrophilum* | https://www.nature.com/articles/nbt1313 |
| GCA_001708385.1 | Psychrophiles | | 4142802 | Bacteria | Bacteroidota | Bacteroidia | Flavobacteriales | Flavobacteriaceae | Flavobacterium | *Flavobacterium psychrophilum_A* | 96345 | *Flavobacterium psychrophilum* | https://www.nature.com/articles/nbt1313 |
| GCA_900129575.1 | Psychrophiles | | 3463995 | Bacteria | Bacteroidota | Bacteroidia | Flavobacteriales | Flavobacteriaceae | Flavobacterium | *Flavobacterium segetis* | 271157 | *Flavobacterium degerlachei* | https://doi.org/10.1099/ijs.0.02857-0 |
| GCA_900110375.1 | Psychrophiles | Alkaliphiles | 3520953 | Bacteria | Bacteroidota | Bacteroidia | Flavobacteriales | Flavobacteriaceae | Flavobacterium | *Flavobacterium sinopsychrotolerans* | 604089 | *Flavobacterium sinopsychrotolerans* | https://doi.org/10.1099/ijs.0.014126-0 |
| GCA_000425485.1 | Psychrophiles | | 3774544 | Bacteria | Bacteroidota | Bacteroidia | Flavobacteriales | Flavobacteriaceae | Flavobacterium | *Flavobacterium tegetincola* | 1121901 | *Flavobacterium tegetincola DSM 22377* | https://www.microbiologyresearch.org/content/journal/ijsem/10.1099/00207713-50-3-1055 |
| GCA_900108015.1 | Psychrophiles | | 3439151 | Bacteria | Bacteroidota | Bacteroidia | Flavobacteriales | Flavobacteriaceae | Flavobacterium | *Flavobacterium urumqiense* | 935224 | *Flavobacterium urumqiense* | https://doi.org/10.1099/ijs.0.030049-0 |
| GCA_003688495.1 | Psychrophiles | | 3397857 | Bacteria | Bacteroidota | Bacteroidia | Flavobacteriales | Flavobacteriaceae | Flavobacterium | *Flavobacterium weaverense* | 271156 | *Flavobacterium weaverense* | https://doi.org/10.1099/ijs.0.64164-0 |

| | | | | | | | | | | | | | |
|---|---|---|---|---|---|---|---|---|---|---|---|---|---|
| GCA_900142695.1 | Psychrophiles | | 3755582 | Bacteria | Bacteroidota | Bacteroidia | Flavobacteriales | Flavobacteriaceae | Flavobacterium | *Flavobacterium xanthum* | 69322 | *Flavobacterium xanthum* | https://www.microbiologyresearch.org/content/journal/ijsem/10.1099/00207713-50-3-1055 |
| GCA_900142885.1 | Psychrophiles | | 3902806 | Bacteria | Bacteroidota | Bacteroidia | Flavobacteriales | Flavobacteriaceae | Flavobacterium | *Flavobacterium xinjiangense* | 178356 | *Flavobacterium xinjiangense* | https://doi.org/10.1099/ijs.0.02310-0 |
| GCA_900112975.1 | Psychrophiles | | 3471047 | Bacteria | Bacteroidota | Bacteroidia | Flavobacteriales | Flavobacteriaceae | Flavobacterium | *Flavobacterium xueshanense* | 935223 | *Flavobacterium xueshanense* | https://doi.org/10.1099/ijs.0.030049-0 |
| GCA_000820505.1 | | Alkaliphiles | 3839692 | Bacteria | Desulfobacterota | Desulfuromonadia | Desulfuromonadales | Geoalkalibacteraceae | Geoalkalibacter | *Geoalkalibacter ferrihydriticus* | 1121915 | *Geoalkalibacter ferrihydriticus DSM 17813* | https://pubmed.ncbi.nlm.nih.gov/17205802/ |
| GCA_000789255.1 | Hyperthermophiles | | 1860815 | Archaea | Halobacteriota | Archaeoglobi | Archaeoglobales | Archaeoglobaceae | Geoglobus | *Geoglobus acetivorans* | 565033 | *Geoglobus acetivorans* | https://doi.org/10.1128/aem.02705-14 |
| GCA_015163485.1 | Hyperthermophiles | | 1901114 | Archaea | Halobacteriota | Archaeoglobi | Archaeoglobales | Archaeoglobaceae | Geoglobus | *Geoglobus acetivorans_A* | 565033 | *Geoglobus acetivorans* | https://doi.org/10.1128/aem.02705-14 |
| GCA_001006045.1 | Hyperthermophiles | | 1770093 | Archaea | Halobacteriota | Archaeoglobi | Archaeoglobales | Archaeoglobaceae | Geoglobus | *Geoglobus ahangari* | 113653 | *Geoglobus ahangari* | https://doi.org/10.1099/00207713-52-3-719 |
| GCA_002952775.1 | | Alkaliphiles | 3313120 | Archaea | Halobacteriota | Halobacteria | Halobacteriales | Haloferacaceae | Halalkaliarchaeum | *Halalkaliarchaeum desulfuricum* | 2055893 | *Halalkaliarchaeum desulfuricum* | https://doi.org/10.1099/ijsem.0.003506 |
| GCA_000423105.1 | | Alkaliphiles | 2707549 | Bacteria | Firmicutes | Bacilli | Bacillales_D | Alkalibacillaceae | Halalkalibacillus | *Halalkalibacillus halophilus* | 1121936 | *Halalkalibacillus halophilus DSM 18494* | https://doi.org/10.1099/ijs.0.64830-0 |
| GCA_000196895.1 | Thermophiles | Alkaliphiles | 3698650 | Archaea | Halobacteriota | Halobacteria | Halobacteriales | Halalkalicoccaceae | Halalkalicoccus | *Halalkalicoccus jeotgali* | 795797 | *Halalkalicoccus jeotgali* | https://pubmed.ncbi.nlm.nih.gov/17911300/ |
| GCA_004799665.1 | Mesophiles | | 3452056 | Archaea | Halobacteriota | Halobacteria | Halobacteriales | Haloarculaceae | Halapricum | *Halapricum salinum* | 1457250 | *Halapricum salinum strain CBA1105* | https://pubmed.ncbi.nlm.nih.gov/24677144/ |
| GCA_014647455.2 | | Acidophiles | 3024665 | Archaea | Halobacteriota | Halobacteria | Halobacteriales | Halobacteriaceae | Halarchaeum | *Halarchaeum grantii* | 1193105 | *Halarchaeum grantii* | https://doi.org/10.1099/ijsem.0.000501 |
| GCA_014647155.1 | | Acidophiles | 3141185 | Archaea | Halobacteriota | Halobacteria | Halobacteriales | Halobacteriaceae | Halarchaeum | *Halarchaeum nitratireducens* | 489913 | *Halarchaeum nitratireducens* | https://doi.org/10.1099/ijs.0.054668-0 |
| GCA_014647115.1 | Mesophiles | Acidophiles | 2813544 | Archaea | Halobacteriota | Halobacteria | Halobacteriales | Halobacteriaceae | Halarchaeum | *Halarchaeum rubridurum* | 489911 | *Halarchaeum rubridurum* | https://doi.org/10.1099/ijs.0.049262-0 |
| GCA_000336615.1 | Thermophiles | Alkaliphiles | 4225424 | Archaea | Halobacteriota | Halobacteria | Halobacteriales | Haloarculaceae | Haloarcula | *Haloarcula amylolytica* | 1227452 | *Haloarcula amylolytica JCM 13557* | https://www.microbiologyresearch.org/content/journal/ijsem/10.1099/ijs.0.64647-0 |
| GCA_000336895.1 | Mesophiles | | 4147107 | Archaea | Halobacteriota | Halobacteria | Halobacteriales | Haloarculaceae | Haloarcula | *Haloarcula argentinensis* | 1230451 | *Haloarcula argentinensis* | https://www.microbiologyresearch.org/content/journal/ijsem/10.1099/00207713-47-1-73 |
| GCA_010119195.1 | Mesophiles | | 3788104 | Archaea | Halobacteriota | Halobacteria | Halobacteriales | Haloarculaceae | Haloarcula | *Haloarcula salina* | 1429914 | *Halomicroarcula salina* | https://pubmed.ncbi.nlm.nih.gov/25721722/ |
| GCA_000337775.1 | Thermophiles | | 3923205 | Archaea | Halobacteriota | Halobacteria | Halobacteriales | Haloarculaceae | Haloarcula | *Haloarcula vallismortis* | 662477 | *Haloarcula vallismortis* | https://www.ncbi.nlm.nih.gov/pmc/articles/PMC545725/ |
| GCA_008124605.1 | Mesophiles | | 2364912 | Archaea | Halobacteriota | Halobacteria | Halobacteriales | Halobacteriaceae | Halobacterium | *Halobacterium salinarum* | 2597657 | *Halobacterium salinarum* | https://doi.org/10.1089%2Fomi.2012.0117 |
| GCA_000336935.1 | Mesophiles | Alkaliphiles | 4199784 | Archaea | Halobacteriota | Halobacteria | Halobacteriales | Halococcaceae | Halococcus | *Halococcus salifodinae* | 1227456 | *Halococcus salifodinae* | https://doi.org/10.1016/j.ijbiomac.2021.02.081 |
| GCA_000337795.1 | Thermophiles | | 3825973 | Archaea | Halobacteriota | Halobacteria | Halobacteriales | Haloferacaceae | Haloferax | *Haloferax denitrificans* | 662478 | *Haloferax denitrificans* | https://www.microbiologyresearch.org/content/journal/ijsem/10.1099/00207713-39-3-359?crawler=true |
| GCA_000336755.1 | Thermophiles | Acidophiles | 3952136 | Archaea | Halobacteriota | Halobacteria | Halobacteriales | Haloferacaceae | Haloferax | *Haloferax elongans* | 1230453 | *Haloferax elongans* | https://www.microbiologyresearch.org/content/journal/ijsem/10.1099/ijs.0.65360-0 |
| GCA_000306765.2 | Thermophiles | | 3904707 | Archaea | Halobacteriota | Halobacteria | Halobacteriales | Haloferacaceae | Haloferax | *Haloferax mediterranei* | 523841 | *Haloferax mediterranei* | https://www.ncbi.nlm.nih.gov/pmc/articles/PMC7915512/ |
| GCA_000337815.1 | Thermophiles | | 3368982 | Archaea | Halobacteriota | Halobacteria | Halobacteriales | Haloferacaceae | Haloferax | *Haloferax mucosum* | 662479 | *Haloferax mucosum* | https://www.microbiologyresearch.org/content/journal/ijsem/10.1099/ijs.0.65360-0 |
| GCA_000025685.1 | Mesophiles | | 4012900 | Archaea | Halobacteriota | Halobacteria | Halobacteriales | Haloferacaceae | Haloferax | *Haloferax volcanii* | 309800 | *Haloferax volcanii* | https://www.ncbi.nlm.nih.gov/pmc/articles/PMC545725/ |
| GCA_000172995.2 | Thermophiles | | 3944467 | Archaea | Halobacteriota | Halobacteria | Halobacteriales | Haloferacaceae | Halogeometricum | *Halogeometricum borinquense* | 469382 | *Halogeometricum borinquense* | https://www.ncbi.nlm.nih.gov/pmc/articles/PMC545725/ |
| GCA_900112175.1 | Thermophiles | | 4187125 | Archaea | Halobacteriota | Halobacteria | Halobacteriales | Haloferacaceae | Halogeometricum | *Halogeometricum rufum* | 553469 | *Halogeometricum rufum* | https://www.microbiologyresearch.org/content/journal/ijsem/10.1099/ijs.0.019463-0 |
| GCA_900110465.1 | Mesophiles | | 5185690 | Archaea | Halobacteriota | Halobacteria | Halobacteriales | Haloferacaceae | Halogranum | *Halogranum amylolyticum* | 660520 | *Halogranum amylolyticum* | https://doi.org/10.1099/ijs.0.024976-0 |
| GCA_900103715.1 | Mesophiles | | 3770187 | Archaea | Halobacteriota | Halobacteria | Halobacteriales | Haloferacaceae | Halogranum | *Halogranum gelatinilyticum* | 660521 | *Halogranum gelatinilyticum* | https://doi.org/10.1099/ijs.0.024976-0 |
| GCA_009791395.1 | Mesophiles | | 4069707 | Archaea | Halobacteriota | Halobacteria | Halobacteriales | Haloarculaceae | Halomarina | *Halomarina oriensis* | 671145 | *Halomarina oriensis* | https://doi.org/10.1099/ijs.0.020677-0 |
| GCA_003862495.1 | Mesophiles | | 3654689 | Archaea | Halobacteriota | Halobacteria | Halobacteriales | Haloarculaceae | Halomarina | *Halomarina oriensis_A* | 671145 | *Halocatena pleomorpha* | https://doi.org/10.1099/ijsem.0.004222 |
| GCA_010119205.1 | Mesophiles | | 3906684 | Archaea | Halobacteriota | Halobacteria | Halobacteriales | Haloarculaceae | Halomicroarcula | *Halomicroarcula limicola* | 1429915 | *Halomicroarcula limicola* | https://pubmed.ncbi.nlm.nih.gov/24554639/ |
| GCA_000379085.1 | Mesophiles | | 3607771 | Archaea | Halobacteriota | Halobacteria | Halobacteriales | Haloarculaceae | Halomicrobium | *Halomicrobium katesii* | 1069082 | *Halomicrobium katesii DSM 19301* | https://doi.org/10.1099/ijs.0.65662-0 |
| GCA_000023965.1 | Mesophiles | | 3332349 | Archaea | Halobacteriota | Halobacteria | Halobacteriales | Haloarculaceae | Halomicrobium | *Halomicrobium mukohataei* | 485914 | *Halomicrobium mukohataei DSM 12286* | https://www.microbiologyresearch.org/content/journal/ijsem/10.1099/00207713-52-5-1831 |
| GCA_900114435.1 | Mesophiles | | 4250330 | Archaea | Halobacteriota | Halobacteria | Halobacteriales | Haloarculaceae | Halomicrobium | *Halomicrobium zhouii* | 767519 | *Halomicrobium zhouii* | https://doi.org/10.1099/ijs.0.031989-0 |
| GCA_002966495.1 | | Alkaliphiles | 3650492 | Bacteria | Proteobacteria | Gammaproteobacteria | Pseudomonadales | Halomonadaceae | Halomonas | *Halomonas sp002966495* | 1118153 | *Halomonas sp. GFAJ-1* | https://www.ncbi.nlm.nih.gov/pmc/articles/PMC6598117/ |
| GCA_900100875.1 | Mesophiles | | 3871751 | Archaea | Halobacteriota | Halobacteria | Halobacteriales | Haloferacaceae | Halopelagius | *Halopelagius longus* | 1236180 | *Halopelagius longus* | https://doi.org/10.1099/ijs.0.051375-0 |
| GCA_000455345.1 | Thermophiles | Alkaliphiles | 3906364 | Archaea | Halobacteriota | Halobacteria | Halobacteriales | Natrialbaceae | Halopiger_A | *Halopiger_A goleimassiliensis* | 1293048 | *Halopiger goleimassiliensis* | http://standardsingenomics.org/content/9/3/956/ |
| GCA_000470655.1 | Mesophiles | | 3146160 | Archaea | Halobacteriota | Halobacteria | Halobacteriales | Haloarculaceae | Halorhabdus | *Halorhabdus tiamatea* | 1033806 | *Halorhabdus tiamatea SARL4B* | https://doi.org/10.1099/ijs.0.65316-0 |
| GCA_000023945.1 | Mesophiles | | 3116795 | Archaea | Halobacteriota | Halobacteria | Halobacteriales | Haloarculaceae | Halorhabdus | *Halorhabdus utahensis* | 519442 | *Halorhabdus utahensis DSM 12940* | https://www.microbiologyresearch.org/content/journal/ijsem/10.1099/00207713-50-1-183 |
| GCA_001542905.1 | Mesophiles | | 3325770 | Archaea | Halobacteriota | Halobacteria | Halobacteriales | Haloferacaceae | Halorubrum | *Halorubrum aethiopicum* | 1758255 | *Halorubrum aethiopicum* | https://doi.org/10.1099/ijsem.0.002525 |
| GCA_000336995.1 | Thermophiles | | 3108525 | Archaea | Halobacteriota | Halobacteria | Halobacteriales | Haloferacaceae | Halorubrum | *Halorubrum aidingense* | 1230454 | *Halorubrum aidingense* | https://www.microbiologyresearch.org/content/journal/ijsem/10.1099/ijs.0.64305-0 |
| GCA_900182635.1 | Mesophiles | | 3176020 | Archaea | Halobacteriota | Halobacteria | Halobacteriales | Haloferacaceae | Halorubrum | *Halorubrum cibi* | 413815 | *Halorubrum cibi* | https://link.springer.com/article/10.1007/s12275-009-0016-y |
| GCA_000337035.1 | Mesophiles | | 3645313 | Archaea | Halobacteriota | Halobacteria | Halobacteriales | Haloferacaceae | Halorubrum | *Halorubrum coriense* | 1227466 | *Halorubrum coriense* | https://bacdive.dsmz.de/strain/5944 |
| GCA_000022205.1 | Psychrophiles | | 3692576 | Archaea | Halobacteriota | Halobacteria | Halobacteriales | Haloferacaceae | Halorubrum | *Halorubrum lacusprofundi* | 416348 | *Halorubrum lacusprofundi ATCC 49239* | https://doi.org/10.1111/1462-2920.13705 |
| GCA_000337375.1 | Thermophiles | | 3425042 | Archaea | Halobacteriota | Halobacteria | Halobacteriales | Haloferacaceae | Halorubrum | *Halorubrum lipolyticum* | 1227482 | *Halorubrum lipolyticum* | https://www.microbiologyresearch.org/content/journal/ijsem/10.1099/ijs.0.64305-0 |
| GCA_000337915.1 | Thermophiles | | 3423703 | Archaea | Halobacteriota | Halobacteria | Halobacteriales | Haloferacaceae | Halorubrum | *Halorubrum saccharovorum* | 1227484 | *Halorubrum saccharovorum* | https://www.ncbi.nlm.nih.gov/pmc/articles/PMC545725/ |
| GCA_900111935.1 | Mesophiles | | 3030553 | Archaea | Halobacteriota | Halobacteria | Halobacteriales | Haloferacaceae | Halorubrum | *Halorubrum sodomense* | 35743 | *Halorubrum sodomense* | https://www.microbiologyresearch.org/content/journal/ijsem/10.1099/00207713-33-2-381 |
| GCA_018228765.1 | Mesophiles | | 3021820 | Archaea | Halobacteriota | Halobacteria | Halobacteriales | Haloferacaceae | Halorubrum | *Halorubrum sp000296615* | 35743 | *Halorubrum ruber* | https://doi.org/10.1007/s12275-022-2173-1 |
| GCA_900188065.1 | Mesophiles | Alkaliphiles | 3477860 | Archaea | Halobacteriota | Halobacteria | Halobacteriales | Haloferacaceae | Halorubrum | *Halorubrum vacuolatum* | 63740 | *Halorubrum vacuolatum* | https://bacdive.dsmz.de/strain/5943 |
| GCA_004765815.2 | Mesophiles | | 4753237 | Archaea | Halobacteriota | Halobacteria | Halobacteriales | Haladaptataceae | Halorussus | *Halorussus salinus* | 1364935 | *Halorussus salinus* | https://doi.org/10.1007/s00203-016-1253-1 |
| GCA_900116205.1 | Mesophiles | | 4108147 | Archaea | Halobacteriota | Halobacteria | Halobacteriales | Natrialbaceae | Halostagnicola | *Halostagnicola kamekurae* | 619731 | *Halostagnicola kamekurae* | https://doi.org/10.1099/ijs.0.014449-0 |
| GCA_005954745.1 | Mesophiles | | 3942449 | Archaea | Halobacteriota | Halobacteria | Halobacteriales | QS-9-68-17 | Halostella | *Halostella pelagica* | 2583824 | *Halostella pelagica* | https://doi.org/10.1099/ijsem.0.004003 |
| GCA_000025325.1 | Thermophiles | | 5440782 | Archaea | Halobacteriota | Halobacteria | Halobacteriales | Natrialbaceae | Haloterrigena | *Haloterrigena turkmenica* | 543526 | *Haloterrigena turkmenica* | https://www.ncbi.nlm.nih.gov/pmc/articles/PMC545725/ |
| GCA_000378345.1 | | Alkaliphiles | 3351270 | Bacteria | Proteobacteria | Alphaproteobacteria | Caulobacterales | Hyphomonadaceae | Hirschia | *Hirschia maritima* | 551275 | *Hirschia maritima DSM 19733* | https://www.sciencedirect.com/science/article/pii/S221359601530101X?via%3Dihub |
| GCA_003574215.1 | Thermophiles | | 2288780 | Bacteria | Proteobacteria | Gammaproteobacteria | Burkholderiales | Rhodocyclaceae | Hydrogenophilus | *Hydrogenophilus thermoluteolus* | 297 | *Hydrogenophilus thermoluteolus* | https://doi.org/10.1099/00207713-49-2-783 |
| GCA_000015145.1 | Hyperthermophiles | | 1667163 | Archaea | Thermoproteota | Thermoproteia | Sulfolobales | Pyrodictiaceae | Hyperthermus | *Hyperthermus butylicus* | 415426 | *Hyperthermus butylicus DSM 5456* | https://doi.org/10.1128/jb.172.7.3959-3965.1990 |
| GCA_001481685.1 | Hyperthermophiles | | 1394664 | Archaea | Thermoproteota | Thermoproteia | Sulfolobales | Ignicoccaceae | Ignicoccus | *Ignicoccus islandicus* | 54259 | *Ignicoccus islandicus DSM 13165* | https://doi.org/10.1099/00207713-50-6-2093 |
| GCA_000017945.1 | Hyperthermophiles | | 1297538 | Archaea | Thermoproteota | Thermoproteia | Sulfolobales | Ignicoccaceae | Ignicoccus_A | *Ignicoccus_A hospitalis* | 453591 | *Ignicoccus hospitalis KIN4/I* | https://www.microbiologyresearch.org/content/journal/ijsem/10.1099/ijs.0.64721-0 |
| GCA_000145985.1 | Hyperthermophiles | | 1875953 | Archaea | Thermoproteota | Thermoproteia | Sulfolobales | Ignisphaeraceae | Ignisphaera | *Ignisphaera aggregans* | 334771 | *Ignisphaera aggregans AQ1.S1, DSM 17230* | https://doi.org/10.1099/ijs.0.63899-0 |
| GCA_000525795.1 | Mesophiles | | 2601148 | Bacteria | Firmicutes | Bacilli | Lactobacillales | Listeriaceae | Listeria | *Listeria aquatica* | 1265818 | *Listeria aquatica FSL S10-1188* | https://www.microbiologyresearch.org/content/journal/ijsem/10.1099/ijs.0.052720-0 |

| | | | | | | | | | | | | | |
|---|---|---|---|---|---|---|---|---|---|---|---|---|---|
| GCA_000344175.1 | Mesophiles | | 2841134 | Bacteria | Firmicutes | Bacilli | Lactobacillales | Listeriaceae | Listeria | *Listeria fleischmannii* | 1430899 | *Listeria fleischmannii 1991* | https://bacdive.dsmz.de/strain/23081 |
| GCA_000525875.1 | Mesophiles | | 2794388 | Bacteria | Firmicutes | Bacilli | Lactobacillales | Listeriaceae | Listeria | *Listeria floridensis* | 1265817 | *Listeria floridensis FSL S10-1187* | https://www.microbiologyresearch.org/content/journal/ijsem/10.1099/ijs.0.052720-0 |
| GCA_000148995.1 | Mesophiles | | 2598321 | Bacteria | Firmicutes | Bacilli | Lactobacillales | Listeriaceae | Listeria | *Listeria grayi* | 1641 | *Listeria grayi* | https://bacdive.dsmz.de/strain/6870 |
| GCA_015276835.1 | Mesophiles | | 2878821 | Bacteria | Firmicutes | Bacilli | Lactobacillales | Listeriaceae | Listeria | *Listeria innocua* | 1642 | *Listeria innocua* | https://bacdive.dsmz.de/strain/136614 |
| GCA_900187025.1 | Mesophiles | | 2919550 | Bacteria | Firmicutes | Bacilli | Lactobacillales | Listeriaceae | Listeria | *Listeria ivanovii* | 881621 | *Listeria ivanovii subsp. ivanovii PAM 55* | https://bacdive.dsmz.de/strain/6872 |
| GCA_013282665.1 | Mesophiles | | 2788056 | Bacteria | Firmicutes | Bacilli | Lactobacillales | Listeriaceae | Listeria | *Listeria monocytogenes_C* | 552536 | *Listeria monocytogenes HCC23* | https://bacdive.dsmz.de/strain/6875 |
| GCA_000027145.1 | Mesophiles | | 2797636 | Bacteria | Firmicutes | Bacilli | Lactobacillales | Listeriaceae | Listeria | *Listeria seeligeri* | 1640 | *Listeria seeligeri* | https://www.microbiologyresearch.org/content/journal/ijsem/10.1099/00207713-33-4-866 |
| GCA_000060285.1 | Mesophiles | | 2814130 | Bacteria | Firmicutes | Bacilli | Lactobacillales | Listeriaceae | Listeria | *Listeria welshimeri* | 1643 | *Listeria welshimeri* | https://www.microbiologyresearch.org/content/journal/ijsem/10.1099/00207713-33-4-866 |
| GCA_000766865.1 | Mesophiles | | 3436956 | Bacteria | Firmicutes | Bacilli | Lactobacillales | Listeriaceae | Listeria_A | *Listeria_A booriae* | 1552123 | *Listeria booriae* | https://www.microbiologyresearch.org/content/journal/ijsem/10.1099/ijs.0.070839-0 |
| GCA_000766145.1 | Mesophiles | | 3515436 | Bacteria | Firmicutes | Bacilli | Lactobacillales | Listeriaceae | Listeria_A | *Listeria_A newyorkensis* | 1497681 | *Listeria newyorkensis* | https://www.microbiologyresearch.org/content/journal/ijsem/10.1099/ijs.0.070839-0 |
| GCA_000525995.1 | Mesophiles | | 3291042 | Bacteria | Firmicutes | Bacilli | Lactobacillales | Listeriaceae | Listeria_A | *Listeria_A riparia* | 1265816 | *Listeria riparia FSL S10-1204* | https://www.microbiologyresearch.org/content/journal/ijsem/10.1099/ijs.0.052720-0 |
| GCA_000525975.1 | Mesophiles | | 3216749 | Bacteria | Firmicutes | Bacilli | Lactobacillales | Listeriaceae | Listeria_A | *Listeria_A rocourtiae* | 647910 | *Listeria rocourtiae* | https://www.microbiologyresearch.org/content/journal/ijsem/10.1099/ijs.0.017376-0 |
| GCA_003534205.1 | Mesophiles | | 3406292 | Bacteria | Firmicutes | Bacilli | Lactobacillales | Listeriaceae | Listeria_A | *Listeria_A weihenstephanensis* | 1006155 | *Listeria weihenstephanensis* | https://www.microbiologyresearch.org/content/journal/ijsem/10.1099/ijs.0.036830-0 |
| GCA_009765685.1 | | Acidophiles | 6380195 | Bacteria | Proteobacteria | Alphaproteobacteria | Acetobacterales | Acetobacteraceae | LMUY01 | *LMUY01 sp009765685* | 1641851 | *Acidisphaera sp. L21* | https://doi.org/10.1099/00207713-50-4-1539 |
| GCA_003970575.1 | Mesophiles | | 2605275 | Bacteria | Firmicutes | Bacilli | Staphylococcales | Staphylococcaceae | Mammaliicoccus | *Mammaliicoccus fleurettii* | 150056 | *Mammaliicoccus fleurettii* | https://bacdive.dsmz.de/strain/14648 |
| GCA_002902755.1 | Mesophiles | | 2546437 | Bacteria | Firmicutes | Bacilli | Staphylococcales | Staphylococcaceae | Mammaliicoccus | *Mammaliicoccus lentus* | 42858 | *Mammaliicoccus lentus* | https://bacdive.dsmz.de/strain/14560 |
| GCA_002901825.1 | Mesophiles | | 2768322 | Bacteria | Firmicutes | Bacilli | Staphylococcales | Staphylococcaceae | Mammaliicoccus | *Mammaliicoccus sciuri* | 1296 | *Mammaliicoccus sciuri* | https://bacdive.dsmz.de/strain/14631 |
| GCA_002902265.1 | Mesophiles | | 2595808 | Bacteria | Firmicutes | Bacilli | Staphylococcales | Staphylococcaceae | Mammaliicoccus | *Mammaliicoccus vitulinus* | 71237 | *Mammaliicoccus vitulinus* | https://bacdive.dsmz.de/strain/138135 |
| GCA_900142385.1 | Psychrophiles | | 3740786 | Bacteria | Proteobacteria | Gammaproteobacteria | Pseudomonadales | Oleiphilaceae | Marinobacter | *Marinobacter antarcticus* | 564117 | *Marinobacter antarcticus* | https://doi.org/10.1099/ijs.0.035774-0 |
| GCA_011049865.1 | Psychrophiles | | 3929789 | Bacteria | Proteobacteria | Gammaproteobacteria | Pseudomonadales | Oleiphilaceae | Marinobacter | *Marinobacter antarcticus_A* | 564117 | *Marinobacter antarcticus* | https://doi.org/10.1099/ijs.0.035774-0 |
| GCA_007671675.1 | Psychrophiles | | 4357025 | Bacteria | Proteobacteria | Gammaproteobacteria | Pseudomonadales | Oleiphilaceae | Marinobacter | *Marinobacter maritimus* | 277961 | *Marinobacter maritimus* | https://doi.org/10.1099/ijs.0.63478-0 |
| GCA_900106945.1 | Mesophiles | | 3971609 | Bacteria | Proteobacteria | Gammaproteobacteria | Pseudomonadales | Oleiphilaceae | Marinobacter | *Marinobacter mobilis* | 488533 | *Marinobacter mobilis* | https://www.microbiologyresearch.org/content/journal/ijsem/10.1099/ijs.0.2008/000786-0 |
| GCA_007671655.1 | Mesophiles | | 3336972 | Bacteria | Proteobacteria | Gammaproteobacteria | Pseudomonadales | Oleiphilaceae | Marinobacter | *Marinobacter piscensis* | 1562308 | *Marinobacter piscensis* | https://doi.org/10.1007/s00284-014-0754-x |
| GCA_001043175.1 | Psychrophiles | | 3998597 | Bacteria | Proteobacteria | Gammaproteobacteria | Pseudomonadales | Oleiphilaceae | Marinobacter | *Marinobacter psychrophilus* | 330734 | *Marinobacter psychrophilus* | https://doi.org/10.1099/ijs.0.65690-0 |
| GCA_000831005.1 | Mesophiles | | 4616532 | Bacteria | Proteobacteria | Gammaproteobacteria | Pseudomonadales | Oleiphilaceae | Marinobacter | *Marinobacter salarius* | 1420917 | *Marinobacter salarius* | https://doi.org/10.1371/journal.pone.0106514 |
| GCA_002806945.1 | | Alkaliphiles | 3820421 | Bacteria | Proteobacteria | Gammaproteobacteria | Pseudomonadales | Oleiphilaceae | Marinobacter | *Marinobacter salexigens* | 1925763 | *Marinobacter salexigens* | https://doi.org/10.1099/ijsem.0.002337 |
| GCA_018860765.1 | | Alkaliphiles | 4133335 | Bacteria | Proteobacteria | Gammaproteobacteria | Pseudomonadales | Oleiphilaceae | Marinobacter | *Marinobacter salexigens_A* | 1925763 | *Marinobacter salexigens* | https://doi.org/10.1099/ijsem.0.002337 |
| GCA_009617755.1 | Mesophiles | | 4150758 | Bacteria | Proteobacteria | Gammaproteobacteria | Pseudomonadales | Oleiphilaceae | Marinobacter | *Marinobacter salsuginis* | 418719 | *Marinobacter salsuginis* | https://doi.org/10.1099/ijs.0.64862-0 |
| GCA_000347775.1 | Mesophiles | | 4033468 | Bacteria | Proteobacteria | Gammaproteobacteria | Pseudomonadales | Oleiphilaceae | Marinobacter | *Marinobacter santoriniensis* | 1288826 | *Marinobacter santoriniensis NKSG1* | https://doi.org/10.1099/ijs.0.003145-0 |
| GCA_900111555.1 | Mesophiles | | 4218891 | Bacteria | Proteobacteria | Gammaproteobacteria | Pseudomonadales | Oleiphilaceae | Marinobacter | *Marinobacter segnicrescens* | 430453 | *Marinobacter segnicrescens* | https://doi.org/10.1099/ijs.0.65030-0 |
| GCA_000364845.1 | Mesophiles | | 5358909 | Bacteria | Proteobacteria | Gammaproteobacteria | Pseudomonadales | Oleiphilaceae | Marinobacter_A | *Marinobacter_A nanhaiticus* | 626887 | *Marinobacter nanhaiticus D15-8W* | https://pubmed.ncbi.nlm.nih.gov/23117603/ |
| GCA_000620065.1 | Thermophiles | | 3034817 | Bacteria | Deinococcota | Deinococci | Deinococcales | Thermaceae | Meiothermus | *Meiothermus cerbereus* | 1122221 | *Meiothermus cerbereus DSM 11376* | https://doi.org/10.1099/00207713-47-4-1225 |
| GCA_003574035.1 | Thermophiles | | 3684677 | Bacteria | Deinococcota | Deinococci | Deinococcales | Thermaceae | Meiothermus | *Meiothermus hypogaeus* | 884155 | *Meiothermus hypogaeus* | https://doi.org/10.1099/ijs.0.028654-0 |
| GCA_003574085.1 | Thermophiles | | 2874609 | Bacteria | Deinococcota | Deinococci | Deinococcales | Thermaceae | Meiothermus | *Meiothermus luteus* | 2026184 | *Meiothermus luteus* | https://doi.org/10.1099/ijsem.0.002040 |
| GCA_000423425.1 | Thermophiles | | 2747076 | Bacteria | Deinococcota | Deinococci | Deinococcales | Thermaceae | Meiothermus | *Meiothermus rufus* | 604331 | *Meiothermus rufus DSM 22234* | https://doi.org/10.1016/j.syapm.2009.05.002 |
| GCA_000482765.1 | Thermophiles | | 3020616 | Bacteria | Deinococcota | Deinococci | Deinococcales | Thermaceae | Meiothermus | *Meiothermus taiwanensis* | 172827 | *Meiothermus taiwanensis* | https://doi.org/10.1099/00207713-52-5-1647 |
| GCA_000092125.1 | Thermophiles | | 3721669 | Bacteria | Deinococcota | Deinococci | Deinococcales | Thermaceae | Meiothermus_B | *Meiothermus_B silvanus* | 526227 | *Meiothermus silvanus DSM 9946* | https://bacdive.dsmz.de/strain/16703 |
| GCA_000204925.1 | Thermophiles | Acidophiles | 1840348 | Archaea | Thermoproteota | Thermoproteia | Sulfolobales | Sulfolobaceae | Metallosphaera | *Metallosphaera cuprina* | 1006006 | *Metallosphaera cuprina Ar-4* | https://doi.org/10.1099/ijs.0.026591-0 |
| GCA_003201675.2 | Thermophiles | Acidophiles | 2544115 | Archaea | Thermoproteota | Thermoproteia | Sulfolobales | Sulfolobaceae | Metallosphaera | *Metallosphaera hakonensis* | 1293036 | *Metallosphaera hakonensis JCM 8857 = DSM 7519* | https://doi.org/10.1099%2F00207713-46-2-377 |
| GCA_000016605.1 | Thermophiles | | 2191517 | Archaea | Thermoproteota | Thermoproteia | Sulfolobales | Sulfolobaceae | Metallosphaera | *Metallosphaera sedula* | 43687 | *Metallosphaera sedula* | https://bacdive.dsmz.de/strain/16645 |
| GCA_013343295.1 | Thermophiles | Acidophiles | 2176897 | Archaea | Thermoproteota | Thermoproteia | Sulfolobales | Sulfolobaceae | Metallosphaera | *Metallosphaera tengchongensis* | 1532350 | *Metallosphaera tengchongensis* | https://doi.org/10.1099/ijs.0.070870-0 |
| GCA_000243315.1 | Thermophiles | Acidophiles | 2817452 | Archaea | Thermoproteota | Thermoproteia | Sulfolobales | Sulfolobaceae | Metallosphaera | *Metallosphaera yellowstonensis* | 671065 | *Metallosphaera yellowstonensis MK1* | https://doi.org/10.1128/aem.03416-13 |
| GCA_017873625.1 | Mesophiles | | 2468550 | Archaea | Methanobacteriota | Methanobacteria | Methanobacteriales | Methanobacteriaceae | Methanobacterium | *Methanobacterium petrolearium* | 710190 | *Methanobacterium petrolearium* | https://doi.org/10.1099/ijs.0.022723-0 |
| GCA_002813695.1 | Mesophiles | | 2515817 | Archaea | Methanobacteriota | Methanobacteria | Methanobacteriales | Methanobacteriaceae | Methanobacterium | *Methanobacterium subterraneum* | 59277 | *Methanobacterium subterraneum* | https://doi.org/10.1099/00207713-48-2-357 |
| GCA_000214725.1 | Mesophiles | | 2546541 | Archaea | Methanobacteriota | Methanobacteria | Methanobacteriales | Methanobacteriaceae | Methanobacterium_C | *Methanobacterium_C paludis* | 868131 | *Methanobacterium paludis* | https://doi.org/10.1099/ijs.0.059964-0 |
| GCA_001639275.1 | Mesophiles | | 2140433 | Archaea | Methanobacteriota | Methanobacteria | Methanobacteriales | Methanobacteriaceae | Methanobrevibacter_A | *Methanobrevibacter_A oralis* | 1415626 | *Methanobrevibacter oralis* | https://bacdive.dsmz.de/strain/6969 |
| GCA_002072215.1 | Mesophiles | | 2445031 | Archaea | Methanobacteriota | Methanobacteria | Methanobacteriales | Methanobacteriaceae | Methanobrevibacter_C | *Methanobrevibacter_C arboriphilus* | 39441 | *Methanobrevibacter arboriphilus* | https://bacdive.dsmz.de/strain/6957 |
| GCA_000513315.1 | Mesophiles | | 2221920 | Archaea | Methanobacteriota | Methanobacteria | Methanobacteriales | Methanobacteriaceae | Methanobrevibacter_C | *Methanobrevibacter_C arboriphilus_A* | 39441 | *Methanobrevibacter arboriphilus* | https://bacdive.dsmz.de/strain/6957 |
| GCA_000739065.1 | Hyperthermophiles | | 1607556 | Archaea | Methanobacteriota_A | Methanococci | Methanococcales | Methanocaldococcaceae | Methanocaldococcus | *Methanocaldococcus bathoardescens* | | *Methanocaldococcus bathoardescens* | https://doi.org/10.1099/ijs.0.000097 |
| GCA_000023985.1 | Thermophiles | | 1507251 | Archaea | Methanobacteriota_A | Methanococci | Methanococcales | Methanocaldococcaceae | Methanocaldococcus | *Methanocaldococcus fervens* | 573064 | *Methanocaldococcus fervens AG86* | https://pubmed.ncbi.nlm.nih.gov/10319479/ |
| GCA_000091665.1 | Thermophiles | | 1739927 | Archaea | Methanobacteriota_A | Methanococci | Methanococcales | Methanocaldococcaceae | Methanocaldococcus | *Methanocaldococcus jannaschii* | 2190 | *Methanocaldococcus jannaschii UBA8849* | https://bacdive.dsmz.de/strain/6981 |
| GCA_000024625.1 | Thermophiles | | 1761737 | Archaea | Methanobacteriota_A | Methanococci | Methanococcales | Methanocaldococcaceae | Methanocaldococcus | *Methanocaldococcus vulcanius* | 579137 | *Methanocaldococcus vulcanius* | https://bacdive.dsmz.de/strain/6983 |
| GCA_000251105.1 | Thermophiles | | 2378438 | Archaea | Halobacteriota | Methanocellia | Methanocellales | Methanocellaceae | Methanocella | *Methanocella conradii* | 1041930 | *Methanocella conradii* | https://doi.org/10.1371/journal.pone.0035279 |
| GCA_000013725.1 | Psychrophiles | | 2575032 | Archaea | Halobacteriota | Methanosarcinia | Methanosarcinales | Methanosarcinaceae | Methanococcoides | *Methanococcoides burtonii* | 259564 | *Methanococcoides burtonii DSM 6242* | https://doi.org/10.1038/ismej.2009.45 |
| GCA_002945325.1 | Mesophiles | | 1714918 | Archaea | Methanobacteriota_A | Methanococci | Methanococcales | Methanococcaceae | Methanococcus | *Methanococcus maripaludis* | 267377 | *Methanococcus maripaludis S2* | https://bacdive.dsmz.de/strain/6991 |
| GCA_000304355.2 | Mesophiles | | 2789774 | Archaea | Halobacteriota | Methanomicrobia | Methanomicrobiales | Methanoculleaceae | Methanoculleus | *Methanoculleus bourgensis* | 1201294 | *Methanoculleus bourgensis* | https://bacdive.dsmz.de/strain/7016 |
| GCA_900095385.1 | Mesophiles | | 2649157 | Archaea | Halobacteriota | Methanomicrobia | Methanomicrobiales | Methanoculleaceae | Methanoculleus | *Methanoculleus chikugoensis_A* | 1293042 | *Methanoculleus chikugoensis* | https://doi.org/10.1099/00207713-51-5-1663 |
| GCA_001602375.1 | Mesophiles | | 2446106 | Archaea | Halobacteriota | Methanomicrobia | Methanomicrobiales | Methanoculleaceae | Methanoculleus | *Methanoculleus horonobensis* | 528314 | *Methanoculleus horonobensis strain T10* | https://doi.org/10.1099/ijs.0.053520-0 |
| GCA_000015825.1 | Mesophiles | | 2478101 | Archaea | Halobacteriota | Methanomicrobia | Methanomicrobiales | Methanoculleaceae | Methanoculleus | *Methanoculleus marisnigri* | 2198 | *Methanoculleus marisnigri isolate 63_41* | https://bacdive.dsmz.de/strain/7020 |
| GCA_002503885.1 | Mesophiles | | 2582043 | Archaea | Halobacteriota | Methanomicrobia | Methanomicrobiales | Methanoculleaceae | Methanoculleus | *Methanoculleus marisnigri_B* | 368407 | *Methanoculleus marisnigri JR1* | https://bacdive.dsmz.de/strain/7020 |

| | | | | | | | | | | | | | |
|---|---|---|---|---|---|---|---|---|---|---|---|---|---|
| GCA_001017125.1 | Mesophiles | | 2489717 | Archaea | Halobacteriota | Methanomicrobia | Methanomicrobiales | Methanoculleaceae | Methanoculleus | Methanoculleus sediminis | 1550566 | Methanoculleus sediminis strain S3Fa | https://doi.org/10.1099/ijs.0.000233 |
| GCA_001571405.1 | Thermophiles | | 2223235 | Archaea | Halobacteriota | Methanomicrobia | Methanomicrobiales | Methanoculleaceae | Methanoculleus | Methanoculleus thermophilus | 2200 | Methanoculleus thermophilus strain CR-1 | https://bacdive.dsmz.de/strain/7022 |
| GCA_004102725.1 | Mesophiles | | 2750720 | Archaea | Halobacteriota | Methanomicrobia | Methanomicrobiales | Methanoculleaceae | Methanoculleus_A | Methanoculleus_A taiwanensis | 1550565 | Methanoculleus taiwanensis strain CYW4 | https://doi.org/10.1099/ijs.0.000062 |
| GCA_000275865.1 | Mesophiles | | 2475100 | Archaea | Halobacteriota | Methanomicrobia | Methanomicrobiales | Methanofollaceae | Methanofollis | Methanofollis liminatans | 28892 | Methanofollis liminatans | https://www.microbiologyresearch.org/content/journal/ijsem/10.1099/00207713-49-1-247 |
| GCA_009914725.1 | Psychrophiles | | 2189363 | Archaea | Halobacteriota | Methanomicrobia | Methanomicrobiales | Methanomicrobiaceae | Methanogenium | Methanogenium sp009914725 | 2599926 | Methanogenium sp. MK-MG | https://www.nature.com/articles/s41586-019-1916-6 |
| GCA_900215215.1 | Thermophiles | | 1940298 | Archaea | Halobacteriota | Methanosarcinia | Methanosarcinales | Methanosarcinaceae | Methanohalophilus | Methanohalophilus euhalobius | 51203 | Methanohalophilus euhalobius strain DSM 10369 | https://link.springer.com/article/10.1023/A:1000103618451 |
| GCA_001889405.1 | Mesophiles | | 2022959 | Archaea | Halobacteriota | Methanosarcinia | Methanosarcinales | Methanosarcinaceae | Methanohalophilus | Methanohalophilus halophilus | 2177 | Methanohalophilus halophilus strain DSM 3094 | https://bacdive.dsmz.de/strain/7064 |
| GCA_017874375.1 | Mesophiles | | 2116010 | Archaea | Halobacteriota | Methanosarcinia | Methanosarcinales | Methanosarcinaceae | Methanohalophilus | Methanohalophilus levihalophilus | 1431282 | Methanohalophilus levihalophilus | https://doi.org/10.1099/ijs.0.063677-0 |
| GCA_000025865.1 | Mesophiles | | 2012424 | Archaea | Halobacteriota | Methanosarcinia | Methanosarcinales | Methanosarcinaceae | Methanohalophilus | Methanohalophilus mahii | 547558 | Methanohalophilus mahii DSM 5219 | https://bacdive.dsmz.de/strain/7068 |
| GCA_002761295.1 | Mesophiles | | 2084975 | Archaea | Halobacteriota | Methanosarcinia | Methanosarcinales | Methanosarcinaceae | Methanohalophilus | Methanohalophilus portucalensis | 523843 | Methanohalophilus portucalensis strain FDF-1T chromosome | https://bacdive.dsmz.de/strain/7074 |
| GCA_004137855.1 | Mesophiles | | 1830088 | Archaea | Halobacteriota | Methanosarcinia | Methanosarcinales | Methanosarcinaceae | Methanohalophilus | Methanohalophilus profundi | 2138083 | Methanohalophilus profundi | https://doi.org/10.1016/j.syapm.2020.126107 |
| GCA_003722075.1 | Mesophiles | | 1969036 | Archaea | Halobacteriota | Methanosarcinia | Methanosarcinales | Methanosarcinaceae | Methanohalophilus | Methanohalophilus sp003722075 | 2485783 | Methanohalophilus sp. RSK | https://www.frontiersin.org/articles/10.3389/fmicb.2019.00839/full |
| GCA_000784355.1 | Mesophiles | | 2791704 | Archaea | Halobacteriota | Methanomicrobia | Methanomicrobiales | Methanomicrobiaceae | Methanolacinia | Methanolacinia paynteri | 694436 | Methanolacinia paynteri | https://bacdive.dsmz.de/strain/7039 |
| GCA_000147875.1 | Mesophiles | | 2843290 | Archaea | Halobacteriota | Methanomicrobia | Methanomicrobiales | Methanomicrobiaceae | Methanolacinia | Methanolacinia petrolearia | 679926 | Methanolacinia petrolearia | https://bacdive.dsmz.de/strain/7043 |
| GCA_017873855.1 | Mesophiles | | 2662345 | Archaea | Halobacteriota | Methanomicrobia | Methanomicrobiales | Methanoregulaceae | Methanolinea_B | Methanolinea_B mesophila | 547055 | Methanolinea mesophila | https://doi.org/10.1099/ijs.0.035048-0 |
| GCA_900114835.1 | Mesophiles | | 3107200 | Archaea | Halobacteriota | Methanosarcinia | Methanosarcinales | Methanosarcinaceae | Methanolobus | Methanolobus profundi | 487685 | Methanolobus profundi strain Mob M | https://doi.org/10.1099/ijs.0.001677-0 |
| GCA_000306725.1 | Psychrophiles | | 3072769 | Archaea | Halobacteriota | Methanosarcinia | Methanosarcinales | Methanosarcinaceae | Methanolobus | Methanolobus psychrophilus | | Methanolobus psychrophilus R15 | https://journals.asm.org/doi/10.1128/AEM.01146-08 |
| GCA_002243045.1 | Psychrophiles | | 3164721 | Archaea | Halobacteriota | Methanosarcinia | Methanosarcinales | Methanosarcinaceae | Methanolobus | Methanolobus psychrotolerans | 1874706 | Methanolobus psychrotolerans strain YSF-03 | https://doi.org/10.1099/ijsem.0.002685 |
| GCA_004745425.1 | Mesophiles | Alkaliphiles | 2592212 | Archaea | Halobacteriota | Methanosarcinia | Methanosarcinales | Methanosarcinaceae | Methanolobus | Methanolobus sp004745425 | 2052935 | Methanolobus halotolerans | https://doi.org/10.1099/ijsem.0.004453 |
| GCA_000504205.1 | Mesophiles | | 3151883 | Archaea | Halobacteriota | Methanosarcinia | Methanosarcinales | Methanosarcinaceae | Methanolobus | Methanolobus tindarius | 1090322 | Methanolobus tindarius DSM 2278 | https://bacdive.dsmz.de/strain/7076 |
| GCA_013388255.1 | Thermophiles | | 2704953 | Archaea | Halobacteriota | Methanosarcinia | Methanosarcinales | Methanosarcinaceae | Methanolobus | Methanolobus zinderi | 536044 | Methanolobus zinderi strain DSM 21339 | https://doi.org/10.1099/ijs.0.003772-0 |
| GCA_000711215.1 | Mesophiles | | 1711791 | Archaea | Halobacteriota | Methanomicrobia | Methanomicrobiales | Methanomicrobiaceae | Methanomicrobium | Methanomicrobium mobile | 694440 | Methanomicrobium mobile | https://bacdive.dsmz.de/strain/7040 |
| GCA_002153915.1 | Thermophiles | Alkaliphiles | 1513137 | Archaea | Halobacteriota | Methanonatronarchaeia | Methanonatronarchaeales | Methanonatronarchaeaceae | Methanonatronarchaeum | Methanonatronarchaeum thermophilum | 1927129 | Methanonatronarchaeum thermophilum | https://doi.org/10.1099/ijsem.0.002810 |
| GCA_000243255.1 | Mesophiles | | 3200946 | Archaea | Halobacteriota | Methanomicrobia | Methanomicrobiales | Methanomicrobiaceae | Methanoplanus | Methanoplanus limicola | 937775 | Methanoplanus limicola | http://standardsingenomics.org/content/9/3/1076/ |
| GCA_000007185.1 | Hyperthermophiles | | 1694969 | Archaea | Methanobacteriota_A | Methanopyri | Methanopyrales | Methanopyraceae | Methanopyrus | Methanopyrus kandleri | 190192 | Methanopyrus kandleri AV19 | https://doi.org/10.1042/bst0320269 |
| GCA_002201895.1 | Hyperthermophiles | | 1421621 | Archaea | Methanobacteriota_A | Methanopyri | Methanopyrales | Methanopyraceae | Methanopyrus | Methanopyrus sp002201895 | 1937004 | Methanopyrus sp. KOL6 | https://www.frontiersin.org/articles/10.3389/fmicb.2017.01278/full |
| GCA_000327485.1 | Mesophiles | | 2820858 | Archaea | Halobacteriota | Methanomicrobia | Methanomicrobiales | Methanoregulaceae | Methanoregula | Methanoregula formicica | 593750 | Methanoregula formicica | https://doi.org/10.1099/ijs.0.014811-0 |
| GCA_000217995.1 | | Alkaliphiles | 2138444 | Archaea | Halobacteriota | Methanosarcinia | Methanosarcinales | Methanosarcinaceae | Methanosalsum | Methanosalsum zhilinae | 679901 | Methanosalsum zhilinae DSM 4017 | https://doi.org/10.1099/ijsem.0.000488 |
| GCA_000970285.1 | Mesophiles | | 5018607 | Archaea | Halobacteriota | Methanosarcinia | Methanosarcinales | Methanosarcinaceae | Methanosarcina | Methanosarcina horonobensis | 1434110 | Methanosarcina horonobensis | https://doi.org/10.1099/ijs.0.028548-0 |
| GCA_000970265.1 | Psychrophiles | | 4139808 | Archaea | Halobacteriota | Methanosarcinia | Methanosarcinales | Methanosarcinaceae | Methanosarcina | Methanosarcina lacustris | 1434111 | Methanosarcina lacustris Z-7289 | https://doi.org/10.1078/0723-2020-00058 |
| GCA_000970205.1 | Mesophiles | | 4142816 | Archaea | Halobacteriota | Methanosarcinia | Methanosarcinales | Methanosarcinaceae | Methanosarcina | Methanosarcina mazei | 192952 | Methanosarcina mazei | https://bacdive.dsmz.de/strain/7096 |
| GCA_000970085.1 | Mesophiles | | 5017558 | Archaea | Halobacteriota | Methanosarcinia | Methanosarcinales | Methanosarcinaceae | Methanosarcina | Methanosarcina siciliae | 1434118 | Methanosarcina siciliae C2J | https://bacdive.dsmz.de/strain/7083 |
| GCA_002287235.1 | Mesophiles | | 5088600 | Archaea | Halobacteriota | Methanosarcinia | Methanosarcinales | Methanosarcinaceae | Methanosarcina | Methanosarcina spelaei | 1036679 | Methanosarcina spelaei | https://doi.org/10.1099/ijs.0.064956-0 |
| GCA_000969885.1 | Thermophiles | | 3127379 | Archaea | Halobacteriota | Methanosarcinia | Methanosarcinales | Methanosarcinaceae | Methanosarcina | Methanosarcina thermophila | 523844 | Methanosarcina thermophila TM-1 | https://bacdive.dsmz.de/strain/7121 |
| GCA_000969905.1 | Mesophiles | | 4563885 | Archaea | Halobacteriota | Methanosarcinia | Methanosarcinales | Methanosarcinaceae | Methanosarcina | Methanosarcina vacuolata | 1434123 | Methanosarcina vacuolata Z-761 | https://bacdive.dsmz.de/strain/7126 |
| GCA_000021965.1 | Mesophiles | | 2922917 | Archaea | Halobacteriota | Methanomicrobia | Methanomicrobiales | Methanosphaerulaceae | Methanosphaerula | Methanosphaerula palustris | 521011 | Methanosphaerula palustris | https://doi.org/10.1099/ijs.0.006890-0 |
| GCA_000013445.1 | Mesophiles | | 3544738 | Archaea | Halobacteriota | Methanomicrobia | Methanomicrobiales | Methanospirillaceae | Methanospirillum | Methanospirillum hungatei | 323259 | Methanospirillum hungatei | https://bacdive.dsmz.de/strain/7132 |
| GCA_019263745.1 | Mesophiles | | 3393136 | Archaea | Halobacteriota | Methanomicrobia | Methanomicrobiales | Methanospirillaceae | Methanospirillum | Methanospirillum sp012729995 | 323259 | Methanospirillum hungatei | https://bacdive.dsmz.de/strain/7132 |
| GCA_003173335.1 | Psychrophiles | | 3740742 | Archaea | Halobacteriota | Methanomicrobia | Methanomicrobiales | Methanospirillaceae | Methanospirillum | Methanospirillum stamsii | 1277351 | Methanospirillum stamsii | https://doi.org/10.1099/ijs.0.056218-0 |
| GCA_000145295.1 | Thermophiles | | 1639135 | Archaea | Methanobacteriota | Methanobacteria | Methanobacteriales | Methanothermobacteraceae | Methanothermobacter | Methanothermobacter marburgensis | 2603820 | Methanothermobacter sp. KEPCO-1 | https://doi.org/10.1016/j.enzmictec.2022.110067 |
| GCA_000828575.1 | Thermophiles | | 1731018 | Archaea | Methanobacteriota | Methanobacteria | Methanobacteriales | Methanothermobacteraceae | Methanothermobacter | Methanothermobacter sp000828575 | 866790 | Methanothermobacter sp. CaT2 | https://doi.org/10.1128/genomea.00672-13 |
| GCA_000008645.1 | Thermophiles | | 1751377 | Archaea | Methanobacteriota | Methanobacteria | Methanobacteriales | Methanothermobacteraceae | Methanothermobacter | Methanothermobacter thermautotrophicus | 187420 | Methanothermobacter thermautotrophicus str. Delta H | https://bacdive.dsmz.de/strain/6882 |
| GCA_900095815.1 | Thermophiles | | 1686891 | Archaea | Methanobacteriota | Methanobacteria | Methanobacteriales | Methanothermobacteraceae | Methanothermobacter | Methanothermobacter wolfeii | 145261 | Methanothermobacter wolfeii SIV6 | https://doi.org/10.1099/00207713-50-1-43 |
| GCA_003264935.1 | Thermophiles | | 1467867 | Archaea | Methanobacteriota | Methanobacteria | Methanobacteriales | Methanothermobacteraceae_A | Methanothermobacter_A | Methanothermobacter_A tenebrarum | 680118 | Methanothermobacter tenebrarum | https://doi.org/10.1099/ijs.0.041681-0 |
| GCA_000017185.1 | Mesophiles | | 1569500 | Archaea | Methanobacteriota_A | Methanococci | Methanococcales | Methanococcaceae | Methanothermococcus_A | Methanothermococcus_A aeolicus | 42879 | Methanococcus aeolicus | https://doi.org/10.1099/ijs.0.64216-0 |
| GCA_000179575.2 | Thermophiles | | 1677455 | Archaea | Methanobacteriota_A | Methanococci | Methanococcales | Methanococcaceae | Methanothermococcus_A | Methanothermococcus_A okinawensis | 647113 | Methanothermococcus okinawensis IH1 | https://doi.org/10.1099/00207713-52-4-1089 |
| GCA_000166095.1 | Hyperthermophiles | | 1243342 | Archaea | Methanobacteriota | Methanobacteria | Methanobacteriales | Methanothermaceae | Methanothermus | Methanothermus fervidus | 523846 | Methanothermus fervidus V24S, DSM 2088 | https://doi.org/10.1128/jb.174.11.3508-3513.1992 |
| GCA_002502785.1 | Mesophiles | | 2512329 | Archaea | Halobacteriota | Methanosarcinia | Methanotrichales | Methanotrichaceae | Methanothrix_A | Methanothrix_A harundinacea_A | 1110509 | Methanosaeta harundinacea | https://bacdive.dsmz.de/strain/131756 |
| GCA_002506535.1 | Mesophiles | | 2342380 | Archaea | Halobacteriota | Methanosarcinia | Methanotrichales | Methanotrichaceae | Methanothrix_A | Methanothrix_A harundinacea_B | 1110509 | Methanosaeta harundinacea | https://bacdive.dsmz.de/strain/131756 |
| GCA_001509375.1 | Mesophiles | | 2382964 | Archaea | Halobacteriota | Methanosarcinia | Methanotrichales | Methanotrichaceae | Methanothrix_A | Methanothrix_A harundinacea_D | 1110509 | Methanosaeta harundinacea | https://bacdive.dsmz.de/strain/131756 |
| GCA_000235565.1 | Mesophiles | | 2571034 | Archaea | Halobacteriota | Methanosarcinia | Methanotrichales | Methanotrichaceae | Methanothrix_A | Methanothrix_A harundinacea_E | 1110509 | Methanosaeta harundinacea | https://bacdive.dsmz.de/strain/131756 |
| GCA_000014945.1 | Thermophiles | | 1879471 | Archaea | Halobacteriota | Methanosarcinia | Methanotrichales | Methanotrichaceae | Methanothrix_B | Methanothrix_B thermoacetophila | 349307 | Methanothrix thermoacetophila | https://doi.org/10.1099/00207713-42-3-463 |
| GCA_000243455.2 | Thermophiles | | 1818783 | Archaea | Methanobacteriota_A | Methanococci | Methanococcales | Methanococcaceae | Methanotorris | Methanotorris formicicus | 647171 | Methanotorris formicicus Mc-S-70 | https://doi.org/10.1099/ijs.0.02887-0 |
| GCA_000214415.1 | Hyperthermophiles | | 1854197 | Archaea | Methanobacteriota_A | Methanococci | Methanococcales | Methanococcaceae | Methanotorris | Methanotorris igneus | 880724 | Methanotorris igneus Kol 5 | https://doi.org/10.1038/s42003-021-01828-5 |
| GCA_902143385.2 | | Acidophiles | 2276790 | Bacteria | Verrucomicrobiota | Verrucomicrobiae | Methylacidiphilales | Methylacidiphilaceae | Methylacidimicrobium | Methylacidimicrobium cyclopophantes | 1041766 | Methylacidimicrobium cyclopophantes | https://doi.org/10.1128/mra.00315-20 |
| GCA_000953475.1 | Thermophiles | Acidophiles | 2476671 | Bacteria | Verrucomicrobiota | Verrucomicrobiae | Methylacidiphilales | Methylacidiphilaceae | Methylacidiphilum | Methylacidiphilum fumariolicum | 591154 | Methylacidiphilum fumariolicum | https://doi.org/10.1038/ismej.2016.171 |
| GCA_007475525.1 | Thermophiles | Acidophiles | 2202032 | Bacteria | Verrucomicrobiota | Verrucomicrobiae | Methylacidiphilales | Methylacidiphilaceae | Methylacidiphilum | Methylacidiphilum kamchatkense | 1202785 | Methylacidiphilum kamchatkense Kam1 | https://doi.org/10.1128/genomea.00065-15 |

| | | | | | | | | | | | | | |
|---|---|---|---|---|---|---|---|---|---|---|---|---|---|
| GCA_004421185.1 | | Acidophiles | 2250350 | Bacteria | Verrucomicrobiota | Verrucomicrobiae | Methylacidiphilales | Methylacidiphilaceae | Methylacidiphilum | *Methylacidiphilum sp004421185* | 1847730 | *Methylacidiphilum sp. Yel* | https://www.mdpi.com/2076-2607/10/1/142 |
| GCA_017310505.1 | Thermophiles | Acidophiles | 2254698 | Bacteria | Verrucomicrobiota | Verrucomicrobiae | Methylacidiphilales | Methylacidiphilaceae | Methylacidiphilum | *Methylacidiphilum sp004421255* | 1847729 | *Methylacidiphilum sp. Phi* | https://biotechnologyforbiofuels.biomedcentral.com/articles/10.1186/s13068-022-02105-1#Bib1 |
| GCA_000968535.1 | | Alkaliphiles | 4796711 | Bacteria | Proteobacteria | Gammaproteobacteria | Methylococcales | Methylomonadaceae | Methylotuvimicrobium | *Methylotuvimicrobium alcaliphilum* | 1091494 | *Methylotuvimicrobium alcaliphilum 20Z* | https://doi.org/10.1007/s00792-021-01228-x |
| GCA_004216855.1 | | Alkaliphiles | 2687802 | Bacteria | Actinobacteriota | Actinomycetia | Actinomycetales | Microbacteriaceae | Microcella | *Microcella alkaliphila* | 279828 | *Microcella alkaliphila* | https://doi.org/10.1099/ijs.0.64320-0 |
| GCA_002355395.1 | | Alkaliphiles | 2702837 | Bacteria | Actinobacteriota | Actinomycetia | Actinomycetales | Microbacteriaceae | Microcella | *Microcella alkaliphila_A* | 279828 | *Microcella alkaliphila* | https://doi.org/10.1099/ijs.0.64320-0 |
| GCA_004216575.1 | | Alkaliphiles | 2544518 | Bacteria | Actinobacteriota | Actinomycetia | Actinomycetales | Microbacteriaceae | Microcella | *Microcella putealis* | 337005 | *Microcella putealis* | https://doi.org/10.1016/j.syapm.2005.03.004 |
| GCA_009735625.1 | Thermophiles | | 3559563 | Bacteria | Firmicutes_B | Moorellia | Moorellales | Moorellaceae | Moorella | *Moorella glycerini* | 55779 | *Moorella glycerini* | https://doi.org/10.1099/00207713-47-4-969 |
| GCA_002995755.1 | Thermophiles | | 2628568 | Bacteria | Firmicutes_B | Moorellia | Moorellales | Moorellaceae | Moorella | *Moorella humiferrea* | 676965 | *Moorella humiferrea* | https://doi.org/10.1099/ijs.0.029009-0 |
| GCA_001594015.1 | Thermophiles | | 2999839 | Bacteria | Firmicutes_B | Moorellia | Moorellales | Moorellaceae | Moorella | *Moorella mulderi* | 1122241 | *Moorella mulderi DSM 14980* | https://doi.org/10.1007/s00203-003-0523-x |
| GCA_002995805.1 | Thermophiles | | 3328173 | Bacteria | Firmicutes_B | Moorellia | Moorellales | Moorellaceae | Moorella | *Moorella stamsii* | 1266720 | *Moorella stamsii* | https://doi.org/10.1099/ijs.0.050369-0 |
| GCA_001267405.1 | Thermophiles | | 2527564 | Bacteria | Firmicutes_B | Moorellia | Moorellales | Moorellaceae | Moorella | *Moorella thermoacetica* | 1325331 | *Moorella thermoacetica Y72* | https://doi.org/10.1016/j.resmic.2004.10.002 |
| GCA_000276805.1 | Psychrophiles | | 4889582 | Bacteria | Proteobacteria | Gammaproteobacteria | Enterobacterales | Moritellaceae | Moritella | *Moritella dasanensis* | 1201293 | *Moritella dasanensis ArB 0140* | https://doi.org/10.1099/ijs.0.65501-0 |
| GCA_008931805.1 | Psychrophiles | | 4760425 | Bacteria | Proteobacteria | Gammaproteobacteria | Enterobacterales | Moritellaceae | Moritella | *Moritella marina* | 1202962 | *Moritella marina ATCC 15381* | https://doi.org/10.1128/jb.01382-12 |
| GCA_000953735.1 | Psychrophiles | | 5093989 | Bacteria | Proteobacteria | Gammaproteobacteria | Enterobacterales | Moritellaceae | Moritella | *Moritella viscosa* | 80854 | *Moritella viscosa* | https://doi.org/10.1016/j.carres.2014.10.007 |
| GCA_900465055.1 | Psychrophiles | | 4433651 | Bacteria | Proteobacteria | Gammaproteobacteria | Enterobacterales | Moritellaceae | Moritella | *Moritella yayanosii* | 69539 | *Moritella yayanosii* | https://www.microbiologyresearch.org/content/journal/mgen/10.1099/mgen.0.000591 |
| GCA_002156705.1 | | Alkaliphiles | 3930546 | Archaea | Halobacteriota | Halobacteria | Halobacteriales | Natrialbaceae | Natrarchaeobaculum | *Natrarchaeobaculum aegyptiacum* | 745377 | *Natrarchaeobaculum aegyptiacum* | https://doi.org/10.1099/ijsem.0.004186 |
| GCA_003430825.1 | | Alkaliphiles | 3789323 | Archaea | Halobacteriota | Halobacteria | Halobacteriales | Natrialbaceae | Natrarchaeobaculum | *Natrarchaeobaculum sulfurireducens* | 2044521 | *Natrarchaeobaculum sulfurireducens* | https://doi.org/10.1099/ijsem.0.004186 |
| GCA_003841505.1 | | Alkaliphiles | 4566486 | Archaea | Halobacteriota | Halobacteria | Halobacteriales | Natrialbaceae | Natrarchaeobius | *Natrarchaeobius chitinivorans* | 1679083 | *Natrarchaeobius chitinivorans* | https://doi.org/10.1016/j.syapm.2019.01.001 |
| GCA_003841465.1 | | Alkaliphiles | 4614480 | Archaea | Halobacteriota | Halobacteria | Halobacteriales | Natrialbaceae | Natrarchaeobius | *Natrarchaeobius chitinivorans_A* | 1679083 | *Natrarchaeobius chitinivorans* | https://doi.org/10.1016/j.syapm.2019.01.001 |
| GCA_008245225.1 | | Alkaliphiles | 4201486 | Archaea | Halobacteriota | Halobacteria | Halobacteriales | Natrialbaceae | Natrarchaeobius | *Natrarchaeobius swarupiae* | 2448032 | *Natrialba swarupiae* | https://doi.org/10.1099/ijsem.0.003986 |
| GCA_000337555.1 | Mesophiles | | 4404175 | Archaea | Halobacteriota | Halobacteria | Halobacteriales | Natrialbaceae | Natrialba | *Natrialba asiatica* | 29540 | *Natrialba asiatica DSM 12278* | https://www.microbiologyresearch.org/content/journal/ijsem/10.1099/00207713-51-3-1133 |
| GCA_000337135.1 | Mesophiles | Alkaliphiles | 4309274 | Archaea | Halobacteriota | Halobacteria | Halobacteriales | Natrialbaceae | Natrialba | *Natrialba chahannaoensis* | 1227492 | *Natrialba chahannaoensis JCM 10990* | https://doi.org/10.1099/00207713-51-5-1693 |
| GCA_000337575.1 | Mesophiles | Alkaliphiles | 4159606 | Archaea | Halobacteriota | Halobacteria | Halobacteriales | Natrialbaceae | Natrialba | *Natrialba hulunbeirensis* | 1227493 | *Natrialba hulunbeirensis JCM 10989* | https://doi.org/10.1099/00207713-51-5-1693 |
| GCA_000025625.1 | | Alkaliphiles | 4443643 | Archaea | Halobacteriota | Halobacteria | Halobacteriales | Natrialbaceae | Natrialba | *Natrialba magadii* | 547559 | *Natrialba magadii ATCC 43099* | https://doi.org/10.1099/00207713-47-3-853 |
| GCA_004217335.1 | Thermophiles | | 4256545 | Archaea | Halobacteriota | Halobacteria | Halobacteriales | Natrialbaceae | Natrinema | *Natrinema hispanicum* | 392421 | *Haloterrigena hispanica* | https://doi.org/10.1099/ijs.0.64895-0 |
| GCA_900111485.1 | Thermophiles | | 3963480 | Archaea | Halobacteriota | Halobacteria | Halobacteriales | Natrialbaceae | Natrinema | *Natrinema hispanicum_A* | 392421 | *Haloterrigena hispanica* | https://doi.org/10.1099/ijs.0.64895-0 |
| GCA_000337475.1 | Thermophiles | | 3522035 | Archaea | Halobacteriota | Halobacteria | Halobacteriales | Natrialbaceae | Natrinema | *Natrinema limicola* | 1230457 | *Haloterrigena limicola* | https://www.microbiologyresearch.org/content/journal/ijsem/10.1099/ijs.0.64372-0 |
| GCA_000690595.2 | Thermophiles | | 3794337 | Archaea | Halobacteriota | Halobacteria | Halobacteriales | Natrialbaceae | Natrinema | *Natrinema mahii* | 1416969 | *Haloterrigena mahii* | https://doi.org/10.1099/ijsem.0.001811 |
| GCA_001953745.1 | Thermophiles | | 3980616 | Archaea | Halobacteriota | Halobacteria | Halobacteriales | Natrialbaceae | Natrinema | *Natrinema saccharevitans* | 301967 | *Haloterrigena saccharevitans* | https://doi.org/10.1099/ijs.0.63761-0 |
| GCA_900110865.1 | Mesophiles | | 4857017 | Archaea | Halobacteriota | Halobacteria | Halobacteriales | Natrialbaceae | Natrinema | *Natrinema salaciae* | 1186196 | *Natrinema salaciae* | https://doi.org/10.1016/j.syapm.2012.06.005 |
| GCA_900110455.1 | Mesophiles | | 4272165 | Archaea | Halobacteriota | Halobacteria | Halobacteriales | Natrialbaceae | Natrinema | *Natrinema salifodinae* | 1202768 | *Halopiger salifodinae* | https://doi.org/10.1099/ijs.0.050971-0 |
| GCA_002572525.1 | Thermophiles | | 5058058 | Archaea | Halobacteriota | Halobacteria | Halobacteriales | Natrialbaceae | Natrinema | *Natrinema sp002572525* | 1608465 | *Natrinema sp. CBA1119* | https://www.nature.com/articles/s41598-018-25887-7 |
| GCA_900215575.1 | Mesophiles | | 3164179 | Archaea | Halobacteriota | Halobacteria | Halobacteriales | Natronoarchaeaceae | Natronoarchaeum | *Natronoarchaeum philippinense* | 558529 | *Natronoarchaeum philippinense* | https://doi.org/10.1099/ijs.0.042549-0 |
| GCA_000230715.3 | | Alkaliphiles | 3788356 | Archaea | Halobacteriota | Halobacteria | Halobacteriales | Natrialbaceae | Natronobacterium | *Natronobacterium gregoryi* | 797304 | *Natronobacterium gregoryi SP2* | https://doi.org/10.1099/00207713-47-3-853 |
| GCA_900104065.1 | Mesophiles | Alkaliphiles | 4009868 | Archaea | Halobacteriota | Halobacteria | Halobacteriales | Natrialbaceae | Natronobacterium | *Natronobacterium texcoconense* | 1095778 | *Natronobacterium texcoconense* | https://doi.org/10.1099/ijs.0.053629-0 |
| GCA_000337675.1 | | Alkaliphiles | 4416525 | Archaea | Halobacteriota | Halobacteria | Halobacteriales | Natrialbaceae | Natronococcus | *Natronococcus amylolyticus* | 1227497 | *Natronococcus amylolyticus DSM 10524* | https://doi.org/10.1099/00207713-45-4-762 |
| GCA_000337695.1 | Mesophiles | | 4496185 | Archaea | Halobacteriota | Halobacteria | Halobacteriales | Natrialbaceae | Natronococcus | *Natronococcus jeotgali* | 1227498 | *Natronococcus jeotgali* | https://doi.org/10.1099/ijs.0.65120-0 |
| GCA_000328685.1 | | Alkaliphiles | 4314118 | Archaea | Halobacteriota | Halobacteria | Halobacteriales | Natrialbaceae | Natronococcus | *Natronococcus occultus* | 694430 | *Natronococcus occultus SP4* | https://doi.org/10.1002/1521-4028(200112)41:6%3C375::aid-jobm375%3E3.0.co;2-0 |
| GCA_008122205.1 | | Alkaliphiles | 5316806 | Archaea | Halobacteriota | Halobacteria | Halobacteriales | Natrialbaceae | Natronococcus | *Natronococcus sp008122205* | 2055836 | *Natronococcus pandeyae* | https://doi.org/10.1007/s00284-021-02740-1 |
| GCA_000026045.1 | | Alkaliphiles | 2749696 | Archaea | Halobacteriota | Halobacteria | Halobacteriales | Haloarculaceae | Natronomonas | *Natronomonas pharaonis* | 348780 | *Natronomonas pharaonis DSM 2160* | https://doi.org/10.1099/00207713-47-3-853 |
| GCA_009392895.1 | | Alkaliphiles | 4348483 | Archaea | Halobacteriota | Halobacteria | Halobacteriales | Natrialbaceae | Natronorubrum | *Natronorubrum aibiense* | 348826 | *Natronorubrum aibiense* | https://doi.org/10.1099/ijs.0.64222-0 |
| GCA_001971705.1 | Mesophiles | Alkaliphiles | 3835796 | Archaea | Halobacteriota | Halobacteria | Halobacteriales | Natrialbaceae | Natronorubrum | *Natronorubrum daqingense* | 588898 | *Haloterrigena daqingensis* | https://doi.org/10.1099/ijs.0.013995-0 |
| GCA_900108095.1 | | Alkaliphiles | 3782545 | Archaea | Halobacteriota | Halobacteria | Halobacteriales | Natrialbaceae | Natronorubrum | *Natronorubrum sediminis* | 640943 | *Natronorubrum sediminis* | https://doi.org/10.1099/ijs.0.015602-0 |
| GCA_000337735.1 | Thermophiles | Alkaliphiles | 3460288 | Archaea | Halobacteriota | Halobacteria | Halobacteriales | Natrialbaceae | Natronorubrum | *Natronorubrum sulfidifaciens* | 1230460 | *Natronorubrum sulfidifaciens* | https://doi.org/10.1099/ijs.0.64651-0 |
| GCA_000383975.1 | | Alkaliphiles | 4934841 | Archaea | Halobacteriota | Halobacteria | Halobacteriales | Natrialbaceae | Natronorubrum | *Natronorubrum tibetense* | 1114856 | *Natronorubrum tibetense GA33* | https://doi.org/10.1099/00207713-49-1-261 |
| GCA_000220175.2 | Mesophiles | | 1607695 | Archaea | Thermoproteota | Nitrososphaeria | Nitrososphaerales | Nitrosopumilaceae | Nitrosarchaeum | *Nitrosarchaeum koreense* | 1088740 | *Nitrosarchaeum koreense MY1* | https://www.microbiologyresearch.org/content/journal/ijsem/10.1099/ijsem.0.002926 |
| GCA_000956175.1 | Mesophiles | | 1803090 | Archaea | Thermoproteota | Nitrososphaeria | Nitrososphaerales | Nitrosopumilaceae | Nitrosopumilus | *Nitrosopumilus adriaticus* | 1580092 | *Nitrosopumilus adriaticus* | https://doi.org/10.1099/ijsem.0.003360 |
| GCA_000018465.1 | Mesophiles | | 1645259 | Archaea | Thermoproteota | Nitrososphaeria | Nitrososphaerales | Nitrosopumilaceae | Nitrosopumilus | *Nitrosopumilus maritimus* | 436308 | *Nitrosopumilus maritimus SCM1* | 10.1099/ijsem.0.002416 |
| GCA_000875775.1 | Mesophiles | | 1713078 | Archaea | Thermoproteota | Nitrososphaeria | Nitrososphaerales | Nitrosopumilaceae | Nitrosopumilus | *Nitrosopumilus piranensis* | 1582439 | *Nitrosopumilus piranensis* | https://doi.org/10.1099/ijsem.0.003360 |
| GCA_000698785.1 | Mesophiles | | 2527938 | Archaea | Thermoproteota | Nitrososphaeria | Nitrososphaerales | Nitrososphaeraceae | Nitrososphaera | *Nitrososphaera viennensis* | 926571 | *Nitrososphaera viennensis EN76* | https://doi.org/10.1099/ijs.0.063172-0 |
| GCA_000967895.1 | Psychrophiles | | 4406383 | Bacteria | Proteobacteria | Gammaproteobacteria | Pseudomonadales | DSM-6294 | Oleispira | *Oleispira antarctica* | 698738 | *Oleispira antarctica RB-8* | https://www.microbiologyresearch.org/content/journal/ijsem/10.1099/ijs.0.02366-0 |
| GCA_000966265.1 | Hyperthermophiles | | 2206431 | Archaea | Methanobacteriota_B | Thermococci | Thermococcales | Thermococcaceae | Palaeococcus | *Palaeococcus ferrophilus* | 588319 | *Palaeococcus ferrophilus DSM 13482* | https://doi.org/10.1099/00207713-50-2-489 |
| GCA_000725425.1 | Thermophiles | | 1859370 | Archaea | Methanobacteriota_B | Thermococci | Thermococcales | Thermococcaceae | Palaeococcus | *Palaeococcus pacificus* | 1343739 | *Palaeococcus pacificus DY20341* | https://doi.org/10.1099/ijs.0.044487-0 |
| GCA_900111865.1 | Thermophiles | | 3448881 | Bacteria | Firmicutes | Bacilli | Bacillales | Anoxybacillaceae | Parageobacillus | *Parageobacillus thermantarcticus* | 186116 | *Parageobacillus thermantarcticus* | https://bacdive.dsmz.de/strain/1438 |
| GCA_001295365.1 | Thermophiles | | 3873116 | Bacteria | Firmicutes | Bacilli | Bacillales | Anoxybacillaceae | Parageobacillus | *Parageobacillus thermoglucosidasius* | 1136178 | *Parageobacillus thermoglucosidasius TNO-09.020* | https://bacdive.dsmz.de/strain/1430 |
| GCA_001029445.1 | | Alkaliphiles | 5078735 | Bacteria | Proteobacteria | Gammaproteobacteria | Enterobacterales | Vibrionaceae | Photobacterium | *Photobacterium aquae* | 1195763 | *Photobacterium aquae* | https://doi.org/10.1099/ijs.0.055020-0 |
| GCA_002954455.1 | Psychrophiles | | 4525475 | Bacteria | Proteobacteria | Gammaproteobacteria | Enterobacterales | Vibrionaceae | Photobacterium | *Photobacterium aquimaris* | 512643 | *Photobacterium aquimaris* | https://www.microbiologyresearch.org/content/journal/ijsem/10.1099/ijs.0.004309-0 |
| GCA_002849605.1 | Psychrophiles | | 4559543 | Bacteria | Proteobacteria | Gammaproteobacteria | Enterobacterales | Vibrionaceae | Photobacterium | *Photobacterium carnosum* | 2023717 | *Photobacterium carnosum* | https://www.sciencedirect.com/science/article/pii/S0723202017301686?via%3Dihub |
| GCA_003026815.1 | Psychrophiles | | 4914916 | Bacteria | Proteobacteria | Gammaproteobacteria | Enterobacterales | Vibrionaceae | Photobacterium | *Photobacterium damselae* | 38293 | *Photobacterium damselae* | https://www.frontiersin.org/articles/10.3389/fmicb.2013.00283/full |
| GCA_003025615.1 | Psychrophiles | | 6446463 | Bacteria | Proteobacteria | Gammaproteobacteria | Enterobacterales | Vibrionaceae | Photobacterium | *Photobacterium frigidiphilum* | 264736 | *Photobacterium frigidiphilum* | https://doi.org/10.1099/ijs.0.63338-0 |
| GCA_003025595.1 | | Alkaliphiles | 5533005 | Bacteria | Proteobacteria | Gammaproteobacteria | Enterobacterales | Vibrionaceae | Photobacterium | *Photobacterium ganghwense* | 320778 | *Photobacterium ganghwense* | https://www.microbiologyresearch.org/content/journal/ijsem/10.1099/ijs.0.63811-0 |

| | | | | | | | | | | | | | |
|---|---|---|---|---|---|---|---|---|---|---|---|---|---|
| GCA_000425165.1 | Psychrophiles | | 4690166 | Bacteria | Proteobacteria | Gammaproteobacteria | Enterobacterales | Vibrionaceae | Photobacterium | *Photobacterium halotolerans* | 1122959 | *Photobacterium halotolerans DSM 18316* | https://www.microbiologyresearch.org/content/journal/ijsem/10.1099/ijs.0.64099-0 |
| GCA_003026395.1 | Psychrophiles | | 4308695 | Bacteria | Proteobacteria | Gammaproteobacteria | Enterobacterales | Vibrionaceae | Photobacterium | *Photobacterium iliopiscarium* | 56192 | *Photobacterium iliopiscarium* | https://bacdive.dsmz.de/strain/17224 |
| GCA_003026355.1 | Psychrophiles | | 4732354 | Bacteria | Proteobacteria | Gammaproteobacteria | Enterobacterales | Vibrionaceae | Photobacterium | *Photobacterium kishitanii* | 318456 | *Photobacterium kishitanii* | https://bacdive.dsmz.de/strain/17235 |
| GCA_003026475.1 | Psychrophiles | | 4943313 | Bacteria | Proteobacteria | Gammaproteobacteria | Enterobacterales | Vibrionaceae | Photobacterium | *Photobacterium lipolyticum* | 266810 | *Photobacterium lipolyticum* | https://www.microbiologyresearch.org/content/journal/ijsem/10.1099/ijs.0.63215-0 |
| GCA_900166965.1 | Psychrophiles | | 4525300 | Bacteria | Proteobacteria | Gammaproteobacteria | Enterobacterales | Vibrionaceae | Photobacterium | *Photobacterium piscicola* | 1378299 | *Photobacterium piscicola* | https://www.sciencedirect.com/science/article/pii/S0723202014000770?via%3Dihub |
| GCA_003026285.1 | Psychrophiles | | 6116719 | Bacteria | Proteobacteria | Gammaproteobacteria | Enterobacterales | Vibrionaceae | Photobacterium | *Photobacterium profundum* | 314280 | *Photobacterium profundum 3TCK* | https://doi.org/10.1007/s007920050036 |
| GCA_001939735.1 | Psychrophiles | | 6484429 | Bacteria | Proteobacteria | Gammaproteobacteria | Enterobacterales | Vibrionaceae | Photobacterium | *Photobacterium proteolyticum* | 1903952 | *Photobacterium proteolyticum* | https://www.microbiologyresearch.org/content/journal/ijsem/10.1099/ijsem.0.001873 |
| GCA_001077885.1 | Psychrophiles | | 5523519 | Bacteria | Proteobacteria | Gammaproteobacteria | Enterobacterales | Vibrionaceae | Photobacterium | *Photobacterium swingsii* | 680026 | *Photobacterium swingsii* | https://www.microbiologyresearch.org/content/journal/ijsem/10.1099/ijs.0.019687-0 |
| GCA_900166975.1 | Psychrophiles | | 4420947 | Bacteria | Proteobacteria | Gammaproteobacteria | Enterobacterales | Vibrionaceae | Photobacterium | *Photobacterium toruni* | 1935446 | *Photobacterium toruni* | https://www.microbiologyresearch.org/content/journal/ijsem/10.1099/ijsem.0.002325 |
| GCA_900176435.1 | Thermophiles | Acidophiles | 1534155 | Archaea | Thermoplasmatota | Thermoplasmata | Thermoplasmatales | Thermoplasmataceae | Picrophilus | *Picrophilus oshimae* | 1122961 | *Picrophilus oshimae DSM 9789* | https://doi.org/10.1023/a:1020525525490 |
| GCA_003719725.1 | | Alkaliphiles | 3335228 | Bacteria | Firmicutes | Bacilli | Bacillales_A | Planococcaceae | Planococcus | *Planococcus salinus* | 1848460 | *Planococcus salinus* | https://doi.org/10.1099/ijsem.0.002548 |
| GCA_001640115.1 | Psychrophiles | | 3942702 | Bacteria | Bacteroidota | Bacteroidia | Flavobacteriales | Flavobacteriaceae | Polaribacter | *Polaribacter atrinae* | 1333662 | *Polaribacter atrinae* | https://www.microbiologyresearch.org/content/journal/ijsem/10.1099/ijs.0.060889-0 |
| GCA_002954665.1 | Psychrophiles | | 4064562 | Bacteria | Bacteroidota | Bacteroidia | Flavobacteriales | Flavobacteriaceae | Polaribacter | *Polaribacter glomeratus* | 102 | *Polaribacter glomeratus* | https://bacdive.dsmz.de/strain/136694 |
| GCA_014784055.1 | Psychrophiles | | 3800626 | Bacteria | Bacteroidota | Bacteroidia | Flavobacteriales | Flavobacteriaceae | Polaribacter | *Polaribacter haliotis* | 1888915 | *Polaribacter haliotis* | https://pubmed.ncbi.nlm.nih.gov/27902190/ |
| GCA_000153225.1 | Psychrophiles | | 2763458 | Bacteria | Bacteroidota | Bacteroidia | Flavobacteriales | Flavobacteriaceae | Polaribacter | *Polaribacter irgensii* | 313594 | *Polaribacter irgensii 23-P* | https://www.microbiologyresearch.org/content/journal/ijsem/10.1099/00207713-48-1-223 |
| GCA_002954685.1 | Psychrophiles | | 3904103 | Bacteria | Bacteroidota | Bacteroidia | Flavobacteriales | Flavobacteriaceae | Polaribacter | *Polaribacter porphyrae* | 1137780 | *Polaribacter porphyrae* | https://www.microbiologyresearch.org/content/journal/ijsem/10.1099/ijs.0.041434-0 |
| GCA_001975665.1 | Psychrophiles | Alkaliphiles | 4125014 | Bacteria | Bacteroidota | Bacteroidia | Flavobacteriales | Flavobacteriaceae | Polaribacter | *Polaribacter reichenbachii* | 996801 | *Polaribacter reichenbachii* | https://link.springer.com/article/10.1007/s00284-012-0200-x |
| GCA_001761365.1 | Psychrophiles | Alkaliphiles | 3809314 | Bacteria | Bacteroidota | Bacteroidia | Flavobacteriales | Flavobacteriaceae | Polaribacter | *Polaribacter vadi* | 1774273 | *Polaribacter vadi* | https://doi.org/10.1099/ijsem.0.001591 |
| GCA_018861005.1 | Psychrophiles | Alkaliphiles | 3934366 | Bacteria | Bacteroidota | Bacteroidia | Flavobacteriales | Flavobacteriaceae | Polaribacter | *Polaribacter vadi_A* | 1774273 | *Polaribacter vadi* | https://doi.org/10.1099/ijsem.0.001591 |
| GCA_000709345.1 | Psychrophiles | | 5284042 | Bacteria | Proteobacteria | Gammaproteobacteria | Burkholderiales | Burkholderiaceae | Polaromonas | *Polaromonas glacialis* | 866564 | *Polaromonas glacialis* | https://doi.org/10.1099/ijs.0.037556-0 |
| GCA_001598235.1 | Psychrophiles | | 5134650 | Bacteria | Proteobacteria | Gammaproteobacteria | Burkholderiales | Burkholderiaceae | Polaromonas | *Polaromonas jejuensis* | 1321608 | *Polaromonas jejuensis NBRC 106434* | https://www.microbiologyresearch.org/content/journal/ijsem/10.1099/ijs.0.65529-0 |
| GCA_000015505.1 | Psychrophiles | | 5366143 | Bacteria | Proteobacteria | Gammaproteobacteria | Burkholderiales | Burkholderiaceae | Polaromonas | *Polaromonas naphthalenivorans* | 365044 | *Polaromonas naphthalenivorans CJ2* | https://pubmed.ncbi.nlm.nih.gov/14742464/ |
| GCA_000507185.2 | Hyperthermophiles | | 6150048 | Bacteria | Proteobacteria | Gammaproteobacteria | Pseudomonadales | Pseudomonadaceae | Pseudomonas_E | *Pseudomonas_E syringae* | 52001 | *Stetteria hydrogenophila* | https://bacdive.dsmz.de/strain/4193 |
| GCA_000217815.1 | Thermophiles | | 2039943 | Bacteria | Thermotogota | Thermotogae | Thermotogales | DSM-5069 | Pseudothermotoga | *Pseudothermotoga thermarum* | 688269 | *Pseudothermotoga thermarum DSM 5069* | https://link.springer.com/article/10.1007/s10482-013-0062-7 |
| GCA_000828655.1 | Thermophiles | | 2014912 | Bacteria | Thermotogota | Thermotogae | Thermotogales | DSM-5069 | Pseudothermotoga_A | *Pseudothermotoga_A caldifontis* | 1408159 | *Thermotoga caldifontis AZM44c09* | https://doi.org/10.1099/ijs.0.060137-0 |
| GCA_000816145.1 | Thermophiles | | 2165416 | Bacteria | Thermotogota | Thermotogae | Thermotogales | DSM-5069 | Pseudothermotoga_A | *Pseudothermotoga_A hypogea* | 1123384 | *Pseudothermotoga hypogea DSM 11164 = NBRC 106472* | https://link.springer.com/article/10.1007/s10482-013-0062-7 |
| GCA_000504085.1 | Thermophiles | | 2169860 | Bacteria | Thermotogota | Thermotogae | Thermotogales | DSM-5069 | Pseudothermotoga_B | *Pseudothermotoga_B elfii* | 416591 | *Pseudothermotoga lettingae TMO* | https://link.springer.com/article/10.1007/s10482-013-0062-7 |
| GCA_000828675.1 | Thermophiles | | 2187612 | Bacteria | Thermotogota | Thermotogae | Thermotogales | DSM-5069 | Pseudothermotoga_B | *Pseudothermotoga_B profunda* | 1408160 | *Thermotoga profunda AZM34c06* | https://doi.org/10.1099/ijs.0.060137-0 |
| GCA_001606025.1 | Psychrophiles | | 3349444 | Bacteria | Proteobacteria | Gammaproteobacteria | Pseudomonadales | Moraxellaceae | Psychrobacter | *Psychrobacter alimentarius_A* | 261164 | *Psychrobacter alimentarius* | https://pubmed.ncbi.nlm.nih.gov/15653872/ |
| GCA_000471625.1 | Psychrophiles | | 3216409 | Bacteria | Proteobacteria | Gammaproteobacteria | Pseudomonadales | Moraxellaceae | Psychrobacter | *Psychrobacter aquaticus* | 1354303 | *Psychrobacter aquaticus CMS 56* | https://doi.org/10.1099/ijs.0.03030-0 |
| GCA_000012305.1 | Psychrophiles | | 2650701 | Bacteria | Proteobacteria | Gammaproteobacteria | Pseudomonadales | Moraxellaceae | Psychrobacter | *Psychrobacter arcticus* | 259536 | *Psychrobacter arcticus 273-4* | https://doi.org/10.1099/ijs.0.64043-0 |
| GCA_016107535.1 | Psychrophiles | | 3242921 | Bacteria | Proteobacteria | Gammaproteobacteria | Pseudomonadales | Moraxellaceae | Psychrobacter | *Psychrobacter cibarius* | 282669 | *Psychrobacter cibarius* | https://doi.org/10.1099/ijs.0.63398-0 |
| GCA_000013905.1 | Psychrophiles | | 3101097 | Bacteria | Proteobacteria | Gammaproteobacteria | Pseudomonadales | Moraxellaceae | Psychrobacter | *Psychrobacter cryohalolentis* | 330922 | *Psychrobacter cryohalolentis* | https://doi.org/10.1099/ijs.0.64043-0 |
| GCA_003217155.1 | Psychrophiles | | 3499844 | Bacteria | Proteobacteria | Gammaproteobacteria | Pseudomonadales | Moraxellaceae | Psychrobacter | *Psychrobacter fozii* | 198480 | *Psychrobacter fozii* | https://doi.org/10.1099/ijs.0.02457-0 |
| GCA_007997305.1 | Psychrophiles | | 2846672 | Bacteria | Proteobacteria | Gammaproteobacteria | Pseudomonadales | Moraxellaceae | Psychrobacter | *Psychrobacter frigidicola* | 45611 | *Psychrobacter frigidicola* | https://doi.org/10.1128/jb.01137-08 |
| GCA_001411745.2 | Psychrophiles | | 3490652 | Bacteria | Proteobacteria | Gammaproteobacteria | Pseudomonadales | Moraxellaceae | Psychrobacter | *Psychrobacter glacincola_A* | 56810 | *Psychrobacter glacincola* | https://doi.org/10.1099/ijs.0.02457-0 |
| GCA_904846215.1 | Psychrophiles | | 3247549 | Bacteria | Proteobacteria | Gammaproteobacteria | Pseudomonadales | Moraxellaceae | Psychrobacter | *Psychrobacter glacincola_B* | 56810 | *Psychrobacter glacincola* | https://doi.org/10.1099/ijs.0.02457-0 |
| GCA_904846075.1 | Psychrophiles | | 3304715 | Bacteria | Proteobacteria | Gammaproteobacteria | Pseudomonadales | Moraxellaceae | Psychrobacter | *Psychrobacter glacincola_C* | 56810 | *Psychrobacter glacincola* | https://doi.org/10.1099/ijs.0.02457-0 |
| GCA_003148585.1 | Psychrophiles | | 3243066 | Bacteria | Proteobacteria | Gammaproteobacteria | Pseudomonadales | Moraxellaceae | Psychrobacter | *Psychrobacter immobilis* | 498 | *Psychrobacter immobilis* | https://bacdive.dsmz.de/strain/8176 |
| GCA_904846265.1 | Psychrophiles | | 3444614 | Bacteria | Proteobacteria | Gammaproteobacteria | Pseudomonadales | Moraxellaceae | Psychrobacter | *Psychrobacter immobilis_A* | 498 | *Psychrobacter immobilis* | https://bacdive.dsmz.de/strain/8176 |
| GCA_904846185.1 | Psychrophiles | | 3473556 | Bacteria | Proteobacteria | Gammaproteobacteria | Pseudomonadales | Moraxellaceae | Psychrobacter | *Psychrobacter immobilis_B* | 498 | *Psychrobacter immobilis* | https://bacdive.dsmz.de/strain/8176 |
| GCA_904846235.1 | Psychrophiles | | 3223472 | Bacteria | Proteobacteria | Gammaproteobacteria | Pseudomonadales | Moraxellaceae | Psychrobacter | *Psychrobacter immobilis_C* | 498 | *Psychrobacter immobilis* | https://bacdive.dsmz.de/strain/8176 |
| GCA_904846285.1 | Psychrophiles | | 3229758 | Bacteria | Proteobacteria | Gammaproteobacteria | Pseudomonadales | Moraxellaceae | Psychrobacter | *Psychrobacter immobilis_D* | 498 | *Psychrobacter immobilis* | https://bacdive.dsmz.de/strain/8176 |
| GCA_904846175.1 | Psychrophiles | | 3123162 | Bacteria | Proteobacteria | Gammaproteobacteria | Pseudomonadales | Moraxellaceae | Psychrobacter | *Psychrobacter immobilis_E* | 498 | *Psychrobacter immobilis* | https://bacdive.dsmz.de/strain/8176 |
| GCA_904846225.1 | Psychrophiles | | 3336711 | Bacteria | Proteobacteria | Gammaproteobacteria | Pseudomonadales | Moraxellaceae | Psychrobacter | *Psychrobacter immobilis_F* | 498 | *Psychrobacter immobilis* | https://bacdive.dsmz.de/strain/8176 |
| GCA_904846145.1 | Psychrophiles | | 3247678 | Bacteria | Proteobacteria | Gammaproteobacteria | Pseudomonadales | Moraxellaceae | Psychrobacter | *Psychrobacter immobilis_G* | 498 | *Psychrobacter immobilis* | https://bacdive.dsmz.de/strain/8176 |
| GCA_000382145.1 | Psychrophiles | | 3176011 | Bacteria | Proteobacteria | Gammaproteobacteria | Pseudomonadales | Moraxellaceae | Psychrobacter | *Psychrobacter lutiphocae* | 1123033 | *Psychrobacter lutiphocae DSM 21542* | https://doi.org/10.1099/ijs.0.008706-0 |
| GCA_900101915.1 | Psychrophiles | | 3062581 | Bacteria | Proteobacteria | Gammaproteobacteria | Pseudomonadales | Moraxellaceae | Psychrobacter | *Psychrobacter pacificensis* | 112002 | *Psychrobacter pacificensis* | https://pubmed.ncbi.nlm.nih.gov/23868329/ |
| GCA_900162825.1 | Psychrophiles | | 2820896 | Bacteria | Proteobacteria | Gammaproteobacteria | Pseudomonadales | Moraxellaceae | Psychrobacter | *Psychrobacter piechaudii* | 1945521 | *Psychrobacter piechaudii* | https://www.microbiologyresearch.org/content/journal/ijsem/10.1099/ijsem.0.002065 |
| GCA_001444505.1 | Psychrophiles | | 3089314 | Bacteria | Proteobacteria | Gammaproteobacteria | Pseudomonadales | Moraxellaceae | Psychrobacter | *Psychrobacter piscatorii* | 554343 | *Psychrobacter piscatorii* | https://doi.org/10.1099/ijs.0.010959-0 |
| GCA_904846415.1 | Psychrophiles | | 3514701 | Bacteria | Proteobacteria | Gammaproteobacteria | Pseudomonadales | Moraxellaceae | Psychrobacter | *Psychrobacter piscatorii_A* | 554343 | *Psychrobacter piscatorii* | https://doi.org/10.1099/ijs.0.010959-0 |
| GCA_003350005.1 | Psychrophiles | | 3031855 | Bacteria | Proteobacteria | Gammaproteobacteria | Pseudomonadales | Moraxellaceae | Psychrobacter | *Psychrobacter proteolyticus_A* | 147825 | *Psychrobacter proteolyticus* | https://doi.org/10.1078/0723-2020-00006 |
| GCA_001298525.1 | Psychrophiles | | 2931580 | Bacteria | Proteobacteria | Gammaproteobacteria | Pseudomonadales | Moraxellaceae | Psychrobacter | *Psychrobacter urativorans_A* | 45610 | *Psychrobacter urativorans* | https://bacdive.dsmz.de/strain/8178 |
| GCA_904846675.1 | Psychrophiles | | 2734405 | Bacteria | Proteobacteria | Gammaproteobacteria | Pseudomonadales | Moraxellaceae | Psychrobacter | *Psychrobacter urativorans_B* | 45610 | *Psychrobacter urativorans* | https://bacdive.dsmz.de/strain/8178 |
| GCA_904846695.1 | Psychrophiles | | 3458123 | Bacteria | Proteobacteria | Gammaproteobacteria | Pseudomonadales | Moraxellaceae | Psychrobacter | *Psychrobacter urativorans_C* | 45610 | *Psychrobacter urativorans* | https://bacdive.dsmz.de/strain/8178 |
| GCA_000153485.2 | Psychrophiles | | 4321832 | Bacteria | Bacteroidota | Bacteroidia | Flavobacteriales | Flavobacteriaceae | Psychroflexus | *Psychroflexus torquis* | 313595 | *Psychroflexus torquis ATCC 700755* | https://doi.org/10.1099/00221287-144-6-1601 |
| GCA_000428725.1 | Psychrophiles | | 5534630 | Bacteria | Proteobacteria | Gammaproteobacteria | Enterobacterales | Psychromonadaceae | Psychromonas | *Psychromonas aquimarina* | 1278312 | *Psychromonas aquimarina ATCC BAA-1526* | https://doi.org/10.1099/ijs.0.65744-0 |
| GCA_000482725.1 | Psychrophiles | | 4745897 | Bacteria | Proteobacteria | Gammaproteobacteria | Enterobacterales | Psychromonadaceae | Psychromonas | *Psychromonas arctica* | 1123036 | *Psychromonas arctica DSM 14288* | https://doi.org/10.1099/ijs.0.02182-0 |
| GCA_000420245.1 | Psychrophiles | | 3979980 | Bacteria | Proteobacteria | Gammaproteobacteria | Enterobacterales | Psychromonadaceae | Psychromonas | *Psychromonas hadalis* | 1278302 | *Psychromonas hadalis ATCC BAA-638* | https://doi.org/10.1099/ijs.0.64933-0 |
| GCA_000015285.1 | Psychrophiles | | 4559598 | Bacteria | Proteobacteria | Gammaproteobacteria | Enterobacterales | Psychromonadaceae | Psychromonas | *Psychromonas ingrahamii* | 357804 | *Psychromonas ingrahamii 37* | https://doi.org/10.1099/ijs.0.64068-0 |

| | | | | | | | | | | | | | |
|---|---|---|---|---|---|---|---|---|---|---|---|---|---|
| GCA_000381745.1 | Psychrophiles | | 5204311 | Bacteria | Proteobacteria | Gammaproteobacteria | Enterobacterales | Psychromonadaceae | Psychromonas | *Psychromonas ossibalaenae* | 1278307 | *Psychromonas ossibalaenae ATCC BAA-1528* | https://doi.org/10.1099/ijs.0.65744-0 |
| GCA_000007225.1 | Hyperthermophiles | | 2222430 | Archaea | Thermoproteota | Thermoproteia | Thermoproteales | Thermoproteaceae | Pyrobaculum | *Pyrobaculum aerophilum* | 13773 | *Pyrobaculum aerophilum* | https://pubmed.ncbi.nlm.nih.gov/19047344/ |
| GCA_000016385.1 | Hyperthermophiles | | 2121076 | Archaea | Thermoproteota | Thermoproteia | Thermoproteales | Thermoproteaceae | Pyrobaculum | *Pyrobaculum arsenaticum* | 121277 | *Pyrobaculum arsenaticum PZ6, DSM 13514* | https://bacdive.dsmz.de/strain/17028 |
| GCA_000015805.1 | Hyperthermophiles | | 2009313 | Archaea | Thermoproteota | Thermoproteia | Thermoproteales | Thermoproteaceae | Pyrobaculum | *Pyrobaculum calidifontis* | 410359 | *Pyrobaculum calidifontis VA1* | https://doi.org/10.1155/2002/616075 |
| GCA_000234805.1 | Hyperthermophiles | | 2467972 | Archaea | Thermoproteota | Thermoproteia | Thermoproteales | Thermoproteaceae | Pyrobaculum | *Pyrobaculum ferrireducens* | 1104324 | *Pyrobaculum ferrireducens* | https://doi.org/10.1099/ijs.0.000027 |
| GCA_000015205.1 | Hyperthermophiles | | 1826402 | Archaea | Thermoproteota | Thermoproteia | Thermoproteales | Thermoproteaceae | Pyrobaculum | *Pyrobaculum islandicum* | 384616 | *Pyrobaculum islandicum DSM 4184* | https://doi.org/10.1271/bbb.120367 |
| GCA_000019805.1 | Hyperthermophiles | | 1769823 | Archaea | Thermoproteota | Thermoproteia | Thermoproteales | Thermoproteaceae | Pyrobaculum | *Pyrobaculum neutrophilum* | 444157 | *Thermoproteus neutrophilus* | https://bacdive.dsmz.de/strain/17030 |
| GCA_000247545.1 | Hyperthermophiles | | 2452920 | Archaea | Thermoproteota | Thermoproteia | Thermoproteales | Thermoproteaceae | Pyrobaculum | *Pyrobaculum oguniense* | 698757 | *Pyrobaculum oguniense TE7, DSM 13380* | https://doi.org/10.1099/00207713-51-2-303 |
| GCA_001189275.1 | Thermophiles | | 1993257 | Archaea | Thermoproteota | Thermoproteia | Thermoproteales | Thermoproteaceae | Pyrobaculum | *Pyrobaculum sp001189275* | 1227555 | *Pyrobaculum yellowstonensis sp WP30* | https://doi.org/10.1128/aem.01095-15 |
| GCA_000195935.2 | Hyperthermophiles | | 1768562 | Archaea | Methanobacteriota_B | Thermococci | Thermococcales | Thermococcaceae | Pyrococcus | *Pyrococcus abyssi* | 272844 | *Pyrococcus abyssi GE5* | https://bacdive.dsmz.de/strain/16859 |
| GCA_002214605.1 | Hyperthermophiles | | 1961979 | Archaea | Methanobacteriota_B | Thermococci | Thermococcales | Thermococcaceae | Pyrococcus | *Pyrococcus chitonophagus* | 54262 | *Pyrococcus chitonophagus DSM 10152* | https://doi.org/10.1128/aem.00319-16 |
| GCA_000007305.1 | Hyperthermophiles | | 1908256 | Archaea | Methanobacteriota_B | Thermococci | Thermococcales | Thermococcaceae | Pyrococcus | *Pyrococcus furiosus* | 186497 | *Pyrococcus furiosus* | https://bacdive.dsmz.de/strain/16854 |
| GCA_000211475.1 | Hyperthermophiles | | 1861320 | Archaea | Methanobacteriota_B | Thermococci | Thermococcales | Thermococcaceae | Pyrococcus | *Pyrococcus sp000211475* | 342949 | *Pyrococcus sp NA2* | https://doi.org/10.1128/jb.05150-11 |
| GCA_000215995.1 | Hyperthermophiles | Acidophiles | 1716818 | Archaea | Methanobacteriota_B | Thermococci | Thermococcales | Thermococcaceae | Pyrococcus | *Pyrococcus yayanosii* | 529709 | *Pyrococcus yayanosii CH1* | https://bacdive.dsmz.de/strain/161812 |
| GCA_001412615.1 | Hyperthermophiles | | 2023836 | Archaea | Thermoproteota | Thermoproteia | Sulfolobales | Pyrodictiaceae | Pyrodictium | *Pyrodictium delaneyi* | 1273541 | *Pyrodictium delaneyi* | https://www.microbiologyresearch.org/content/journal/ijsem/10.1099/ijsem.0.001201 |
| GCA_001462395.1 | Hyperthermophiles | | 1621727 | Archaea | Thermoproteota | Thermoproteia | Sulfolobales | Pyrodictiaceae | Pyrodictium | *Pyrodictium occultum* | 2309 | *Pyrodictium occultum* | https://doi.org/10.1128/jb.177.8.2164-2177.1995 |
| GCA_000223395.1 | Hyperthermophiles | Acidophiles | 1843267 | Archaea | Thermoproteota | Thermoproteia | Sulfolobales | Pyrodictiaceae | Pyrolobus | *Pyrolobus fumarii* | 694429 | *Pyrolobus fumarii 1A* | https://doi.org/10.1007/s007920050010 |
| GCA_009765975.1 | | Acidophiles | 8370829 | Bacteria | Proteobacteria | Alphaproteobacteria | Acetobacterales | Acetobacteraceae | Rhodopila | *Rhodopila sp009765975* | 1747223 | *Acidisphaera sp. S103* | https://doi.org/10.1099/00207713-50-4-1539 |
| GCA_000024845.1 | Thermophiles | | 3386737 | Bacteria | Bacteroidota | Rhodothermia | Rhodothermales | Rhodothermaceae | Rhodothermus | *Rhodothermus marinus* | 518766 | *Rhodothermus marinus DSM 4252* | https://bacdive.dsmz.de/strain/17794 |
| GCA_900142415.1 | Thermophiles | | 3139689 | Bacteria | Bacteroidota | Rhodothermia | Rhodothermales | Rhodothermaceae | Rhodothermus | *Rhodothermus profundi* | 633813 | *Rhodothermus profundi* | https://doi.org/10.1099/ijs.0.012724-0 |
| GCA_003568865.1 | Mesophiles | | 3078689 | Bacteria | Actinobacteriota | Rubrobacteria | Rubrobacterales | Rubrobacteraceae | Rubrobacter | *Rubrobacter indicoceani* | 2051957 | *Rubrobacter indicoceani SCSIO 08198* | https://www.microbiologyresearch.org/content/journal/ijsem/10.1099/ijsem.0.003018 |
| GCA_900175965.1 | | Alkaliphiles | 3398074 | Bacteria | Actinobacteriota | Rubrobacteria | Rubrobacterales | Rubrobacteraceae | Rubrobacter | *Rubrobacter radiotolerans* | 42256 | *Rubrobacter radiotolerans* | https://www.ncbi.nlm.nih.gov/pmc/articles/PMC4148983/ |
| GCA_000014185.1 | Thermophiles | Alkaliphiles | 3225748 | Bacteria | Actinobacteriota | Rubrobacteria | Rubrobacterales | Rubrobacteraceae | Rubrobacter_B | *Rubrobacter_B xylanophilus* | 266117 | *Rubrobacter xylanophilus DSM 9941* | https://bacdive.dsmz.de/strain/14036 |
| GCA_000632715.1 | Thermophiles | | 3798752 | Bacteria | Firmicutes | Bacilli | Bacillales | Anoxybacillaceae | Saccharococcus | *Saccharococcus caldoxylosilyticus* | 81408 | *Parageobacillus caldoxylosilyticus* | https://doi.org/10.1016/j.dib.2021.107764 |
| GCA_000022485.1 | Hyperthermophiles | Acidophiles | 2854410 | Archaea | Thermoproteota | Thermoproteia | Sulfolobales | Sulfolobaceae | Saccharolobus | *Saccharolobus islandicus* | 425944 | *Sulfolobus islandicus L.D.8.5* | https://doi.org/10.1007/978-1-0716-2445-6_10 |
| GCA_900079115.1 | Hyperthermophiles | Acidophiles | 3034024 | Archaea | Thermoproteota | Thermoproteia | Sulfolobales | Sulfolobaceae | Saccharolobus | *Saccharolobus solfataricus* | 2287 | *Saccharolobus solfataricus* | https://doi.org/10.1099/ijsem.0.002665 |
| GCA_001719125.1 | Thermophiles | Acidophiles | 2688317 | Archaea | Thermoproteota | Thermoproteia | Sulfolobales | Sulfolobaceae | Saccharolobus | *Saccharolobus sp001719125* | 1891280 | *Sulfolobus sp. A20* | https://www.frontiersin.org/articles/10.3389/fmicb.2016.01902/full |
| GCA_007004735.1 | Thermophiles | | 4196798 | Archaea | Halobacteriota | Halobacteria | Halobacteriales | Haloferacaceae | Salinigranum | *Salinigranum halophilum* | 2565931 | *Salinigranum halophilum* | https://www.microbiologyresearch.org/content/journal/ijsem/10.1099/ijsem.0.003951 |
| GCA_003226325.1 | | Alkaliphiles | 4150426 | Bacteria | Firmicutes | Bacilli | Bacillales_H | Salisediminibacteriaceae | Salipaludibacillus | *Salipaludibacillus keqinensis* | 2045207 | *Salipaludibacillus keqinensis* | https://pubmed.ncbi.nlm.nih.gov/30788630/ |
| GCA_009183365.2 | Mesophiles | | 4924764 | Bacteria | Proteobacteria | Gammaproteobacteria | Enterobacterales | Shewanellaceae | Shewanella | *Shewanella algae* | 38313 | *Shewanella algae* | https://bacdive.dsmz.de/strain/14062 |
| GCA_003605125.1 | Psychrophiles | | 4203325 | Bacteria | Proteobacteria | Gammaproteobacteria | Enterobacterales | Shewanellaceae | Shewanella | *Shewanella algidipiscicola* | 614070 | *Shewanella algidipiscicola* | https://doi.org/10.1099/ijs.0.64708-0 |
| GCA_003966265.1 | Psychrophiles | | 5392127 | Bacteria | Proteobacteria | Gammaproteobacteria | Enterobacterales | Shewanellaceae | Shewanella | *Shewanella atlantica* | 271099 | *Shewanella atlantica* | https://doi.org/10.1099/ijs.0.64708-0 |
| GCA_900456975.1 | Psychrophiles | | 5300842 | Bacteria | Proteobacteria | Gammaproteobacteria | Enterobacterales | Shewanellaceae | Shewanella | *Shewanella baltica* | 693974 | *Shewanella baltica BA175* | https://doi.org/10.1016/j.jprot.2019.103419 |
| GCA_003966225.1 | Psychrophiles | | 5676915 | Bacteria | Proteobacteria | Gammaproteobacteria | Enterobacterales | Shewanellaceae | Shewanella | *Shewanella canadensis* | 271096 | *Shewanella canadensis* | https://doi.org/10.1099/ijs.0.64596-0 |
| GCA_002777975.1 | Mesophiles | | 4375287 | Bacteria | Proteobacteria | Gammaproteobacteria | Enterobacterales | Shewanellaceae | Shewanella | *Shewanella carassii* | 1987584 | *Shewanella carassii* | https://bacdive.dsmz.de/strain/158312 |
| GCA_002836945.1 | | Alkaliphiles | 4399136 | Bacteria | Proteobacteria | Gammaproteobacteria | Enterobacterales | Shewanellaceae | Shewanella | *Shewanella chilikensis* | 558541 | *Shewanella chilikensis* | https://doi.org/10.1099/ijs.0.010918-0 |
| GCA_000518705.1 | Mesophiles | | 4575622 | Bacteria | Proteobacteria | Gammaproteobacteria | Enterobacterales | Shewanellaceae | Shewanella | *Shewanella colwelliana* | 1336240 | *Shewanella colwelliana ATCC 39565* | https://bacdive.dsmz.de/strain/14082 |
| GCA_003353085.1 | Mesophiles | | 5215037 | Bacteria | Proteobacteria | Gammaproteobacteria | Enterobacterales | Shewanellaceae | Shewanella | *Shewanella corallii* | 560080 | *Shewanella corallii* | https://doi.org/10.1099/ijs.0.015768-0 |
| GCA_000013765.1 | Mesophiles | | 4545906 | Bacteria | Proteobacteria | Gammaproteobacteria | Enterobacterales | Shewanellaceae | Shewanella | *Shewanella denitrificans* | 318161 | *Shewanella denitrificans OS217* | https://www.microbiologyresearch.org/docserver/fulltext/ijsem/52/6/0522211a.pdf?expires=1677115145&id=id&accname=guest&checksum=9A4347E7FD89240A16EF15F868F657D3 |
| GCA_007567505.1 | Psychrophiles | | 4912773 | Bacteria | Proteobacteria | Gammaproteobacteria | Enterobacterales | Shewanellaceae | Shewanella | *Shewanella donghaensis* | 238836 | *Shewanella donghaensis* | https://doi.org/10.1099/ijs.0.64469-0 |
| GCA_000518605.1 | Mesophiles | | 4798688 | Bacteria | Proteobacteria | Gammaproteobacteria | Enterobacterales | Shewanellaceae | Shewanella | *Shewanella fidelis* | 1336247 | *Shewanella fidelis ATCC BAA-318* | https://www.microbiologyresearch.org/content/journal/ijsem/10.1099/ijs.0.02198-0 |
| GCA_014651955.1 | Mesophiles | | 3905718 | Bacteria | Proteobacteria | Gammaproteobacteria | Enterobacterales | Shewanellaceae | Shewanella | *Shewanella fodinae* | 552357 | *Shewanella fodinae* | https://doi.org/10.1099/ijs.0.017046-0 |
| GCA_003797125.1 | Psychrophiles | | 4784071 | Bacteria | Proteobacteria | Gammaproteobacteria | Enterobacterales | Shewanellaceae | Shewanella | *Shewanella frigidimarina* | 56812 | *Shewanella frigidimarina* | https://www.microbiologyresearch.org/content/journal/ijsem/10.1099/00207713-52-1-195 |
| GCA_000019185.1 | Psychrophiles | | 5226917 | Bacteria | Proteobacteria | Gammaproteobacteria | Enterobacterales | Shewanellaceae | Shewanella | *Shewanella halifaxensis* | 271098 | *Shewanella halifaxensis* | https://doi.org/10.1099/ijs.0.63829-0 |
| GCA_007197645.1 | Psychrophiles | | 5939027 | Bacteria | Proteobacteria | Gammaproteobacteria | Enterobacterales | Shewanellaceae | Shewanella | *Shewanella hanedai* | 25 | *Shewanella hanedai* | https://www.microbiologyresearch.org/content/journal/ijsem/10.1099/ijsem.0.005152 |
| GCA_002836975.1 | | Alkaliphiles | 4402806 | Bacteria | Proteobacteria | Gammaproteobacteria | Enterobacterales | Shewanellaceae | Shewanella | *Shewanella indica* | 768528 | *Shewanella indica* | https://pubmed.ncbi.nlm.nih.gov/20851908/ |
| GCA_002075795.1 | Mesophiles | | 4975677 | Bacteria | Proteobacteria | Gammaproteobacteria | Enterobacterales | Shewanellaceae | Shewanella | *Shewanella japonica* | 93973 | *Shewanella japonica* | https://www.microbiologyresearch.org/content/journal/ijsem/10.1099/00207713-51-3-1027 |
| GCA_003855395.1 | Psychrophiles | | 4839879 | Bacteria | Proteobacteria | Gammaproteobacteria | Enterobacterales | Shewanellaceae | Shewanella | *Shewanella livingstonensis* | 150120 | *Shewanella livingstonensis* | https://doi.org/10.1099/00207713-52-1-195 |
| GCA_000016065.1 | Psychrophiles | | 4602594 | Bacteria | Proteobacteria | Gammaproteobacteria | Enterobacterales | Shewanellaceae | Shewanella | *Shewanella loihica* | 323850 | *Shewanella loihica PV-4* | https://doi.org/10.1099/ijs.0.64354-0 |
| GCA_000753795.1 | Psychrophiles | | 4215794 | Bacteria | Proteobacteria | Gammaproteobacteria | Enterobacterales | Shewanellaceae | Shewanella | *Shewanella mangrovi* | 1515746 | *Shewanella mangrovi* | https://doi.org/10.1099/ijs.0.000313 |
| GCA_000614975.1 | Psychrophiles | | 4424648 | Bacteria | Proteobacteria | Gammaproteobacteria | Enterobacterales | Shewanellaceae | Shewanella | *Shewanella marina* | 1236542 | *Shewanella marina JCM 15074* | https://www.microbiologyresearch.org/content/journal/ijsem/10.1099/ijs.0.005470-0 |
| GCA_002215585.1 | Psychrophiles | | 4321452 | Bacteria | Proteobacteria | Gammaproteobacteria | Enterobacterales | Shewanellaceae | Shewanella | *Shewanella marisflavi* | 260364 | *Shewanella marisflavi* | https://doi.org/10.1099/ijs.0.63198-0 |
| GCA_900156405.1 | Psychrophiles | | 4190369 | Bacteria | Proteobacteria | Gammaproteobacteria | Enterobacterales | Shewanellaceae | Shewanella | *Shewanella morhuae* | 365591 | *Shewanella morhuae* | https://www.microbiologyresearch.org/content/journal/ijsem/10.1099/ijs.0.63931-0 |
| GCA_000018285.1 | Mesophiles | | 5174581 | Bacteria | Proteobacteria | Gammaproteobacteria | Enterobacterales | Shewanellaceae | Shewanella | *Shewanella pealeana* | 398579 | *Shewanella pealeana ATCC 700345* | https://doi.org/10.1099/00207713-49-4-1341 |
| GCA_000014885.1 | Psychrophiles | | 5396476 | Bacteria | Proteobacteria | Gammaproteobacteria | Enterobacterales | Shewanellaceae | Shewanella | *Shewanella piezotolerans* | 225849 | *Shewanella piezotolerans WP3* | https://doi.org/10.1099/ijs.0.64500-0 |
| GCA_002005305.1 | Psychrophiles | | 6353406 | Bacteria | Proteobacteria | Gammaproteobacteria | Enterobacterales | Shewanellaceae | Shewanella | *Shewanella psychrophila* | 225848 | *Shewanella psychrophila* | https://doi.org/10.1099/ijs.0.64500-0 |
| GCA_016406325.1 | Mesophiles | | 4386330 | Bacteria | Proteobacteria | Gammaproteobacteria | Enterobacterales | Shewanellaceae | Shewanella | *Shewanella putrefaciens* | 24 | *Shewanella putrefaciens* | https://bacdive.dsmz.de/strain/14054 |
| GCA_016406305.1 | Mesophiles | | 4575397 | Bacteria | Proteobacteria | Gammaproteobacteria | Enterobacterales | Shewanellaceae | Shewanella | *Shewanella putrefaciens_C* | 24 | *Shewanella putrefaciens* | https://bacdive.dsmz.de/strain/14054 |
| GCA_000018025.1 | Psychrophiles | | 5517674 | Bacteria | Proteobacteria | Gammaproteobacteria | Enterobacterales | Shewanellaceae | Shewanella | *Shewanella sediminis* | 425104 | *Shewanella sediminis HAW-EB3* | https://doi.org/10.1099/ijs.0.63604-0 |
| GCA_003797885.1 | Psychrophiles | | 4722637 | Bacteria | Proteobacteria | Gammaproteobacteria | Enterobacterales | Shewanellaceae | Shewanella | *Shewanella vesiculosa* | 518738 | *Shewanella vesiculosa* | https://doi.org/10.1099/ijs.0.000737-0 |
| GCA_000091325.1 | Psychrophiles | | 4962103 | Bacteria | Proteobacteria | Gammaproteobacteria | Enterobacterales | Shewanellaceae | Shewanella | *Shewanella violacea* | 637905 | *Shewanella violacea DSS12* | https://doi.org/10.1039/c000396d |

| | | | | | | | | | | | | | |
|---|---|---|---|---|---|---|---|---|---|---|---|---|---|
| GCA_000518805.1 | Mesophiles | | 4971480 | Bacteria | Proteobacteria | Gammaproteobacteria | Enterobacterales | Shewanellaceae | Shewanella | *Shewanella waksmanii* | 1336233 | *Shewanella waksmanii ATCC BAA-643* | https://pubmed.ncbi.nlm.nih.gov/13130035/ |
| GCA_000019525.1 | Psychrophiles | | 5935403 | Bacteria | Proteobacteria | Gammaproteobacteria | Enterobacterales | Shewanellaceae | Shewanella | *Shewanella woodyi* | 392500 | *Shewanella woodyi ATCC 51908* | https://www.microbiologyresearch.org/content/journal/ijsem/10.1099/00207713-47-4-1034 |
| GCA_014647135.1 | Mesophiles | | 4600673 | Bacteria | Proteobacteria | Gammaproteobacteria | Enterobacterales | Shewanellaceae | Shewanella | *Shewanella xiamenensis* | 332186 | *Shewanella xiamenensis* | https://bacdive.dsmz.de/strain/14084 |
| GCA_004123295.1 | | Acidophiles | 5116187 | Bacteria | Acidobacteriota | Acidobacteriae | Acidobacteriales | Acidobacteriaceae | Silvibacterium | *Silvibacterium dinghuense* | 1560006 | *Silvibacterium dinghuense* | https://doi.org/10.1099/ijsem.0.005415 |
| GCA_900105965.1 | Psychrophiles | | 4622641 | Bacteria | Actinobacteriota | Actinomycetia | Actinomycetales | Micrococcaceae | Specibacter | *Specibacter alpinus* | 656366 | *Arthrobacter alpinus* | https://doi.org/10.1099/ijs.0.017178-0 |
| GCA_001294625.1 | Psychrophiles | | 4046453 | Bacteria | Actinobacteriota | Actinomycetia | Actinomycetales | Micrococcaceae | Specibacter | *Specibacter alpinus_A* | 656366 | *Arthrobacter alpinus* | https://doi.org/10.1099/ijs.0.017178-0 |
| GCA_001445575.1 | Psychrophiles | | 4333648 | Bacteria | Actinobacteriota | Actinomycetia | Actinomycetales | Micrococcaceae | Specibacter | *Specibacter alpinus_C* | 656366 | *Arthrobacter alpinus* | https://doi.org/10.1099/ijs.0.017178-0 |
| GCA_002909445.1 | Psychrophiles | | 4469153 | Bacteria | Actinobacteriota | Actinomycetia | Actinomycetales | Micrococcaceae | Specibacter | *Specibacter glacialis* | 1664 | *Arthrobacter glacialis* | https://doi.org/10.1016/j.syapm.2018.10.005 |
| GCA_003219815.1 | Psychrophiles | | 5042614 | Bacteria | Actinobacteriota | Actinomycetia | Actinomycetales | Micrococcaceae | Specibacter | *Specibacter livingstonensis* | 670078 | *Arthrobacter livingstonensis* | https://doi.org/10.1099/ijs.0.021022-0 |
| GCA_003185915.1 | Psychrophiles | | 4049680 | Bacteria | Actinobacteriota | Actinomycetia | Actinomycetales | Micrococcaceae | Specibacter | *Specibacter psychrochitiniphilus* | 291045 | *Arthrobacter psychrochitiniphilus* | https://doi.org/10.1099/ijs.0.008912-0 |
| GCA_003219795.1 | Psychrophiles | | 3871146 | Bacteria | Actinobacteriota | Actinomycetia | Actinomycetales | Micrococcaceae | Specibacter | *Specibacter psychrolactophilus* | 92442 | *Arthrobacter psychrolactophilus* | https://doi.org/10.1007/s002030050722 |
| GCA_000242595.3 | | Alkaliphiles | 3285855 | Bacteria | Spirochaetota | Spirochaetia | DSM-27196 | DSM-8902 | Spirochaeta_B | *Spirochaeta_B africana* | 889378 | *Spirochaeta africana DSM 8902* | https://doi.org/10.1099/00207713-46-1-305 |
| GCA_002901865.1 | Mesophiles | | 2491359 | Bacteria | Firmicutes | Bacilli | Staphylococcales | Staphylococcaceae | Staphylococcus | *Staphylococcus agnetis* | 985762 | *Staphylococcus agnetis* | https://bacdive.dsmz.de/strain/14677 |
| GCA_002902305.1 | Mesophiles | | 2452468 | Bacteria | Firmicutes | Bacilli | Staphylococcales | Staphylococcaceae | Staphylococcus | *Staphylococcus argensis* | 1607738 | *Staphylococcus argensis* | https://bacdive.dsmz.de/strain/132229 |
| GCA_000236925.1 | Mesophiles | | 2787638 | Bacteria | Firmicutes | Bacilli | Staphylococcales | Staphylococcaceae | Staphylococcus | *Staphylococcus argenteus* | 985002 | *Staphylococcus argenteus* | https://bacdive.dsmz.de/strain/130706 |
| GCA_002902345.1 | Mesophiles | | 2665344 | Bacteria | Firmicutes | Bacilli | Staphylococcales | Staphylococcaceae | Staphylococcus | *Staphylococcus arlettae* | 29378 | *Staphylococcus arlettae* | https://bacdive.dsmz.de/strain/14441 |
| GCA_001027105.1 | Mesophiles | | 2782562 | Bacteria | Firmicutes | Bacilli | Staphylococcales | Staphylococcaceae | Staphylococcus | *Staphylococcus aureus* | 1280 | *Staphylococcus aureus* | https://bacdive.dsmz.de/strain/14487 |
| GCA_001500315.1 | Mesophiles | | 2201708 | Bacteria | Firmicutes | Bacilli | Staphylococcales | Staphylococcaceae | Staphylococcus | *Staphylococcus auricularis* | 29379 | *Staphylococcus auricularis* | https://bacdive.dsmz.de/strain/14503 |
| GCA_002902325.1 | Mesophiles | | 2434909 | Bacteria | Firmicutes | Bacilli | Staphylococcales | Staphylococcaceae | Staphylococcus | *Staphylococcus capitis* | 904334 | *Staphylococcus capitis VCU116* | https://bacdive.dsmz.de/strain/14508 |
| GCA_002902725.1 | Mesophiles | | 2606761 | Bacteria | Firmicutes | Bacilli | Staphylococcales | Staphylococcaceae | Staphylococcus | *Staphylococcus caprae* | 29380 | *Staphylococcus caprae* | https://bacdive.dsmz.de/strain/14509 |
| GCA_900458435.1 | Mesophiles | | 2576833 | Bacteria | Firmicutes | Bacilli | Staphylococcales | Staphylococcaceae | Staphylococcus | *Staphylococcus carnosus* | 1281 | *Staphylococcus carnosus* | https://bacdive.dsmz.de/strain/14516 |
| GCA_002901945.1 | Mesophiles | | 2276768 | Bacteria | Firmicutes | Bacilli | Staphylococcales | Staphylococcaceae | Staphylococcus | *Staphylococcus chromogenes* | 46126 | *Staphylococcus chromogenes* | https://bacdive.dsmz.de/strain/14517 |
| GCA_900636325.1 | Mesophiles | | 2798161 | Bacteria | Firmicutes | Bacilli | Staphylococcales | Staphylococcaceae | Staphylococcus | *Staphylococcus delphini* | 53344 | *Staphylococcus delphini* | https://bacdive.dsmz.de/strain/14521 |
| GCA_002902625.1 | Mesophiles | | 2379883 | Bacteria | Firmicutes | Bacilli | Staphylococcales | Staphylococcaceae | Staphylococcus | *Staphylococcus devriesei* | 586733 | *Staphylococcus devriesei* | https://www.microbiologyresearch.org/content/journal/ijsem/10.1099/ijs.0.015982-0 |
| GCA_003035445.1 | Mesophiles | | 2394070 | Bacteria | Firmicutes | Bacilli | Staphylococcales | Staphylococcaceae | Staphylococcus | *Staphylococcus devriesei_A* | 586733 | *Staphylococcus devriesei* | https://www.microbiologyresearch.org/content/journal/ijsem/10.1099/ijs.0.015982-0 |
| GCA_900458565.1 | Mesophiles | | 2758007 | Bacteria | Firmicutes | Bacilli | Staphylococcales | Staphylococcaceae | Staphylococcus | *Staphylococcus equorum* | 1357294 | *Staphylococcus equorum UMC-CNS-924* | https://bacdive.dsmz.de/strain/14662 |
| GCA_003012915.1 | Mesophiles | | 2479423 | Bacteria | Firmicutes | Bacilli | Staphylococcales | Staphylococcaceae | Staphylococcus | *Staphylococcus felis* | 46127 | *Staphylococcus felis* | https://bacdive.dsmz.de/strain/14607 |
| GCA_000875895.1 | Mesophiles | | 3171720 | Bacteria | Firmicutes | Bacilli | Staphylococcales | Staphylococcaceae | Staphylococcus | *Staphylococcus gallinarum* | 1293 | *Staphylococcus gallinarum* | https://bacdive.dsmz.de/strain/14542 |
| GCA_006094395.1 | Mesophiles | | 2572027 | Bacteria | Firmicutes | Bacilli | Staphylococcales | Staphylococcaceae | Staphylococcus | *Staphylococcus haemolyticus* | 1283 | *Staphylococcus haemolyticus* | https://bacdive.dsmz.de/strain/14544 |
| GCA_002901845.1 | Mesophiles | | 2204528 | Bacteria | Firmicutes | Bacilli | Staphylococcales | Staphylococcaceae | Staphylococcus | *Staphylococcus hominis* | 1290 | *Staphylococcus hominis* | https://bacdive.dsmz.de/strain/14639 |
| GCA_000816085.1 | Mesophiles | | 2472129 | Bacteria | Firmicutes | Bacilli | Staphylococcales | Staphylococcaceae | Staphylococcus | *Staphylococcus hyicus* | 1284 | *Staphylococcus hyicus* | https://bacdive.dsmz.de/strain/14553 |
| GCA_002902385.1 | Mesophiles | | 2801199 | Bacteria | Firmicutes | Bacilli | Staphylococcales | Staphylococcaceae | Staphylococcus | *Staphylococcus intermedius* | 1285 | *Staphylococcus intermedius* | https://bacdive.dsmz.de/strain/14555 |
| GCA_003019255.1 | Mesophiles | | 2639038 | Bacteria | Firmicutes | Bacilli | Staphylococcales | Staphylococcaceae | Staphylococcus | *Staphylococcus kloosii* | 29384 | *Staphylococcus kloosii* | https://bacdive.dsmz.de/strain/14556 |
| GCA_002901705.1 | Mesophiles | | 2519514 | Bacteria | Firmicutes | Bacilli | Staphylococcales | Staphylococcaceae | Staphylococcus | *Staphylococcus lugdunensis* | 904354 | *Staphylococcus lugdunensis VCU150* | https://bacdive.dsmz.de/strain/14562 |
| GCA_000298075.1 | Mesophiles | | 2366595 | Bacteria | Firmicutes | Bacilli | Staphylococcales | Staphylococcaceae | Staphylococcus | *Staphylococcus massiliensis* | 1229783 | *Staphylococcus massiliensis S46* | https://bacdive.dsmz.de/strain/14673 |
| GCA_000934465.1 | Mesophiles | | 2381859 | Bacteria | Firmicutes | Bacilli | Staphylococcales | Staphylococcaceae | Staphylococcus | *Staphylococcus microti* | 569857 | *Staphylococcus microti* | https://bacdive.dsmz.de/strain/14672 |
| GCA_003019275.1 | Mesophiles | | 2095131 | Bacteria | Firmicutes | Bacilli | Staphylococcales | Staphylococcaceae | Staphylococcus | *Staphylococcus muscae* | 1294 | *Staphylococcus muscae* | https://bacdive.dsmz.de/strain/14605 |
| GCA_014635045.1 | Mesophiles | | 2849797 | Bacteria | Firmicutes | Bacilli | Staphylococcales | Staphylococcaceae | Staphylococcus | *Staphylococcus nepalensis* | 214473 | *Staphylococcus nepalensis* | https://bacdive.dsmz.de/strain/14652 |
| GCA_003970495.1 | Mesophiles | | 2462952 | Bacteria | Firmicutes | Bacilli | Staphylococcales | Staphylococcaceae | Staphylococcus | *Staphylococcus pasteuri* | 45972 | *Staphylococcus pasteuri* | https://bacdive.dsmz.de/strain/14617 |
| GCA_002902685.1 | Mesophiles | | 2455272 | Bacteria | Firmicutes | Bacilli | Staphylococcales | Staphylococcaceae | Staphylococcus | *Staphylococcus pettenkoferi* | 170573 | *Staphylococcus pettenkoferi* | https://bacdive.dsmz.de/strain/14671 |
| GCA_900186985.1 | Mesophiles | | 2613271 | Bacteria | Firmicutes | Bacilli | Staphylococcales | Staphylococcaceae | Staphylococcus | *Staphylococcus piscifermentans* | 70258 | *Staphylococcus piscifermentans* | https://bacdive.dsmz.de/strain/14606 |
| GCA_001792775.2 | Mesophiles | | 2519048 | Bacteria | Firmicutes | Bacilli | Staphylococcales | Staphylococcaceae | Staphylococcus | *Staphylococcus pseudintermedius* | 283734 | *Staphylococcus pseudintermedius* | https://bacdive.dsmz.de/strain/14670 |
| GCA_000010125.1 | Mesophiles | | 2577899 | Bacteria | Firmicutes | Bacilli | Staphylococcales | Staphylococcaceae | Staphylococcus | *Staphylococcus saprophyticus* | 29385 | *Staphylococcus saprophyticus* | https://bacdive.dsmz.de/strain/14626 |
| GCA_002902405.1 | Mesophiles | | 2743713 | Bacteria | Firmicutes | Bacilli | Staphylococcales | Staphylococcaceae | Staphylococcus | *Staphylococcus schweitzeri* | 1654388 | *Staphylococcus schweitzeri* | https://bacdive.dsmz.de/strain/130707 |
| GCA_002902285.1 | Mesophiles | | 2735408 | Bacteria | Firmicutes | Bacilli | Staphylococcales | Staphylococcaceae | Staphylococcus | *Staphylococcus simulans* | 1286 | *Staphylococcus simulans* | https://bacdive.dsmz.de/strain/14572 |
| GCA_003043455.1 | Mesophiles | | 2556508 | Bacteria | Firmicutes | Bacilli | Staphylococcales | Staphylococcaceae | Staphylococcus | *Staphylococcus simulans_A* | 1286 | *Staphylococcus simulans* | https://bacdive.dsmz.de/strain/14572 |
| GCA_002994445.1 | Mesophiles | | 2595226 | Bacteria | Firmicutes | Bacilli | Staphylococcales | Staphylococcaceae | Staphylococcus | *Staphylococcus simulans_B* | 1286 | *Staphylococcus simulans* | https://bacdive.dsmz.de/strain/14572 |
| GCA_001006765.1 | Mesophiles | | 2887686 | Bacteria | Firmicutes | Bacilli | Staphylococcales | Staphylococcaceae | Staphylococcus | *Staphylococcus succinus* | 61015 | *Staphylococcus succinus* | https://www.microbiologyresearch.org/content/journal/ijsem/10.1099/00207713-48-2-511 |
| GCA_002902235.1 | Mesophiles | | 2670427 | Bacteria | Firmicutes | Bacilli | Staphylococcales | Staphylococcaceae | Staphylococcus | *Staphylococcus ureilyticus* | 94138 | *Staphylococcus ureilyticus* | https://bacdive.dsmz.de/strain/136260 |
| GCA_900636385.1 | Mesophiles | | 2427576 | Bacteria | Firmicutes | Bacilli | Staphylococcales | Staphylococcaceae | Staphylococcus | *Staphylococcus warneri* | 904338 | *Staphylococcus warneri VCU121* | https://bacdive.dsmz.de/strain/14587 |
| GCA_002732165.1 | Mesophiles | | 2734769 | Bacteria | Firmicutes | Bacilli | Staphylococcales | Staphylococcaceae | Staphylococcus | *Staphylococcus xylosus* | 1288 | *Staphylococcus xylosus* | https://bacdive.dsmz.de/strain/14599 |
| GCA_000338275.1 | Mesophiles | | 2939263 | Bacteria | Firmicutes | Bacilli | Staphylococcales | Staphylococcaceae | Staphylococcus | *Staphylococcus xylosus_B* | 1288 | *Staphylococcus xylosus* | https://bacdive.dsmz.de/strain/14599 |
| GCA_000015945.1 | Hyperthermophiles | | 1570485 | Archaea | Thermoproteota | Thermoproteia | Sulfolobales | Desulfurococcaceae | Staphylothermus | *Staphylothermus marinus* | 399550 | *Staphylothermus marinus F1* | https://www.sciencedirect.com/science/article/pii/S0168165606008595?via%3Dihub#aep-section-id32 |
| GCA_013343115.1 | Mesophiles | | 2358846 | Bacteria | Firmicutes | Bacilli | Lactobacillales | Streptococcaceae | Streptococcus | *Streptococcus sanguinis_H* | 888825 | *Streptococcus sanguinis VMC66* | https://bacdive.dsmz.de/strain/14767 |
| GCA_010604095.1 | Mesophiles | | 2173350 | Bacteria | Firmicutes | Bacilli | Lactobacillales | Streptococcaceae | Streptococcus | *Streptococcus sp000187445* | 1343 | *Streptococcus vestibularis* | https://bacdive.dsmz.de/strain/14794 |
| GCA_900095845.1 | Mesophiles | | 1925331 | Bacteria | Firmicutes | Bacilli | Lactobacillales | Streptococcaceae | Streptococcus | *Streptococcus timonensis* | 1852387 | *Streptococcus timonensis* | https://www.sciencedirect.com/science/article/pii/S2052297516301251?via%3Dihub |
| GCA_002355215.1 | Mesophiles | | 2097874 | Bacteria | Firmicutes | Bacilli | Lactobacillales | Streptococcaceae | Streptococcus | *Streptococcus troglodytae* | 1111760 | *Streptococcus troglodytae* | https://www.microbiologyresearch.org/content/journal/ijsem/10.1099/ijs.0.039388-0 |
| GCA_900475595.1 | Mesophiles | | 1975601 | Bacteria | Firmicutes | Bacilli | Lactobacillales | Streptococcaceae | Streptococcus | *Streptococcus uberis* | 1349 | *Streptococcus uberis* | https://bacdive.dsmz.de/strain/14787 |
| GCA_000785785.1 | Mesophiles | | 2149440 | Bacteria | Firmicutes | Bacilli | Lactobacillales | Streptococcaceae | Streptococcus | *Streptococcus uberis_A* | 1349 | *Streptococcus uberis* | https://bacdive.dsmz.de/strain/14787 |
| GCA_000188055.3 | Mesophiles | | 2130431 | Bacteria | Firmicutes | Bacilli | Lactobacillales | Streptococcaceae | Streptococcus | *Streptococcus urinalis* | 764291 | *Streptococcus urinalis 2285-97* | https://bacdive.dsmz.de/strain/14822 |
| GCA_001375655.1 | Mesophiles | | 2460376 | Bacteria | Firmicutes | Bacilli | Lactobacillales | Streptococcaceae | Streptococcus | *Streptococcus varani* | 1608583 | *Streptococcus varani* | https://bacdive.dsmz.de/strain/139780 |
| GCA_000188295.1 | Mesophiles | | 1872773 | Bacteria | Firmicutes | Bacilli | Lactobacillales | Streptococcaceae | Streptococcus | *Streptococcus vestibularis* | 1343 | *Streptococcus vestibularis* | https://bacdive.dsmz.de/strain/14794 |
| GCA_009729035.1 | Thermophiles | Acidophiles | 1987069 | Archaea | Thermoproteota | Thermoproteia | Sulfolobales | Sulfolobaceae | Stygiolobus | *Stygiolobus azoricus* | 1313299 | *Stygiolobus azoricus* | https://doi.org/10.1128/mra.01490-19 |
| GCA_006716115.1 | Psychrophiles | | 4214389 | Bacteria | Actinobacteriota | Actinomycetia | Actinomycetales | Microbacteriaceae | Subtercola | *Subtercola boreus* | 120213 | *Subtercola boreus* | https://doi.org/10.1099/00207713-50-5-1731 |
| GCA_003399645.1 | Psychrophiles | | 3670876 | Bacteria | Actinobacteriota | Actinomycetia | Actinomycetales | Microbacteriaceae | Subtercola | *Subtercola boreus_A* | 120213 | *Subtercola boreus* | https://doi.org/10.1099/00207713-50-5-1731 |
| GCA_003399635.1 | Psychrophiles | | 3848592 | Bacteria | Actinobacteriota | Actinomycetia | Actinomycetales | Microbacteriaceae | Subtercola | *Subtercola boreus_B* | 120213 | *Subtercola boreus* | https://doi.org/10.1099/00207713-50-5-1731 |
| GCA_003399625.1 | Psychrophiles | | 4121892 | Bacteria | Actinobacteriota | Actinomycetia | Actinomycetales | Microbacteriaceae | Subtercola | *Subtercola boreus_C* | 120213 | *Subtercola boreus* | https://doi.org/10.1099/00207713-50-5-1731 |

| | | | | | | | | | | | | | |
|---|---|---|---|---|---|---|---|---|---|---|---|---|---|
| GCA_004923255.1 | Psychrophiles | | 4043135 | Bacteria | Actinobacteriota | Actinomycetia | Actinomycetales | Microbacteriaceae | Subtercola | *Subtercola vilae* | 2056433 | *Subtercola vilae* | https://pubmed.ncbi.nlm.nih.gov/29214367/ |
| GCA_003023725.1 | | Acidophiles | 4556669 | Bacteria | Firmicutes_E | Sulfobacillia | Sulfobacillales | Sulfobacillaceae | Sulfobacillus | *Sulfobacillus benefaciens_A* | 453960 | *Sulfobacillus benefaciens* | https://doi.org/10.1007/s00792-008-0184-4 |
| GCA_900176145.1 | Thermophiles | Acidophiles | 3861015 | Bacteria | Firmicutes_E | Sulfobacillia | Sulfobacillales | Sulfobacillaceae | Sulfobacillus | *Sulfobacillus thermosulfidooxidans* | 28034 | *Sulfobacillus thermosulfidooxidans* | https://www.ncbi.nlm.nih.gov/pmc/articles/PMC7000362/ |
| GCA_001280565.1 | Thermophiles | Acidophiles | 3828023 | Bacteria | Firmicutes_E | Sulfobacillia | Sulfobacillales | Sulfobacillaceae | Sulfobacillus | *Sulfobacillus thermosulfidooxidans_A* | 28034 | *Sulfobacillus thermosulfidooxidans* | https://www.ncbi.nlm.nih.gov/pmc/articles/PMC7000362/ |
| GCA_000237975.1 | | Acidophiles | 3557831 | Bacteria | Firmicutes_E | Sulfobacillia | Sulfobacillales | Sulfobacillaceae | Sulfobacillus_A | *Sulfobacillus_A acidophilus* | 1051632 | *Sulfobacillus acidophilus TPY* | https://doi.org/10.4056/sigs.2736042 |
| GCA_003023695.1 | | Acidophiles | 4108624 | Bacteria | Firmicutes_E | Sulfobacillia | Sulfobacillales | Sulfobacillaceae | Sulfobacillus_C | *Sulfobacillus_C benefaciens_A* | 453960 | *Sulfobacillus benefaciens* | https://doi.org/10.1007/s00792-008-0184-4 |
| GCA_003967175.1 | Thermophiles | Acidophiles | 2353189 | Archaea | Thermoproteota | Thermoproteia | Sulfolobales | Sulfolobaceae | Sulfodiicoccus | *Sulfodiicoccus acidiphilus* | 1670455 | *Sulfodiicoccus acidiphilus* | https://doi.org/10.1099/ijsem.0.001881 |
| GCA_000012285.1 | Thermophiles | Acidophiles | 2225959 | Archaea | Thermoproteota | Thermoproteia | Sulfolobales | Sulfolobaceae | Sulfolobus | *Sulfolobus acidocaldarius* | 2285 | *Sulfolobus acidocaldarius* | https://pubmed.ncbi.nlm.nih.gov/2478523/ |
| GCA_000508305.1 | Thermophiles | Acidophiles | 2061920 | Archaea | Thermoproteota | Thermoproteia | Sulfolobales | Sulfolobaceae | Sulfolobus | *Sulfolobus acidocaldarius_A* | 2285 | *Sulfolobus acidocaldarius* | https://pubmed.ncbi.nlm.nih.gov/2478523/ |
| GCA_009938015.1 | | Acidophiles | 3452774 | Bacteria | Proteobacteria | Gammaproteobacteria | Burkholderiales | Sulfuriferulaceae | Sulfuriferula | *Sulfuriferula plumbiphila* | 171865 | *Sulfuriferula plumbiphila* | https://www.microbiologyresearch.org/content/journal/ijsem/10.1099/ijsem.0.004166 |
| GCA_009729055.1 | Thermophiles | Acidophiles | 2803915 | Archaea | Thermoproteota | Thermoproteia | Sulfolobales | Sulfolobaceae | Sulfurisphaera | *Sulfurisphaera ohwakuensis* | 69656 | *Sulfurisphaera ohwakuensis* | https://doi.org/10.1099/00207713-48-2-451 |
| GCA_000011205.1 | Hyperthermophiles | | 2694756 | Archaea | Thermoproteota | Thermoproteia | Sulfolobales | Sulfolobaceae | Sulfurisphaera | *Sulfurisphaera tokodaii* | 273063 | *Sulfurisphaera tokodaii str. 7* | https://pubmed.ncbi.nlm.nih.gov/30304218/ |
| GCA_002754935.1 | Thermophiles | | 2659739 | Bacteria | Cyanobacteria | Cyanobacteriia | Thermosynechococcales | Thermosynechococcaceae | Synechococcus_A | *Synechococcus_A lividus* | 1917166 | *Thermostichus lividus PCC 6715* | https://pubmed.ncbi.nlm.nih.gov/35602029/ |
| GCA_004634385.1 | Hyperthermophiles | | 3039102 | Bacteria | Firmicutes_E | Thermaerobacteria | Thermaerobacterales | Thermaerobacteraceae | Thermaerobacter | *Thermaerobacter sp004634385* | 2546351 | *Thermaerobacter sp. FW80* | http://www.kjom.org/journal/view.html?uid=176&pn=lastest&vmd=Full |
| GCA_000183545.3 | | Alkaliphiles | 2888741 | Bacteria | Firmicutes_E | Thermaerobacteria | Thermaerobacterales | Thermaerobacteraceae | Thermaerobacter | *Thermaerobacter subterraneus* | 867903 | *Thermaerobacter subterraneus DSM 13965* | https://pubmed.ncbi.nlm.nih.gov/12054240/ |
| GCA_900176005.1 | Thermophiles | | 3334124 | Bacteria | Firmicutes_B | Moorellia | Moorellales | Moorellaceae | Thermanaeromonas | *Thermanaeromonas toyohensis* | 698762 | *Thermanaeromonas toyohensis ToBE* | https://doi.org/10.1099/00207713-52-5-1675 |
| GCA_003722315.1 | Thermophiles | | 2911280 | Bacteria | Firmicutes_A | Thermoanaerobacteria | Thermoanaerobacterales | Thermoanaerobacteraceae | Thermoanaerobacter | *Thermoanaerobacter ethanolicus* | 509192 | *Thermoanaerobacter ethanolicus JW 200* | https://doi.org/10.1128/aem.01773-20 |
| GCA_000763575.1 | Thermophiles | | 2397824 | Bacteria | Firmicutes_A | Thermoanaerobacteria | Thermoanaerobacterales | Thermoanaerobacteraceae | Thermoanaerobacter | *Thermoanaerobacter kivui* | 2325 | *Thermoanaerobacter kivui* | https://bacdive.dsmz.de/strain/16823 |
| GCA_000019085.1 | Thermophiles | | 2362816 | Bacteria | Firmicutes_A | Thermoanaerobacteria | Thermoanaerobacterales | Thermoanaerobacteraceae | Thermoanaerobacter | *Thermoanaerobacter pseudethanolicus* | 509193 | *Thermoanaerobacter brockii subsp. finnii Ako-1* | https://doi.org/10.1099/00207713-45-4-783 |
| GCA_001310975.1 | Thermophiles | | 2452939 | Bacteria | Firmicutes_A | Thermoanaerobacteria | Thermoanaerobacterales | Thermoanaerobacteraceae | Thermoanaerobacter | *Thermoanaerobacter thermocopriae* | 580331 | *Thermoanaerobacter thermocopriae* | https://bacdive.dsmz.de/strain/165540 |
| GCA_900129115.1 | Thermophiles | Acidophiles | 2409754 | Bacteria | Firmicutes_A | Thermoanaerobacteria | Thermoanaerobacterales | Thermoanaerobacteraceae | Thermoanaerobacter | *Thermoanaerobacter uzonensis* | 1123369 | *Thermoanaerobacter uzonensis DSM 18761* | https://doi.org/10.1099/ijs.0.65343-0 |
| GCA_000147695.3 | Thermophiles | | 2785056 | Bacteria | Firmicutes_A | Thermoanaerobacteria | Thermoanaerobacterales | Thermoanaerobacteraceae | Thermoanaerobacter | *Thermoanaerobacter wiegelii* | 697303 | *Thermoanaerobacter wiegelii Rt8.B1* | https://doi.org/10.1007/s007920070014 |
| GCA_000145615.1 | Thermophiles | Acidophiles | 2785752 | Bacteria | Firmicutes_A | Thermoanaerobacteria | Thermoanaerobacterales | Thermoanaerobacteraceae | Thermoanaerobacterium | *Thermoanaerobacterium thermosaccharolyticum* | 1517 | *Thermoanaerobacterium thermosaccharolyticum* | https://bacdive.dsmz.de/strain/18044 |
| GCA_001516585.1 | Thermophiles | | 1876950 | Archaea | Thermoproteota | Thermoproteia | Thermoproteales | Thermocladiaceae | Thermocladium | *Thermocladium sp001516585* | 1714261 | *Thermocladium sp. ECH_B* | https://doi.org/10.1099/00207713-48-3-879 |
| GCA_002214465.1 | Hyperthermophiles | | 1922421 | Archaea | Methanobacteriota_B | Thermococci | Thermococcales | Thermococcaceae | Thermococcus | *Thermococcus barossii* | 54077 | *Thermococcus barossii* | https://doi.org/10.1016/s0723-2020(98)80007-6 |
| GCA_002214365.1 | Hyperthermophiles | | 1866819 | Archaea | Methanobacteriota_B | Thermococci | Thermococcales | Thermococcaceae | Thermococcus | *Thermococcus celer* | 1293037 | *Thermococcus celer Vu 13 = JCM 8558* | https://doi.org/10.1016/s0167-7012(99)00092-5 |
| GCA_001484195.1 | Hyperthermophiles | | 2337139 | Archaea | Methanobacteriota_B | Thermococci | Thermococcales | Thermococcaceae | Thermococcus | *Thermococcus celericrescens* | 2598455 | *Thermococcus celericrescens* | https://doi.org/10.1099/ijs.0.64597-0 |
| GCA_000265525.1 | Hyperthermophiles | | 1950313 | Archaea | Methanobacteriota_B | Thermococci | Thermococcales | Thermococcaceae | Thermococcus | *Thermococcus cleftensis* | 163003 | *Thermococcus cleftensis CL1* | https://doi.org/10.1099/ijs.0.066100-0 |
| GCA_000769655.1 | Hyperthermophiles | | 2126164 | Archaea | Methanobacteriota_B | Thermococci | Thermococcales | Thermococcaceae | Thermococcus | *Thermococcus eurythermalis* | 1505907 | *Thermococcus eurythermalis* | https://doi.org/10.1099/ijs.0.067942-0 |
| GCA_000022365.1 | Hyperthermophiles | | 2045438 | Archaea | Methanobacteriota_B | Thermococci | Thermococcales | Thermococcaceae | Thermococcus | *Thermococcus gammatolerans* | 593117 | *Thermococcus gammatolerans EJ3* | https://doi.org/10.1099/ijs.0.02503-0 |
| GCA_002214385.1 | Thermophiles | | 1674122 | Archaea | Methanobacteriota_B | Thermococci | Thermococcales | Thermococcaceae | Thermococcus | *Thermococcus gorgonarius* | 71997 | *Thermococcus gorgonarius W-12* | https://doi.org/10.1099/00207713-48-1-23 |
| GCA_000816105.1 | Hyperthermophiles | | 1920914 | Archaea | Methanobacteriota_B | Thermococci | Thermococcales | Thermococcaceae | Thermococcus | *Thermococcus guaymasensis* | 1432656 | *Thermococcus guaymasensis DSM 11113* | https://doi.org/10.1099/00207713-48-4-1181 |
| GCA_000009965.1 | Hyperthermophiles | | 2088737 | Archaea | Methanobacteriota_B | Thermococci | Thermococcales | Thermococcaceae | Thermococcus | *Thermococcus kodakarensis* | 69014 | *Thermococcus kodakarensis KOD1* | https://doi.org/10.1007/978-1-0716-2445-6_5 |
| GCA_000585495.1 | Hyperthermophiles | | 1976356 | Archaea | Methanobacteriota_B | Thermococci | Thermococcales | Thermococcaceae | Thermococcus | *Thermococcus nautili* | 195522 | *Thermococcus nautili 30-1* | https://doi.org/10.1099/ijs.0.060376-0 |
| GCA_000018365.1 | Hyperthermophiles | Alkaliphiles | 1847607 | Archaea | Methanobacteriota_B | Thermococci | Thermococcales | Thermococcaceae | Thermococcus | *Thermococcus onnurineus* | 523850 | *Thermococcus onnurineus NA1* | https://bacdive.dsmz.de/strain/161372 |
| GCA_002214485.1 | Thermophiles | | 1785673 | Archaea | Methanobacteriota_B | Thermococci | Thermococcales | Thermococcaceae | Thermococcus | *Thermococcus pacificus* | 71998 | *Thermococcus pacificus P-4* | https://doi.org/10.1099/00207713-48-1-23 |
| GCA_001592435.1 | Thermophiles | | 1896106 | Archaea | Methanobacteriota_B | Thermococci | Thermococcales | Thermococcaceae | Thermococcus | *Thermococcus peptonophilus* | 53952 | *Thermococcus peptonophilus* | https://pubmed.ncbi.nlm.nih.gov/7545383/ |
| GCA_001647085.1 | Hyperthermophiles | | 1928800 | Archaea | Methanobacteriota_B | Thermococci | Thermococcales | Thermococcaceae | Thermococcus | *Thermococcus piezophilus* | 1712654 | *Thermococcus piezophilus* | https://doi.org/10.1016/j.syapm.2016.08.003 |
| GCA_002214565.1 | Hyperthermophiles | | 1868990 | Archaea | Methanobacteriota_B | Thermococci | Thermococcales | Thermococcaceae | Thermococcus | *Thermococcus radiotolerans* | 187880 | *Thermococcus radiotolerans EJ2* | https://doi.org/10.1007/s00792-004-0380-9 |
| GCA_000151205.2 | Hyperthermophiles | | 2086428 | Archaea | Methanobacteriota_B | Thermococci | Thermococcales | Thermococcaceae | Thermococcus | *Thermococcus sp000151205* | 246969 | *Thermococcus sp. AM4* | https://doi.org/10.1128/jb.06259-11 |
| GCA_900198835.1 | Hyperthermophiles | | 2155760 | Archaea | Methanobacteriota_B | Thermococci | Thermococcales | Thermococcaceae | Thermococcus | *Thermococcus sp900198835* | 2016361 | *Thermococcus henrietii* | https://doi.org/10.1099/ijsem.0.004895 |
| GCA_017873335.1 | Hyperthermophiles | | 2034780 | Archaea | Methanobacteriota_B | Thermococci | Thermococcales | Thermococcaceae | Thermococcus | *Thermococcus stetteri* | 49900 | *Thermococcus stetteri* | https://doi.org/10.1093/protein/10.8.905 |
| GCA_002214545.1 | Hyperthermophiles | | 2065932 | Archaea | Methanobacteriota_B | Thermococci | Thermococcales | Thermococcaceae | Thermococcus | *Thermococcus thioreducens* | 277988 | *Thermococcus thioreducens* | https://doi.org/10.1099/ijs.0.65057-0 |
| GCA_000258515.1 | Hyperthermophiles | | 1764559 | Archaea | Methanobacteriota_B | Thermococci | Thermococcales | Thermococcaceae | Thermococcus | *Thermococcus zilligii* | 1151117 | *Thermococcus zilligii AN1* | https://doi.org/10.1128/jb.182.16.4632-4636.2000 |
| GCA_000246985.3 | Hyperthermophiles | | 2215172 | Archaea | Methanobacteriota_B | Thermococci | Thermococcales | Thermococcaceae | Thermococcus_A | *Thermococcus_A litoralis* | 523849 | *Thermococcus litoralis DSM 5473* | https://bacdive.dsmz.de/strain/16862 |
| GCA_000022545.1 | Hyperthermophiles | | 1845800 | Archaea | Methanobacteriota_B | Thermococci | Thermococcales | Thermococcaceae | Thermococcus_A | *Thermococcus_A sibiricus* | 604354 | *Thermococcus sibiricus MM 739* | https://doi.org/10.3390/ijms22189894 |
| GCA_000151105.2 | Hyperthermophiles | | 2064237 | Archaea | Methanobacteriota_B | Thermococci | Thermococcales | Thermococcaceae | Thermococcus_B | *Thermococcus_B barophilus* | 391623 | *Thermococcus barophilus* | https://doi.org/10.1099/00207713-49-2-351 |
| GCA_000025605.1 | Thermophiles | | 1500577 | Bacteria | Aquificota | Aquificae | Aquificales | Aquificaceae | Thermocrinis | *Thermocrinis albus* | 638303 | *Thermocrinis albus DSM 14484* | https://link.springer.com/article/10.1007/s00792-001-0259-y |
| GCA_000512735.1 | Hyperthermophiles | | 1521037 | Bacteria | Aquificota | Aquificae | Aquificales | Aquificaceae | Thermocrinis | *Thermocrinis ruber* | 75906 | *Thermocrinis ruber* | https://doi.org/10.1128/aem.64.10.3576-3583.1998 |
| GCA_900142435.1 | Thermophiles | | 1367921 | Bacteria | Aquificota | Aquificae | Aquificales | Aquificaceae | Thermocrinis_A | *Thermocrinis_A minervae* | 381751 | *Thermocrinis minervae* | https://pubmed.ncbi.nlm.nih.gov/19651724/ |
| GCA_003057965.1 | Thermophiles | Acidophiles | 1774794 | Bacteria | Thermodesulfobiota | Thermodesulfobiia | Thermodesulfobiales | Thermodesulfobiaceae | Thermodesulfobium | *Thermodesulfobium acidiphilum* | 1794699 | *Thermodesulfobium acidiphilum* | https://doi.org/10.1099/ijsem.0.001745 |
| GCA_000212395.1 | Thermophiles | | 1898865 | Bacteria | Thermodesulfobiota | Thermodesulfobiia | Thermodesulfobiales | Thermodesulfobiaceae | Thermodesulfobium | *Thermodesulfobium narugense* | 747365 | *Thermodesulfobium narugense DSM 14796* | https://doi.org/10.1007/s00792-003-0320-0 |
| GCA_000446015.1 | Hyperthermophiles | | 1750259 | Archaea | Thermoproteota | Thermoproteia | Thermofilales | Thermofilaceae | Thermofilum_B | *Thermofilum_B adornatum* | 1365176 | *Thermofilum adornatus 1910b* | https://doi.org/10.3389/fmicb.2019.02972 |
| GCA_000015225.1 | Hyperthermophiles | | 1813393 | Archaea | Thermoproteota | Thermoproteia | Thermofilales | Thermofilaceae | Thermofilum_B | *Thermofilum_B pendens_A* | 368408 | *Thermofilum pendens Hrk 5* | https://doi.org/10.1007/s00253-009-2190-6 |
| GCA_000264495.1 | Hyperthermophiles | | 1356318 | Archaea | Thermoproteota | Thermoproteia | Sulfolobales | Desulfurococcaceae | Thermogladius | *Thermogladius calderae* | 1184251 | *Thermogladius calderae 1633* | https://doi.org/10.1099/ijsem.0.000916 |
| GCA_000195915.1 | Thermophiles | Acidophiles | 1564906 | Archaea | Thermoplasmatota | Thermoplasmata | Thermoplasmatales | Thermoplasmataceae | Thermoplasma | *Thermoplasma acidophilum* | 273075 | *Thermoplasma acidophilum DSM 1728* | https://bacdive.dsmz.de/strain/17017 |
| GCA_000011185.1 | Thermophiles | Acidophiles | 1584804 | Archaea | Thermoplasmatota | Thermoplasmata | Thermoplasmatales | Thermoplasmataceae | Thermoplasma | *Thermoplasma volcanium* | 273116 | *Thermoplasma volcanium GSS1* | https://doi.org/10.1128/jb.178.3.902-905.1996 |
| GCA_000253055.1 | Hyperthermophiles | Acidophiles | 1841542 | Archaea | Thermoproteota | Thermoproteia | Thermoproteales | Thermoproteaceae | Thermoproteus | *Thermoproteus tenax* | 768679 | *Thermoproteus tenax Kra 1* | https://bacdive.dsmz.de/strain/17031 |
| GCA_000193375.1 | Thermophiles | Acidophiles | 1936063 | Archaea | Thermoproteota | Thermoproteia | Thermoproteales | Thermoproteaceae | Thermoproteus | *Thermoproteus uzoniensis* | 999630 | *Thermoproteus uzoniensis 768-20* | https://doi.org/10.1128/jb.00409-11 |
| GCA_000008545.1 | Hyperthermophiles | | 1860725 | Bacteria | Thermotogota | Thermotogae | Thermotogales | Thermotogaceae | Thermotoga | *Thermotoga maritima* | 243274 | *Thermotoga maritima MSB8* | https://bacdive.dsmz.de/strain/17060 |
| GCA_000018945.1 | Hyperthermophiles | | 1884562 | Bacteria | Thermotogota | Thermotogae | Thermotogales | Thermotogaceae | Thermotoga | *Thermotoga neapolitana* | 309803 | *Thermotoga neapolitana DSM 4359* | https://bacdive.dsmz.de/strain/17061 |

| | | | | | | | | | | | | | |
|---|---|---|---|---|---|---|---|---|---|---|---|---|---|
| GCA_000016785.1 | Hyperthermophiles | | 1823511 | Bacteria | Thermotogota | Thermotogae | Thermotogales | Thermotogaceae | Thermotoga | *Thermotoga petrophila* | 590168 | *Thermotoga petrophila* | https://doi.org/10.1099/00207713-51-5-1901 |
| GCA_000744885.1 | Thermophiles | | 2160855 | Bacteria | Deinococcota | Deinococci | Deinococcales | Thermaceae | Thermus | *Thermus amyloliquefaciens* | 1449080 | *Thermus amyloliquefaciens* | https://doi.org/10.1099/ijs.0.000289 |
| GCA_000423905.1 | Thermophiles | | 2165150 | Bacteria | Deinococcota | Deinococci | Deinococcales | Thermaceae | Thermus | *Thermus antranikianii* | 1123386 | *Thermus antranikianii DSM 12462* | https://bacdive.dsmz.de/strain/16730 |
| GCA_001280255.1 | Thermophiles | | 2248795 | Bacteria | Deinococcota | Deinococci | Deinococcales | Thermaceae | Thermus | *Thermus aquaticus* | 271 | *Thermus aquaticus* | https://doi.org/10.1128/jb.98.1.289-297.1969 |
| GCA_900102145.1 | Thermophiles | | 2442297 | Bacteria | Deinococcota | Deinococci | Deinococcales | Thermaceae | Thermus | *Thermus arciformis* | 482827 | *Thermus arciformis* | https://doi.org/10.1099/ijs.0.007690-0 |
| GCA_001880325.1 | Thermophiles | | 2388273 | Bacteria | Deinococcota | Deinococci | Deinococcales | Thermaceae | Thermus | *Thermus brockianus* | 56956 | *Thermus brockianus* | https://bacdive.dsmz.de/strain/161175 |
| GCA_003336745.1 | Thermophiles | | 2163786 | Bacteria | Deinococcota | Deinococci | Deinococcales | Thermaceae | Thermus | *Thermus caldifontis* | 1930763 | *Thermus caldifontis* | https://doi.org/10.1099/ijsem.0.002037 |
| GCA_000745065.1 | Thermophiles | | 2218114 | Bacteria | Deinococcota | Deinococci | Deinococcales | Thermaceae | Thermus | *Thermus calditerrae* | 1330700 | *Thermus calditerrae* | https://doi.org/10.1099/ijs.0.056838-0 |
| GCA_000376265.1 | Thermophiles | | 2225983 | Bacteria | Deinococcota | Deinococci | Deinococcales | Thermaceae | Thermus | *Thermus igniterrae* | 1123388 | *Thermus igniterrae ATCC 700962* | https://doi.org/10.1099/00207713-50-1-209 |
| GCA_000421625.1 | Thermophiles | Acidophiles | 2263010 | Bacteria | Deinococcota | Deinococci | Deinococcales | Thermaceae | Thermus | *Thermus islandicus* | 1123389 | *Thermus islandicus DSM 21543* | https://doi.org/10.1099/ijs.0.007013-0 |
| GCA_000373145.1 | Thermophiles | | 2260954 | Bacteria | Deinococcota | Deinococci | Deinococcales | Thermaceae | Thermus | *Thermus oshimai* | 1123390 | *Thermus oshimai DSM 12092* | https://doi.org/10.1099/00207713-46-2-403 |
| GCA_001535545.1 | Hyperthermophiles | | 2016098 | Bacteria | Deinococcota | Deinococci | Deinococcales | Thermaceae | Thermus | *Thermus parvatiensis* | 456163 | *Thermus parvatiensis* | https://pubmed.ncbi.nlm.nih.gov/26543260/ |
| GCA_000381045.1 | Thermophiles | | 2070699 | Bacteria | Deinococcota | Deinococci | Deinococcales | Thermaceae | Thermus | *Thermus scotoductus* | 1123391 | *Thermus scotoductus DSM 8553* | https://bacdive.dsmz.de/strain/16724 |
| GCA_000744175.1 | Thermophiles | | 2562314 | Bacteria | Deinococcota | Deinococci | Deinococcales | Thermaceae | Thermus | *Thermus tengchongensis* | 1214928 | *Thermus tengchongensis* | https://pubmed.ncbi.nlm.nih.gov/23104072/ |
| GCA_002964845.1 | Thermophiles | | 2261036 | Bacteria | Deinococcota | Deinococci | Deinococcales | Thermaceae | Thermus | *Thermus tenuipuniceus* | 2078690 | *Thermus tenuipuniceus* | https://bacdive.dsmz.de/strain/163768 |
| GCA_000091545.1 | Thermophiles | | 2116056 | Bacteria | Deinococcota | Deinococci | Deinococcales | Thermaceae | Thermus | *Thermus thermophilus* | 274 | *Thermus thermophilus* | https://doi.org/10.1016/j.ymben.2017.10.007 |
| GCA_002355995.1 | Thermophiles | | 2140665 | Bacteria | Deinococcota | Deinococci | Deinococcales | Thermaceae | Thermus | *Thermus thermophilus_C* | 274 | *Thermus thermophilus* | https://doi.org/10.1016/j.ymben.2017.10.007 |
| GCA_000771745.2 | Thermophiles | | 2386081 | Bacteria | Deinococcota | Deinococci | Deinococcales | Thermaceae | Thermus_A | *Thermus_A filiformis* | 276 | *Thermus filiformis* | https://bacdive.dsmz.de/strain/16723 |
| GCA_000376425.1 | Thermophiles | | 3609948 | Bacteria | Proteobacteria | Gammaproteobacteria | Burkholderiales | Thiobacillaceae | Thiobacillus | *Thiobacillus denitrificans* | 1123392 | *Thiobacillus denitrificans DSM 12475* | https://bacdive.dsmz.de/strain/6143 |
| GCA_000012745.1 | Thermophiles | | 2909809 | Bacteria | Proteobacteria | Gammaproteobacteria | Burkholderiales | Thiobacillaceae | Thiobacillus | *Thiobacillus denitrificans_B* | 292415 | *Thiobacillus denitrificans ATCC 25259* | https://bacdive.dsmz.de/strain/6143 |
| GCA_900113635.1 | | Alkaliphiles | 3083807 | Bacteria | Firmicutes_A | Clostridia | Peptostreptococcales | Tindalliaceae | Tindallia | *Tindallia magadiensis* | 69895 | *Tindallia magadiensis* | https://doi.org/10.1007/s002849900345 |
| GCA_000092425.1 | Thermophiles | Alkaliphiles | 3260398 | Bacteria | Deinococcota | Deinococci | Deinococcales | Trueperaceae | Truepera | *Truepera radiovictrix* | 649638 | *Truepera radiovictrix DSM 17093* | https://pubmed.ncbi.nlm.nih.gov/15927420/ |
| GCA_000148385.1 | Hyperthermophiles | Acidophiles | 2374137 | Archaea | Thermoproteota | Thermoproteia | Thermoproteales | Thermocladiaceae | Vulcanisaeta | *Vulcanisaeta distributa* | 572478 | *Vulcanisaeta distributa DSM 14429* | https://doi.org/10.1099/00207713-52-4-1097 |
| GCA_000190315.1 | Hyperthermophiles | | 2298983 | Archaea | Thermoproteota | Thermoproteia | Thermoproteales | Thermocladiaceae | Vulcanisaeta | *Vulcanisaeta moutnovskia* | 985053 | *Vulcanisaeta moutnovskia 768-28* | https://doi.org/10.1128/jb.00237-11 |
| GCA_014646555.1 | Hyperthermophiles | Acidophiles | 2424395 | Archaea | Thermoproteota | Thermoproteia | Thermoproteales | Thermocladiaceae | Vulcanisaeta | *Vulcanisaeta souniana* | 1293586 | *Vulcanisaeta souniana JCM 11219* | https://doi.org/10.1099/00207713-52-4-1097 |
| GCA_001748385.1 | Hyperthermophiles | Acidophiles | 2022594 | Archaea | Thermoproteota | Thermoproteia | Thermoproteales | Thermocladiaceae | Vulcanisaeta_B | *Vulcanisaeta_B thermophila* | 867917 | *Vulcanisaeta thermophila* | https://doi.org/10.1099/ijs.0.065862-0 |
| GCA_002094855.1 | Psychrophiles | | 4780723 | Bacteria | Bacteroidota | Bacteroidia | Flavobacteriales | Flavobacteriaceae | Zunongwangia | *Zunongwangia atlantica* | 1185767 | *Zunongwangia atlantica 22II14-10F7* | https://doi.org/10.1099/ijs.0.054007-0 |
| GCA_000023465.1 | Psychrophiles | | 5128187 | Bacteria | Bacteroidota | Bacteroidia | Flavobacteriales | Flavobacteriaceae | Zunongwangia | *Zunongwangia profunda* | 655815 | *Zunongwangia profunda SM-A87* | https://doi.org/10.1186%2F1471-2164-11-247 |
