## Supplementary Table S2 for "Environment and taxonomy shape the genomic signature of prokaryotic extremophiles"

| Supplementary Table S2-Dataset Metadata | | | |
|---|---|---|---|
| **Assembly Accession** | **Submission Date** | **Last Updated** | **Submission Organization** |
| GCF_000091545.1 | 2004/11/15 | 2020/03/06 | National Institute of Advanced Industrial Science and Technology, Japan |
| GCF_000236925.1 | 2011/11/10 | 2013/09/23 | The Wellcome Trust Sanger Institute |
| GCF_000025685.1 | 2010/03/23 | 2016/11/29 | TIGR |
| GCF_002945325.1 | 2018/02/12 | 2018/03/02 | Goettingen Genomics Laboratory |
| GCF_000196795.1 | 2010/06/21 | 2016/11/29 | Division of Environmental Science and Ecological Engineering, Korea University |
| GCF_000009965.1 | 2005/01/05 | 2016/11/29 | Kyoto University, Japan |
| GCF_009662475.1 | 2019/11/14 | 2019/11/16 | Shandong University(Qingdao) |
| GCF_000022325.1 | 2009/02/04 | 2016/11/29 | US DOE Joint Genome Institute (JGI-PGF) |
| GCF_000027145.1 | 2010/02/18 | 2016/11/29 | PathoGenoMik |
| GCF_000816085.1 | 2015/01/07 | 2015/01/23 | University of Missouri - Columbia |
| GCF_000304355.2 | 2014/02/04 | 2015/07/16 | irena.maus |
| GCF_001277215.2 | 2016/01/13 | 2016/01/17 | Nestle Institute of Health Sciences |
| GCF_001975665.1 | 2017/01/30 | 2017/02/03 | Kyung Hee University |
| GCF_000014885.1 | 2008/11/13 | 2016/11/29 | James D. Watson Institute of Genome Sciences |
| GCF_000016065.1 | 2007/03/07 | 2016/11/29 | US DOE Joint Genome Institute |
| GCF_000091665.1 | 1999/12/22 | 2016/06/03 | TIGR |
| GCF_000018465.1 | 2007/11/15 | 2019/08/31 | US DOE Joint Genome Institute (JGI-PGF) |
| GCF_000195915.1 | 2003/05/06 | 2016/11/29 | Max-Plank-Institute |
| GCF_000172995.2 | 2010/11/22 | 2016/11/29 | US DOE Joint Genome Institute (JGI-PGF) |
| GCF_000007225.1 | 2002/01/24 | 2019/01/29 | UCLA/CalTech |
| GCF_001277235.1 | 2015/09/01 | 2015/09/02 | Nestle Institute of Health Sciences |
| GCF_000202835.1 | 2011/03/01 | 2017/04/07 | National Institute of Technology and Evaluation |
| GCF_000024845.1 | 2009/11/03 | 2016/11/29 | US DOE Joint Genome Institute (JGI-PGF) |
| GCF_000023965.1 | 2009/09/03 | 2016/11/29 | US DOE Joint Genome Institute (JGI-PGF) |
| GCF_000024285.1 | 2009/09/09 | 2016/11/29 | US DOE Joint Genome Institute (JGI-PGF) |
| GCF_000022205.1 | 2009/02/04 | 2021/04/08 | US DOE Joint Genome Institute (JGI-PGF) |
| GCF_003012915.1 | 2018/03/21 | 2018/03/23 | Illinois Institute of Technology |

| | | | |
|---|---|---|---|
| GCF_008931805.1 | 2019/10/09 | 2019/10/13 | University hospital Essen |
| GCF_000968535.2 | 2011/10/14 | 2021/12/27 | CEA/DSV/IG/Genoscope |
| GCF_000230715.2 | 2012/12/10 | 2012/12/21 | DOE Joint Genome Institute |
| GCF_000196895.1 | 2010/06/23 | 2016/11/29 | Kyung Hee University |
| GCF_000017945.1 | 2007/08/29 | 2016/11/29 | US DOE Joint Genome Institute |
| GCF_001277175.1 | 2015/09/01 | 2015/09/02 | Nestle Institute of Health Sciences |
| GCF_000014185.1 | 2006/06/09 | 2016/11/29 | DOE Joint Genome Institute |
| GCF_000698785.1 | 2014/06/06 | 2016/12/26 | University of Vienna |
| GCF_000019085.1 | 2008/02/05 | 2021/06/08 | US DOE Joint Genome Institute |
| GCF_000018945.1 | 2009/01/27 | 2016/11/29 | Genotech Corp. |
| GCF_000018365.1 | 2008/11/06 | 2016/11/29 | Korea Ocean Research & Development Institute |
| GCF_000011125.1 | 2006/10/11 | 2019/08/31 | NITE |
| GCF_001277195.1 | 2015/09/01 | 2015/09/02 | Nestle Institute of Health Sciences |
| GCF_000016545.1 | 2007/04/19 | 2016/11/29 | US DOE Joint Genome Institute |
| GCF_000235405.2 | 2012/04/02 | 2016/11/30 | DOE Joint Genome Institute |
| GCF_000151105.2 | 2010/12/17 | 2016/11/29 | Moore Foundation |
| GCF_000195935.2 | 2003/05/06 | 2014/02/03 | Genoscope |
| GCF_002075795.1 | 2017/03/31 | 2017/04/03 | Korea Polar Research Institute |
| GCF_014784055.1 | 2020/09/29 | 2020/09/29 | Kyung Hee University |
| GCF_001971705.1 | 2017/01/26 | 2017/02/15 | Heilongjiang Academy |
| GCF_003344925.1 | 2018/07/30 | 2018/08/01 | Shandong University |
| GCF_001634285.1 | 2016/04/29 | 2016/08/18 | Los Alamos National Laboratory |
| GCF_000470655.1 | 2013/08/01 | 2021/02/10 | MPI BREMEN |
| GCF_003019255.1 | 2018/03/26 | 2018/03/29 | Illinois Institute of Technology |
| GCF_000214725.1 | 2011/05/24 | 2016/11/29 | US DOE Joint Genome Institute |
| GCF_000763575.1 | 2014/10/06 | 2015/07/25 | Georg-August-University Goettingen, Genomic and Applied Microbiology, Goettingen Genomics Laboratory |
| GCF_003574215.1 | 2018/06/21 | 2018/09/20 | Department of Biotechnology, The University of Tokyo |
| GCF_001412615.1 | 2015/10/26 | 2015/10/27 | Kyung Hee University |
| GCF_001889405.1 | 2016/12/06 | 2016/12/18 | UBO |

| | | | |
|---|---|---|---|
| GCF_002078355.1 | 2017/04/04 | 2017/04/05 | Max Planck Institute for Chemical Ecology |
| GCF_000023465.1 | 2010/04/21 | 2016/11/29 | ShanDong University |
| GCF_000970285.1 | 2015/04/07 | 2015/06/26 | University of Illinois at Urbana-Champaign |
| GCF_000015505.1 | 2007/01/04 | 2016/11/29 | DOE Joint Genome Institute |
| GCF_000022565.1 | 2009/03/30 | 2016/11/29 | J. Craig Venter Institute |
| GCF_000025625.1 | 2010/02/24 | 2016/11/29 | US DOE Joint Genome Institute (JGI-PGF) |
| GCF_000012305.1 | 2005/07/20 | 2016/11/29 | DOE Joint Genome Institute |
| GCF_000013725.1 | 2006/04/10 | 2016/11/29 | DOE Joint Genome Institute |
| GCF_000145215.1 | 2010/08/11 | 2016/11/29 | US DOE Joint Genome Institute |
| GCF_000025505.1 | 2010/02/16 | 2016/11/29 | US DOE Joint Genome Institute |
| GCF_000504085.1 | 2013/10/23 | 2015/11/23 | National Institute of Technology and Evaluation |
| GCF_000223395.1 | 2011/08/12 | 2012/11/23 | DOE Joint Genome Institute |
| GCF_000007185.1 | 2002/04/03 | 2019/08/31 | Fidelity Systems |
| GCF_000591035.1 | 2013/09/27 | 2019/09/01 | Laboratory of Marine Microbiology, Division of Applied Biosciences, Graduate School of Agriculture, Kyoto University |
| GCF_000019185.1 | 2008/02/07 | 2016/11/29 | US DOE Joint Genome Institute |
| GCF_001277255.1 | 2015/09/01 | 2015/09/02 | Nestle Institute of Health Sciences |
| GCF_000025325.1 | 2010/01/19 | 2016/11/29 | US DOE Joint Genome Institute (JGI-PGF) |
| GCF_000013445.1 | 2006/02/17 | 2016/11/29 | DOE Joint Genome Institute |
| GCF_000969885.1 | 2015/04/07 | 2015/06/26 | University of Illinois at Urbana-Champaign |
| GCF_000226295.1 | 2011/09/23 | 2016/11/30 | The Pennsylvania State University |
| GCF_000148385.1 | 2010/09/22 | 2016/11/29 | US DOE Joint Genome Institute (JGI-PGF) |
| GCF_000016385.1 | 2007/04/16 | 2019/01/29 | DOE Joint Genome Institute |
| GCF_000022365.1 | 2009/06/12 | 2016/11/29 | IGM/ Universit Paris-Sud, France |
| GCF_000020965.1 | 2008/09/26 | 2016/11/29 | The J. Craig Venter Institute |
| GCF_000253055.1 | 2011/10/06 | 2012/10/11 | University of Duisburg-Essen, Faculty of Chemistry, Molecular Enzyme Technology and Biochemistry |
| GCF_000016785.1 | 2007/05/18 | 2016/11/29 | US DOE Joint Genome Institute |
| GCF_000186365.1 | 2011/01/20 | 2016/11/29 | US DOE Joint Genome Institute (JGI-PGF) |
| GCF_000166095.1 | 2010/11/05 | 2016/11/29 | US DOE Joint Genome Institute (JGI-PGF) |

| | | | |
|---|---|---|---|
| GCF_003430825.1 | 2018/08/28 | 2018/09/02 | C.N.R. - Institute for Coastal Marine Environment (IAMC) |
| GCF_003201835.2 | 2020/03/10 | 2020/04/16 | North Carolina State University |
| GCF_000585495.1 | 2014/03/05 | 2015/02/16 | Institute de Genetique et Microbiologie |
| GCF_001535545.1 | 2016/01/25 | 2016/01/26 | University of Delhi |
| GCF_000021645.1 | 2008/12/12 | 2020/03/06 | US DOE Joint Genome Institute (JGI-PGF) |
| GCF_000875775.1 | 2015/02/13 | 2019/09/01 | University of Vienna |
| GCF_000512735.1 | 2014/01/03 | 2015/02/04 | JGI |
| GCF_000019525.1 | 2008/03/13 | 2016/11/29 | US DOE Joint Genome Institute |
| GCF_000018025.1 | 2007/09/18 | 2016/11/29 | US DOE Joint Genome Institute (JGI-PGF) |
| GCF_000018285.1 | 2007/10/04 | 2016/11/29 | US DOE Joint Genome Institute |
| GCF_000970085.1 | 2015/04/07 | 2015/06/26 | University of Illinois at Urbana-Champaign |
| GCF_000091325.1 | 2010/04/01 | 2016/11/29 | Nara Institute of Science and Technology |
| GCF_000255115.2 | 2012/06/15 | 2012/10/18 | DOE Joint Genome Institute |
| GCF_000015285.1 | 2006/12/22 | 2016/11/29 | DOE Joint Genome Institute |
| GCF_000013765.1 | 2006/04/10 | 2016/11/29 | DOE Joint Genome Institute |
| GCF_000153485.2 | 2012/10/09 | 2018/04/04 | The Gordon and Betty Moore Foundation Marine Microbiology Initiative |
| GCF_000328685.1 | 2012/12/28 | 2012/12/31 | JGI |
| GCF_000242595.2 | 2012/04/02 | 2016/11/30 | DOE Joint Genome Institute |
| GCF_000092425.1 | 2010/05/28 | 2016/11/29 | US DOE Joint Genome Institute (JGI-PGF) |
| GCF_000092125.1 | 2010/05/26 | 2016/11/29 | US DOE Joint Genome Institute (JGI-PGF) |
| GCF_000023945.1 | 2009/08/27 | 2016/11/29 | US DOE Joint Genome Institute (JGI-PGF) |
| GCF_000021965.1 | 2009/01/05 | 2016/11/29 | US DOE Joint Genome Institute (JGI-PGF) |
| GCF_000166775.1 | 2010/11/17 | 2016/11/29 | US DOE Joint Genome Institute (JGI-PGF) |
| GCF_000147875.1 | 2010/09/17 | 2019/12/27 | US DOE Joint Genome Institute (JGI-PGF) |
| GCF_000327485.1 | 2012/12/21 | 2012/12/21 | DOE Joint Genome Institute |
| GCF_000147695.2 | 2011/08/23 | 2018/04/04 | US DOE Joint Genome Institute (JGI-PGF) |
| GCF_000166355.1 | 2010/11/10 | 2016/11/29 | US DOE Joint Genome Institute |
| GCF_000011205.1 | 2004/05/11 | 2016/11/29 | NITE |
| GCF_000026045.1 | 2005/09/28 | 2016/11/29 | Max Planck Institute |
| GCF_000015825.1 | 2007/02/16 | 2016/11/29 | DOE Joint Genome Institute |

| | | | |
|---|---|---|---|
| GCF_000015025.1 | 2006/11/09 | 2016/11/29 | DOE Joint Genome Institute |
| GCF_000166335.1 | 2010/11/10 | 2016/11/29 | US DOE Joint Genome Institute |
| GCF_000251105.1 | 2012/03/07 | 2016/11/30 | China Agricultural University |
| GCF_000190315.1 | 2011/02/22 | 2016/11/29 | Centre Bioengineering RAS |
| GCF_000246985.2 | 2013/08/13 | 2015/07/20 | New England Biolabs, Inc. |
| GCF_000217995.1 | 2011/06/13 | 2016/11/30 | US DOE Joint Genome Institute (JGI-PGF) |
| GCF_000024605.1 | 2009/10/08 | 2016/11/29 | US DOE Joint Genome Institute |
| GCF_000018305.1 | 2007/10/05 | 2016/11/29 | US DOE Joint Genome Institute (JGI-PGF) |
| GCF_000217815.1 | 2011/06/10 | 2016/11/30 | US DOE Joint Genome Institute (JGI-PGF) |
| GCF_000025865.1 | 2010/03/31 | 2016/11/29 | US DOE Joint Genome Institute (JGI-PGF) |
| GCF_000015805.1 | 2007/02/27 | 2020/10/25 | DOE Joint Genome Institute |
| GCF_000152265.2 | 2013/06/26 | 2013/06/29 | DOE Joint Genome Institute |
| GCF_000212395.1 | 2011/05/05 | 2016/11/29 | US DOE Joint Genome Institute (JGI-PGF) |
| GCF_000014945.1 | 2006/10/25 | 2020/06/19 | US DOE Joint Genome Institute |
| GCF_000725425.1 | 2014/07/14 | 2015/06/26 | The third institute of oceanography of SOA,PRChina |
| GCF_000214415.1 | 2011/05/23 | 2016/11/29 | US DOE Joint Genome Institute |
| GCF_000022545.1 | 2009/06/29 | 2016/11/29 | Centre |
| GCF_000204925.1 | 2011/04/19 | 2016/11/29 | Environmental Microbiology Research Center (EMRC) |
| GCF_000015205.1 | 2006/12/20 | 2016/11/29 | DOE Joint Genome Institute |
| GCF_000015225.1 | 2006/12/18 | 2016/11/29 | DOE Joint Genome Institute |
| GCF_000019805.1 | 2008/03/27 | 2019/08/31 | US DOE Joint Genome Institute (JGI-PGF) |
| GCF_000024625.1 | 2009/10/15 | 2016/11/29 | US DOE Joint Genome Institute (JGI-PGF) |
| GCF_000215995.1 | 2011/06/07 | 2016/11/30 | Shanghai JiaoTong University |
| GCF_000015145.1 | 2007/01/22 | 2016/11/29 | Univ. Copenhagen |
| GCF_000179575.2 | 2011/06/06 | 2018/04/04 | US DOE Joint Genome Institute (JGI-PGF) |
| GCF_000145295.1 | 2010/08/11 | 2016/11/29 | Georg-August-University Goettingen |
| GCF_000011185.1 | 2004/05/11 | 2016/11/29 | National Institute of Advanced Industrial Science and Technology, Japan |
| GCF_000015945.1 | 2007/02/21 | 2020/05/06 | US DOE Joint Genome Institute |
| GCF_000017185.1 | 2007/06/29 | 2016/11/29 | US DOE Joint Genome Institute (JGI-PGF) |
| GCF_000317795.1 | 2012/12/10 | 2012/12/10 | JGI |

| | | | |
|---|---|---|---|
| GCF_000025605.1 | 2010/02/19 | 2016/11/29 | US DOE Joint Genome Institute (JGI-PGF) |
| GCF_000144915.1 | 2010/08/06 | 2016/11/29 | Centre Bioengineering RAS |
| GCF_000023985.1 | 2009/08/28 | 2016/11/29 | US DOE Joint Genome Institute (JGI-PGF) |
| GCF_001481685.1 | 2015/12/23 | 2016/12/26 | Oak Ridge National Laboratory |
| GCF_002005305.1 | 2017/02/21 | 2017/02/23 | Shanghai Jiao Tong University |
| GCF_017298635.1 | 2021/03/08 | 2022/05/13 | Institute of Microbiology |
| GCF_007567505.1 | 2019/07/29 | 2019/08/03 | Korea Institute of Ocean Science and Technology |
| GCF_003855395.1 | 2018/12/02 | 2018/12/04 | Korea Polar Research Institute |
| GCF_000969905.1 | 2015/04/07 | 2015/06/26 | University of Illinois at Urbana-Champaign |
| GCF_900465055.1 | 2018/06/24 | 2018/07/13 | Genoscope CEA |
| GCF_002499975.2 | 2020/05/11 | 2020/05/13 | NCAOR |
| GCF_001043175.1 | 2015/06/30 | 2015/07/02 | Beijing Institute of Genomics, Chinese Academy of Sciences |
| GCF_002156705.1 | 2017/05/23 | 2017/05/25 | Chinese Academy of Agricultural Sciences |
| GCF_003352165.1 | 2018/08/01 | 2018/08/03 | institute of microbiology |
| GCF_001761365.1 | 2016/10/12 | 2016/10/14 | Korea University |
| GCF_009938015.1 | 2020/01/17 | 2020/02/03 | Hokkaido University |
| GCF_009392895.1 | 2019/10/30 | 2019/11/01 | Jiangsu University |
| GCF_002952775.1 | 2018/02/01 | 2018/09/26 | C.N.R. - Institute for Coastal Marine Environment (IAMC) |
| GCF_018228765.1 | 2021/05/03 | 2021/05/11 | Incheon National University |
| GCF_003568865.1 | 2018/09/13 | 2018/09/18 | South China Sea Institute of Oceanology, Chinese Academy of Sciences |
| GCF_009729055.1 | 2019/12/04 | 2019/12/09 | North Carolina State University |
| GCF_013388255.1 | 2020/07/04 | 2020/07/04 | National Chung Hsing University |
| GCF_002754935.1 | 2017/11/06 | 2017/11/09 | Peking University Shenzhen Graduate School |
| GCF_900186985.1 | 2017/08/15 | 2021/12/15 | SC |
| GCF_003201675.2 | 2020/03/10 | 2020/04/16 | North Carolina State University |
| GCF_002813695.1 | 2017/12/07 | 2017/12/09 | Japan Agency for Marine-Earth Science and technology |
| GCF_000234805.1 | 2011/11/10 | 2019/08/31 | Centre Bioengineering RAS |
| GCF_003201765.2 | 2020/03/10 | 2020/04/16 | North Carolina State University |
| GCF_009729015.1 | 2019/12/04 | 2019/12/09 | North Carolina State University |
| GCF_000828675.1 | 2014/03/21 | 2015/01/30 | National Institute of Technology and Evaluation |

| | | | |
|---|---|---|---|
| GCF_013343295.1 | 2020/06/15 | 2020/06/15 | Institute of microbiology |
| GCF_900198835.1 | 2017/10/17 | 2017/10/28 | Universite de Bretagne Occidentale |
| GCF_000769655.1 | 2014/10/28 | 2015/08/04 | Laboratory of Marine Oceanography |
| GCF_002355215.1 | 2015/02/05 | 2019/01/29 | Tsurumi University, School of Dental Medicine |
| GCF_000828655.1 | 2014/03/21 | 2015/01/30 | National Institute of Technology and Evaluation |
| GCF_009729035.1 | 2019/12/04 | 2019/12/09 | North Carolina State University |
| GCF_000265525.1 | 2012/06/11 | 2016/11/30 | Kyung Hee University |
| GCF_001647085.1 | 2016/05/18 | 2016/08/19 | Institut National des Sciences Appliquees de Lyon |
| GCF_002214465.1 | 2017/07/05 | 2017/07/07 | Ecole Normale Superieure de Lyon |
| GCF_000816105.1 | 2015/01/07 | 2016/03/28 | University of Waterloo |
| GCF_002214565.1 | 2017/07/05 | 2017/07/07 | Ecole Normale Superieure de Lyon |
| GCF_002214365.1 | 2017/07/05 | 2017/07/07 | Ecole Normale Superieure de Lyon |
| GCF_001592435.1 | 2016/03/15 | 2016/03/28 | Ecole Normale Superieure de Lyon |
| GCF_000956175.1 | 2015/03/19 | 2015/06/26 | University of Vienna |
| GCF_002214485.1 | 2017/07/05 | 2017/07/07 | Ecole Normale Superieure de Lyon |
| GCF_003057965.1 | 2018/04/19 | 2018/04/25 | Immanuel Kant Baltic Federal University |
| GCF_001006045.1 | 2015/05/11 | 2015/06/26 | Michigan State University |
| GCF_002214385.1 | 2017/07/05 | 2017/07/07 | Ecole Normale Superieure de Lyon |
| GCF_000739065.1 | 2014/08/12 | 2015/06/26 | Kyung Hee University |
| GCF_013340765.1 | 2019/05/09 | 2020/06/14 | Japan Collection of Microorganisms |
| GCF_000264495.1 | 2012/05/25 | 2016/11/30 | Centre Bioengineering RAS |
| GCF_002215585.1 | 2017/07/06 | 2017/07/08 | Hubei Polytechnic University |
| GCF_003534205.1 | 2017/03/13 | 2019/09/18 | Technische Universitaet Muenchen |
| GCF_000970265.1 | 2015/04/07 | 2015/06/26 | University of Illinois at Urbana-Champaign |
| GCF_000235565.1 | 2011/11/16 | 2016/11/30 | State Key Laboratory of Microbial Resources, Institute of Microbiology, Chinese Academy of Sciences |
| GCF_007475525.1 | 2019/07/25 | 2019/07/31 | University of Bergen |
| GCF_000193375.1 | 2011/03/28 | 2016/11/29 | Centre Bioengineering RAS |
| GCF_001606025.1 | 2016/04/04 | 2016/04/07 | Korea Polar Research Institute |
| GCF_001298525.1 | 2015/09/24 | 2016/03/28 | University of Malaya |

| | | | |
|---|---|---|---|
| GCF_002116695.1 | 2017/05/01 | 2019/09/01 | Cental South University, China |
| GCF_002761295.1 | 2017/11/07 | 2017/11/09 | Macumba |
| GCF_004117075.1 | 2019/01/29 | 2019/02/01 | Kyungpook National University |
| GCF_900176005.1 | 2017/04/06 | 2017/04/17 | DOE - JOINT GENOME INSTITUTE |
| GCF_009735625.1 | 2019/11/28 | 2019/12/11 | Georg-August-University Goettingen |
| GCF_900142435.1 | 2016/12/02 | 2017/03/18 | DOE - JOINT GENOME INSTITUTE |
| GCF_000243255.1 | 2012/01/24 | 2016/11/30 | JGI |
| GCF_000275865.1 | 2012/07/13 | 2012/07/23 | DOE Joint Genome Institute |
| GCF_004346035.1 | 2019/03/11 | 2019/03/16 | DOE Joint Genome Institute |
| GCF_900458435.1 | 2018/08/01 | 2018/08/07 | SC |
| GCF_900101915.1 | 2016/10/21 | 2017/07/29 | DOE - JOINT GENOME INSTITUTE |
| GCF_900176145.1 | 2017/04/07 | 2017/04/17 | DOE - JOINT GENOME INSTITUTE |
| GCF_014196195.1 | 2020/08/14 | 2020/08/17 | DOE Joint Genome Institute |
| GCF_900107035.1 | 2016/10/22 | 2016/12/22 | DOE - JOINT GENOME INSTITUTE |
| GCF_900129125.1 | 2016/12/03 | 2016/12/18 | DOE - JOINT GENOME INSTITUTE |
| GCF_000376425.1 | 2013/04/20 | 2014/07/12 | DOE Joint Genome Institute |
| GCF_006716115.1 | 2019/07/08 | 2019/07/12 | DOE Joint Genome Institute |
| GCF_004123295.1 | 2019/01/30 | 2019/02/07 | Sun Yat-sen University |
| GCF_010119205.1 | 2020/02/03 | 2020/02/10 | Anhui Normal University |
| GCF_900100875.1 | 2016/10/21 | 2017/07/29 | DOE - JOINT GENOME INSTITUTE |
| GCF_003185895.1 | 2018/06/04 | 2018/06/07 | Institute of Microbiology, Chinese Academy of Sciences |
| GCF_900110455.1 | 2016/10/31 | 2016/12/23 | DOE - JOINT GENOME INSTITUTE |
| GCF_000771745.2 | 2015/02/24 | 2015/03/20 | Centro Nacional de Pesquisa em Energia e Materiais |
| GCF_900111935.1 | 2016/11/02 | 2017/07/29 | DOE - JOINT GENOME INSTITUTE |
| GCF_010119195.1 | 2020/02/03 | 2020/02/10 | Anhui Normal University |
| GCF_003259835.1 | 2018/06/21 | 2018/06/23 | DOE Joint Genome Institute |
| GCF_002954685.1 | 2018/02/27 | 2018/03/04 | Atmosphere and Ocean Research Institute, The University of Tokyo |
| GCF_003185915.1 | 2018/06/04 | 2018/06/07 | Institute of Microbiology, Chinese Academy of Sciences |
| GCF_004216855.1 | 2019/02/20 | 2019/02/22 | DOE Joint Genome Institute |
| GCF_003797885.1 | 2018/11/19 | 2018/11/26 | Korea Polar Research Institute |

| | | | |
|---|---|---|---|
| GCF_900143245.1 | 2016/12/03 | 2016/12/18 | DOE - JOINT GENOME INSTITUTE |
| GCF_003312425.1 | 2018/07/10 | 2018/07/15 | Konggi University |
| GCF_002954665.1 | 2018/02/27 | 2018/03/04 | Atmosphere and Ocean Research Institute, The University of Tokyo |
| GCF_009791395.1 | 2019/12/22 | 2020/01/02 | CSIR-IMTECH |
| GCF_003219795.1 | 2018/06/12 | 2018/06/16 | Institute of Microbiology, Chinese Academy of Sciences |
| GCF_002836945.1 | 2017/12/12 | 2017/12/14 | National Institute for Communicable Disease Control and Prevention, China CDC |
| GCF_900102145.1 | 2016/10/21 | 2017/07/29 | DOE - JOINT GENOME INSTITUTE |
| GCF_001402945.1 | 2015/10/22 | 2015/10/23 | RIPCM |
| GCF_000383975.1 | 2013/04/26 | 2014/07/11 | JGI |
| GCF_000337795.1 | 2013/02/04 | 2013/02/07 | University of California, Davis |
| GCF_000337815.1 | 2013/02/04 | 2013/02/07 | University of California, Davis |
| GCF_900176435.1 | 2017/04/08 | 2021/06/08 | DOE - JOINT GENOME INSTITUTE |
| GCF_900111075.1 | 2016/10/31 | 2016/12/22 | DOE - JOINT GENOME INSTITUTE |
| GCF_900108085.1 | 2016/10/24 | 2018/02/10 | DOE - JOINT GENOME INSTITUTE |
| GCF_000368025.1 | 2013/04/19 | 2013/05/09 | Broad Institute |
| GCF_900175965.1 | 2017/04/07 | 2017/04/17 | DOE - JOINT GENOME INSTITUTE |
| GCF_000423905.1 | 2013/07/11 | 2014/07/12 | DOE Joint Genome Institute |
| GCF_000337915.1 | 2013/02/04 | 2013/02/07 | University of California, Davis |
| GCF_000337555.1 | 2013/02/04 | 2013/02/07 | University of California, Davis |
| GCF_000298075.1 | 2012/09/21 | 2014/07/12 | URMITE |
| GCF_000688455.1 | 2014/04/08 | 2014/07/11 | DOE Joint Genome Institute |
| GCF_007997305.1 | 2019/08/14 | 2019/08/17 | University of Tasmania |
| GCF_004217335.1 | 2019/02/20 | 2019/02/22 | DOE Joint Genome Institute |
| GCF_008369445.1 | 2019/09/10 | 2019/09/16 | Chung-Ang University |
| GCF_004216575.1 | 2019/02/20 | 2019/02/22 | DOE Joint Genome Institute |
| GCF_001571405.1 | 2016/01/19 | 2016/08/18 | Bioproduction Research Institute, National Institute of Advanced Industrial Science and Technology (AIST) |
| GCF_003350545.1 | 2018/07/31 | 2018/08/02 | DOE Joint Genome Institute |
| GCF_003148585.1 | 2018/05/22 | 2018/05/26 | DOE Joint Genome Institute |

| | | | |
|---|---|---|---|
| GCF_900116805.1 | 2016/11/02 | 2016/12/23 | DOE - JOINT GENOME INSTITUTE |
| GCF_008245225.1 | 2019/09/03 | 2019/09/16 | NCMR- NCCS |
| GCF_007988865.1 | 2019/08/01 | 2019/08/16 | National Institute of Technology and Evaluation |
| GCF_000934755.1 | 2015/02/26 | 2015/06/09 | DOE Joint Genome Institute |
| GCF_001950255.1 | 2016/12/09 | 2017/01/16 | Laboratory of Marine Microbiology, Graduate School of Agriculture, Kyoto University |
| GCF_000504205.1 | 2013/12/05 | 2013/12/10 | JGI |
| GCF_000744825.1 | 2014/08/28 | 2014/12/23 | DOE Joint Genome Institute |
| GCF_000364845.1 | 2013/04/15 | 2013/05/09 | First Institute of Oceanography, State Oceanic Administration of China |
| GCF_000220175.1 | 2011/07/07 | 2016/11/30 | Korea Research Institute of Bioscience and Biotechnology |
| GCF_000183545.2 | 2012/11/05 | 2016/04/19 | US DOE Joint Genome Institute (JGI-PGF) |
| GCF_000258515.1 | 2012/04/11 | 2014/07/12 | Korea Research Institute of Bioscience and Biotechnology |
| GCF_000153225.1 | 2006/05/04 | 2016/11/29 | The Gordon and Betty Moore Foundation Marine Microbiology Initiative |
| GCF_000378345.1 | 2013/04/23 | 2014/07/11 | DOE Joint Genome Institute |
| GCF_000419685.1 | 2013/07/09 | 2014/07/11 | DOE Joint Genome Institute |
| GCF_000379085.1 | 2013/04/23 | 2014/07/11 | JGI |
| GCF_000336755.1 | 2013/02/04 | 2013/02/07 | University of California, Davis |
| GCF_000347775.1 | 2013/03/21 | 2013/03/29 | University of Manchester |
| GCF_000373205.1 | 2013/04/23 | 2014/07/11 | DOE Joint Genome Institute |
| GCF_000425505.1 | 2013/07/11 | 2014/07/11 | DOE Joint Genome Institute |
| GCF_000336615.1 | 2013/02/04 | 2013/02/07 | University of California, Davis |
| GCF_000421625.1 | 2013/07/09 | 2014/07/12 | DOE Joint Genome Institute |
| GCF_000337675.1 | 2013/02/04 | 2013/02/07 | University of California, Davis |
| GCF_000518605.1 | 2014/01/13 | 2014/07/11 | DOE Joint Genome Institute |
| GCF_000966265.1 | 2015/03/30 | 2015/06/26 | DOE Joint Genome Institute |
| GCF_000336995.1 | 2013/02/04 | 2013/02/07 | University of California, Davis |
| GCF_000472905.1 | 2013/09/30 | 2014/07/11 | DOE Joint Genome Institute |
| GCF_000471625.1 | 2013/09/24 | 2013/09/26 | Centre for Cellular and Molecular Biology |
| GCF_001598235.1 | 2016/03/11 | 2016/08/21 | National Institute of Technology and Evaluation |
| GCF_000337735.1 | 2013/02/04 | 2013/02/07 | University of California, Davis |

| | | | |
|---|---|---|---|
| GCF_000276805.1 | 2012/07/09 | 2014/07/11 | Korea Polar Research Institute |
| GCF_000337575.1 | 2013/02/04 | 2013/02/07 | University of California, Davis |
| GCF_000425485.1 | 2013/07/11 | 2014/07/11 | DOE Joint Genome Institute |
| GCF_000423105.1 | 2013/07/11 | 2014/07/11 | DOE Joint Genome Institute |
| GCF_000337375.1 | 2013/02/04 | 2013/02/07 | University of California, Davis |
| GCF_000430585.1 | 2013/07/16 | 2014/07/11 | DOE Joint Genome Institute |
| GCF_000378465.1 | 2013/04/23 | 2014/07/11 | DOE Joint Genome Institute |
| GCF_000428725.1 | 2013/07/15 | 2014/07/11 | DOE Joint Genome Institute |
| GCF_001552655.1 | 2016/02/06 | 2016/08/19 | National Institute of Technology and Evaluation |
| GCF_000337135.1 | 2013/02/04 | 2013/02/07 | University of California, Davis |
| GCF_000337475.1 | 2013/02/04 | 2013/02/07 | University of California, Davis |
| GCF_000336935.1 | 2013/02/04 | 2013/02/07 | University of California, Davis |
| GCF_000518805.1 | 2014/01/13 | 2014/07/11 | DOE Joint Genome Institute |
| GCF_000337035.1 | 2013/02/04 | 2013/02/07 | University of California, Davis |
| GCF_000337695.1 | 2013/02/04 | 2013/02/07 | University of California, Davis |
| GCF_000376225.1 | 2013/04/20 | 2014/07/11 | DOE Joint Genome Institute |
| GCF_000381745.1 | 2013/04/24 | 2014/07/11 | DOE Joint Genome Institute |
| GCF_001552255.1 | 2016/02/03 | 2016/08/19 | National Institute of Technology and Evaluation |
| GCF_000482725.1 | 2013/10/25 | 2014/07/11 | DOE Joint Genome Institute |
| GCF_000429525.1 | 2013/07/15 | 2014/07/11 | DOE Joint Genome Institute |
| GCF_000376265.1 | 2013/04/20 | 2014/07/12 | DOE Joint Genome Institute |
| GCF_001544355.1 | 2016/01/29 | 2016/08/18 | National Institute of Technology and Evaluation |
| GCF_000429345.1 | 2013/09/05 | 2014/07/11 | DOE Joint Genome Institute |
| GCF_000430045.1 | 2013/07/15 | 2014/07/11 | JGI |
| GCF_001552675.1 | 2016/02/06 | 2016/03/28 | National Institute of Technology and Evaluation |
| GCF_000423425.1 | 2013/07/11 | 2014/07/11 | DOE Joint Genome Institute |
| GCF_000243455.1 | 2012/01/23 | 2016/11/30 | JGI |
| GCF_000420245.1 | 2013/07/09 | 2014/07/11 | DOE Joint Genome Institute |
| GCF_004365915.1 | 2019/03/22 | 2019/03/27 | DOE Joint Genome Institute |
| GCF_001953745.1 | 2017/01/18 | 2017/01/20 | National Chung Hsing University |

| | | | |
|---|---|---|---|
| GCF_001542905.1 | 2016/01/29 | 2016/08/18 | Silla university |
| GCF_000955725.1 | 2015/03/17 | 2015/03/19 | DOE Joint Genome Institute |
| GCF_900142385.1 | 2016/12/03 | 2016/12/18 | DOE - JOINT GENOME INSTITUTE |
| GCF_017873855.1 | 2021/04/08 | 2021/04/10 | DOE Joint Genome Institute |
| GCF_009729545.1 | 2019/12/04 | 2019/12/09 | North Carolina State University |
| GCF_000955745.1 | 2015/03/17 | 2015/03/19 | DOE Joint Genome Institute |
| GCF_000744885.1 | 2014/08/28 | 2014/12/25 | DOE Joint Genome Institute |
| GCF_003634755.1 | 2018/10/11 | 2018/10/18 | DOE Joint Genome Institute |
| GCF_001462395.1 | 2015/12/08 | 2015/12/10 | Oak Ridge National Laboratory |
| GCF_900095845.1 | 2016/09/08 | 2017/01/11 | URMITE |
| GCF_900114435.1 | 2016/11/02 | 2017/07/29 | DOE - JOINT GENOME INSTITUTE |
| GCF_000955735.1 | 2015/03/17 | 2015/03/19 | DOE Joint Genome Institute |
| GCF_900116205.1 | 2016/11/02 | 2017/07/29 | DOE - JOINT GENOME INSTITUTE |
| GCF_900104065.1 | 2016/10/21 | 2017/07/29 | DOE - JOINT GENOME INSTITUTE |
| GCF_007671655.1 | 2019/07/30 | 2019/08/06 | Third Institute of State Oceanic Administration |
| GCF_007671675.1 | 2019/07/30 | 2019/08/06 | Third Institute of State Oceanic Administration |
| GCF_900112175.1 | 2016/11/02 | 2017/07/29 | DOE - JOINT GENOME INSTITUTE |
| GCF_003226325.1 | 2018/06/13 | 2018/06/16 | Chinese Academy of Agricultural Sciences |
| GCF_900103715.1 | 2016/10/21 | 2017/07/29 | DOE - JOINT GENOME INSTITUTE |
| GCF_900108095.1 | 2016/10/27 | 2017/07/29 | DOE - JOINT GENOME INSTITUTE |
| GCF_900215575.1 | 2017/09/28 | 2017/10/12 | DOE - JOINT GENOME INSTITUTE |
| GCF_003688495.1 | 2018/10/25 | 2018/10/27 | DOE Joint Genome Institute |
| GCF_014647455.2 | 2020/09/11 | 2020/10/31 | WFCC-MIRCEN World Data Centre for Microorganisms (WDCM) |
| GCF_900110865.1 | 2016/10/31 | 2017/07/29 | DOE - JOINT GENOME INSTITUTE |
| GCF_017874375.1 | 2021/04/08 | 2021/04/10 | DOE Joint Genome Institute |
| GCF_007004735.1 | 2019/07/12 | 2019/07/18 | Jiangsu Universiry |
| GCF_001748385.1 | 2016/07/28 | 2016/09/29 | Korea Basic Science Institute |
| GCF_000455345.1 | 2013/08/20 | 2015/08/02 | ebi |
| GCF_004102725.1 | 2019/01/22 | 2019/01/26 | National Chung Hsing University |
| GCF_900166965.1 | 2017/03/17 | 2017/03/24 | UVEG |

| | | | |
|---|---|---|---|
| GCF_001602375.1 | 2016/01/19 | 2016/08/18 | Bioproduction Research Institute, National Institute of Advanced Industrial Science and Technology (AIST) |
| GCF_001375655.1 | 2015/04/22 | 2017/07/29 | URMITE |
| GCF_002806945.1 | 2017/12/04 | 2017/12/06 | Shandong Uniwersity at Weihai |
| GCF_014647115.1 | 2020/09/11 | 2020/09/20 | WFCC-MIRCEN World Data Centre for Microorganisms (WDCM) |
| GCF_002153915.1 | 2017/05/22 | 2017/05/24 | National Center for Biotechnology Information |
| GCF_015223105.1 | 2020/11/03 | 2020/11/05 | Guangdong Academy of Sciences |
| GCF_002954225.1 | 2018/02/27 | 2018/03/04 | Institute of Microbiology, Chinese Academy of Sciences |
| GCF_003966225.1 | 2018/12/23 | 2018/12/28 | Third Institute of Oceanography, State of Oceanic Administration |
| GCF_014647155.1 | 2020/09/11 | 2020/09/20 | WFCC-MIRCEN World Data Centre for Microorganisms (WDCM) |
| GCF_900129575.1 | 2016/12/03 | 2016/12/18 | DOE - JOINT GENOME INSTITUTE |
| GCF_001758465.1 | 2016/10/12 | 2016/10/13 | Second Institute of Oceanography, State Oceanic Administration |
| GCF_017873335.1 | 2021/04/08 | 2021/04/10 | DOE Joint Genome Institute |
| GCF_900114835.1 | 2016/11/02 | 2017/07/29 | DOE - JOINT GENOME INSTITUTE |
| GCF_004765815.2 | 2020/01/27 | 2020/02/07 | Jiangsu University |
| GCF_003719725.1 | 2018/11/10 | 2018/11/14 | Northwest Institute of Eco-Environment and Resources,CAS |
| GCF_900106945.1 | 2016/10/22 | 2017/07/29 | DOE - JOINT GENOME INSTITUTE |
| GCF_000711215.1 | 2014/06/24 | 2015/02/16 | DOE Joint Genome Institute |
| GCF_003217155.1 | 2018/06/12 | 2018/06/16 | DOE Joint Genome Institute |
| GCF_003966265.1 | 2018/12/23 | 2018/12/28 | Third Institute of Oceanography, State of Oceanic Administration |
| GCF_900110375.1 | 2016/10/31 | 2017/07/29 | DOE - JOINT GENOME INSTITUTE |
| GCF_004137855.1 | 2019/02/05 | 2019/02/16 | UBO |
| GCF_000766865.1 | 2014/10/10 | 2014/12/24 | Cornell University |
| GCF_008122205.1 | 2019/08/28 | 2019/09/03 | NCMR- NCCS |
| GCF_013760845.1 | 2020/07/27 | 2020/08/03 | DOE Joint Genome Institute |
| GCF_000687875.1 | 2014/04/08 | 2014/07/11 | DOE Joint Genome Institute |
| GCF_001029285.1 | 2015/06/17 | 2015/06/18 | Goettingen Genomics Laboratory |
| GCF_900107635.1 | 2016/10/22 | 2017/07/29 | DOE - JOINT GENOME INSTITUTE |
| GCF_900113635.1 | 2016/11/02 | 2017/07/29 | DOE - JOINT GENOME INSTITUTE |
| GCF_900142415.1 | 2016/12/03 | 2016/12/18 | DOE - JOINT GENOME INSTITUTE |

| | | | |
|---|---|---|---|
| GCF_000934745.1 | 2015/02/26 | 2015/03/19 | DOE Joint Genome Institute |
| GCF_900182635.1 | 2019/07/11 | 2019/07/20 | DOE - JOINT GENOME INSTITUTE |
| GCF_000686525.1 | 2014/05/08 | 2014/07/11 | DOE Joint Genome Institute |
| GCF_002909375.1 | 2018/01/30 | 2018/02/01 | Institute of Microbiology, Chinese Academy of Sciences |
| GCF_900142885.1 | 2016/12/03 | 2016/12/18 | DOE - JOINT GENOME INSTITUTE |
| GCF_003026475.1 | 2018/03/31 | 2018/04/05 | FDA/CFSAN |
| GCF_000753795.1 | 2014/09/10 | 2014/12/24 | 85303 |
| GCF_002094855.1 | 2017/04/17 | 2017/04/21 | The Third Institute of State Oceanic Administration (SOA) |
| GCF_900108015.1 | 2016/10/24 | 2018/02/11 | DOE - JOINT GENOME INSTITUTE |
| GCF_001017125.1 | 2015/06/01 | 2015/06/26 | National Chung Hsing University |
| GCF_004745425.1 | 2019/04/08 | 2019/04/11 | National Chung Hsing University |
| GCF_017873625.1 | 2021/04/08 | 2021/04/10 | DOE Joint Genome Institute |
| GCF_001640115.1 | 2016/05/06 | 2016/08/19 | Korea University |
| GCF_900129585.1 | 2016/12/03 | 2016/12/18 | DOE - JOINT GENOME INSTITUTE |
| GCF_900106645.1 | 2016/10/22 | 2017/07/29 | DOE - JOINT GENOME INSTITUTE |
| GCF_002954245.1 | 2018/02/27 | 2018/03/04 | Institute of Microbiology, Chinese Academy of Sciences |
| GCF_900156755.1 | 2017/01/14 | 2017/01/21 | DOE - JOINT GENOME INSTITUTE |
| GCF_014201585.1 | 2020/08/14 | 2020/08/17 | DOE Joint Genome Institute |
| GCF_001594015.1 | 2016/03/18 | 2016/03/28 | Goettingen Genomics Laboratory |
| GCF_900129115.1 | 2016/12/03 | 2016/12/18 | DOE - JOINT GENOME INSTITUTE |
| GCF_003336745.1 | 2018/07/24 | 2018/07/26 | SUN YAT-SEN UNIVERSITY |
| GCF_900104215.1 | 2016/10/21 | 2017/07/29 | DOE - JOINT GENOME INSTITUTE |
| GCF_001029445.1 | 2015/06/18 | 2015/06/19 | Technical University of Denmark |
| GCF_014201825.1 | 2020/08/14 | 2020/08/17 | DOE Joint Genome Institute |
| GCF_001939735.1 | 2017/01/09 | 2017/01/12 | Yantai Institute of Coastal Zone Research,Chinese Academy of Sciences |
| GCF_003841505.1 | 2018/11/26 | 2018/11/30 | Winogradsky Institute of Microbiology |
| GCF_900099915.1 | 2016/10/21 | 2017/07/29 | DOE - JOINT GENOME INSTITUTE |
| GCF_900112975.1 | 2016/11/02 | 2017/07/29 | DOE - JOINT GENOME INSTITUTE |
| GCF_004366795.1 | 2019/03/22 | 2019/03/27 | DOE Joint Genome Institute |
| GCF_900111555.1 | 2016/10/31 | 2017/07/29 | DOE - JOINT GENOME INSTITUTE |

| | | | |
|---|---|---|---|
| GCF_900188065.1 | 2017/07/16 | 2017/07/21 | DOE - JOINT GENOME INSTITUTE |
| GCF_900110465.1 | 2016/11/17 | 2017/07/29 | DOE - JOINT GENOME INSTITUTE |
| GCF_001719065.1 | 2016/09/02 | 2016/09/07 | IRNAS-CSIC |
| GCF_003219815.1 | 2018/06/12 | 2018/06/16 | Institute of Microbiology, Chinese Academy of Sciences |
| GCF_900112455.1 | 2016/11/02 | 2017/07/29 | DOE - JOINT GENOME INSTITUTE |
| GCF_900128885.1 | 2016/12/03 | 2016/12/18 | DOE - JOINT GENOME INSTITUTE |
| GCF_000614975.1 | 2014/03/31 | 2014/07/11 | The University of Tokyo |
| GCF_003317055.1 | 2018/07/12 | 2018/07/18 | DOE Joint Genome Institute |
| GCF_007197645.1 | 2019/07/20 | 2019/08/01 | Third Institute of Oceanography, State of Oceanic Administration |
| GCF_003025615.1 | 2018/03/31 | 2018/04/05 | FDA/CFSAN |
| GCF_003862495.1 | 2018/12/04 | 2018/12/08 | institute of microbial technology |
| GCF_003173335.1 | 2018/05/28 | 2018/05/31 | Aalborg University |
| GCF_000525995.1 | 2014/01/31 | 2014/12/24 | Cornell University |
| GCF_001399675.1 | 2015/10/19 | 2015/10/21 | California Institute of Technology |
| GCF_900142255.1 | 2016/12/03 | 2016/12/18 | DOE - JOINT GENOME INSTITUTE |
| GCF_003731635.1 | 2018/11/15 | 2018/11/21 | Zhejiang Sci-Tech University |
| GCF_900156265.1 | 2017/01/14 | 2021/06/08 | DOE - JOINT GENOME INSTITUTE |
| GCF_000620065.1 | 2014/04/08 | 2014/07/11 | DOE Joint Genome Institute |
| GCF_000711975.1 | 2014/06/25 | 2014/08/05 | DOE Joint Genome Institute |
| GCF_002995805.1 | 2018/03/14 | 2018/03/16 | Goettingen Genomics Laboratory |
| GCF_016107535.1 | 2020/12/17 | 2020/12/19 | INRAE |
| GCF_002995755.1 | 2018/03/14 | 2018/03/16 | Goettingen Genomics Laboratory |
| GCF_004923255.1 | 2019/04/29 | 2019/05/03 | GEOMAR - Helmholtz Centre for Ocean Research Kiel |
| GCF_003368535.1 | 2018/08/09 | 2018/08/15 | Winogradsky Institute of Microbiology |
| GCF_000525875.1 | 2014/01/31 | 2014/12/24 | Cornell University |
| GCF_000615945.1 | 2014/03/31 | 2014/07/11 | The University of Tokyo |
| GCF_900142695.1 | 2016/12/03 | 2016/12/18 | DOE - JOINT GENOME INSTITUTE |
| GCF_002243045.1 | 2017/08/07 | 2017/08/09 | National Chung Hsing University |
| GCF_900111865.1 | 2016/11/02 | 2017/07/29 | DOE - JOINT GENOME INSTITUTE |
| GCF_000784355.1 | 2014/11/17 | 2015/06/26 | Leibniz Institute DSMZ Braunschweig |

| | | | |
|---|---|---|---|
| GCF_001484195.1 | 2016/01/04 | 2016/01/06 | Kyungpook National University, Shin's Lab. |
| GCF_900128965.1 | 2016/12/03 | 2016/12/18 | DOE - JOINT GENOME INSTITUTE |
| GCF_002964845.1 | 2018/02/28 | 2018/03/12 | SUN YAT-SEN UNIVERSITY |
| GCF_900111795.1 | 2016/11/02 | 2017/07/29 | DOE - JOINT GENOME INSTITUTE |
| GCF_900109085.1 | 2016/10/31 | 2017/07/29 | DOE - JOINT GENOME INSTITUTE |
| GCF_000949295.1 | 2015/03/06 | 2015/08/19 | Georg-August-University Goettingen, Genomic and Applied Microbiology, Goettingen Genomics Laboratory |
| GCF_003574095.1 | 2018/09/18 | 2018/09/20 | Biocant |
| GCF_900101165.1 | 2016/10/21 | 2017/07/29 | DOE - JOINT GENOME INSTITUTE |
| GCF_002287235.1 | 2017/09/06 | 2017/09/08 | University of California Santa Barbara |
| GCF_003574085.1 | 2018/09/18 | 2018/09/20 | Biocant |
| GCF_902143385.2 | 2019/09/17 | 2019/09/21 | RADBOUD UNIVERSITY NIJMEGEN |
| GCF_009617755.1 | 2019/10/25 | 2019/11/12 | National Institute of Technology and Evaluation (NITE) |
| GCF_001447355.1 | 2015/11/23 | 2015/11/25 | Yunnan normal university |
| GCF_000243315.1 | 2012/01/23 | 2016/11/30 | JGI |
| GCF_000745065.1 | 2014/08/28 | 2014/12/25 | DOE Joint Genome Institute |
| GCF_904846075.1 | 2020/11/21 | 2021/01/22 | MAX PLANCK INSTITUTE FOR DEVELOPMENTAL BIOLOGY |
| GCF_003350005.1 | 2018/07/31 | 2018/08/02 | University of Helsinki |
| GCF_000709345.1 | 2014/06/17 | 2014/07/11 | China Center for Type Culture Collection |
| GCF_000632495.1 | 2014/04/15 | 2015/06/26 | INDEAR |
| GCF_001950325.1 | 2016/12/09 | 2017/01/16 | Laboratory of Marine Microbiology, Graduate School of Agriculture, Kyoto University |
| GCF_001444505.1 | 2015/11/17 | 2015/11/19 | Third Institute of Oceanography, State Oceanic Administration |
| GCF_900162825.1 | 2017/02/10 | 2017/02/20 | INSTITUT PASTEUR |
| GCF_001027105.1 | 2015/06/03 | 2015/06/22 | Okinawa Institute of Advanced Sciences |
| GCF_009759685.1 | 2019/12/17 | 2019/12/19 | Monash University |
| GCF_000742895.1 | 2014/08/21 | 2015/01/21 | Los Alamos National Laboratory |
| GCF_006094395.1 | 2019/06/06 | 2019/06/08 | Zhejiang Tianke High Technology Development Co.Ltd. |
| GCF_000982825.1 | 2015/04/30 | 2015/05/06 | Food-borne Pathogen Omics Research Center |
| GCF_900636385.1 | 2018/12/20 | 2018/12/28 | SC |

| | | | |
|---|---|---|---|
| GCF_000021485.1 | 2008/12/19 | 2016/11/29 | TIGR |
| GCF_009183365.2 | 2021/03/24 | 2021/03/26 | Karolinska Institutet |
| GCF_000010125.1 | 2005/08/12 | 2016/11/29 | Kitasato Institute for Life Sciences, Japan |
| GCF_000026225.1 | 2008/12/18 | 2016/11/29 | Genoscope |
| GCF_000022485.1 | 2009/04/29 | 2016/11/29 | US DOE Joint Genome Institute (JGI-PGF) |
| GCF_900079115.1 | 2016/04/11 | 2016/04/29 | Institute of Biology |
| GCF_006757745.1 | 2019/06/14 | 2019/07/12 | National Institute of Technology and Evaluation |
| GCF_016406325.1 | 2020/12/27 | 2020/12/28 | Ocean University of China |
| GCF_000012285.1 | 2005/07/05 | 2016/11/29 | Danish Archaea Centre |
| GCF_001295365.1 | 2015/09/22 | 2016/01/27 | Institute of Microbiology, Chinese Academy of Sciences |
| GCF_900475595.1 | 2018/06/17 | 2018/06/25 | SC |
| GCF_000007305.1 | 2002/02/27 | 2016/11/29 | Utah Genome Center |
| GCF_000970205.1 | 2015/04/07 | 2015/06/25 | University of Illinois at Urbana-Champaign |
| GCF_900187025.1 | 2017/08/15 | 2017/08/28 | SC |
| GCF_000008545.1 | 2001/01/09 | 2020/03/06 | TIGR |
| GCF_000175575.2 | 2014/05/27 | 2022/02/12 | Center for Bioinformatics and Genome Biology |
| GCF_000145615.1 | 2010/08/13 | 2016/11/29 | US DOE Joint Genome Institute (JGI-PGF) |
| GCF_002900365.1 | 2018/01/26 | 2018/01/31 | Institute of Microbiology Chinese Academy of Sciences |
| GCF_001267405.1 | 2015/08/13 | 2015/08/22 | Goettingen Genomics Laboratory |
| GCF_000306765.2 | 2014/05/27 | 2015/03/09 | Institute of Microbiology, Chinese Academy of Sciences |
| GCF_001543205.1 | 2016/02/01 | 2016/02/02 | Slagelse Hospital |
| GCF_900636325.1 | 2018/12/20 | 2018/12/28 | SC |
| GCF_000816145.1 | 2015/01/07 | 2016/05/29 | University of Waterloo |
| GCF_000008665.1 | 1999/12/22 | 2016/11/29 | TIGR |
| GCF_000008645.1 | 1999/12/22 | 2016/11/29 | Genome Therapeutics Corporation |
| GCF_004799665.1 | 2019/04/18 | 2019/04/24 | Wuhan University |
| GCF_003967175.1 | 2018/08/01 | 2018/12/28 | Laboratory of Extremophiles, Environmental Engineering for Symbiosis, Graduate school Soka University |
| GCF_002214545.1 | 2017/07/05 | 2017/07/07 | Ecole Normale Superieure de Lyon |
| GCF_000819445.1 | 2015/01/16 | 2015/01/24 | CeBiTec, Bielefeld University |

| | | | |
|---|---|---|---|
| GCF_000020585.3 | 2013/09/13 | 2017/07/10 | The Gordon and Betty Moore Foundation Marine Microbiology Initiative |
| GCF_000013905.1 | 2006/04/18 | 2016/11/29 | DOE Joint Genome Institute |
| GCF_000060285.1 | 2006/10/27 | 2017/02/26 | Universitat Giessen |
| GCF_000446015.1 | 2013/08/13 | 2015/08/20 | Baltic Federal University |
| GCF_003019275.1 | 2018/03/26 | 2018/03/29 | Illinois Institute of Technology |
| GCF_002214605.1 | 2017/07/05 | 2017/07/07 | Ecole Normale Superieure de Lyon |
| GCF_013343115.1 | 2020/06/15 | 2020/06/15 | US Food and Drug Administration |
| GCF_016406305.1 | 2020/12/27 | 2020/12/28 | Ocean University of China |
| GCF_001445575.1 | 2015/11/18 | 2016/03/28 | CSIR-Institute of Himalayan Bioresource Technology |
| GCF_001294625.1 | 2015/09/18 | 2016/03/28 | University of Malaya |
| GCF_000219875.1 | 2011/07/05 | 2016/11/30 | Key Laboratory of Genome Science and Information, Beijing Institute of Genomics, Chinese Academy of Sciences |
| GCF_001644665.1 | 2016/05/16 | 2016/05/17 | University of Manitoba |
| GCF_000012745.1 | 2005/09/01 | 2016/11/29 | DOE Joint Genome Institute |
| GCF_001679725.1 | 2016/07/05 | 2016/08/18 | Korea Polar Research Institute |
| GCF_002355395.1 | 2016/01/15 | 2017/09/27 | Graduate School of Agriculture, Kindai University |
| GCF_001880325.1 | 2016/11/17 | 2016/11/18 | Hamburg University of Technology |
| GCF_000828575.1 | 2012/12/03 | 2015/06/26 | National Institute of Genetics |
| GCF_019263745.1 | 2021/07/16 | 2021/07/18 | University of Minnesota |
| GCF_000025665.1 | 2010/02/24 | 2016/11/29 | US DOE Joint Genome Institute |
| GCF_002966495.1 | 2018/03/01 | 2018/03/07 | Wuhan University |
| GCF_004634385.1 | 2019/04/04 | 2019/04/06 | Chungbuk national university |
| GCF_001719125.1 | 2016/09/06 | 2016/09/07 | Institute of Microbiology, Chinese Academy of Sciences |
| GCF_000195575.1 | 2011/04/08 | 2016/11/29 | Georg-August-University Goettingen |
| GCF_017310505.1 | 2021/03/09 | 2021/03/13 | Korea Institute of Geoscience and Mineral Resources |
| GCF_000151205.2 | 2011/09/08 | 2018/04/04 | Moore Foundation |
| GCF_000211475.1 | 2011/04/29 | 2016/11/29 | Korea Ocean Research and Development Institute |
| GCF_013282665.1 | 2020/06/06 | 2020/06/07 | PulseNet Next Generation Subtyping Methods Unit |
| GCF_002201895.1 | 2017/06/20 | 2019/09/01 | Zhejiang University of Technology |
| GCF_000148995.1 | 2010/11/05 | 2016/11/29 | Baylor College of Medicine |

| | | | |
|---|---|---|---|
| GCF_001544255.1 | 2016/01/28 | 2016/02/03 | National Institute of Technology and Evaluation |
| GCF_000507185.2 | 2015/07/20 | 2015/08/14 | KRIBB |
| GCF_001792775.2 | 2018/06/30 | 2018/07/13 | University of Tennessee |
| GCF_002901845.1 | 2018/01/27 | 2018/01/31 | Modernising Medical Microbiology |
| GCF_002901825.1 | 2018/01/27 | 2018/01/31 | Modernising Medical Microbiology |
| GCF_002732165.1 | 2017/10/26 | 2017/10/28 | Masaryk University |
| GCF_008124605.1 | 2019/08/29 | 2019/09/03 | DOE Joint Genome Institute |
| GCF_002902325.1 | 2018/01/27 | 2018/01/31 | Modernising Medical Microbiology |
| GCF_000759775.1 | 2014/09/25 | 2015/02/09 | National Institute of Technology and Evaluation (NITE) |
| GCF_002901705.1 | 2018/01/27 | 2018/01/31 | Modernising Medical Microbiology |
| GCF_002217405.1 | 2017/07/12 | 2017/07/14 | FDA |
| GCF_003026815.1 | 2018/03/31 | 2018/04/05 | FDA/CFSAN |
| GCF_003970495.1 | 2018/12/24 | 2021/07/17 | Modernising Medical Microbiology |
| GCF_900456975.1 | 2018/08/01 | 2018/08/07 | SC |
| GCF_000369065.1 | 2013/04/19 | 2013/05/09 | Broad Institute |
| GCF_900458565.1 | 2018/08/01 | 2018/08/04 | SC |
| GCF_014647135.1 | 2020/09/12 | 2020/09/20 | WFCC-MIRCEN World Data Centre for Microorganisms (WDCM) |
| GCF_004345675.1 | 2019/03/11 | 2019/03/16 | DOE Joint Genome Institute |
| GCF_000875895.1 | 2015/02/13 | 2015/02/17 | Zhejiang University |
| GCF_002901865.1 | 2018/01/27 | 2018/01/31 | Modernising Medical Microbiology |
| GCF_002902755.1 | 2018/01/27 | 2018/01/31 | Modernising Medical Microbiology |
| GCF_002902345.1 | 2018/01/27 | 2018/01/31 | Modernising Medical Microbiology |
| GCF_900105965.1 | 2016/10/22 | 2016/12/22 | DOE - JOINT GENOME INSTITUTE |
| GCF_002902285.1 | 2018/01/27 | 2018/01/31 | Modernising Medical Microbiology |
| GCF_002902235.1 | 2018/01/27 | 2018/01/31 | Modernising Medical Microbiology |
| GCF_002902725.1 | 2018/01/27 | 2018/01/31 | Modernising Medical Microbiology |
| GCF_002901945.1 | 2018/01/27 | 2018/01/31 | Modernising Medical Microbiology |
| GCF_001280255.1 | 2015/09/08 | 2015/09/11 | University of Gdansk |
| GCF_002072215.1 | 2017/03/24 | 2017/03/28 | Goettingen Genomics Laboratory |
| GCF_000381045.1 | 2013/04/24 | 2014/07/12 | DOE Joint Genome Institute |

| | | | |
|---|---|---|---|
| GCF_000368825.1 | 2013/04/19 | 2013/05/09 | Broad Institute |
| GCF_000444055.1 | 2013/08/05 | 2014/07/17 | Agricultural Research Organisation |
| GCF_014635045.1 | 2020/09/11 | 2020/09/19 | WFCC-MIRCEN World Data Centre for Microorganisms (WDCM) |
| GCF_001500315.1 | 2016/01/06 | 2016/01/08 | Harbin Medical University |
| GCF_000766145.1 | 2014/10/10 | 2014/12/24 | Cornell University |
| GCF_003970575.1 | 2018/12/24 | 2021/07/17 | Modernising Medical Microbiology |
| GCF_003797125.1 | 2018/11/19 | 2018/11/26 | Korea Polar Research Institute |
| GCF_002902685.1 | 2018/01/27 | 2018/01/31 | Modernising Medical Microbiology |
| GCF_000337775.1 | 2013/02/04 | 2013/02/07 | University of California, Davis |
| GCF_014646555.1 | 2020/09/11 | 2020/09/20 | WFCC-MIRCEN World Data Centre for Microorganisms (WDCM) |
| GCF_000632715.1 | 2014/04/11 | 2015/02/09 | National Institute of Technology and Evaluation |
| GCF_000188295.1 | 2011/02/10 | 2016/11/29 | Baylor College of Medicine |
| GCF_014200405.1 | 2020/08/14 | 2020/08/17 | DOE Joint Genome Institute |
| GCF_003025595.1 | 2018/03/31 | 2018/04/05 | FDA/CFSAN |
| GCF_004402405.1 | 2019/03/26 | 2019/03/29 | Institute of Microbiology, Chinese Academy of Sciences |
| GCF_900156405.1 | 2017/01/14 | 2017/01/21 | DOE - JOINT GENOME INSTITUTE |
| GCF_900109325.1 | 2016/10/31 | 2017/07/29 | DOE - JOINT GENOME INSTITUTE |
| GCF_002836975.1 | 2017/12/12 | 2017/12/14 | National Institute for Communicable Disease Control and Prevention, China CDC |
| GCF_000833605.1 | 2015/02/09 | 2015/02/10 | Universiti Teknologi Malaysia |
| GCF_900109825.1 | 2016/10/31 | 2017/07/29 | DOE - JOINT GENOME INSTITUTE |
| GCF_000820505.1 | 2015/01/16 | 2015/01/21 | University of Minnesota |
| GCF_001310975.1 | 2015/10/01 | 2015/10/15 | The University of Tokyo |
| GCF_001639275.1 | 2016/04/29 | 2016/05/13 | Goettingen Genomics Laboratory |
| GCF_003574035.1 | 2018/09/18 | 2018/09/20 | Biocant |
| GCF_002902265.1 | 2018/01/27 | 2018/01/31 | Modernising Medical Microbiology |
| GCF_000188055.2 | 2011/11/18 | 2016/11/30 | JCVI |
| GCF_000513855.1 | 2014/01/07 | 2015/02/14 | DOE Joint Genome Institute |
| GCF_001006765.1 | 2015/05/14 | 2015/05/15 | Jiamusi University |
| GCF_000425165.1 | 2013/07/11 | 2014/07/11 | DOE Joint Genome Institute |

| | | | |
|---|---|---|---|
| GCF_000373145.1 | 2013/04/23 | 2014/07/12 | DOE Joint Genome Institute |
| GCF_000421725.1 | 2013/07/09 | 2014/07/11 | DOE Joint Genome Institute |
| GCF_001570745.1 | 2016/02/10 | 2016/08/18 | National Institute of Technology and Evaluation |
| GCF_000525975.1 | 2014/01/31 | 2014/12/24 | Cornell University |
| GCF_000344175.1 | 2013/03/04 | 2013/03/29 | Cornell University |
| GCF_900101115.1 | 2016/10/21 | 2017/07/29 | DOE - JOINT GENOME INSTITUTE |
| GCF_000934465.1 | 2015/02/25 | 2015/03/04 | Zhejiang University |
| GCF_000518705.1 | 2014/01/13 | 2014/07/11 | DOE Joint Genome Institute |
| GCF_000382145.1 | 2013/04/24 | 2014/07/11 | DOE Joint Genome Institute |
| GCF_002902385.1 | 2018/01/27 | 2018/01/31 | Modernising Medical Microbiology |
| GCF_003026395.1 | 2018/03/31 | 2018/04/05 | FDA/CFSAN |
| GCF_002777975.1 | 2017/11/16 | 2017/11/22 | National Institute for Communicable Disease Control and Prevention, China CDC |
| GCF_900103805.1 | 2016/10/21 | 2017/07/29 | DOE - JOINT GENOME INSTITUTE |
| GCF_002909445.1 | 2018/01/30 | 2018/02/01 | Institute of Microbiology, Chinese Academy of Sciences |
| GCF_003605125.1 | 2018/10/01 | 2018/10/04 | National Institute for Communicable Disease Control and Prevention, China CDC |
| GCF_003026355.1 | 2018/03/31 | 2018/04/05 | FDA/CFSAN |
| GCF_014651955.1 | 2020/09/12 | 2020/09/21 | WFCC-MIRCEN World Data Centre for Microorganisms (WDCM) |
| GCF_001077885.1 | 2015/07/14 | 2015/07/16 | CIAD, A.C. |
| GCF_900110125.1 | 2016/10/31 | 2017/07/29 | DOE - JOINT GENOME INSTITUTE |
| GCF_000525795.1 | 2014/01/31 | 2014/12/24 | Cornell University |
| GCF_002954455.1 | 2018/02/27 | 2018/03/04 | Atmosphere and Ocean Research Institute, The University of Tokyo |
| GCF_002849605.1 | 2018/01/02 | 2018/01/11 | Technische Universitaet Muenchen |
| GCF_900166975.1 | 2017/03/17 | 2017/03/24 | UVEG |
| GCF_002902305.1 | 2018/01/27 | 2018/01/31 | Modernising Medical Microbiology |
| GCF_002902405.1 | 2018/01/27 | 2018/01/31 | Modernising Medical Microbiology |
| GCF_002902625.1 | 2018/01/27 | 2018/01/31 | Modernising Medical Microbiology |
| GCF_000338275.1 | 2013/02/07 | 2014/07/12 | Zhejiang University |
| GCF_018854495.1 | 2021/06/11 | 2021/06/13 | Fundacion Ciencia y Vida |

| | | | |
|---|---|---|---|
| GCF_015276835.1 | 2020/11/06 | 2020/11/10 | Cornell University |
| GCF_002994445.1 | 2018/03/12 | 2018/03/16 | Helsinki University |
| GCF_003043455.1 | 2018/04/06 | 2018/04/18 | University of Calgary |
| GCF_000513315.1 | 2013/12/19 | 2015/02/16 | URMITE |
| GCF_007991715.1 | 2019/07/31 | 2019/08/16 | National Institute of Technology and Evaluation Biological Resource Center |
| GCF_001280565.1 | 2015/09/09 | 2015/09/21 | Fundacion Ciencia & Vida |
| GCF_003026285.1 | 2018/03/31 | 2018/04/05 | FDA/CFSAN |
| GCF_003399645.1 | 2018/08/16 | 2018/08/21 | Korea Polar Research Institute |
| GCF_003399625.1 | 2018/08/16 | 2018/08/21 | Korea Polar Research Institute |
| GCF_003264935.1 | 2018/06/25 | 2018/07/12 | Agharkar Research Institute |
| GCF_003399635.1 | 2018/08/16 | 2018/08/21 | Korea Polar Research Institute |
| GCF_900095385.1 | 2016/12/16 | 2016/12/22 | CEBITEC |
| GCF_000744175.1 | 2014/08/28 | 2014/12/25 | DOE Joint Genome Institute |
| GCF_000252125.1 | 2012/03/08 | 2016/11/30 | University of Nevada Las Vegas |
| GCF_904846175.1 | 2020/11/21 | 2021/01/22 | MAX PLANCK INSTITUTE FOR DEVELOPMENTAL BIOLOGY |
| GCF_904846235.1 | 2020/11/21 | 2021/01/22 | MAX PLANCK INSTITUTE FOR DEVELOPMENTAL BIOLOGY |
| GCF_900215215.1 | 2017/09/21 | 2017/09/29 | DOE - JOINT GENOME INSTITUTE |
| GCF_904846285.1 | 2020/11/21 | 2021/01/22 | MAX PLANCK INSTITUTE FOR DEVELOPMENTAL BIOLOGY |
| GCF_900111485.1 | 2016/11/17 | 2017/07/29 | DOE - JOINT GENOME INSTITUTE |
| GCF_904846225.1 | 2020/11/21 | 2021/01/22 | MAX PLANCK INSTITUTE FOR DEVELOPMENTAL BIOLOGY |
| GCF_904846185.1 | 2020/11/21 | 2021/01/22 | MAX PLANCK INSTITUTE FOR DEVELOPMENTAL BIOLOGY |
| GCF_000702485.1 | 2014/06/11 | 2014/07/11 | DOE Joint Genome Institute |
| GCF_002572525.1 | 2017/10/17 | 2017/10/20 | World Institute of Kimchi |
| GCF_902712915.1 | 2020/03/05 | 2020/06/14 | Friedrich Schiller University Jena |
| GCF_018860765.1 | 2021/06/12 | 2021/06/14 | Massachusetts Institute of Technology |
| GCF_003841465.1 | 2018/11/26 | 2018/11/30 | Winogradsky Institute of Microbiology |
| GCF_018861005.1 | 2021/06/12 | 2021/06/14 | Massachusetts Institute of Technology |
| GCF_003722075.1 | 2018/11/12 | 2018/11/14 | King Abdullah University of Science and Technology (KAUST) |
| GCF_003097355.1 | 2018/05/07 | 2018/08/29 | Florida A&M University |
| GCF_904846695.1 | 2020/11/21 | 2021/01/22 | MAX PLANCK INSTITUTE FOR DEVELOPMENTAL BIOLOGY |

| | | | |
|---|---|---|---|
| GCF_904846215.1 | 2020/11/21 | 2021/01/22 | MAX PLANCK INSTITUTE FOR DEVELOPMENTAL BIOLOGY |
| GCF_904846675.1 | 2020/11/21 | 2021/01/22 | MAX PLANCK INSTITUTE FOR DEVELOPMENTAL BIOLOGY |
| GCF_904846415.1 | 2020/11/21 | 2021/01/22 | MAX PLANCK INSTITUTE FOR DEVELOPMENTAL BIOLOGY |
| GCF_003035445.1 | 2018/04/05 | 2018/04/12 | University of Calgary |
| GCF_001707885.1 | 2016/08/18 | 2016/08/21 | Institute of Bioengineering, Research Center of Biotechnology of RAS |
| GCF_009765975.1 | 2019/12/18 | 2020/01/12 | Institute of Microbiology of the AS CR |
| GCF_004421185.1 | 2019/03/27 | 2019/03/29 | University of Bergen |
| GCF_009765685.1 | 2019/12/18 | 2020/01/12 | Institute of Microbiology of the AS CR |
| GCF_009914725.1 | 2020/01/22 | 2020/03/08 | National Institute of Advanced Industrial Science and Technology |
| GCF_002355995.1 | 2016/11/26 | 2020/12/30 | Kyushu University |
| GCF_001708385.1 | 2016/08/23 | 2016/12/20 | BioInformatics Center |
| GCF_000508305.1 | 2013/12/16 | 2016/12/30 | Center for Genomic Sciences |
| GCF_000016605.1 | 2007/04/25 | 2021/04/14 | US DOE Joint Genome Institute |
| GCF_000953475.1 | 2014/10/06 | 2020/12/16 | LEIDEN UNIVERSITY MEDICAL CENTER |
| GCF_003722315.1 | 2018/11/13 | 2021/04/14 | University of Exeter |
| GCF_001562115.1 | 2016/02/17 | 2021/04/14 | University Miguel Hernandez |
| GCF_000027065.2 | 2011/02/17 | 2020/12/16 | Technische Universitaet Muenchen |
| GCF_000831005.1 | 2014/02/05 | 2020/12/30 | Swinburne University of Technology |
| GCF_000789255.1 | 2014/12/04 | 2021/04/05 | Centre Bioengineering RAS |
| GCF_000967895.1 | 2013/05/31 | 2020/12/16 | Max Planck Institute for Molecular Genetics |
| GCF_000237975.1 | 2011/12/14 | 2015/08/25 | DOE Joint Genome Institute |
| GCF_000306725.1 | 2012/10/18 | 2019/10/03 | Beijing Institute of Genomics |
| GCF_000247545.1 | 2012/02/10 | 2019/01/25 | UCSC-Lowelab |
| GCF_000145985.1 | 2010/08/24 | 2015/08/25 | US DOE Joint Genome Institute (JGI-PGF) |
| GCF_017298655.1 | 2021/03/08 | 2022/01/27 | Institute of Microbiology |
| GCF_000953735.1 | 2014/09/30 | 2020/12/16 | University of Tromso |
| GCF_005954745.1 | 2019/06/04 | 2021/04/14 | Jiangsu University |
| GCA_001189275.1 | 2015/07/30 | 2015/07/30 | Montana State University |
| GCF_900095815.1 | 2016/12/16 | 2020/09/30 | CEBITEC |
| GCF_000200715.1 | 2006/11/20 | 2018/07/13 | DOE Joint Genome Institute |

| | | | |
|---|---|---|---|
| GCF_000785785.1 | 2014/11/21 | 2022/12/22 | CIAD, A.C. |
| GCF_018853935.1 | 2021/06/11 | 2021/09/02 | Fundacion Ciencia y Vida |
| GCF_000482765.1 | 2013/10/25 | 2021/04/14 | DOE Joint Genome Institute |
| GCF_010604095.1 | 2020/02/14 | 2022/05/10 | Ares Genetics |
| GCF_015163485.1 | 2020/11/01 | 2021/04/05 | The third Institute of Oceanography, Ministry of natural resources |
| GCF_000745905.1 | 2014/08/28 | 2022/05/10 | DOE Joint Genome Institute |
| GCF_000336895.1 | 2013/02/04 | 2021/04/14 | University of California, Davis |
| GCF_904846145.1 | 2020/11/21 | 2022/02/16 | MAX PLANCK INSTITUTE FOR DEVELOPMENTAL BIOLOGY |
| GCA_904846265.1 | 2020/11/21 | 2020/11/23 | MAX PLANCK INSTITUTE FOR DEVELOPMENTAL BIOLOGY |
| GCA_002506535.1 | 2017/10/10 | 2017/10/10 | University of Queensland |
| GCA_001509375.1 | 2016/01/08 | 2016/01/08 | Lawrence Berkeley National Lab |
| GCA_002502785.1 | 2017/10/10 | 2017/10/10 | University of Queensland |
| GCA_002503885.1 | 2017/10/10 | 2017/10/10 | University of Queensland |
| GCF_001411745.2 | 2015/10/23 | 2021/04/14 | Universidad de Santiago de Chile |
| GCF_003353085.1 | 2018/08/01 | 2022/04/25 | UFRJ |
| GCF_000690595.1 | 2016/05/26 | 2019/11/20 | National Chung Hsing University, Taichung City, Taiwan R.O.C. |
| GCA_002279355.1 | 2017/08/16 | 2017/08/30 | None |
| GCA_003023725.1 | 2018/03/27 | 2018/03/27 | UC Berkeley |
| GCA_003023695.1 | 2018/03/27 | 2018/03/27 | UC Berkeley |
| GCA_011049865.1 | 2020/03/02 | 2022/06/27 | The University of Hong Kong |
| GCA_001516585.1 | 2016/01/15 | 2016/01/15 | Montana State University |
