## Supplementary Table S3 for "Environment and taxonomy shape the genomic signature of prokaryotic extremophiles"

**Supplementary Table S3A. Accuracy summary for an alternative random sampling with pseudo-labels.** Additional computational experiments for supervised training with random labeling in the restriction-free scenario (*overlapping genera*), where the artificial pseudo-labels are sampled according to the distribution of the true environmental labels.

| Dataset | k -value | Classification Model Accuracy (%) | | | | | |
|---|---|---|---|---|---|---|---|
| | | RBF SVM | Random Forrest | ANN | MLDSP 1 | MLDSP 2 | MLDSP 3 |
| Temperature | k = 1 | 22.26 | 29.42 | 31.77 | 22.58 | 32.27 | 29.60 |
| | k = 2 | 23.25 | 28.10 | 27.09 | 33.61 | 31.61 | 33.61 |
| | k = 3 | 21.57 | 30.93 | 32.29 | 33.44 | 28.93 | 33.78 |
| | k = 4 | 23.08 | 24.73 | 27.60 | 28.43 | 28.93 | 26.09 |
| | k = 5 | 23.42 | 28.59 | 28.93 | 24.08 | 28.60 | 26.59 |
| | k = 6 | 21.23 | 30.10 | 31.77 | 26.09 | 27.59 | 29.10 |
| pH | k = 1 | 51.20 | 46.61 | 50.53 | 47.85 | 44.09 | 46.77 |
| | k = 2 | 51.14 | 52.72 | 51.67 | 50.00 | 41.94 | 50.54 |
| | k = 3 | 44.71 | 50.09 | 52.25 | 56.99 | 52.15 | 56.99 |
| | k = 4 | 44.68 | 51.99 | 50.61 | 51.61 | 51.61 | 47.85 |
| | k = 5 | 42.95 | 51.02 | 53.74 | 48.39 | 51.61 | 56.45 |
| | k = 6 | 50.14 | 54.72 | 51.63 | 47.85 | 53.23 | 55.91 |

**Supplementary Table S3B**. Additional computational experiments for supervised training with random labeling in the restricted scenario (non-*overlapping genera*), where the artificial pseudo-labels are sampled according to the distribution of the true environmental labels.

| Dataset | k -value | Classification Model Accuracy (%) | | | | | |
|---|---|---|---|---|---|---|---|
| | | RBF SVM | Random Forrest | ANN | MLDSP 1 | MLDSP 2 | MLDSP 3 |
| Temperature | k = 1 | 23.91 | 32.93 | 30.15 | 26.30 | 32.40 | 28.10 |
| | k = 2 | 22.59 | 27.56 | 32.80 | 34.60 | 33.30 | 34.40 |
| | k = 3 | 26.90 | 29.09 | 32.60 | 30.80 | 30.60 | 30.60 |
| | k = 4 | 26.24 | 28.23 | 29.94 | 28.40 | 29.40 | 25.40 |
| | k = 5 | 21.42 | 28.30 | 28.31 | 27.30 | 29.40 | 24.90 |
| | k = 6 | 25.07 | 29.23 | 27.58 | 26.60 | 30.60 | 27.90 |
| pH | k = 1 | 41.78 | 55.89 | 48.60 | 47.80 | 46.20 | 46.80 |
| | k = 2 | 51.52 | 54.14 | 45.79 | 49.50 | 44.60 | 48.90 |
| | k = 3 | 45.33 | 50.11 | 47.06 | 57.00 | 54.80 | 55.90 |
| | k = 4 | 52.29 | 50.66 | 50.25 | 53.80 | 54.80 | 48.40 |
| | k = 5 | 47.35 | 52.33 | 54.80 | 44.60 | 54.80 | 49.50 |
| | k = 6 | 54.08 | 47.78 | 46.11 | 44.10 | 56.50 | 51.10 |
