## Supplementary Table S4 for "Environment and taxonomy shape the genomic signature of prokaryotic extremophiles"

**Supplementary Table S4. Referenced articles supporting consistencies in k-mer associations for the temperature dataset**. The following table describes the articles referenced in Table 5, which correlate to literature supporting the observed associated k-mers for each of the temperature-adapted extremophilic classes in the context of codon bias or amino acid compositional patterns.

| Reference Link | Observation |
|---|---|
| https://doi.org/10.1186%2F1471-2164-15-1120 | Increased valine, alanine, glycine, proline abundance in psychrophilic proteins |
| https://doi.org/10.1186/1471-2164-9-210 | Decreased methionine, arginine, cysteine, histidine, increased asparagine in psychrophilic proteins |
| https://doi.org/10.1110%2Fps.072947007 | Increase in alanine, cysteine, valine, glycine, and methionine in psychrophilic proteins, and a decrease in Asn, Lys, Tyr, Phe, and Glu |
| https://doi.org/10.1101/gr.1180903 | Increase in glutamine and threonine, and decrease in leucine in psychrophilic proteins. Also, decreased Gln/Thr/His/Ser at higher OGT (thermophilic range) and increased Glu (glutamic acid) |
| https://doi.org/10.1074/jbc.m802158200 | Increase glutamate for cold stability in psychrophilic proteins |
| https://doi.org/10.1093/femsec/fiy023 | Increase in glycine/serine, decrease in aspartic/glutamic acids |
| https://doi.org/10.1016/S0969-2126(00)00133-7 | Low serine, lower isoleucine abundance in mesophiles |
| https://doi.org/10.1038/s41598-020-58825-7 | Gln and Met have increased abundance in mesophiles |
| https://doi.org/10.1093/protein/13.3.179 | Increased proline abundance in mesophiles relative to thermophiles |
| https://doi.org/10.1371/journal.pcbi.0030005 | Support for the IVYWREL adaptation set for increase OGT |
| https://doi.org/10.1002/prot.25866 | Increased proline (codon=CCN) due to increased GC% in thermophiles |
| https://doi.org/10.1186/gb-2002-3-8-preprint0006 | Gly, Lys, Tyr, and Ile are preferred in thermophilic organisms |
| https://doi.org/10.1002/prot.25866 | Threonine and serine are rare in thermophiles and frequent in hyperthermophiles, while prolines are rare in hyperthermophiles |
| https://doi.org/10.1186/gb-2002-3-8-preprint0006 | More charged (Glu, Arg, Lys) and fewer uncharged polar residues (Ser, Thr, Asn, Gln, His, Cys), increased residue hydrophobicity (Ile, Val), and increased residue volume (Tyr) |
