## Supplementary Table S5 for "Environment and taxonomy shape the genomic signature of prokaryotic extremophiles"

**Supplementary Table S5. Referenced articles supporting consistencies in k-mer associations for the pH dataset.** The following table describes the articles referenced in Table 6, which correlate to literature supporting the observed associated k-mers for each of the pH-adapted extremophilic classes in the context of codon bias or amino acid compositional patterns.

| Reference | Reference Link | Observation |
|---|---|---|
| Khan, M. F. & Patra, S. (2018) | DOI: 10.1038/s41598-018-33476-x | "TGG in acidophiles, GAG in alkaliphiles and GAC in halophiles had highest preference. In (alkaliphile-acidophile) dataset, the codons like AAC (Asn), TGG (Trp), TAC (Tyr), ACT (Thr), TTC (Phe), ACC (Thr), AAT (Asn), TCC (Ser), TAT (Tyr), CAA (Gln), etc. had higher abundance and GAG (Glu), GAA (Glu), CTG (Leu), AAA (Lys), GTG (Val), GCG (Ala), AAG (Lys), CGC (Arg), etc. had lowest abundance in acidophiles and vice versa for alkaliphiles. Codons for small, polar and aromatic amino acid had higher frequency in acidophilic proteins whereas, charged and aliphatic amino acid codons had higher frequencies in alkaliphilic proteins. New finding was obtained in the A-B dataset, as TGG (Trp) and TAC (Tyr) codon ranked highest in acidophiles whereas, GAG (Glu) codon got highest rank in alkaliphiles." |
| Horikoshi, K. (1999) | DOI: 10.1128/mmbr.63.4.735-750.1999 | "This analysis revealed a decrease in the number of negatively charged amino acids (aspartic acid and glutamic acid) and lysine residues, and an increase in arginine and neutral hydrophilic amino acids (histidine, asparagine, and glutamine) residues during the course of adaptation (in alkaliphiles)." |
| Mukhtar, S., *et al.* (2022) | https://doi.org/10.1016/B978-0-12-822945-3.00010-5 | "Use of increased prolines for maintenance of low pH in acidophiles" |
