## Supplementary Table S6 for "Environment and taxonomy shape the genomic signature of prokaryotic extremophiles"

**Supplementary Table S6. Characterizing phenotypic traits of candidate species and their isolating environment.** The following table describes various phenotypic traits attributed to each of the candidate species, and the geochemical information associated with the environments the microbes were initially isolated from. Coordinates are described in the Longitude/Latitude format.

| **Phenotypic Trait** | ***Pyrococcus furiosus*** | ***Thermococcus litoralis*** | ***Pyrococcus chitonophagus*** | ***Thermocrinis ruber*** |
|---|---|---|---|---|
| References | •Fiala, G. & Stetter, K. O. (1986) DOI: 10.1007/BF00413027<br>•Capaccioni, B., Tassi, F. & Vaselli, O. (2001) DOI: 10.1016/S0377-0273(00)00284-5 | •Neuner, A., Jannasch, H. W., Belkin, S. & Stetter, K. O. (1990) DOI: 10.1007/BF00247822<br>•Orlando, V., Franco, T., Dario, T., Robert, P. J. & Antonio, C. (2011) DOI: 10.1016/j.proeps.2011.11.007 | •Huber, R. et al. (1995) DOI: 10.1007/BF02529959<br>•Bazylinski, D. A., Farrington, J. W. & Jannasch, H. W. (1988) DOI: 10.1016/0146-6380(88)90146-5 | •Clifton, C., Walters, C. & Simoneit, B. (1990) DOI: 10.1016/0883-2927(90)90047-9<br>•Huber, R. et al. (1998) DOI: 10.1128/AEM.64.10.3576-3583.1998 |
| Domain | Archaea | Archaea | Archaea | Bacteria |
| DSM Strain ID | 3638 | 5473 | 10152 | 12173 |
| pH Range | pH 5.0 to 9.0 (OGpH=7.0) | pH 4.0 to 8.0 (OGpH=6) | pH 3.5 to 9 (OGpH=6.7) | pH 7 to 8.5 |
| Temperature Range | 70 to 103ºC (OGT=100ºC) | 55 to 98ºC (OGT=85ºC) | 60 to 93ªC (OGT=75ºC) | 44 to 89ºC (OGT=80ºC) |
| Salinity | 0.5 to 5% NaCl | 1.8 to 6.5% NaCl (optimum 2.5%) | 1.8 to 6.5% NaCl (optimum 2.5%) | 0 to 0.4% NaCl |
| Cell Shape | Slightly irregular cocci | Round to irregular cocci | Round to irregular cocci | Rod-shaped cells |
| Gram Stain | Negative | Negative | Negative | Negative |
| Motility | Monopolar polytrichous flagella | Non-flagellated, non-motile | Monopolar polytrichous flagella | Monopolar polytrichous flagella |
| S-Layer Protein Composition | Hexagonal lattice | Hexagonal lattice | Hexagonal lattice | No evidence of a regularly arrayed SLP |
| Oxygen Tolerance | Anaerobic | Anaerobic | Anaerobic | Aerobic |
| Genome Size | 1.89 Mb | 1.82 Mb | 1.95 Mb | 1.52 Mb |
| Intergenic Sequence Composition | Pseudogenes and IS elements present | Pseudogenes and IS elements present | Pseudogenes and IS elements present | Pseudogenes present |
| Geographic Source | Submarine solfataric field in the bay of Porto di Levante, Vulcano Island, Italy. | Shallow submarine solfataras near the beach of Lucrino, Bay of Naples, Italy | Smoker site, Guaymas Basin, Gulf of California, Mexico | Octopus Spring, Yellowstone National Park, Wyoming, USA |
| Coordinates of Isolating Environment | 38.41341, 14.9620271 | 40.822111, 14.090833 | 20.8333, -109.1 | 44.543652, -110.79792 |
| Hydrocarbon Emissions | Evidence for gas discharges containing light hydrocarbons | Evidence for gas discharges containing light hydrocarbons | Evidence for gas discharges containing petroleum-like substances | Evidence for hydrothermal petroleum-like substances in other hot springs atop the caldera |
