## Supplementary Table S7 for "Environment and taxonomy shape the genomic signature of prokaryotic extremophiles"

**Supplementary Table S7. Analyzing the co-occurrence of *Thermococcus litoralis* and *Pyrococcus chitonophagus* along the Guaymas Basin.** The following table describes the initial geographic location that *T. litoralis* and *P. chitonophagus* were isolated from. The hydrothermal landscape of Guaymas Basin, Mexico, and Lucrino Bay, Italy are also described. The locations they are isolated from along the Guaymas Basin, Gulf of California, Mexico, are also described.

| | ***Pyrococcus chitonophagus*** | ***Thermococcus litoralis*** |
|---|---|---|
| Initial Isolation Source | 20°50’00.0"N 109°06’00.0"W, Guaymas Basin, Gulf of California, Mexico (Huber, R. *et al.* 1995) | 40°49’19.6"N 14°05’27.0"E, Lucrino Bay, Pozzuoli, Italy (Neuner, A. *et al.* 1990) |
| JGI GOLD Ecosystem Classifier | 4027 (Mukherjee, S. *et al.* 2023) | 3991 (Mukherjee, S. *et al.* 2023) |
| Hydrothermal Landscape of Initial Isolation Source | Actively circulating hydrothermal system venting metal-enriched hydrothermal fluids to the ocean (Sherman, L. *et al.* 2009) | Deep hydrothermal system, including at least five submarine solfataric fields (Orlando, V. *et al.* 2011) |
| Coordinates of Co-Isolation in Guaymas Basin | 20°50’00.0"N 109°06’00.0"W (Huber, R. *et al.* 1995) | 27°00’28.1"N 111°24’25.6"W (Zhou, Z. *et al.* 2020) |
